# Reconstructing the history of founder events using genome-wide patterns of allele sharing across individuals

**DOI:** 10.1101/2020.09.07.286450

**Authors:** Rémi Tournebize, Gillian Chu, Priya Moorjani

**Affiliations:** Department of Molecular and Cell Biology, University of California, Berkeley, CA; Department of Electrical Engineering and Computer Science, University of California, Berkeley, CA; Center for Computational Biology, University of California, Berkeley, CA

## Abstract

Founder events play a critical role in shaping genetic diversity, impacting the fitness of a species and disease risk in humans. Yet our understanding of the prevalence and distribution of founder events in humans and other species remains incomplete, as most existing methods for characterizing founder events require large sample sizes or phased genomes. To learn about the frequency and evolutionary history of founder events, we introduce *ASCEND* (Allele Sharing Correlation for the Estimation of Non-equilibrium Demography), a flexible two-locus method to infer the age and strength of founder events. This method uses the correlation in allele sharing across the genome between pairs of individuals to recover signatures of past bottlenecks. By performing coalescent simulations, we show that *ASCEND* can reliably estimate the parameters of founder events under a range of demographic scenarios, with genotype or sequence data. We apply *ASCEND* to ~5,000 worldwide human samples (~3,500 present-day and ~1,500 ancient individuals), and ~1,000 domesticated dog samples. In both species, we find pervasive evidence of founder events in the recent past. In humans, over half of the populations surveyed in our study had evidence for a founder events in the past 10,000 years, associated with geographic isolation, modes of sustenance, and historical invasions and epidemics. We document that island populations have historically maintained lower population sizes than continental groups, ancient hunter-gatherers had stronger founder events than Neolithic Farmers or Steppe Pastoralists, and periods of epidemics such as smallpox were accompanied by major population crashes. Many present-day groups--including Central & South Americans, Oceanians and South Asians--have experienced founder events stronger than estimated in Ashkenazi Jews who have high rates of recessive diseases due to their history of founder events. In dogs, we uncovered extreme founder events in most groups, more than ten times stronger than the median strength of founder events in humans. These founder events occurred during the last 25 generations and are likely related to the establishment of dog breeds during Victorian times. Our results highlight a widespread history of founder events in humans and dogs, and provide insights about the demographic and cultural processes underlying these events.

## Introduction

A founder event occurs when a new population is formed by a subset of individuals from a larger one or when the original population goes through a reduction in size due to a bottleneck^1^. Founder events have played a critical role in shaping genetic diversity in many species, including humans. For instance, anatomically modern humans spread worldwide in the past ~50,000 years, following periods of successive bottlenecks and mixtures^2^. Many human populations have further undergone severe founder events in the recent past (last hundreds of generations) due to geographical isolation (e.g. Finns^3^ and Roma^4^) or historical migrations or cultural practices (e.g. Amish^5^ and Ashkenazi Jews^6^).

Founder events reduce genetic variation in a population, decrease the efficacy of selection to remove deleterious variants, and increase the risk of recessive diseases^1^. Understanding the history of founder events can thus be helpful for learning about the cultural and demographic events leading to population bottlenecks, and also for mapping functional and disease variants. Gene mapping efforts in founder populations—including Ashkenazi Jews, Finns, Amish, and French Canadians—have resulted in the discovery of numerous disease-causing mutations in each group^7^.

Despite the importance of founder events in evolutionary and disease studies, we still only have a limited comprehension of the number, tempo and properties (i.e. age and intensity) of founder events. Characterizing the timing and strength of founder events is the first step towards improving our understanding of the impact of founder events on neutral and deleterious genetic variation. In particular, the estimated timing of the founder event (referred to as *founder age*, henceforth) can inform us about the expected length of genomic segments that are inherited identical-by-descent (IBD) among individuals in a population and the cultural or environmental factors underlying the founder events. Founder intensity—a function of the effective population size and the duration of the bottleneck—measures the strength of genetic drift during the bottleneck and is informative about the probability of fixation of deleterious variants and ultimately, the risk of recessive diseases in a population^8^. Thus, together these parameters reveal the evolutionary history and impact of founder events in shaping genetic diversity.

There are two main classes of methods currently available for characterizing founder events: polymorphism-based and IBD-based approaches. Polymorphism-based approaches leverage the observed patterns of genetic variation–either by studying the density of heterozygous sites in a region (e.g., PSMC and MSMC^9,10^) or by analyzing the allele frequencies of markers in a population (such as dadi, PopSizeABC or fastsimcoal^11,12,13^)–to recover the time to the most recent common ancestor across the genome. These methods make inferences based on the mutation clock and thus have low resolution at recent timescales (in the past hundreds of generations)^9,10^. On the other hand, IBD-based methods use the distribution of IBD segments in a population to infer recent demographic history^14,15^. The length of IBD segments is inversely proportional to the age of the founder event—the older the event, the more recombination events have occurred and hence shorter IBD segments remain in the target population. Most IBD-based methods require large sample sizes and phased data for reliably detecting IBD segments. While accurate and precise for recent timescales, the inference can become unstable or noisy in sparse datasets or for older timescales (as most methods rely on segments >2 cM).

A third class of methods, recently introduced by Reich et al. (2009), characterizes the average allele sharing correlation across individuals in a population to infer the time of the founder event^16^. This approach uses the insight that a founder event introduces long-range linkage disequilibrium (LD) or allelic correlation in nearby loci co-inherited from a common ancestor by a pair of individuals in the population. As recombination occurs in each generation, it breaks down these associations. Thus, by measuring the decay of allelic association or LD across the genome at sites that are shared between pairs of individuals (i.e., identical by state (IBS)), we can infer how long ago the founder event occurred. A major advantage of this approach is that it does not require explicit identification of IBD segments and hence does not require phased data, making it applicable to sparse datasets that contain few individuals or low coverage samples (such as ancient genomes).

Here, we introduce *ASCEND* (Allele Sharing Correlation for the Estimation of Nonequilibrium Demography) that extends the idea introduced in Reich et al. (2009) and comprehensively develops the use of allele sharing correlation to not only estimate the age of the founder event, but also its intensity. We provide theoretical expectations for leveraging allele sharing correlation for estimating the parameters of the founder event. Finally, we implement the method using fast Fourier transform (FFT) to make the approach computationally tractable for large datasets. We report extensive simulations under a range of demographic events and apply the method to empirical datasets from two species–humans (using present-day and ancient samples) and modern dog breeds–to characterize the spatio-temporal patterns of founder events in both species.

## Results

### Overview of *ASCEND*

*ASCEND* measures the parameters (age, intensity) of a founder event by leveraging the distribution of LD across the genome at sites that are shared between pairs of individuals in a population. Briefly, assume we have two populations, one target population (population *A*) with a history of founder events (with varying age and intensity of bottlenecks) and an outgroup population (population *O*) for comparison (Figure 1). To estimate the age and intensity of the founder event, *ASCEND* computes the allele sharing correlation between pairs of individuals within the target population *A*, and then subtracts the cross-population correlation (with an outgroup, *O*) to remove the effects of ancestral allele sharing inherited from the common ancestor (see Methods). This statistic is expected to decay exponentially as a function of the genetic distance due to recombination events breaking down the correlation between pairs of sites over successive generations^16,17^ (see Methods, ***ASCEND*: Model and Theory**). Intuitively, if the founder event is recent, fewer recombination events have occurred and hence the rate of decay is slower. In addition, the stronger the founder event, the more correlated the shared alleles will be at shorter genetic distances and hence the amplitude of the exponential decay will be higher. By fitting an exponential distribution to the empirical decay of the allele sharing correlation, *ASCEND* simultaneously infers the age and the intensity of the founder event in population *A*.

**Figure 1.**
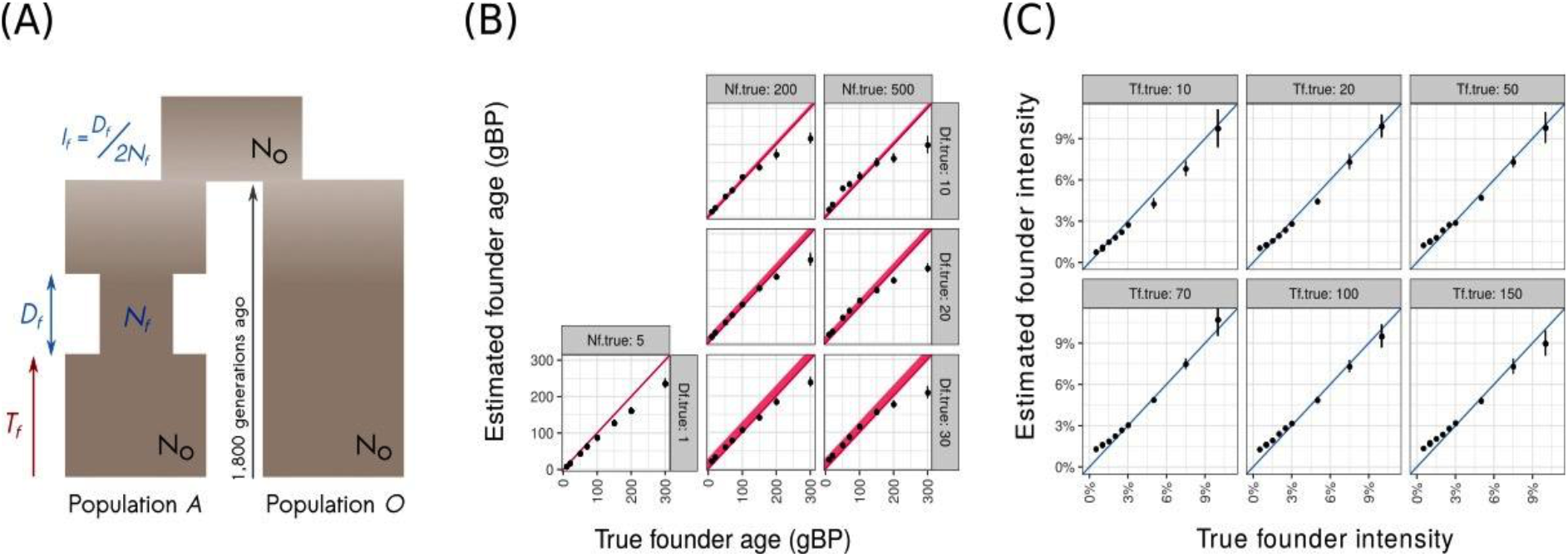
*ASCEND* simulation results. *(A) Model of founder event*. We simulated two populations, population *A* (target) and population *O* (outgroup) that diverged 1,800 generations ago. Population *A* had a founder event where the population size decreased to *N_f_* for a duration of *D_f_* generations that occurred *T_f_* generations before present. After the bottleneck, the population size recovered to *N_o_*. We ran *ASCEND* and compared the estimated parameters with the true parameters of the founder event in population *A*. *(B) Accuracy in estimating founder age*. The x-axis shows the true founder age that was simulated in generation before present (gBP) and y-axis shows the founder age estimated by *ASCEND*. The diagonal represents the expectation (i.e., the case where the estimated values are exactly the same as the true values). We note that for *D_f_* > 0 we show a thick band for the diagonal, proportional to duration of the founder event*. (C) Accuracy in estimating founder intensity*. We define the founder intensity as the ratio of the bottleneck duration over twice the effective population size during the bottleneck, i.e. *I_f_*=*D_f_*(2*N_f_*). The x-axis shows the true founder intensity and the y-axis shows the estimated founder intensity. The diagonal represents the expectation (i.e., the case where the estimated values are exactly the same as the true values).

To characterize the reliability of *ASCEND*, we simulated data under a range of demographic scenarios. First, we focused on the epoch model where the founder event in population *A* occurred *T_f_* generations ago (10 and 300 generations ago) such that the population size reduced to a few individuals (*N_f_* = 5 to 500) for a short duration (*D_f_* = 1 to 30 generations). After the founder event, the population recovered to its original size. Applying *ASCEND* to population *A* and accounting for the cross-population allele sharing with the outgroup *O*, we found that *ASCEND* reliably inferred the age and intensity of the bottleneck, when the founder event occurred within the past ~200 generations (Figure 1). Next, we simulated more complex models where the target *A* had a history of admixture (or gene flow) and founder event. Specifically, we simulated data from population *A* that derived 60% ancestry from population *A*’ and 40% ancestry from population *B*’ (Figure S2.4.1). Population *A*’ and *B*’ diverged 1,800 generations ago. The admixture event occurred 110 generations ago and was then followed by an extreme founder event (*N_f_* = 5) lasting for one generation that occurred between 10 to 100 generations ago (Figure S2.4.1, Note S2.4). Applying *ASCEND* to the target population and using one of the ancestral groups as the outgroup (to compute cross-population allele sharing), we obtained accurate estimates for the age and intensity of the founder event, and observed no noticeable impact of admixture on the parameters of the founder event (Figure S2.4.2).

Thus far, our simulations assumed a single epoch model of founder events. However, in reality the demographic history of a population can involve multiple epochs of founder events, a bottleneck with gradual (instead of instantaneous) exponential recovery over time or a bottleneck without recovery (i.e., maintenance of a small population size from *T_f_* to present). To test the impact of these alternate models, we simulated data under each of these three scenarios (Note S2). We found that in the case of two successive founder events, *ASCEND* recovered the parameters of the founder event with the strongest intensity (Figure S2.3.2). In the case of no recovery, *ASCEND* underestimated the founder age but reliably estimated the founder intensity. However, in the case of the gradual exponential recovery model, *ASCEND* tended to underestimate the founder age and overestimate the founder intensity. We note that this is unsurprising as this model has three key parameters: age, population size at the start of the bottleneck and rate of exponential increase in population size over time. In *ASCEND*, we fit a simple epoch model based on just two parameters, age and intensity. This means that summary statistics like the rate of exponential increase are not captured (but assumed to occur over a short duration). The estimated intensity in *ASCEND* represents a harmonic mean over the duration of the bottleneck and thus the results should be interpreted with caution, if there is evidence supporting the exponential recovery model (Note S2.5).

An important feature of *ASCEND* is that it does not require phased data, which makes it applicable to datasets with small sample sizes and low coverage such as ancient genomes. To test the reliability of *ASCEND* to make inferences with small datasets, we performed simulations to characterize the impact of the following parameters: (i) sample size; (ii) proportion of missing genotypes, and (iii) features of ancient DNA samples, including missing data, low sample sizes and use of pseudo-homozygous genotype calls–where the genotype is based on choosing a single random allele observed in the reads mapped at a particular site (Note S2.7.3). We observed that *ASCEND* estimated the parameters of the founder event accurately for low sample sizes, even with only 5 samples from the target population (Figure S2.7.1.2). When considering missing data, we found no bias even when the dataset had a large proportion of missing genotypes (e.g. 60% missing genotypes in a dataset with ~500,000 simulated SNPs) (Figure S2.7.2.2). Finally, simulations mimicking features of ancient DNA samples showed that *ASCEND* reliably recovered the parameters of the founder events (Figure S2.7.3.2). These results highlight a major strength of *ASCEND*, in that it works reliably even with limited data and hence is applicable to sparse datasets.

### Founder events in present-day human populations

We applied *ASCEND* to 5,637 present-day individuals from the Human Origins dataset to learn about the frequency and distribution of founder events in worldwide populations. For two groups that had been previously reported to have a history of founder events, South Asians and Ashkenazi Jews, we also analyzed two additional datasets with 1,662 South Asians (IndiaHO dataset)^18^ and 21 Ashkenazi Jews (Behar dataset)^19^. For all three datasets, we limited our analysis to all groups with a minimum of 5 samples. To ensure we are characterizing founder events, and not consanguinity that can also lead to long-range IBD sharing among individuals, we removed all individuals with evidence of recent relatedness (see Methods). After filtering, we retained 2,310 present-day individuals (184 groups) in the Human Origins Dataset, 1,253 individuals (116 groups) in the IndiaHO dataset, and 21 individuals in the Behar dataset (see Table S1 and Note S3).

We applied *ASCEND* to study the global patterns of founder events in recent human history. We found that 61% of the groups (113 out of 184) experienced significant founder events (see Methods) that occurred in the past 200 generations (choice of threshold is based on simulations (Figure 1B)). The ages of founder events ranged from ~10 generations (in Aleuts) to 195 generations (in Icelandic people), with a median of 27 generations (~750 years ago, assuming 28 years per generation^20^) (Figure 2B). The intensity of founder events ranged from 0.6% in Maasai from Kenya to 13.4% in Gimi islanders from Papua New Guinea, with a median intensity of 2.2% across populations in our dataset (Figure 2A). We found no correlation between founder age and founder intensity, suggesting that we can reliably disentangle the estimation of both parameters (Pearson’s r=0.05, *P*=0.60).

**Figure 2.**
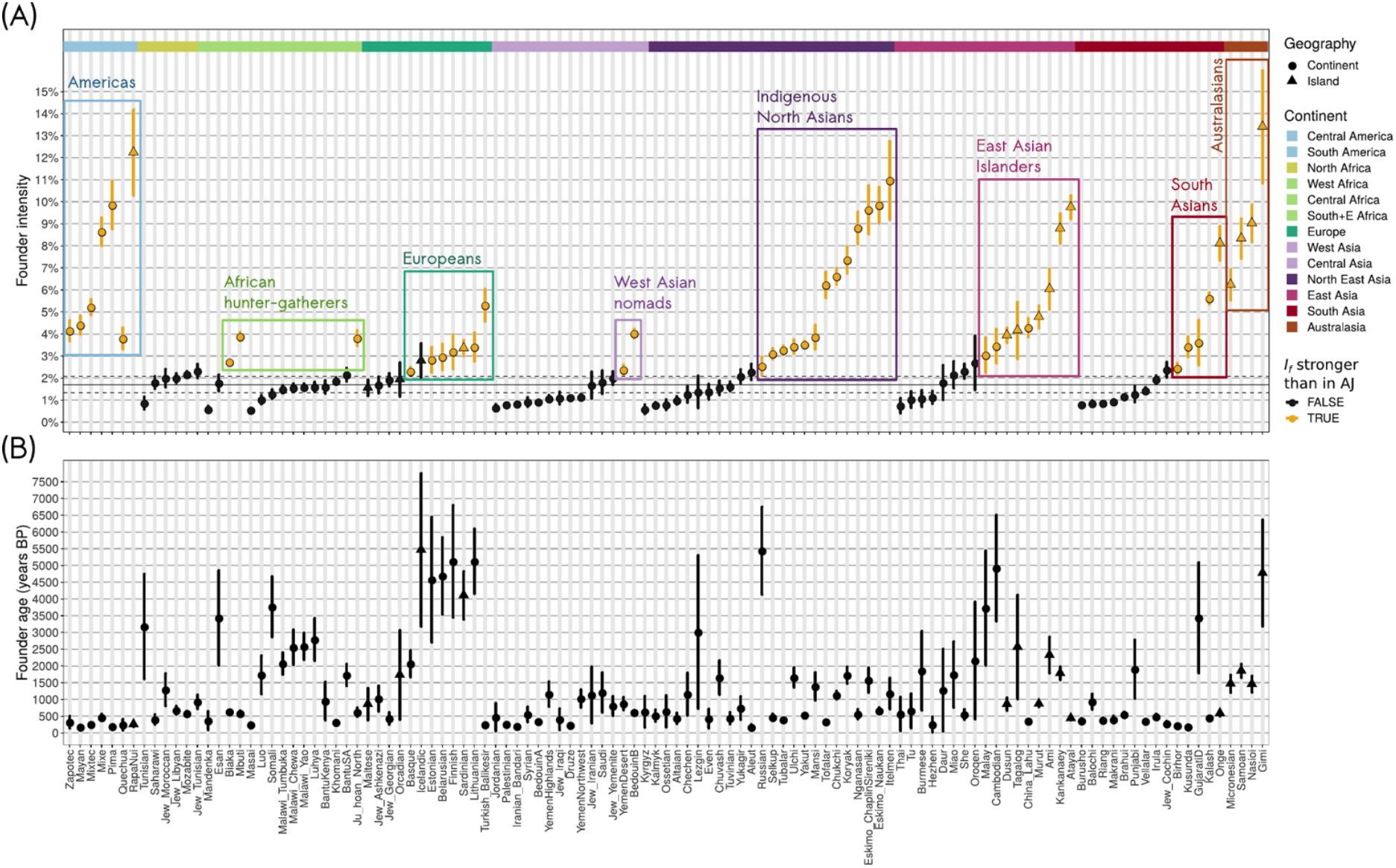
History of founder events in present-day human populations. Results of *ASCEND* for present-day populations in the Human Origins v37 dataset that passed filtering criteria and showed significant evidence of founder event (see Methods). Each point shown represents a population and the shape of the points indicates whether the group was sampled from an island (triangle) or a continental region (circle). The top coloured ribbon represents the various subcontinents where the populations are located. *(A) Distribution of the estimated founder intensities*. We show the founder intensity and the associated 95% confidence interval for all populations. The populations are ordered by their subcontinent and by increasing order of estimated founder intensity. The black horizontal line shows the inferred founder intensity in Ashkenazi Jews (AJ) (1.7% [1.3%– 2.1%]). Populations that have an estimated founder intensity that is significantly higher than AJ (i.e. where the intensity is greater than the upper-bound of the 95% confidence interval in AJs) are colored in gold, else they are shown in black. The colored boxes show population groups that share certain common features, based on geography or modes of sustenance and have an estimated founder intensity higher than AJ. *(B) Distribution of the estimated founder ages*. We show the estimated founder ages and the associated 95% confidence intervals. The estimated ages were converted from generations to years by assuming an average generation time of 28 years (Moorjani et al. 2016).

Next, we investigated the geographical patterns of founder events by studying the distribution of the age and intensity of bottlenecks across various continental groups. We found that founder intensities differed significantly across geographical continents (Kruskal-Wallis test, *P*=4×10^−5^). We estimated the highest founder intensities in Central America and Oceania, followed by groups in Asia, Europe and Africa (Figure 2A). Moreover, the founder events in Central America were also significantly more recent (occurring within the past 300 years) than in other geographical regions. We observed that island populations had more extreme founder events compared to continental groups. Using data from 16 island and 97 continental groups, we found that on average island groups had a ~2.5-fold higher founder intensity than estimated in continental groups (*P*=4×10^−4^, bootstrap test using 15 populations from each group) (Figure 3A).

**Figure 3.**
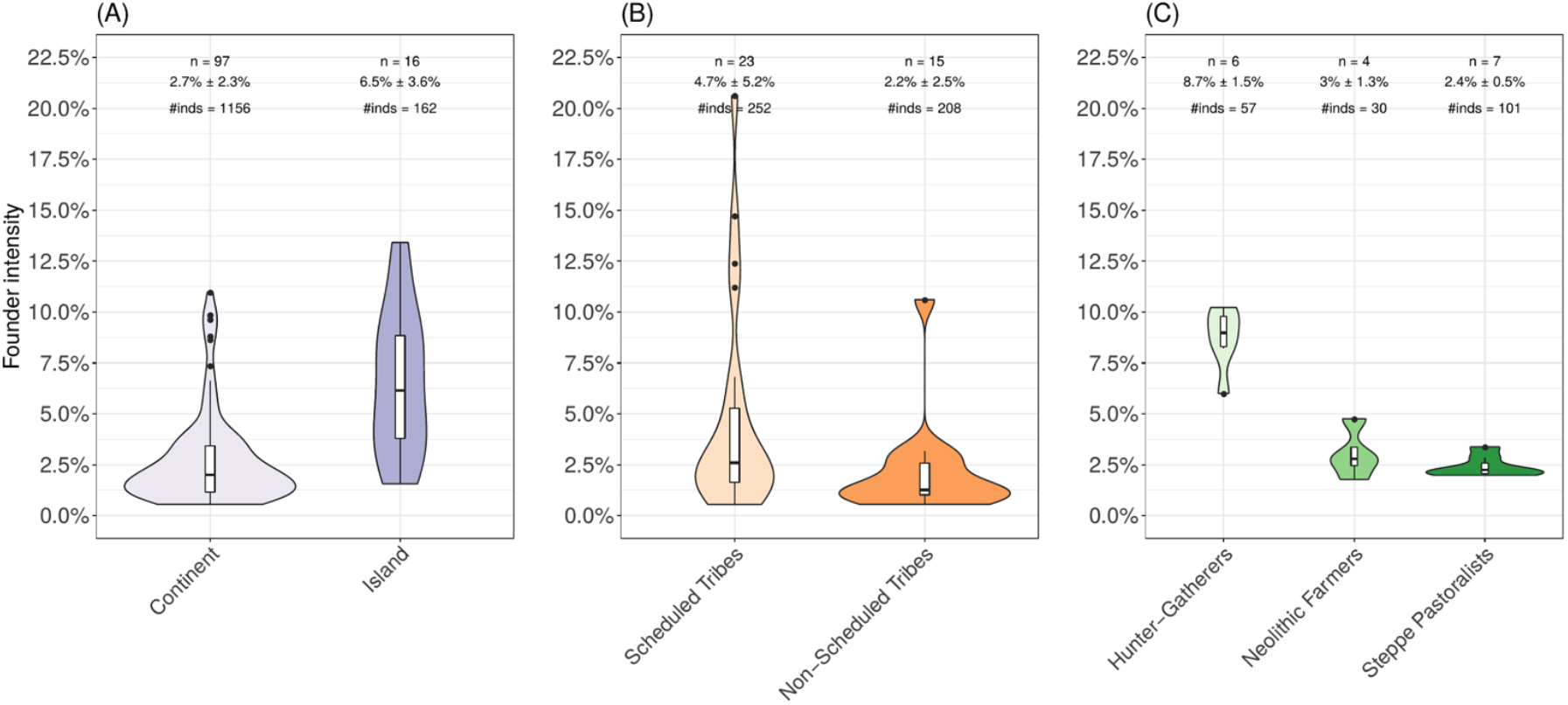
Geographic and cultural practices impact founder events in humans. We show the variation in estimated founder intensity as a violin plot across groups, classified in three main categories. Each violin plot includes a boxplot and the number of populations (*n*) in each group along with the mean ± standard deviation and the total number of individuals used in the analysis (# inds). *(A) Continental vs. island populations*. This plot shows the variation in founder intensities estimated for present-day populations in the HO37 dataset classified according to geography. *(B) Tribal vs. non-tribal groups in South Asia*. This plot shows the variation in founder intensities estimated in South Asian groups from the IndiaHO dataset, classified by caste group. *(C) Ancient hunter-gatherers, Neolithic Farmers and Steppe Pastoralists*. This plot shows the variation in estimated founder intensities for ancient groups in the HO42 dataset, classified based on their mode of sustenance. We note that in order to increase the number of groups in each category, we considered populations where the estimated founder age dated to less than 250 generations (default for other analyses was below 200 generations).

Previous studies have documented a pervasive history of founder events in Jewish groups^21,22^. To characterize the impact of founder events in diverse Jewish populations, we applied *ASCEND* to 11 groups, including Ashkenazi Jews (AJs). We observed significant evidence of founder events in most Jewish groups, except in Ethiopian and Turkish Jews (Figure S2). The strongest founder event was inferred in Cochin Jews (with intensity of ~2.4%) and the weakest in Iraqi Jews (intensity of ~1.1%). The most recent founder event among Jewish communities was also detected in Cochin Jews (~300 years ago), in agreement with previous results^23^. We inferred that a founder event in AJs occurred 37 generations ago with the 95% confidence interval (CI) of 23–51 generations, and an intensity of ~1.7% [1.3%–2.1%] (Figure S2). These results are consistent with previous studies based on whole genome sequences^21,24^, demonstrating the reliability of *ASCEND* to characterize founder events even with datasets containing few samples and only SNP genotypes.

The evidence of founder events in AJs has been linked to an increased risk of recessive diseases^25^. Across worldwide populations, we identified 50 groups that have a more extreme history of founder events than observed in AJs (Table S2). Below we describe these populations in detail.

#### Central and South America

In our dataset, all populations from Central and South America showed evidence for strong, recent founder events. The median estimated founder intensity was ~5.2%, which is almost three-fold higher than in AJs. The highest founder intensity (~12.2%) was observed in Rapa Nui (Easter Islands), followed by Pima (Arizona) and the populations of the Gulf of Mexico (Mixe, Mixtec, Mayan, Zapotec), and the lowest was estimated in Quechuas (~3.8%) (Figure 2). These founder events occurred recently–ranging between ~7 generations ago (in Mayans) to ~17 generations ago (in Mixe), translating to ~200-500 years ago–concordant with the European colonization and following smallpox epidemic in the Americas^26^.

Given that European colonization in the Americas was also accompanied by European admixture into Native American groups^26^, we were concerned that the similarity in dates of founder events could be driven by some technical artifact related to admixture. Admixture introduces long-range LD in the target population^27^, which could spuriously confound the long-range allele sharing correlation that we aim to characterize in *ASCEND*. As noted earlier, we did not observe any bias of admixture in simulations (Note S2.4). To further confirm this result empirically, we compared the dates of founder events (inferred using *ASCEND*) with previously published dates of admixture (inferred using *ALDER* or *GLOBETROTTER*). We found no overlap in the dates of admixture reported by *GLOBETROTTER* for Maya and Pima with the age of founder events in our study^28^. Importantly, comparison of the two dates in 64 worldwide populations did not yield significant correlations (*r*=-0.04, *P*=0.77 for *GLOBETROTTER* dates; *r*=-0.29, *P*=0.10 for *ALDER* dates) (Table S3)^28,29^. This suggests that the founder ages estimated with *ASCEND* do not seem to be impacted by the history of recent admixture.

#### Hunter-gatherers, indigenous and nomadic groups

We detected strong founder events in African hunter-gatherer groups, including Biaka pygmies, Mbuti pygmies and Ju/ʾhoan hunter-gatherers from South Africa (Figure 2). We inferred these founder events occurred recently and concomitantly in the three groups, ~20 generations ago (i.e. ~560 years ago). Our estimate for Mbuti pygmies of ~21 [18–24] generations ago is consistent but more precise than the previous published estimate of ~10–100 generations^30^. These results are interesting in light of the fact that most African populations, including hunter-gatherers, have high diversity and historically large population sizes. However, recent demographic events such as the Bantu expansion and destruction of hunter-gatherer habitats could have led to recent declines in population size of hunter-gatherer groups in Africa.

We also documented strong founder events in Bedouins and populations of the Yemen Desert (Figure 2). Previous studies have reported exceptionally high rates of consanguineous marriages in Bedouins^31,32^, with consequent founder events^33^. We note that our results should be interpreted as mainly reflecting the founder events in this group, as we minimize the impact of consanguinity by removing close relatives (see Methods). Indeed, we do not observe an excess of runs of homozygosity in these populations (as expected from consanguinity) and the timing of the founder events is not very recent (~20 generations ago) (Figure 2). We also detected strong founder events in most indigenous populations from North Asia, including Aleut, Chukchi, Eskimos and Yakut. The median age of the founder events in North Asia was estimated ~900 years ago, with a median intensity of ~6.4% (Figure 2). The strong founder events in these groups are consistent with their nomadic lifestyle and history of small population sizes and geographic isolation.

#### West Eurasia

We inferred strong founder events in the history of seven West Eurasian present-day populations, including in Western Europe (Basque country and Sardinia), Eastern and Northern Europe (Belarus, Estonia, Finland and Lithuania) and Anatolia (Turkey) (Figure 2). The timing of the founder events in Basque people (~1,700–2,500 years ago) overlaps with the Roman colonization of the Basque country. Strikingly, in Eastern and Northern Europe, the timing coincides with the spread of Steppe Pastoralists and the appearance of Beaker and Corded Ware cultures. The founder event in Sardinia occurred ~3,400-4,900 years ago and is coincident with the appearance of the Beaker culture on the island^34^. Similarly, in Eastern Europe (Lithuania, Estonia and Belarus) and Northern Europe (Finland), the estimated founder ages of ~4,500-5,000 years ago overlaps with the arrival of the Corded Ware culture^35,36^.

#### East Asia

We found evidence for significant founder events only in two East Asian groups, Lahu and Cambodians. In these groups, the estimated intensities were nearly twice as high as in AJs (> 3.5%). A previous study had documented strong drift in Lahu people, though it did not include any characterization of the founder event^37^. We inferred that founder events occurred very recently in the Lahu people (~310–420 years ago) and more anciently in Cambodians (3,340–6,520 years ago). The Lahu founder age coincides with their migration southward and to higher elevations, in response to the expansion of the Qing Dynasty^38^.

#### South Asia

Our *ASCEND* analysis revealed that 64% of the South Asian groups in the HO37 dataset have a history of founder events, including ~23% with higher intensity than AJs (Figure 2A). This is in line with results of a recent study that documented a widespread history of founder events in South Asia^18^. By measuring IBD sharing across individuals, Nakatsuka et al. (2017) identified that many groups in South Asia had a history of strong founder events, more extreme than in AJs or Finns. However, the timing and demographic events related to these founder events are unclear.

To learn about the age and intensity of founder events in South Asian groups, we applied *ASCEND* to the IndiaHO dataset^18^ to 116 groups that passed the filtering procedure. Similar to the HO37 dataset, 66 groups (57%) showed significant evidence of founder events, with ~25% of the groups experiencing more extreme founder events than AJs (Figure S3-S4). Comparison of our founder intensity with the IBD score reported in Nakatsuka et al. (2017) shows strong correlation (Pearson’s r=0.95, *P* < 10^−20^), highlighting the reliability of our inference without explicitly identifying IBD segments (Note S5.2). The strongest founder event was observed in the Onge population (~20.6%, ~10-fold higher than in AJs) that has a census size of ~100 individuals today (http://censusindia.gov.in/). Moreover, we found that Scheduled Tribes (4.7% [2.6%–6.8%]) have significantly stronger founder events than non-tribal groups (2.2% [0.9%–3.4%], comprising of Scheduled Castes, Backward Classes, Most Backward Classes, Other Backward Classes, Forward Castes and Upper Castes (Kruskal-Wallis test: *P*=0.024)) (Figure 3B), consistent with other studies^39,40^.

To understand the demographic events leading to such extreme founder events in South Asia, we investigated the timing of the founder events using *ASCEND*. Reich *et al*. (2009) had previously inferred the age of founder events in some South Asian groups using a similar statistic (Methods). For the 25 groups previously analyzed, we obtained statistically consistent results (Note S5.1). Across the 66 groups with evidence for founder events in the IndiaHO dataset, we inferred that the founder ages ranged between ~115 years ago (Gujjar) to ~3,210 years ago (Kondh_TN). In Gujjar, the very recent founder event within the past 10 generations suggests a possible confounding due to recent consanguineous marriages as we observed long runs of homozygosity in this population (Notes S5.3). There was no significant difference in the mean founder age across the speakers of the four major language families (i.e., Austro-Asiatic, Dravidian, Indo-European, Tibeto-Burmese) in India, suggesting that these events are uncorrelated to the spread of languages on the Indian subcontinent (Kruskal-Wallis test: *P*=0.21).

### History of founder events in ancient human past

To investigate founder events deeper in our evolutionary past, we applied *ASCEND* to ancient DNA samples from the Human Origins HO37 dataset. For some ancient groups (including hunter-gatherers, Neolithic Farmers and Steppe Pastoralists), we enriched our samples using the Humans Origins HO42 dataset (see Methods for filtering details, Table S1). We limited the analysis to the populations with at least 5 individuals. For ancient DNA samples, it is difficult to match the outgroup population as samples are from different timescales and geographic locations. To avoid any bias introduced by the choice of outgroup, we did not subtract the cross-population allele sharing correlation (but used only the within-population correlation to measure the parameters of the founder events). We performed simulations to test the reliability of this choice and found the inference was not impacted (Note S2.7.3).

We analyzed 111 worldwide ancient populations dated between ~100-15,000 years ago using radiocarbon dates or dates inferred based on the cultural context of the sample. Applying *ASCEND*, we discovered that ~34% of the ancient groups had significant evidence of founder events that occurred within 200 generations before the sampling age of the ancient specimen (Table S2). Accounting for the mean generation time and the sampling age of the ancient specimen, this translates to an average age of founder events of ~8,400 [7,327–9,516] years before present (BP). The intensity of the founder events was on average estimated to be 5.5% [4.2%–6.8%] that is about three times higher than in present-day AJs, suggesting that population sizes were fairly small in the past (Table S2). Investigating the geographic patterns of founder events, we observed that ancient populations from the Americas (n=6, intensity of 12.4% [9.2%–15.5%]) experienced significantly stronger founder events than those from Eurasia (n=31, intensity of 4.1% [3.3%–5.0%]) (*P* < 10^−20^, based on the bootstrap test using 5 populations from each group). Interestingly, we found that the timing of founder events in three of the oldest Native American groups (dated between 7000–10,000 years BP) was ~12,000 years BP which is consistent with the timing of the founding of the Americas^41^.

Previous analyses have suggested that Paleolithic, Mesolithic and Neolithic hunter-gatherers in Europe had a lower diversity than expanding Neolithic Farmers^42^. However, these results were based on low coverage data, where calling diploid genotypes is challenging^42,43^. We applied *ASCEND* to hunter-gatherers, Neolithic Farmers and Steppe Pastoralists and compared the intensity of founder events (a proxy for genetic diversity in the population). To maximize sensitivity for this analysis, we considered founder events that were dated between 0 and 250 generations before sampling (we note, this threshold differs from the usual setup where we report dates between 0–200 generations, which is the most reliable time range based on our simulations). We observed that founder events experienced by hunter-gatherers (~8.7% [7.5%–9.9%]) were significantly stronger than Neolithic Farmers (3.0% [1.8%–4.3%]) (*P* < 10^−20^, based on a bootstrap test using 4 populations from each group) or Steppe Pastoralists (2.4% [2.0%–2.8%]) (*P* < 10^−20^, based on bootstrap analysis with 4 populations per group) (Figure 3C). We note that the founder intensities were also significantly higher in hunter-gatherers when using the standard thresholds (*P* < 10^−20^, for both comparisons). The hunter-gatherers were also found to have experienced significantly older founder events (relative to their sampling age) compared to Neolithic Farmers or Steppe Pastoralists (*P*=0.023 and *P*=0.01, respectively) (Figure S5).

### History of recent and extreme founder events in canids

While human populations have been extensively studied, to demonstrate the broader applicability of our method, we also applied *ASCEND* to the domesticated dog species. Recent studies have suggested that dogs were domesticated between ~10,000–40,000 years ago^44–46^. The domestication of dogs was followed by the establishment of various breeds that likely occurred in the past 200 years, though the history of different breeds remains poorly understood. These events were accompanied by severe founder events and selection to propagate breeds with selected phenotypes^47^.

To reconstruct the history of founder events in dogs, we applied *ASCEND* to data from ~6,000 domesticated dogs (~200 breeds, including populations of village dogs and mixed breeds) from two publicly available datasets from the Sams^48^ and Hayward^49^ studies. We excluded individuals with evidence of recent inbreeding or close relatedness and considered only breeds with more than 5 individuals (see Methods). We used the standard parameters to run *ASCEND* with the exception that the maximum genetic distance was set to 40 cM (instead of 30 cM) as we observed long-range LD extending to large distances in many breeds. After filtering, we retained 50 populations belonging to 40 unique breeds and two village dog populations (see Methods, Table S4).

Application of *ASCEND* to canids revealed strong evidence of founder events in all breeds analyzed as well as two village dog populations, with average intensity of ~25.3% (ranging between 1.3% in Village_dogs_dr to 77.7% in Boxers) (Figure 4, Figure S6). The estimated founder events in dogs were substantially more extreme than in humans, with median intensity in dogs about ten-fold higher than in present-day humans. Interestingly, we found that founder intensities differed significantly across the traditional roles of dog breeds, with higher founder intensities estimated in traditionally agricultural or sedentary breeds (non-sporting dogs: 31.4% [23.5%–39.4%] or working dogs: 43.2% [31.6%–54.8%]) than breeds used for hunting or sports (hounds: 14.4% [12.5%–16.3%] or sporting dogs: 19.8% [17.2%–22.5%]) (Kruskal-Wallis test, *P*=6×10^−4^) (Figure 4, Table S5). We note this is unlikely to be due to sampling bias, as we obtained highly correlated results in the Sams and Hayward datasets (founder age: Pearson’s r=0.9, *P*=4×10^−4^, and founder intensity: r=0.92, *P*=1.9×10^−4^) (Table S6). Founder events in all breeds occurred very recently, within the past 25 generations (Figure 4B). The most recent occurred ~6 generations ago (in Gordon Setter) and the oldest occurred ~24 generations ago (in Bulldogs) (Figure 4B). Assuming a generation time of 3-5 years^50,51^, this translates to ~75–125 years ago, concordant with the timing of establishment of several modern breeds during the Victorian era^52^.

**Figure 4.**
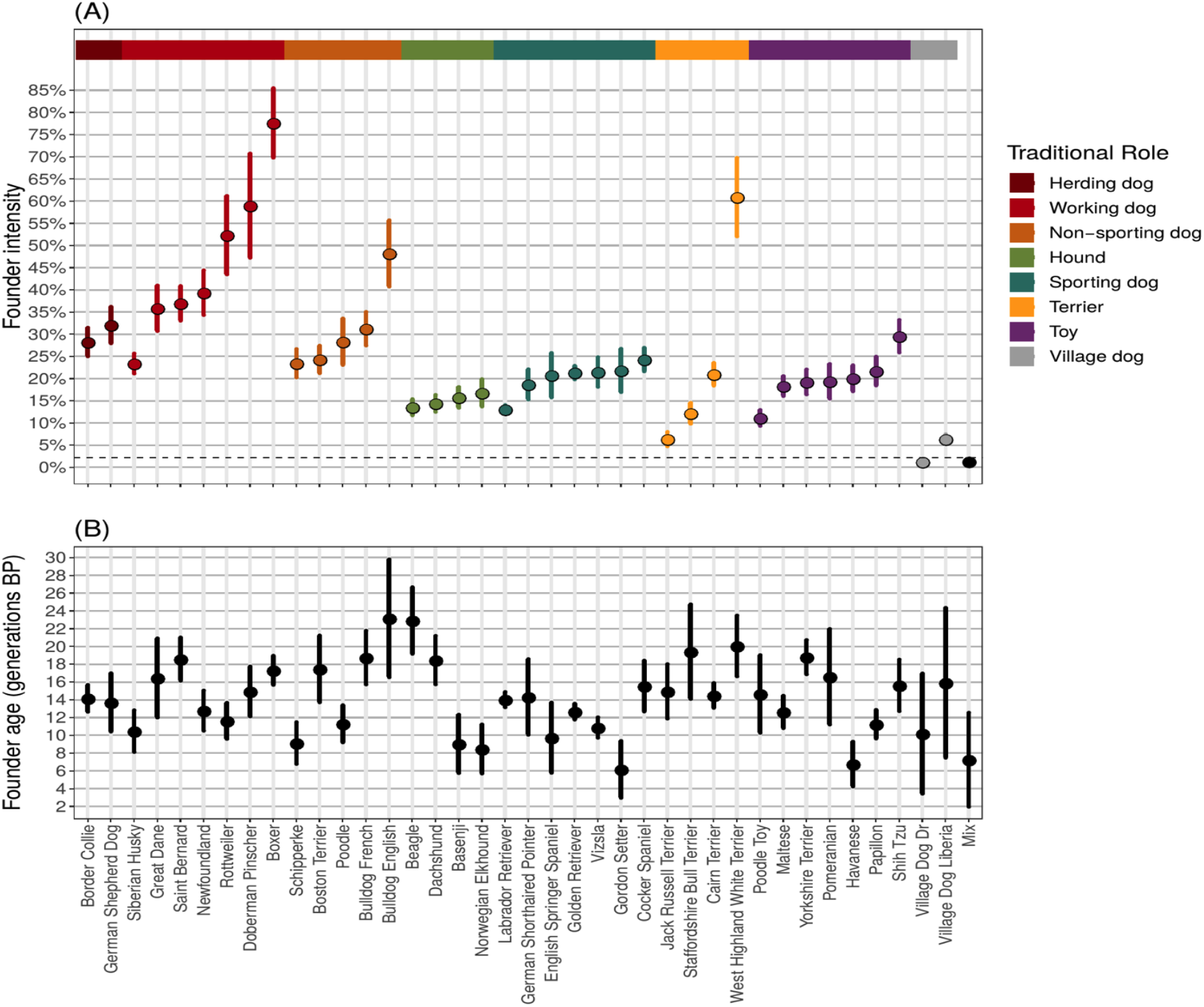
History of extreme founder events in dogs. Results of *ASCEND* for all dog breeds that passed the filtering criteria and showed evidence for significant founder events in the Hayward dataset (see Methods). The top colored ribbon represents the traditional roles of the breeds. Note that the role associated to village dogs is not defined and this population is shown in black. *(A) Distribution of the estimated founder intensities*. We show the mean founder intensity (point) and the associated 95% confidence interval. The breeds are ordered by their traditional role and in increasing order of estimated founder intensity. The horizontal dashed line shows the median of founder intensities estimated in the present-day human populations from the HO37 dataset (2.2%). *(B) Distribution of the estimated founder ages*. We show the estimated founder ages in generations, with their associated 95% confidence intervals.

## Discussion

We introduce *ASCEND*, a two-locus approach that leverages the allele sharing correlation across the genome to infer the time and strength of the founder event in a population. *ASCEND* is complementary to IBD-based methods^14,15^ such as DoRIS and IBDNe in its time range but is more flexible as it does not require phased data and hence is applicable to sparse datasets, including ancient DNA. By applying *ASCEND* to around 300 present-day and 100 ancient human populations, we document that more than half the groups in our study experienced a strong founder event during the past 10,000 years. To our knowledge, this is the first comprehensive survey of founder events across worldwide human populations and provides insights about the frequency and demographic processes underlying population bottlenecks during human evolution. We note that the sampling of human populations in our study was not random and the Human Origins dataset may be enriched for small, isolated groups and thus these frequencies should be interpreted with caution and could differ in other worldwide surveys of human populations.

We recover previously reported signals of founder events in AJs, Finns and South Asians, as well as uncharacterized events in many groups such as Papua New Guinean Nasioi and Gimi people, African hunter-gatherers, and Northeast Asian indigenous groups. Our results suggest that geographic isolation, modes of sustenance and cultural practices are notable predictors of founder intensity. Specifically, we document that populations living on islands have experienced founder events twice as strong as those estimated in continental groups. This could be because island populations were formed by small groups of individuals, or because they tend to maintain a small population size due to limited resources. Across diverse groups, our results highlighted that modes of sustenance (hunter-gatherers vs. farmers) and lifestyle choices (nomadic vs. settled) can have a substantial impact on shaping the genetic variation of the population. Finally, cultural practices such as endogamous marriages may contribute to the strong founder events observed in Ashkenazi Jews and South Asians.

The temporal distribution of founder events provides insights about the demographic events related to founder events. Notably, we found that many epidemics and major migrations of invading groups were accompanied by severe population bottlenecks. This is exemplified by founder events in the Americas related to the arrival of European colonizers, and the following smallpox epidemic (~1500 AD). In Europe, many founder events date to ~3,500–5,000 years ago, coincident with the spread of Steppe Pastoralists and the appearance of Beaker and Corded Ware cultures in Eastern and Northern Europe. This period has been referred to as the “Neolithic decline” as it was accompanied by permanent abandonment of previously heavily-populated settlements. The causes are still disputed and could have involved warfare with invading Steppe Pastoralists, environmental overexploitation^56^, and/or the spread of infectious diseases like plague^57^. For instance, recent findings suggest there were multiple lineages of *Yersinia pestis* expanding through early trade networks across Eurasia during this period^57^.

The majority of the South Asian populations in our survey showed evidence for strong founder events. Some groups had nearly ten-fold higher intensity than estimated in AJs, despite large census sizes (with over a million individuals) living today. This suggests low rates of gene exchange among neighboring groups in South Asia, likely due to strong rules about marrying within castes (endogamy) that has maintained the genetic signal of the founder events. We inferred the founder events occurred in the last ~3,500 years, which postdates the admixture between the Ancestral North Indians and Ancestral South Indians and the arrival of Steppe Pastoralists to South Asia^29,58^. We found tribal populations tend to have significantly stronger founder events than non-tribal groups. This history of extreme founder events in South Asians predicts a high burden of deleterious variants and increased risk of recessive diseases, as seen in Finns and AJs^7^.

Applying *ASCEND* to ancient human genomes, we found founder intensities varied substantially across populations but on average were significantly higher than estimated in present-day populations, as expected due to limitations of resources in the past. We found ancient hunter-gatherers had stronger founder events than Neolithic Farmers or Steppe Pastoralists. These results provide an independent line of evidence that the local population sizes increased following the Neolithic Expansion, consistent with previous results based on measuring genetic diversity in low coverage data^42^ and archaeological evidence^59–61^. These analyses highlight the power of our method to use the recombination clock to make reliable inferences with limited data available in ancient DNA studies.

We also applied *ASCEND* to the domesticated dog species. Among the 42 populations that we analyzed, all breeds and two village dog populations had significant evidence for a recent founder event, with intensities of around ten times stronger than the median strength of founder events in humans. These founder events may be due to: (i) *the sire effect* whereby only a very few highly valued individuals (based on selected phenotypic traits) are bred repeatedly and contribute disproportionately to the next generations^62^, or (ii) *inbreeding* that involves mating between closely related individuals. Such strong founder events can lead to high rates of homozygosity, and in turn increased risk of diseases. In accordance, some breeds like Boxers and Bulldogs that have among the strongest founder events in our analysis are also known to be affected by high rates of genetic disorders. For instance, arrhythmogenic right ventricular cardiomyopathy that is a cardiac disease causing sudden death is seen at high frequency in both breeds^63,64^. The temporal distribution of founder events in dogs (~75–125 years ago) overlaps with the Victorian era when a large number of modern dog breeds were created in Great Britain in the context of popularizing dog-fancying and showing^52^.

### Outlook

*ASCEND* is applicable to any species–for which genotype data and recombination maps are available. In this study, we analyzed autosomal data and inferred the sex-averaged founder events in humans and dogs. However, in many populations, the genetic diversity and demographic events impacting males and females differ markedly. By applying *ASCEND* to autosomes and X chromosomes separately, we can shed light on sex-specific founder event parameters. Further, applying *ASCEND* to admixed populations, whereby gene flow between divergent groups produces chromosomes of mosaic ancestry, we can gain insights about the population size of different ancestral groups, and learn about the dynamics of population migrations and invasions in human history and prehistory.

## Methods

### *ASCEND*: model and theory

*ASCEND* measures the distribution of LD across the genome at sites that are shared between pairs of individuals in a population to recover the age and intensity of the founder event. Below we describe the theoretical and implementation details of *ASCEND*.

#### Basic model and notation

Assume we have a target population *A* that has a history of recent founder event that occurred *T_f_* generations ago where the population size reduced from *N_o_* to *N_f_* (*N_f_* ≪ *N_o_*) (Figure 5). To estimate the properties of the founder event in *A*, we compute the average correlation in allele sharing *z_w_*(*d*) across pairs of individuals in *A* (referred to as *within-population allele sharing correlation*). Specifically, for each marker *a*, we record if pairs of individual share 0,1 or 2 allele(s) (i.e., number of IBS alleles), a quantity we denote as *N_i,j,a_*. For heterozygous sites, we assume the individuals share one allele, regardless of haplotype phase. We then average the correlation coefficient *r*(·, ·) across all pairs of neighboring markers (*a, b*) located at a genetic distance of *d* Morgans apart as follows:

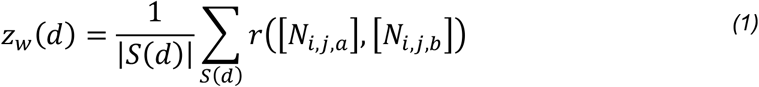

where *S_d_* is the set of unique pairs of single nucleotide polymorphisms (SNPs) located *d* Morgans apart and |*S*(*d*)| is the number of pairs of markers in the set *S_d_*, [*N_i,j,a_*] is the vector of allele sharing values across all possible pairs of individuals *i* and *j* (*i* ≠ *j*) in population *A*.

**Figure 5.**
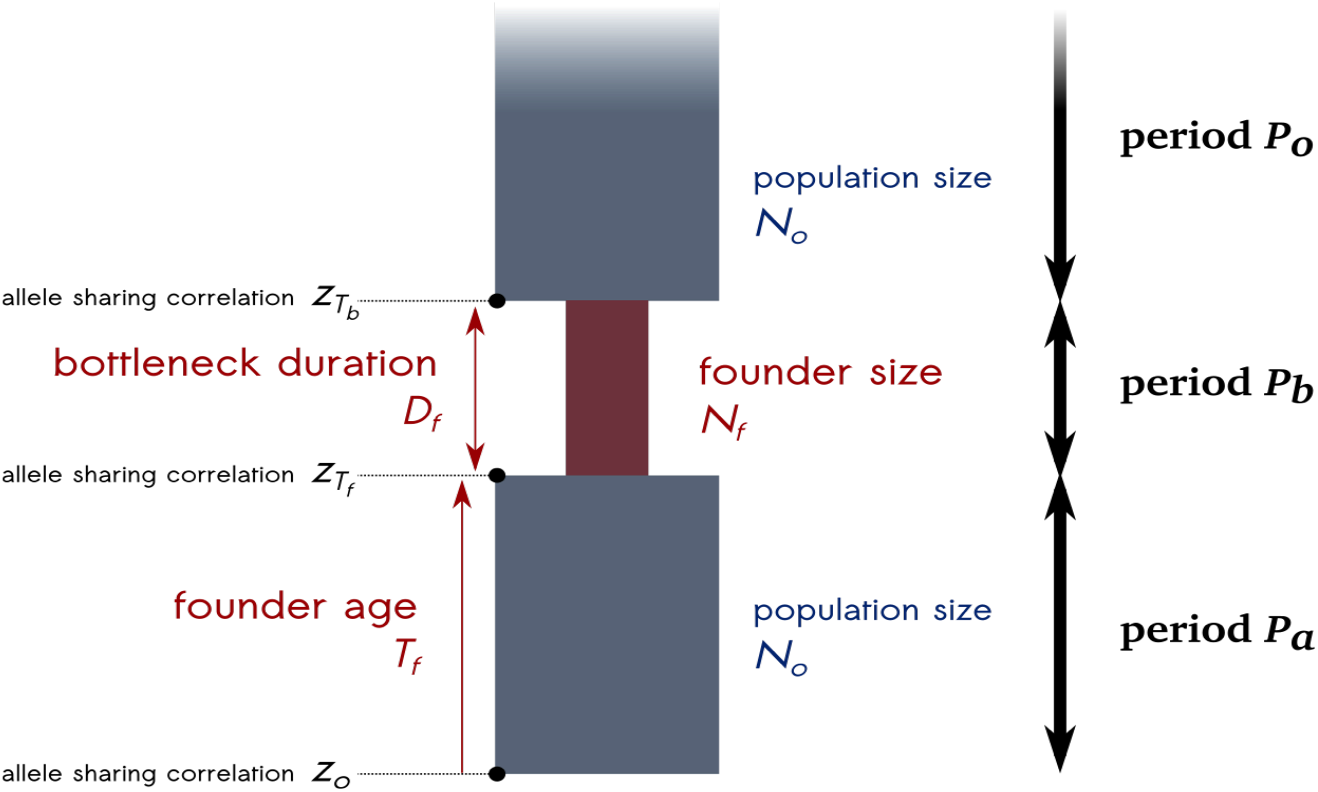
Demographic model with founder event. Consider a population with a history of founder event. This history can be divided into three main periods (from the most ancient to the most recent): a period *P_a_* where the population has a constant effective population size of *N_o_*, followed by a period *P_b_* where the effective population size reduces to *N_f_* for the duration of *D_f_* generations till *T_f_* generations before present. Then, during period *P_o_*, the population recovers and the population size returns to *N_o_*.

Some background LD or ancestral allelic correlation is expected at short genetic distances (even in the absence of a founder event) and can confound the signal of founder event in *A*. To minimize the impact of ancestral allele sharing, we compute the *cross-population allele sharing correlation z_c_*(*d*), which measures the correlation across pairs of individuals in *A*, and an outgroup population (*O*) (Figure 5) as follows:

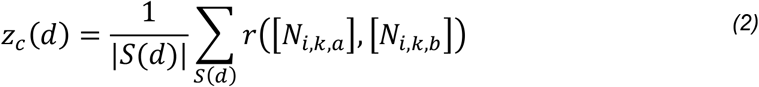

where *i* is the index of any individual from the target *A* and *k* the index of any individual from the outgroup *O*. The outgroup population is ideally a population that (i) has limited gene flow with the target in the recent past, and (ii) a population that separated from the target before the founder event. In simulations, we observed that the estimated founder parameters showed little sensitivity to the choice of the outgroup and hence we choose to pick a random set of *n* individuals to compute the ancestral LD in empirical analysis.

We define the corrected allele sharing correlation *z*(*d*) for any genetic distance *d* as:

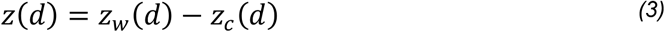

#### Relating allele sharing correlation to founder event parameters

Assume if individuals reproduce at random in a large population without selection or migration, the correlation across neighboring markers *z* is expected to decay due to recombination over successive generations^17^. Thus *z_n_*, the correlation in allele sharing (which is a proxy of the linkage disequilibrium) at generation *n*, can be expressed recursively as a function of the previous generation *z*_*n*+1_ (backward in time, time increases towards the past), the rate of recombination *d* and the effective population size *N_o_*. Specifically, the amount of allele sharing correlation between two loci is expected to decrease due to (i) crossing-overs that break down the allelic associations between markers, occurring as a function of the genetic distance *d* between neighboring markers and (ii) loss of allelic variation (ie. heterozygosity) by genetic drift due to finite population size, occurring at a rate, 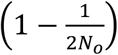. By combining these two effects, we get the recurrence relation for *z_n_* over one generation:

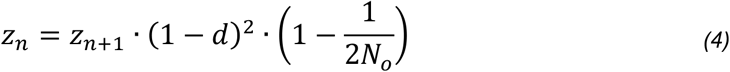

where (1 − *d*)^2^ is the probability that no recombination occurred between the two loci located *d* Morgans apart in the past generation and 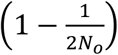 is the probability that the alleles at the locus *b* do not coalesce in the previous generation. For all analysis, we assume a single recombination event occurred between two neighboring markers. While in practice, more than one event could occur, we consider *d* to be small enough that the probability of multiple events is exceedingly low.

We now consider the period *P*_0_ (Figure 5) where the effective population size is constant, with size *N_o_* from present to the end of the bottleneck period, *T_f_* generations ago. We note *z*_0_ is the autocorrelation in allele sharing at the time of sampling (i.e., present = 0 generation ago). Thus we have, by recurrence:

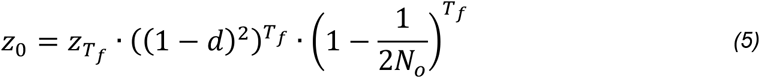

where *z_T_f__* is the allele sharing correlation at the end of the bottleneck period. For large *N_o_* and small genetic distances *d*, we can approximate this equation by:

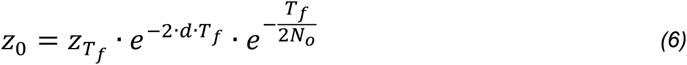

Assuming 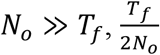, tends towards 0, so we can write:

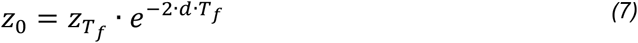

The allele sharing correlation thus decays exponentially with a rate proportional to the age of the founder event^16^.

Now, considering the bottleneck period *P_b_* (Figure 5) we can express *z_T_b__* as a function of *z_T_b__*, the correlation before the onset of the bottleneck (*T_b_ = T_f_ + D_f_* by definition of our model):

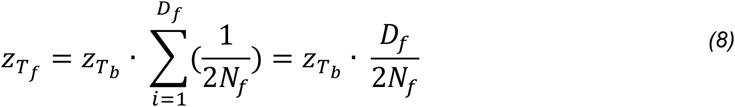

where *z_T_b__* is the correlation between two adjacent loci before the onset of the bottleneck and 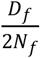 is the probability that the two alleles at the second locus share a common ancestor in any of the generations during the bottleneck period *P_b_*.

We define the intensity of the bottleneck, 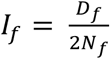. Thus, the equation above can be written as a function of the founder intensity as:

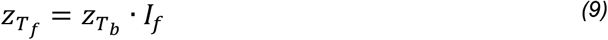

Now consider the period *P_a_* before the bottleneck (Figure 5). Our model assumes that the demography is at equilibrium before the onset of the bottleneck, with the population size approximately equal to *N_o_* so that *z_T_b__* equals to the probability that the two adjacent loci do not share a common ancestor for any generation till the upper limit of the genome-wide expected time to the most recent common ancestor between any two loci, i.e., (*T_MRCA_*) = 2*N_o_* (if and only if *N_f_ < N_o_*), thus:

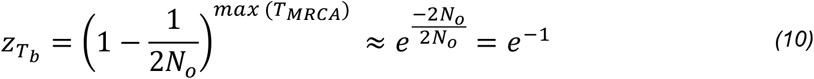

Integrating the information from the above equations, we obtain the allele sharing correlation computed at the sampling time (*z*_0_) as a function of founder age *T_f_* and founder intensity *I_f_* as follows:

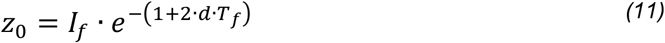

#### Implementation of ASCEND

We estimate the allele sharing correlation using two approaches: a Naïve approach and an approach based on fast Fourier transform (FFT). In the Naïve approach, we perform pairwise correlations across hundreds of thousands of markers (*n*) in the genome. This has a runtime of *O*(*n*^2^), which can be exceedingly slow in large datasets. In a recent study, Loh et al. (2013) introduced an approach to compute genetic correlations using FFT to dramatically speed up the computation of admixture LD^65^. Briefly, the method performs an algebraic transformation of the admixture LD statistic and computes the FFT convolution in discrete equally sized bins (referred to as mesh points). This provides a speedup from *O*(*n*^2^) to *O*(*n log log n*), which reduces the typical runtimes from hours to seconds. Here, we extend this approach to compute allele sharing correlation in the target and outgroup populations. Below we describe the details of the two approaches.

##### Naïve implementation

We first construct an allele sharing matrix whose elements contain the counts 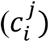 of alleles shared for any *j*^th^ pair of individuals at any SNP *i*. This matrix has dimensions *N×P*, where *N* is the number of SNPs and *P* is the number of individual pairs. We then compute the correlation across pairs of SNPs using the following equation:

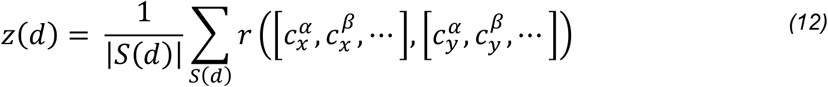

where *z*(*d*) is the allele sharing correlation for a genetic distance *d* cM, *S*(*d*) is the set of pairs of SNPs (*x, y*) located *d* cM apart, |*S*(*d*)| is the size of this set, 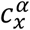 is the allele sharing where *α* is the index of the pair of individuals and *x* is the SNP index. In the above equation, *r*(·, ·) is the Pearson’s correlation coefficient.

##### FFT-based implementation

The FFT implementation calculates the autocorrelation in alleles shared between each pair of individuals along mesh points. Mesh points are equally-spaced genetic positions located every *δ* centiMorgans along the genome (such that *δ* ≪ *d*). Specifically, the FFT approach computes the following quantity:

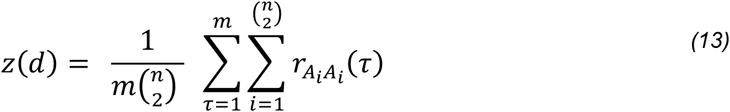

where *n* is the number of individuals, *m* is the number of mesh points in the bin *d, A_i_* is the vector of allele sharing in the individual pair *i* across all mesh points (the length of *A_i_* is thus equal to the number of mesh points), and *r_A_i_A_i__*(*τ*) is the autocorrelation coefficient for genetic distance bins (non-overlapping shifts) of *τ* cM.

We note that the use of the mesh is the only source of approximation in the FFT implementation compared to the Naïve approach. The genetic distance between mesh points, *δ*, depends on the dataset size and leads to a trade-off between runtime and accuracy. To improve accuracy, *δ* should be smaller than the bin size *d* used in the Naïve implementation. Empirically, we observe that setting the mesh size as 0.001 cM gives nearly identical allele sharing correlation values between the Naïve and FFT implementations (Notes S2.8). With this mesh size, we obtain a runtime speed up of a hundred-fold, allowing the analysis of population genome-wide data to be completed in a few minutes rather than taking hours with the Naïve approach. Further details of the FFT implementation are provided in Notes S1.

#### Inference of parameters of the founder event

To estimate the age and intensity of a founder event in a target population *A, ASCEND* computes the allele sharing correlation *z*(*d*) as the difference between the within-population allele sharing *z_w_*(*d*) across pairs of individuals in *A* and the cross-population correlation *z_c_*(*d*) across the individuals in the target and outgroup populations. In the empirical datasets (except ancient DNA samples), outgroup populations are defined as a random set of 15 non-overlapping individuals, sampled across all the populations present in the curated and filtered dataset. For ancient DNA samples, we do not use an outgroup as it is difficult to match samples based on the time of sampling and geographical location. All other parameters are similar for ancient and present-day samples, even when performing the analysis with pseudo-homozygous samples. We ran *ASCEND* for sites having at least one variant present in at least one sample across the target and outgroup populations, and for pairs of SNPs with at least one non-missing genotype in common across the individuals. We compute *z*(*d*) for genetic distances ranging from 0.1 to 30 cM, with a bin size of 0.1 cM and setting the number of mesh points per bin to 100 for the FFT computation and then fit an exponential distribution to estimate the rate and amplitude of *z*(*d*) using non-linear least squares:

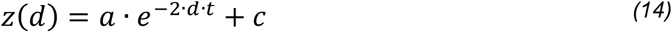

Here, the affine term, *c*, is used to account for noise in the fit of the exponential decay, which is a function of the sampling variance. We then simultaneously estimate the parameters of interest: 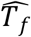, the age of the founder event and 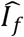, the intensity of the founder event as follows:

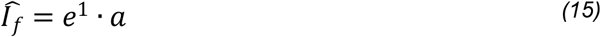

and

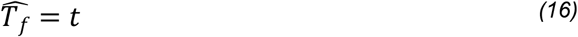

In order to assess the quality of the exponential fit, we calculate a normalized root-mean-square deviation (NRMSD) between the empirical allele sharing correlation values *z* and the fitted ones 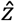, across the *D* genetic distance bins:

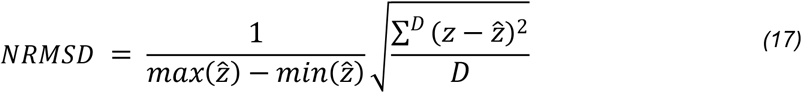

Based on the empirical distribution of NRMSD values across all the present-day human populations for which a fit was obtained, we use a cut-off of NRMSD=0.29 to identify reliable exponential fits (Note S1.3), though in practice other thresholds may also be valid. We compute standard errors by performing a weighted block-jackknife procedure^66^, considering each chromosome as an independent block and setting the weights proportional to their respective SNP counts. For all analyses, we report the 95% confidence interval of the estimated parameters.

Unless specified otherwise, founder events were considered significant if the following criteria are met: (i) the 95% confidence intervals of the estimated founder age and intensity does not include 0, (ii) the estimated founder age is lower than 200 generations and its associated standard error is lower than 50 generations, (iii) the estimated founder intensity is greater than 0.5%, and (iv) the NRMSD is lower than 0.29. These thresholds are based on simulation results (see Note S1 and S2 for details).

#### Software availability

*ASCEND* is written in Python and is available for download (with tutorial and examples) at: https://github.com/sunyatin/ASCEND. The released version includes both the Naïve and FFT implementations.

### Simulations

In order to investigate the accuracy of *ASCEND* in estimating the age and intensity of the founder event, we performed simulations under a range of demographic models using the coalescent simulator *msprime*^67^. All simulations involved at least two populations (one target, *A* and one outgroup, *O*) which diverged 1,800 generations ago (Figure 1). Unless stated otherwise, for each simulation and each population, we generated data for 30 haploid individuals for a total of 20 chromosomes of size 50 megabases each. We assumed a mutation rate of 1.2×10^−8^ per base pair per generation^68^ and a recombination rate of 1×10^−8^ per base pair per generation^69^. Except during the founder event, the two populations had an effective population size of 12,500. We combined two haploid chromosomes at random without replacement to generate one diploid chromosome. Details are described in the Notes S2.

### Datasets

#### Human datasets

We applied *ASCEND* to data from worldwide human populations containing both present-day and ancient individuals using four datasets: (i) Human Origins version 37.2 dataset (HO37): This datasets comprises of 5,637 present-day and 2,104 ancient individuals genotyped on the Human Origins array (499,158 SNPs) (https://reich.hms.harvard.edu/datasets); (ii) IndiaHO: This includes 1,662 individuals from 249 ethno-linguistic groups genotyped on the Human Origins array^18^; (iii) Human Origins version 42.4 (HO42): We used 27 ancient DNA populations including 6 hunter-gatherer, 10 Neolithic Farmer and 10 Steppe Pastoralist populations from the recent release of the Human Origins Dataset (https://reich.hms.harvard.edu/datasets); (iv) Behar dataset: We used 21 present-day Ashkenazi Jewish individuals genotyped on the Illumina 610K and 660K bead arrays^19^. We used the genetic positions from the 1000 Genomes recombination map^70^.

For all four datasets, we limited our analysis to all groups with a minimum of 5 samples. We removed individuals that were marked as duplicates, low coverage or outliers in the original studies. We filtered the datasets to remove any potential close relatives, i.e. individuals defined as having either (i) pairwise genomic sharing (*π*) greater than 0.45 with any other individual, or (ii) having both *π* > 0.125 (third-degree relatives) and at least one IBD segment greater than 65 cM (on average, half the chromosome). Note, *π* is an estimator of the proportion of genome-wide IBD (for example, *π* = ½ for first-degree relatives). We computed pairwise *π* using PLINK v1.90b6.2 *genome* module^71^. For identification of IBD segments, we first phased the samples from each dataset using EAGLE 2.4.1 with default parameters^72^, using the 1000 Genomes Project phased samples as a reference panel to increase the phasing accuracy^70^. The IBD segments were then called using GERMLINE 1.5.3^73^ with default parameters (-*bits 75, -err_hom 0, -err_het 0, -min_m 3*) in the -*genotype* extension mode. We applied the HaploScore algorithm to remove false positive IBD segments using the recommended parameters, namely genotype error of 0.75%, switch error of 0.3% and the threshold matrix for a mean overlap of 80%^74^. For details of data curation, see Note S3.

#### Dog datasets

We downloaded two publicly available genome-wide datasets of dogs: (i) Sams dataset: This includes 1,792 individuals from 11 breeds genotyped on 175,123 SNPs^48^, and (ii) Hayward dataset: This contains 4,342 individuals from 198 breeds genotyped on 160,723 SNPs^49^. We used the genetic positions from CanFam3.1 genetic map^75^.

For both datasets, we filtered out SNPs and individuals with missingness greater than 1% and 5% respectively. To remove close relatives, we computed *π* between all pairs of individuals using PLINK v1.90b6.2 *genome* module^71^. For each pair of individuals with *π* greater than 0.45, we excluded one of the individuals out of the pair. Compared to the human datasets, we did not include the filter criterion based on the IBD segments as phasing would be less reliable with small sample size and without the use of a reference panel^76^. Instead, to control for inbreeding, we excluded any individual with at least one run of homozygosity (ROH) longer than 30 cM (on average, half the chromosome). To compute ROH, we used PLINK *homozyg* module. We limited our analysis to all groups with a minimum of 5 samples.

## Acknowledgments

We are grateful for support from the Burroughs Wellcome Fund Careers at the Scientific Interface and the Sloan Research fellowship awarded to PM. RT was supported by the UC Departmental Startup funds awarded to PM. We thank Monty Slatkin, Shai Carmi, Shamam Waldman, Manjusha Chintalapati and members of the Moorjani lab for helpful discussions. We thank Nick Patterson for help with implementation of the fast Fourier transform for computing allele sharing correlations, and Kumarasamy Thangaraj and Pratheusa Machha for information about the group affiliation of present-day South Asian populations. We thank Monty Slatkin, Nick Patterson, Sonal Singhal, Kelsey Witt, Shai Carmi and Emilia Huerta-Sanchez for helpful comments on the manuscript.

## Supplementary Materials

### Supplementary Figures

**Figure S1 - Decay curves for present-day human populations from the HO37, Behar and IndiaHO datasets.** The x-axis represents the genetic distance (in cM) and the y-axis represents the average allele sharing correlation. The legend shows the mean and 95% confidence interval for the founder age (*T_f_*) in generations before present (gBP) and the founder intensity (*I_f_*), as well as the NRMSD (see Methods). The panels are greyed when the exponential fitting failed or when the evidence for the founder event was not significant (see Methods). The specific reason is highlighted in red in the legend.

**Figure S2 - Distribution of the founder age and intensity estimated in present-day Jewish communities from the HO37 dataset.** The x-axis shows the analyzed Jewish groups, ordered by increasing founder intensity. The y-axis shows the estimated founder intensity. Vertical bars show the 95% confidence intervals. The colors are proportional to the founder age that was estimated in generations.

**Figure S3 - History of founder events in present-day South Asian populations from the IndiaHO dataset.** Panel (A) shows the distribution of founder intensity and panel (B) shows the distribution of founder ages for all present-day populations that passed our filtering criteria and showed evidence for a significant founder event in the IndiaHO dataset (see Methods). Each point represents a population and the shape of the points indicates whether the group lives on an island (triangle) or in a continental region (circular). The top colored ribbon represents the linguistic affiliation of the group. *(A) Distribution of the estimated founder intensities*. We show the founder intensity (point) and the associated 95% confidence interval. The populations are ordered by their linguistic affiliation and by increasing order of estimated founder intensity. The black horizontal line shows the inferred founder intensity in Ashkenazi Jews in the IndiaHO dataset (2.0%). Populations that have experienced a founder intensity significantly stronger than in Ashkenazi Jews (i.e. whose founder intensity is greater than the upper-bound of the 95% confidence interval of Ashkenazi Jews) are colored in gold, else they are shown in black. *(B) Distribution of the estimated founder ages*. We show the estimated founder ages and the associated 95% confidence intervals. The estimated ages were converted from generations to years assuming an average generation time of 28 years^20^.

**Figure S4 - Decay curves for ancient human genomes from the HO37 and HO42 datasets.** The x-axis represents the genetic distance (in cM) and the y-axis represents the average allele sharing correlation. The legend shows the mean and 95% confidence interval for the founder age (*T_f_*) in generations before sampling age of the ancient specimen (gBS) and the founder intensity (*I_f_*), as well as the NRMSD (see Methods). The panels are greyed when the exponential fitting failed or when the evidence for the founder event was not significant (see Methods). The specific reason is highlighted in red in the legend.

**Figure S5 - History of founder events in ancient human populations from the Human Origins v37 (HO37) and v42 (HO42) datasets.** Panel (A) shows the distribution of founder intensity, and panel (B) shows the distribution of founder ages for all ancient populations that passed our filtering criteria in the HO37 and HO42 datasets and showed evidence for a significant founder event (see Methods). Each point represents a population and the colour of the points indicates the geographical location of the population. *(A) Distribution of the estimated founder intensities*. We show the mean founder intensity (point) and the associated 95% confidence interval. The populations are ordered by their subcontinent and in increasing order of founder intensity. The black horizontal line shows the inferred founder intensity in Ashkenazi Jews (1.7% [1.3%–2.1%]). *(B) Distribution of the estimated founder ages*. We show the estimated founder ages and the associated 95% confidence intervals. The estimated ages were converted from generations to years before present by using a generation time of 28 years^20^ and by adding the sampling age of the specimens (shown as the black diamond-shaped points).

**Figure S6 - Decay curves for all dog breeds surveyed in the Sams and Hayward datasets.** The x-axis represents the genetic distance (in cM) and the y-axis represents the average allele sharing correlation. The legend shows the mean and 95% confidence interval for the founder age (*T_f_*) and the founder intensity (*I_f_*), as well as the NRMSD (see Methods). The panels are greyed when the exponential fitting failed or when the evidence for the founder event was not significant (see Methods). The specific reason is highlighted in red in the legend.

### Supplementary Tables

**Table S1 - Data curation for all present-day and ancient human populations from the HO37, IndiaHO, Behar and HO42 datasets.**

**Table S2 - Founder event estimates for all present-day and ancient human populations from the HO37, IndiaHO, Behar and HO42 datasets.**

**Table S3 - Comparison of founder event ages estimated in present-day human populations from the HO37 dataset with published dates of admixture.**

**Table S4 - Data curation for dog breeds from the Sams and Hayward datasets.**

**Table S5 - Founder parameter estimates for dog breeds from the Sams and Hayward datasets.**

**Table S6 - Comparison of the founder parameter estimates for dog breeds between the Sams and Hayward datasets.**

