## Supplementary Notes and Figures for "Reconstructing the history of founder events using genome-wide patterns of allele sharing across individuals"

**S1. Implementation of *ASCEND*** 1

**S2. Simulations to test the performance of *ASCEND*** 9
- **S2.1 Single-generation epoch model** 9
- **S2.2 Multi-generation epoch model** 12
- **S2.3 Two-epoch bottleneck model** 14
- **S2.4 Model with founder event and admixture** 16
- **S2.5 Gradual exponential growth model** 18
- **S2.6 No recovery founder event model** 20
- **S2.7 Robustness of the inference to data quality issues** 22
  - **S2.7.1 Impact of sample size** 22
  - **S2.7.2 Impact of missing data** 24
  - **S2.7.3 Impact of ancient DNA data features** 26
- **S2.8. Comparison between Naïve and FFT implementation** 28
- **S2.9. *msprime* commands** 29

**S3. Data curation for human datasets** 37

**S4. Comparison of overlapping groups across human datasets** 40

**S5. Comparison of our results with published estimates** 44
- **S5.1 Comparison of dates of founder event with Reich et al. 2009** 44
- **S5.2. Comparison of founder event intensity and IBD scores with Nakatsuka et al. (2017)** 48
- **S5.3. Comparison of founder intensity with the ROH statistic** 51

**S6. History of founder events in dogs** 54

**Figure S1** 57

**Figure S2** 84

**Figure S3** 86

**Figure S4** 88

**Figure S5** 99

**Figure S6** 101

### S1. Implementation of *ASCEND*

We implemented *ASCEND* (Allele Sharing Correlation for the Estimation of Nonequilibrium Demography) in two ways: a Naive approach and a Fast Fourier Transform (FFT) approach. We illustrate these two methods with an example containing three SNPs (say, $s_1$, $s_2$ and $s_3$) and three individuals (referred to as, $A$, $B$ and $C$).

#### S1.1 Naive implementation of *ASCEND*

In the Naive implementation, we first construct an allele sharing matrix whose elements are counts ($c_i^j$) of the alleles shared between a pair $j$ of individuals at each SNP $i$. Specifically, this is a $N \times P$ matrix, where $N$ is the number of SNPs and $P$ is the number of individual pairs.

$$\begin{array}{c} \\ s_1 \\ s_2 \\ s_3 \end{array} \begin{array}{c} \begin{array}{ccc} A,B & A,C & B,C \end{array} \\ \left[ \begin{array}{ccc} c_1^\alpha & c_1^\beta & c_1^\gamma \\ c_2^\alpha & c_2^\beta & c_2^\gamma \\ c_3^\alpha & c_3^\beta & c_3^\gamma \end{array} \right] \end{array}$$

For simplicity, we consider that the three SNPs are equally spaced, $d = 0.001$ cM apart, with the genetic positions of $s_1$, $s_2$ and $s_3$ as 0.001, 0.002 and 0.003 cM respectively.

The Naive implementation loops over the SNP pairs, and calculates the following equation:

$$\boxed{z(d) = \frac{1}{|S(d)|} \sum_{S(d)} r([c_x^\alpha, c_x^\beta, ...], [c_y^\alpha, c_y^\beta, ...])} \tag{1}$$

where $S(d)$ is the set of SNP pairs located $d$ cM apart, $|S(d)|$ is the size of this set. To simplify the computations, we standardize the allele sharing vectors:

$$s_i' = \frac{s_i - \bar{s}_i}{\sigma_{s_i}} \tag{2}$$

And we then compute the Pearson's population correlation coefficient as follows, considering the genetic distance bin [0.001-0.003] cM for our application:

$$
\begin{aligned}
z &= \frac{1}{3} \cdot \Big( r(s'_1, s'_2) + r(s'_1, s'_3) + r(s'_2, s'_3) \Big) \\
&= \frac{1}{3} \cdot \Big( \frac{cov(s'_1, s'_2)}{\sigma_{s'_1}\sigma_{s'_2}} + \frac{cov(s'_1, s'_3)}{\sigma_{s'_1}\sigma_{s'_3}} + \frac{cov(s'_2, s'_3)}{\sigma_{s'_2}\sigma_{s'_3}} \Big)
\end{aligned}
\tag{3}
$$

Since the $s'_i$ vectors are standardized, i.e., with mean = 0 and variance = 1, this equation reduces to:

$$
\begin{aligned}
z &= \frac{1}{3} \cdot \Big( cov(s'_1, s'_2) + cov(s'_1, s'_3) + cov(s'_2, s'_3) \Big) \\
&= \frac{1}{3} \cdot \Big( \frac{\sum s'_1 s'_2}{3} - \bar{s'_1}\bar{s'_2} + \frac{\sum s'_1 s'_3}{3} - \bar{s'_1}\bar{s'_3} + \frac{\sum s'_2 s'_3}{3} - \bar{s'_2}\bar{s'_3} \Big) \\
&= \frac{1}{9} \cdot \Big( \sum s'_1 s'_2 + \sum s'_1 s'_3 + \sum s'_2 s'_3 \Big)
\end{aligned}
\tag{4}
$$

Thus:

$$
\boxed{9z = (c_1^{\alpha'} c_2^{\alpha'} + c_1^{\beta'} c_2^{\beta'} + c_1^{\gamma'} c_2^{\gamma'}) + (c_1^{\alpha'} c_3^{\alpha'} + c_1^{\beta'} c_3^{\beta'} + c_1^{\gamma'} c_3^{\gamma'}) + (c_2^{\alpha'} c_3^{\alpha'} + c_2^{\beta'} c_3^{\beta'} + c_2^{\gamma'} c_3^{\gamma'})}
\tag{5}
$$

#### S1.2 FFT implementation of *ASCEND*

In brief, the FFT approach provides a significant speedup to Naive implementation because it performs an FFT convolution to compute pairwise LD. To implement this, we calculate the autocorrelation of alleles shared between each pair of individuals along equally-spaced mesh points (the "grid mesh") in the genome. The main implementation difference with the Naive implementation is the assignment of SNPs to these pre-defined mesh points, which we describe below.

##### S1.2.1 Grid mesh

The use of FFT in *ASCEND* requires dividing the genome into equally-spaced bins such that data points are distributed evenly located $\delta$ cM apart across the genome (Figure S1.2.1). To improve accuracy, $\delta$ should be smaller than the genetic bin size $d$ used in the Naive approach, i.e. $\delta = d/m$ where $m \in \mathbb{N}^*$ is defined as the number of grid points per genetic distance bin. Typically, $\delta$ is set to be very small, say $10^{-3}$ cM.

Each SNP, located at a particular genetic position $g$ cM, is then assigned to the nearest mesh point with genetic position $g_m$ so that $g \in [g_m; g_m+\delta]$. In the case where multiple SNPs map to the same mesh point $g_m$, only the SNP with the lowest genetic position is retained, the others are excluded (note that if $\delta$ is small, this exclusion would affect only a very minor proportion of the SNPs in the dataset). Given the density of markers, some regions $[g_m; g_m + \delta]$ may not contain any SNPs. These mesh regions are considered as "missing" and discussed in the next section.

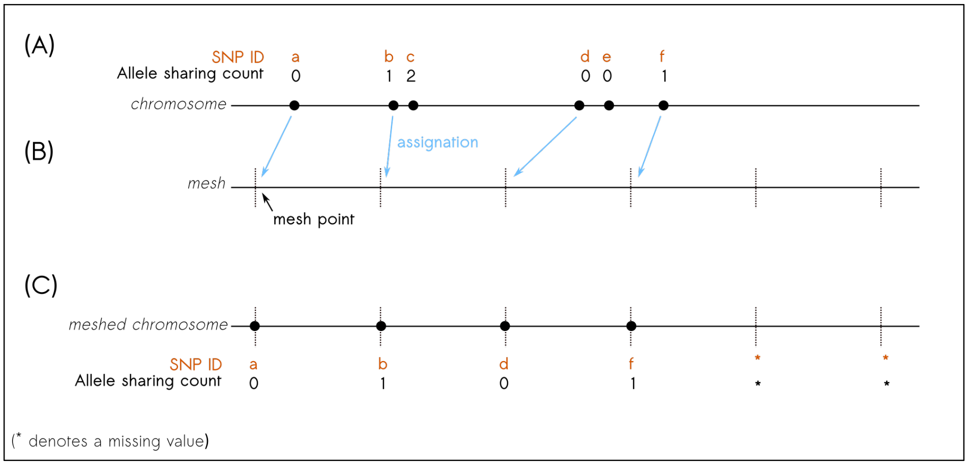

Figure S1.2.1: **Representation of the mesh procedure.** (A) A chromosome segment with SNPs (with IDs $a$, $b$, $c$, $d$, $e$, $f$) and their associated allele sharing count for a given pair of individuals. (B) A mesh is placed upon the chromosome (vertical dotted lines) and each SNP is assigned to a mesh point (the first SNP, a, is on the right of the mesh point). (C) The resulting "meshed chromosome" with the associated vector of allele sharing counts that will be used for the FFT computation. Comparison of (A) and (C) shows the difference in the non-meshed and meshed allele sharing vectors.

##### S1.2.2 Computing allele sharing correlation using FFT

In the FFT implementation, we loop over all possible mesh points and all pairs of individuals. In each loop, we estimate the autocorrelation of the allele sharing vector, instead of the Pearson's correlation coefficient as in the Naive implementation. Thus, $z(d)$ becomes:

$$\boxed{z(d) = \frac{1}{m\binom{n}{2}} \sum_{\tau=1}^{m} \sum_{i=1}^{\binom{n}{2}} r_{A_i A_i}(\tau)} \qquad (6)$$

Where $n$ is the number of individuals, $m$ is the number of mesh points in the distance bin $d$, $A_i$ is the vector of allele sharing for the pair of individuals indexed by $i$ (the length of $A_i$ is equal to the number of mesh points), and $r_{A_i A_i}(\tau)$ is the convolution function with a shift of $\tau$ cM. Note that here we are summing over the convolution coefficients for the values of $\tau$ which leads to the mesh points being spaced by $[d; d+\Delta d]$. The convolution computation is performed in Python using the function *correlate* from SciPy v1.5.2 in *fft* mode.

Let us now consider the allele sharing matrix that was described earlier for the Naive approach. Since the SNPs in our toy example are equally spaced (at 0.001 cM intervals), they meet the criteria for FFT and hence we use them as mesh points for the computation of autocorrelation. Below we describe the transposed and standardized version of the allele sharing matrix for FFT:

$$\begin{array}{c} \\ A,B \\ A,C \\ B,C \end{array} \begin{array}{c} \begin{array}{ccc} s_1 & s_2 & s_3 \end{array} \\ \begin{bmatrix} c_1^{\alpha\prime} & c_2^{\alpha\prime} & c_3^{\alpha\prime} \\ c_1^{\beta\prime} & c_2^{\beta\prime} & c_3^{\beta\prime} \\ c_1^{\gamma\prime} & c_2^{\gamma\prime} & c_3^{\gamma\prime} \end{bmatrix} \end{array}$$

The autocorrelation is calculated as such:

$$\begin{aligned} &\text{For } \tau = 1,\ z = \frac{1}{3} \cdot \frac{1}{3}\Big((c_1^{\alpha\prime} c_2^{\alpha\prime} + c_2^{\alpha\prime} c_3^{\alpha\prime}) + (c_1^{\beta\prime} c_2^{\beta\prime} + c_2^{\beta\prime} c_3^{\beta\prime}) + (c_1^{\gamma\prime} c_2^{\gamma\prime} + c_2^{\gamma\prime} c_3^{\gamma\prime})\Big) \\ &\text{For } \tau = 2,\ z = \frac{1}{3} \cdot \frac{1}{3}\Big((c_1^{\alpha\prime} c_3^{\alpha\prime}) + (c_1^{\beta\prime} c_3^{\beta\prime}) + (c_1^{\gamma\prime} c_3^{\gamma\prime})\Big) \end{aligned} \tag{7}$$

Summing over the shifts $\tau$ and rearranging the terms, we get:

$$\boxed{9z = (c_1^{\alpha\prime} c_2^{\alpha\prime} + c_1^{\beta\prime} c_2^{\beta\prime} + c_1^{\gamma\prime} c_2^{\gamma\prime}) + (c_1^{\alpha\prime} c_3^{\alpha\prime} + c_1^{\beta\prime} c_3^{\beta\prime} + c_1^{\gamma\prime} c_3^{\gamma\prime}) + (c_2^{\alpha\prime} c_3^{\alpha\prime} + c_2^{\beta\prime} c_3^{\beta\prime} + c_2^{\gamma\prime} c_3^{\gamma\prime})} \tag{8}$$

This shows that, within the limits of resolution of the mesh grid, the FFT (Eq. 8) converges to the Naive implementation (Eq. 5). This identity is valid if and only if the SNPs are equally spaced. If not, then Eq. 8 is an approximation of Eq. 5. Note, Eq. 8 is asymptotically equal to Eq. 5 if $m$ tends towards infinity. When the allele sharing value is missing for a particular mesh point, we assign to it the value of 0 so that the convolution is equal to 0 and has no impact on the final summing of convolution coefficients for a given shift $\tau$. However, when taking the average of the convolution coefficients for all $\tau$ within a particular genetic distance bin $d$, we make sure to not include these missing mesh points at the denominator so that the average autocorrelation of allele sharing is not biased.

Our analyses show that the FFT implementation can lead to runtime speed up by a hundred-fold, allowing the analysis of population genome-wide data in a few minutes, rather than hours using the Naive approach. We note that the smaller the mesh, the lower the approximation (but the higher the runtime and memory usage) and the closer the convergence between the Naive and FFT implementations. Empirically, we find that using a resolution of $10^{-3}$ cM leads to nearly equal allele sharing correlation values between the FFT and Naive implementations.

#### S1.3 Assessment of the fitted decay curves

The allele sharing correlation decay curves in empirical data can be noisy, either because of technical issues (such as dataset size) or population history (for example, there is a high variance in coalescent rates in case of long, weak bottlenecks). This can impede reliable inference of the founder event parameters as it is challenging to reliably fit an exponential model in noisy data (Bromage 1983; Groen et al. 1987).

In order to systematically assess if we can make reliable inference in real data, we calculated the Normalized Root-Mean-Square Deviation (NRMSD) between the empirical and fitted decay curves for each population (Ramasubramanian and Singh 2017). Specifically, we calculated the NRMSD as follows:

$$NRMSD = \frac{1}{max(\hat{z}) - min(\hat{z})} \sqrt{\frac{\sum^{D} (z_o - \hat{z})^2}{D}} \tag{9}$$

where $z_o$ and $\hat{z}$ are the empirical allele sharing correlation and the fitted allele sharing correlation values for any particular genetic distance ($d$) respectively. $D$ is the number of genetic distance bins.

We calculated NRMSD for each population of the present-day HO37 human populations and the distribution of these values is shown in Figure S1.3.1. Based on visual inspection, we suggest that the value of NRMSD=0.29 at the right tail of the NRMSD distribution is an appropriate threshold above which the empirical decay curves become too noisy to be reliably interpreted (Figure S1.3.2).

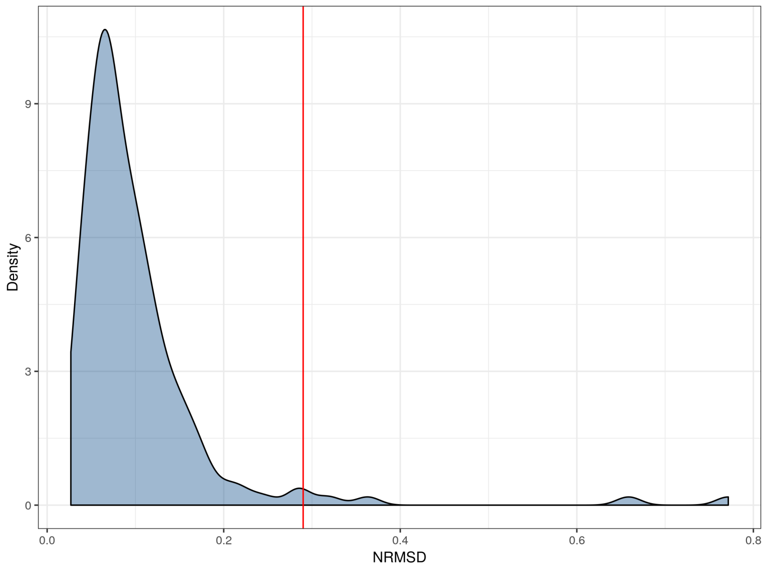

Figure S1.3.1: Density of the NRMSD computed as the normalized residual between the empirical and fitted decay curves, for all the present-day HO37 human populations. To decide the threshold, we focused on present-day human populations as they have a better data quality than ancient DNA samples. The red vertical line represents the value NRMSD=0.29, which we used as the threshold to exclude populations from our analysis, because inspection of fitted curves above this threshold suggest the results are too noisy to make reliable inference (see Figure S1.3.2 below).

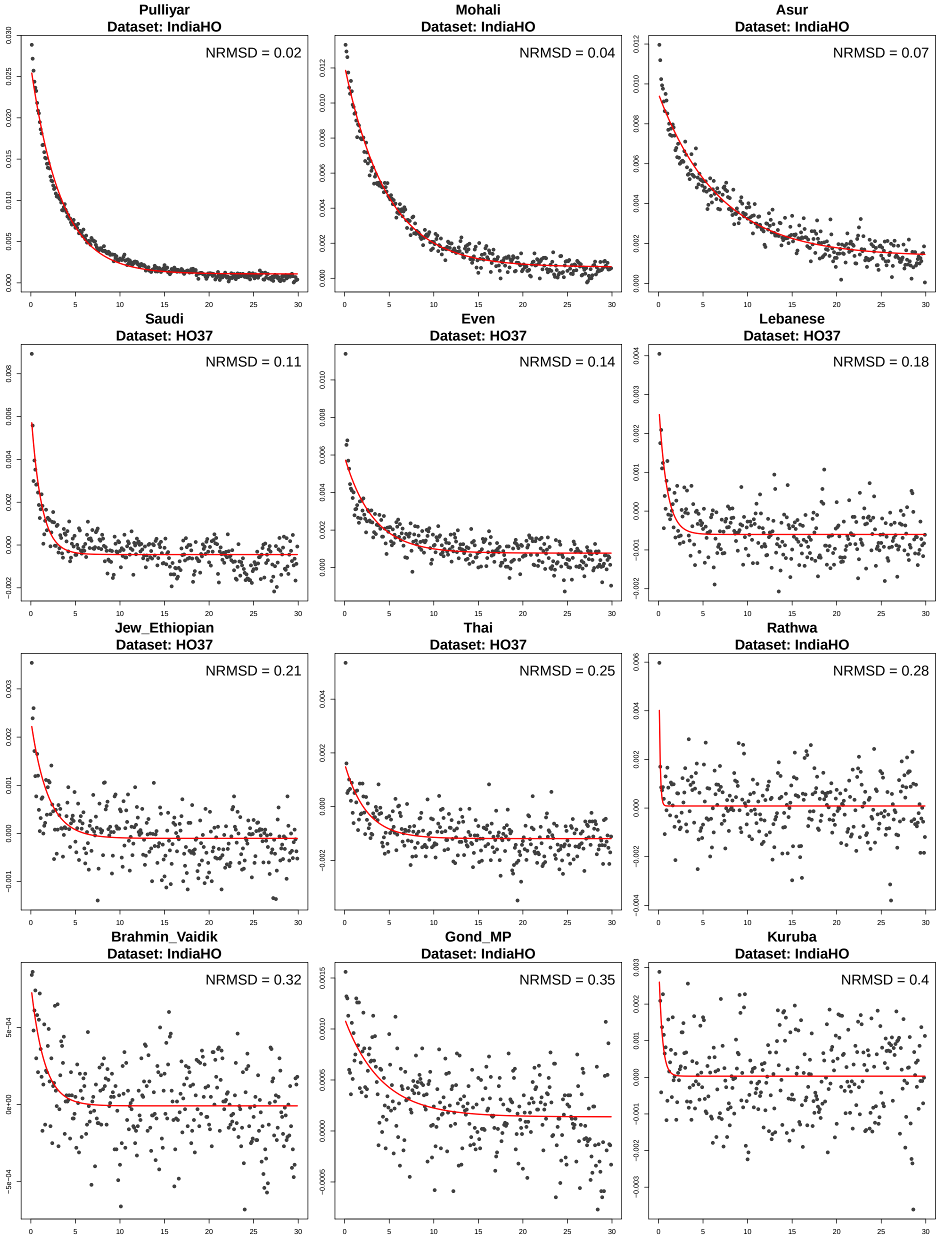

Figure S1.3.2: Allele sharing correlation curves for selected populations with NRMSD values ranging from NRMSD=0.02 to 0.40. Based on visual inspection, we exclude populations with an NRMSD>0.29. The three plots on the last row have a NRMSD above this threshold and hence were excluded from further analysis.

### S2. Simulations to test the performance of *ASCEND*

In order to characterize the accuracy of the method in estimating the age and intensity of founder events, we simulated datasets using *msprime* (Kelleher et al. 2016) under a variety of demographic models.

For all simulations, we generated data for two populations (target $A$, outgroup $O$) that diverged 1,800 generations ago (similar to Africans and Europeans) with effective population sizes of $N_o$=12,500 (Fig. S2.1.1). Unless stated otherwise, we simulated 20 chromosomes of size 50 megabases each for sample size of $n = 30$ haploid individuals in each population, assuming a mutation rate of $1.2 \times 10^{-8}$ per base pair per generation (Jonsson et al. 2017) and a recombination rate of $1.0 \times 10^{-8}$ per base pair per generation (Halldorsson et al. 2019). We assumed a uniform recombination rate across the chromosomes. We combined two haploid chromosomes at random to generate one diploid chromosome.

For the inference, unless stated otherwise, we computed $z_w(d)$ as the correlation between pairs of individuals in the target population $A$, and then subtracted the cross-population correlation $z_c(d)$ estimated across pairs of individuals in $(A, O)$ to remove the effects of ancestral allele sharing (Fig. S2.1.1). We computed the correlation in windows of genetic distance of 0.1 cM, starting from 0.1 cM to 30 cM. For the FFT implementation, we set the number of sub-bins to 100.

#### S2.1 Single-generation epoch model

In the single generation epoch model, the target population $A$ experienced a severe bottleneck happening $T_f$ generations ago (ranging between 10 to 300 generations ago), such that the population size reduced to $N_f$=5 for a single generation. After $T_f$, the population recovered to the original population size of $N_o$ (Figure S2.1.1).

Applying *ASCEND* to this simulated dataset, we accurately recovered the age and intensity of the founder events that occurred up to 200 generations ago (Figure S2.1.2). We observed a slight underestimation of the dates beyond 200 generations, largely because the exponential fitting starts at 0.1 cM (to guard against fine-scale errors in the recombination map in real data (Sankaraman et al. 2012)) which can lead to lower resolution for older dates.

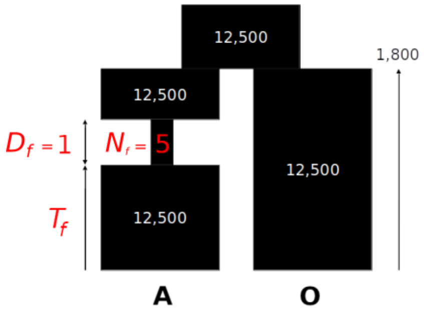

**Figure S2.1.1** - Demographic model used for single-generation epoch model. In the figure, we show two simulated populations $(A, O)$ that diverged 1,800 generations ago. The target population $A$ experienced a severe bottleneck $T_f$ generations ago, such that the population size reduced to $N_f$=5 for the duration $D_f$ of a single generation.

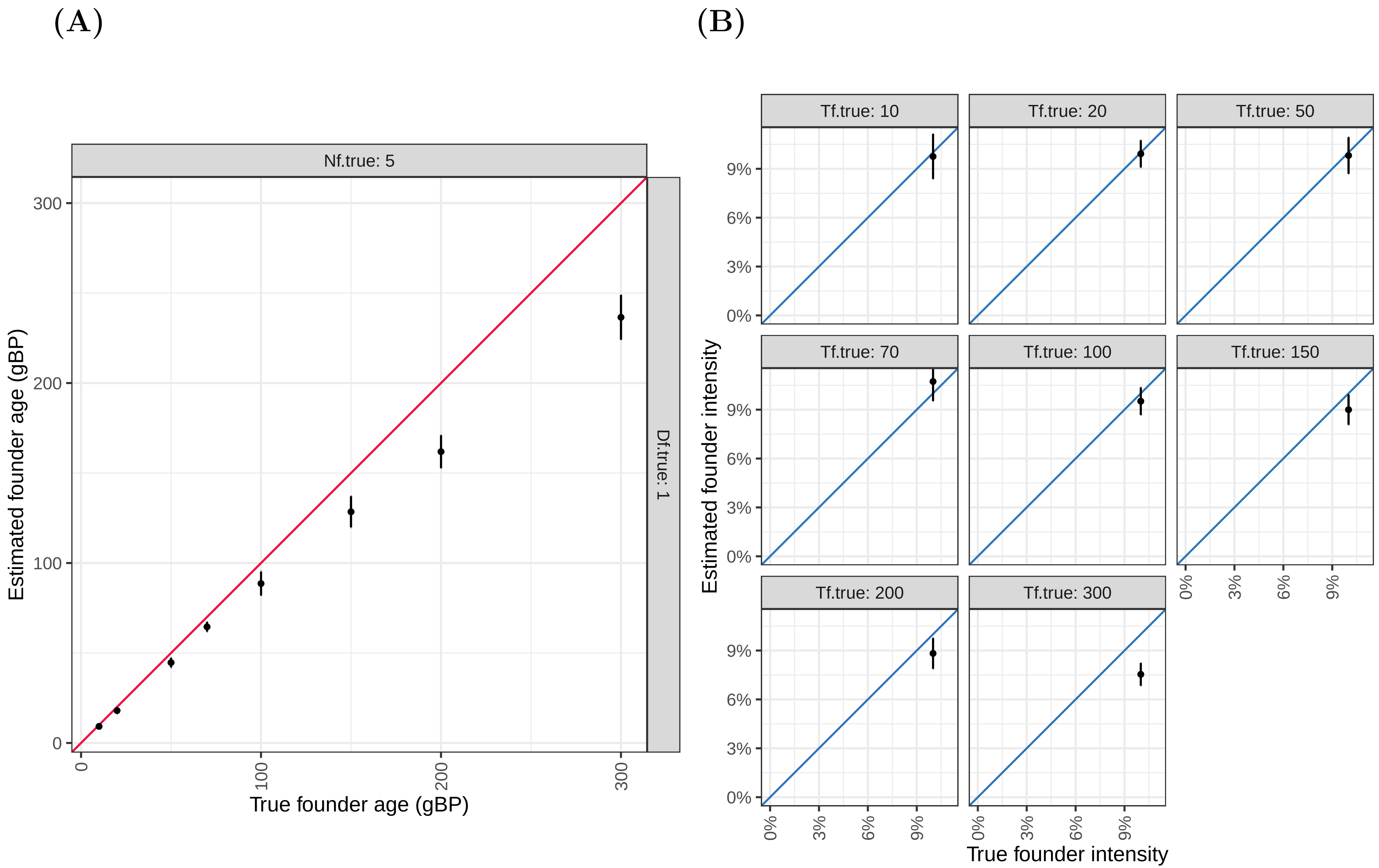

**Figure S2.1.2** Performance of *ASCEND* for the single-generation epoch model: **(A)** Founder age $T_f$ and **(B)** Founder intensity $I_f$.
The x-axis shows the true simulated parameter values and the y-axis shows the parameter values estimated by *ASCEND*.
The diagonal represents the expectation under the model. $N_f$ refers to the population size during the bottleneck and $D_f$ to the duration of the bottleneck.

#### S2.2 Multi-generation epoch model

We simulated data for a population $A$ that experienced a bottleneck for a duration $D_f$ ranging from 10 to 30 generations. During the bottleneck (Figure S2.2.1) period, the population size reduced to $N_f$, ranging from 200 to 1,000.

Applying *ASCEND* to this dataset, we observed that the estimated age and intensity of the founder event was accurate up to 200 generations ago, even in case of less intense founder events where the population size was $N_f = 1,000$ (Figure S2.2.2). However, we note that beyond 200 generations, *ASCEND* underestimated the age of the founder event. The estimated intensity appeared to be relatively unbiased for all ages.

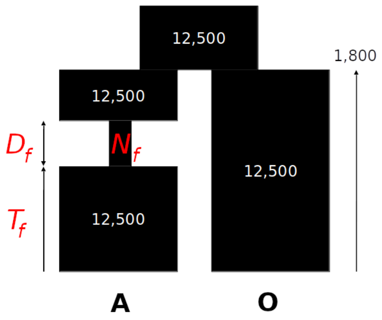

**Figure S2.2.1** - Demographic model for multi-generation epoch model. In the figure, we show two simulated populations $(A, O)$ that diverged 1,800 generations ago. The target population $A$ experienced a severe bottleneck $T_f$ generations ago, such that the population size reduced to $N_f$ for a duration of $D_f$ generations.

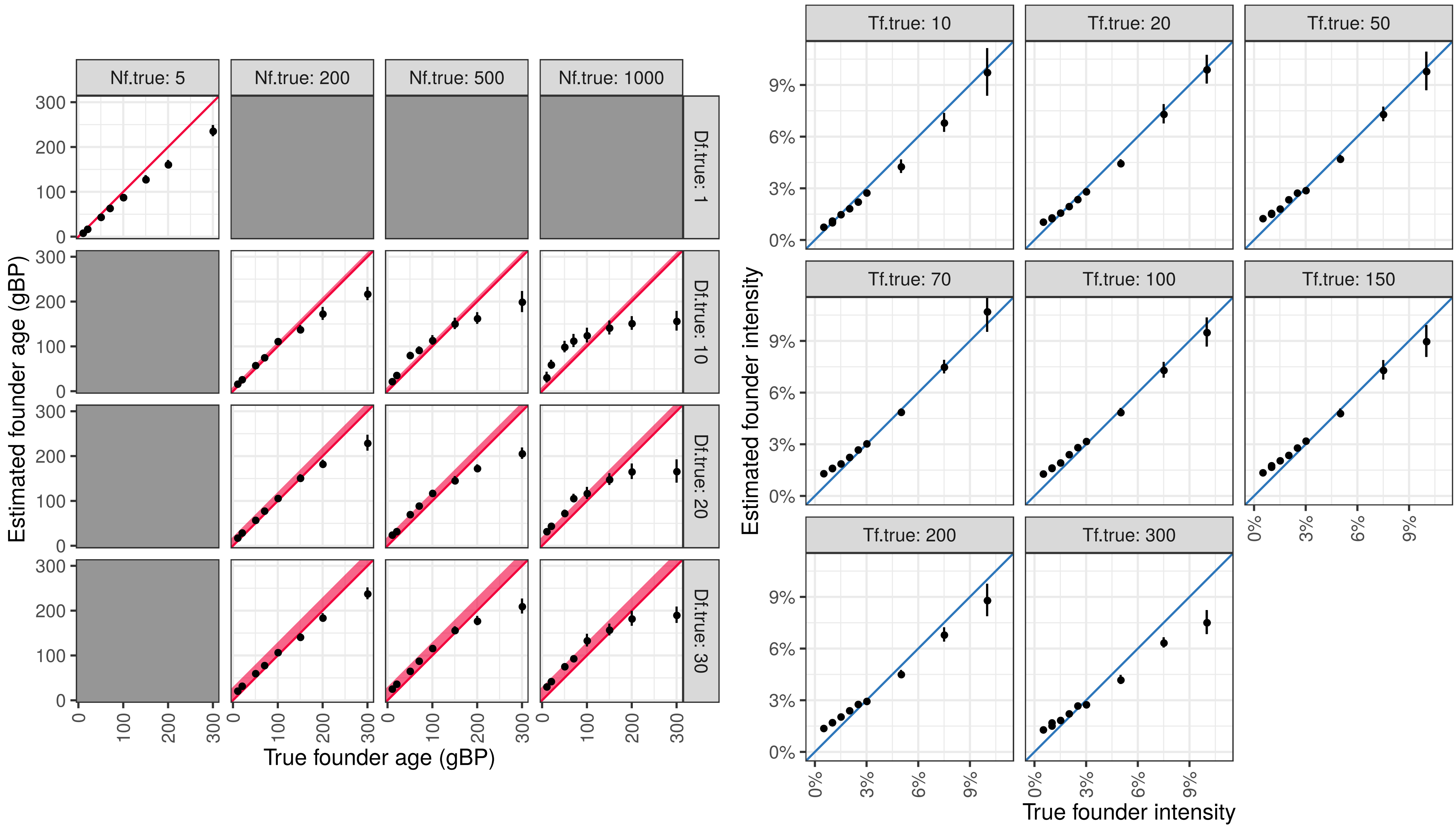

**Figure S2.2.2** - Performance of *ASCEND* for the multi-generation epoch model: **(A)** Founder age $T_f$ and **(B)** Founder intensity $I_f$.
The x-axis shows the true simulated parameter values and the y-axis shows the parameter values estimated by *ASCEND*.
The diagonal represents the expectation under the model. $N_f$ refers to the population size during the bottleneck and $D_f$ refers to the duration of the bottleneck.
Grey boxes indicate cases where no data were simulated.

#### S2.3 Two-epoch bottleneck model

In real data, population history can be more complex, involving multiple bottlenecks separated by long periods of time. To study the behavior of *ASCEND* under this setup, we generated data for a target population $A$ that experienced two founder events, one $T_f + \Delta T$ generations ago (with $T_f$ ranging between 10 and 200 generations) where population size reduced to $N_f = 5$ during a single generation, then the population recovered to its original size $N_o$. This was followed by a second bottleneck $T_f$ generations ago where the population size reduced to $N_f$ (ranging from 5 to 500) before recovering to $N_o$ again (Figure S2.3.1). The parameter $\Delta T$ (noted as the $TimeBetweenFEs$ in the Figure S2.3.2) represents the time lapsed between the two bottlenecks.

Figure S2.3.2 shows that we reliably recovered the intensity of the strongest founder event (here, the oldest one with $I_f = 10\%$). For severe bottlenecks ($N_f = 5$), we reliably recovered the age of the most recent founder event, but for less severe bottlenecks ($N_f \geq 200$), the estimated age is roughly the weighted average of the two founder ages, weighted by their respective intensities.

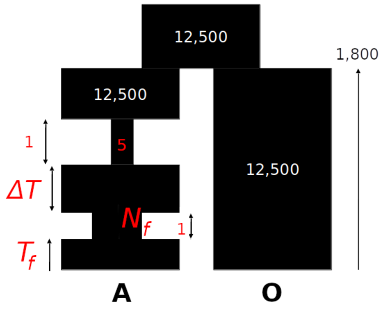

**Figure S2.3.1** - Demographic model for the two-epoch bottleneck model. The target population $A$ experienced two founder events, one $T_f + \Delta T$ generations ago where population size reduced to $N_f = 5$ for one generation, then the population recovered to its original size $N_o$. This was followed by a second bottleneck $T_f$ generations ago where the population size reduced to $N_f$ for a single generation, before recovering to $N_o$ again. The parameter $\Delta T$ represents the time lapsed between the two bottlenecks.

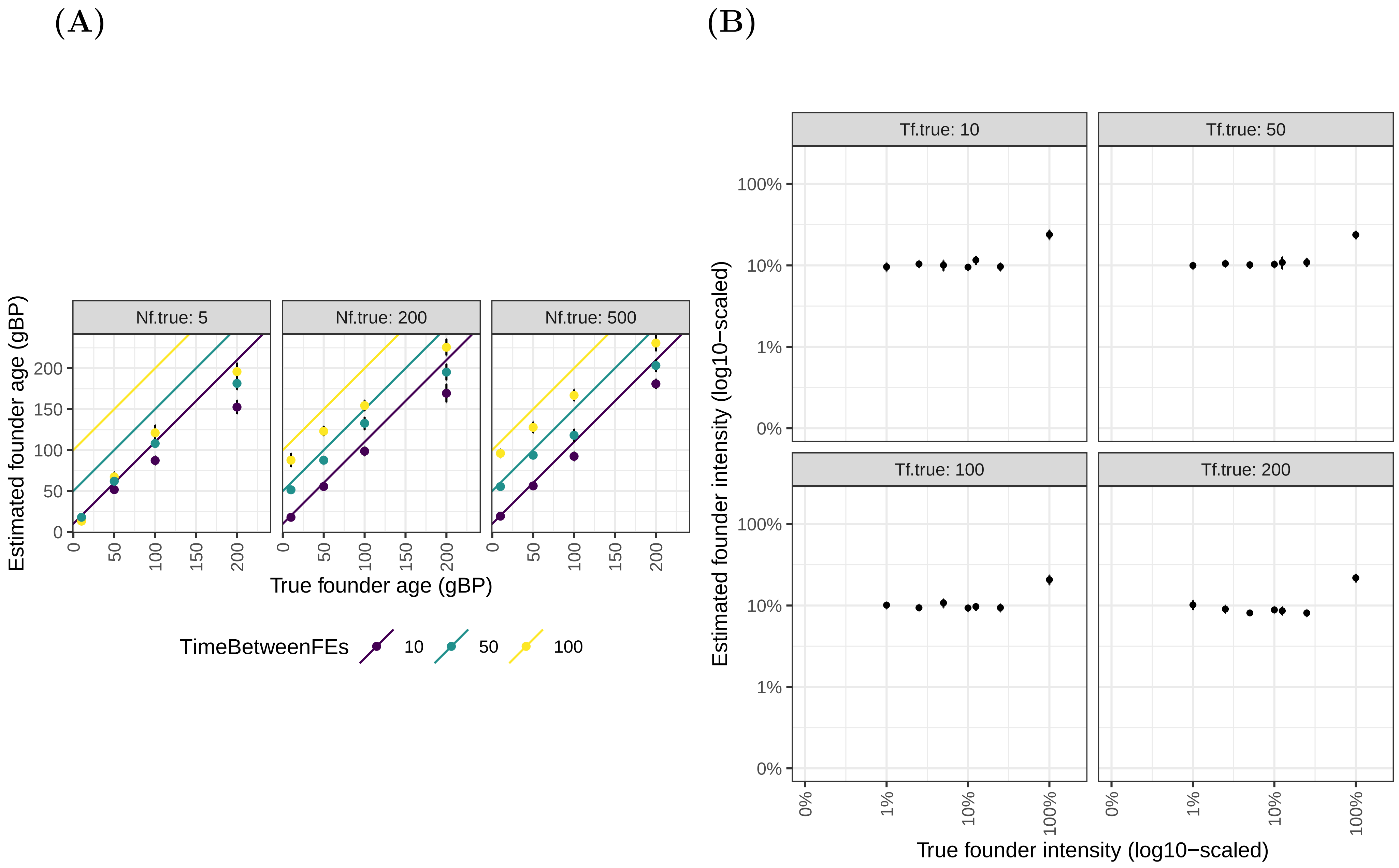

**Figure S2.3.2** - Performance of *ASCEND* for the two-epoch bottleneck model: **(A)** Founder age $T_f$ and **(B)** Founder intensity $I_f$.
The x-axis shows the true simulated parameter values and the y-axis shows the parameter values estimated by *ASCEND*.
The red line indicates the timing of most recent founder event. $N_f$ is the size of the last founder event.
The diagonal lines (in purple, green and yellow) represent the age of the oldest founder event (as a function of the time elapsed between the two events, *TimeBetweenFEs*).

#### S2.4 Model with founder event and admixture

Like a founder event, admixture introduces long-range allele correlation across the genome in the target population. Thus, under the scenario where a population has experienced both admixture and founder event, it is possible that admixture can confound the inference of the founder event. To explore the effect of admixture on *ASCEND*, we simulated a target population $A$ that derived ancestry from two ancestral populations $A'$ and $B'$, with ancestry proportions of 60% and 40% respectively. The two ancestral populations diverged 1,800 generations ago and the admixture occurred 110 generations ago. The target population $A$ then experienced a severe bottleneck $T_f$ generations ago (ranging between 10 to 100 generations) where the population size reduced to $N_f = 5$ (Figure S2.4.1).

Applying *ASCEND* to the target population and using one of the ancestral groups as the outgroup (to compute cross-population allele sharing), we observed that the admixture had no impact on the inference of the parameters (age, intensity) of the founder event (Figure S2.4.2).

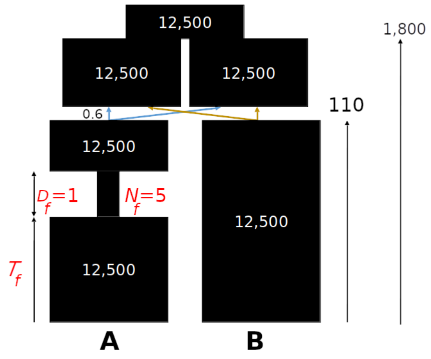

**Figure S2.4.1** - Demographic model for admixture and founder event scenario. The target population $A$ derived ancestry from two ancestral populations $A'$ and $B'$ (that are ancestral to present-day groups, $A$ and $B$), with ancestry proportions of 60% and 40% respectively. The two ancestral populations diverged 1,800 generations ago and the admixture occurred 110 generations ago. The target population $A$ then experienced a severe bottleneck $T_f$ generations ago where the population size reduced to $N_f = 5$ for a duration $D_f$ of single generation.

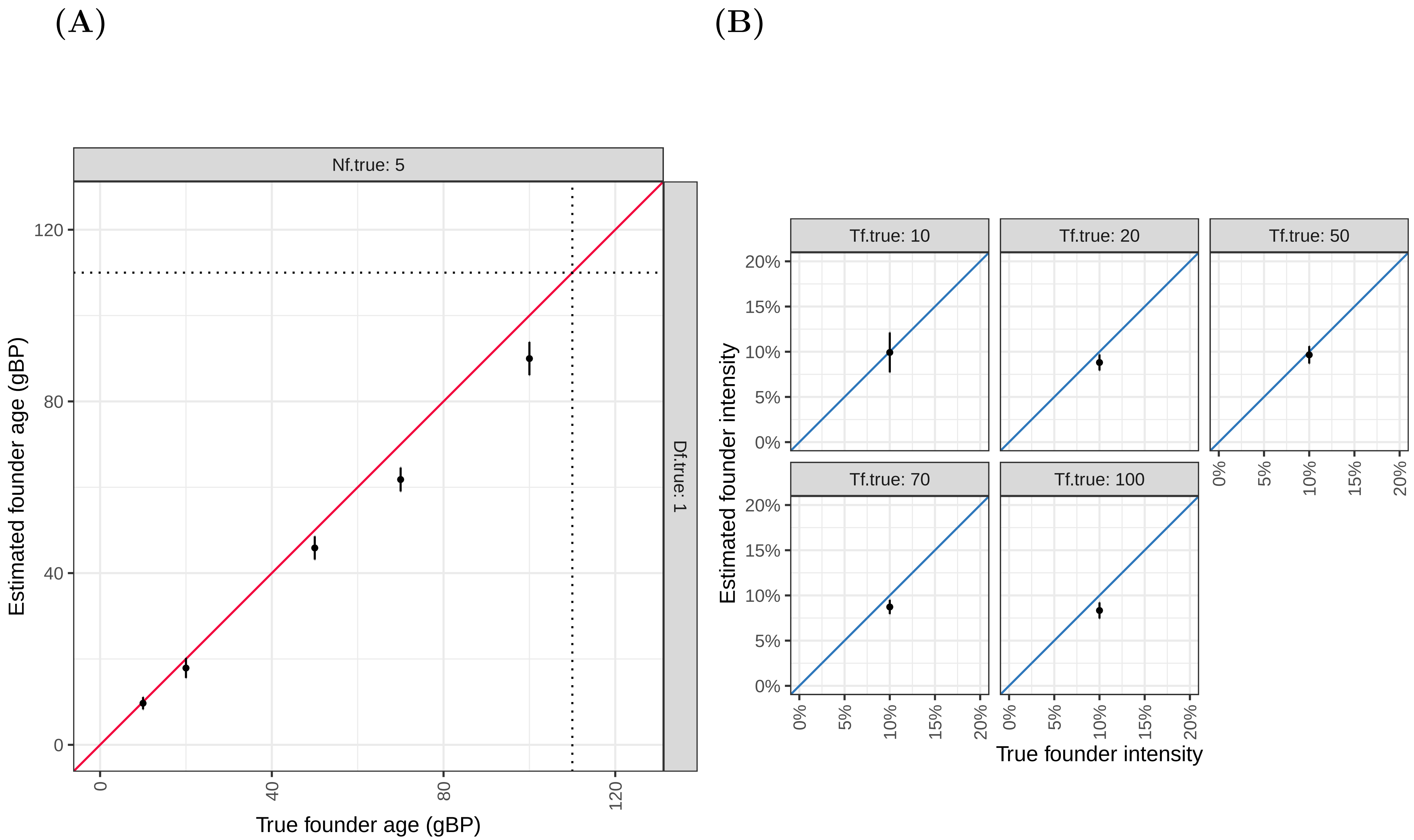

**Figure S2.4.2** - Performance of *ASCEND* for the model with admixture and founder event: **(A)** Founder age $T_f$ and **(B)** Founder intensity $I_f$.
The x-axis shows the true simulated parameter values and the y-axis shows the parameter values estimated by *ASCEND*.
The diagonal represents the expectation under the model. $N_f$ refers to the population size during the bottleneck and $D_f$ refers to the duration of the bottleneck.
The dotted black line in (A) shows the time of admixture (110 generations).

#### S2.5 Gradual exponential growth model

In the gradual exponential growth model, we generated data for a target population $A$ that experienced a founder event $T_f$ generations ago (from 10 to 300 generations) where the population size reduced to $N_f$ (from 5 to 1,000). This population then exponentially recovered with a rate $\lambda$ to reach $N_o = 12,500$ at present (Figure 2.5.1). The population size at time $t$ after the founder event is thus equal to $N_f \cdot e^{\lambda t}$.

Applying *ASCEND* under the standard setup, which assumes the epoch model of founder event, we observed that *ASCEND* tended to underestimate the founder age and to overestimate the founder intensity. However, we note that under the gradual epoch model, our parameters of age and intensity are ill-defined as our two parameter model does not capture important summary statistics such as rate of exponential increase in population size. Our inferred parameters can be assumed as the harmonic mean of the age and intensity over the duration the bottleneck (Figure 2.5.2B). By leveraging additional moments (such as variance) of the two-point allele sharing statistics, we may be able to infer the rate of recovery of the bottleneck and improve the reliability of the parameter estimation.

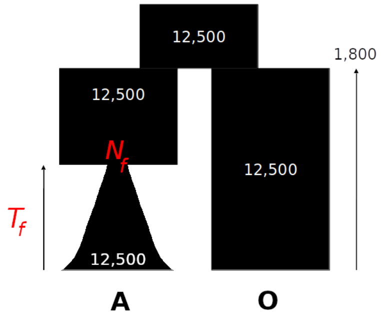

**Figure S2.5.1** - Demographic model for gradual exponential growth model. The target population $A$ experienced a founder event $T_f$ generations ago where the population size reduced to $N_f$. This population then exponentially recovered with a rate $\lambda$ to reach $N_o = 12,500$ at present.

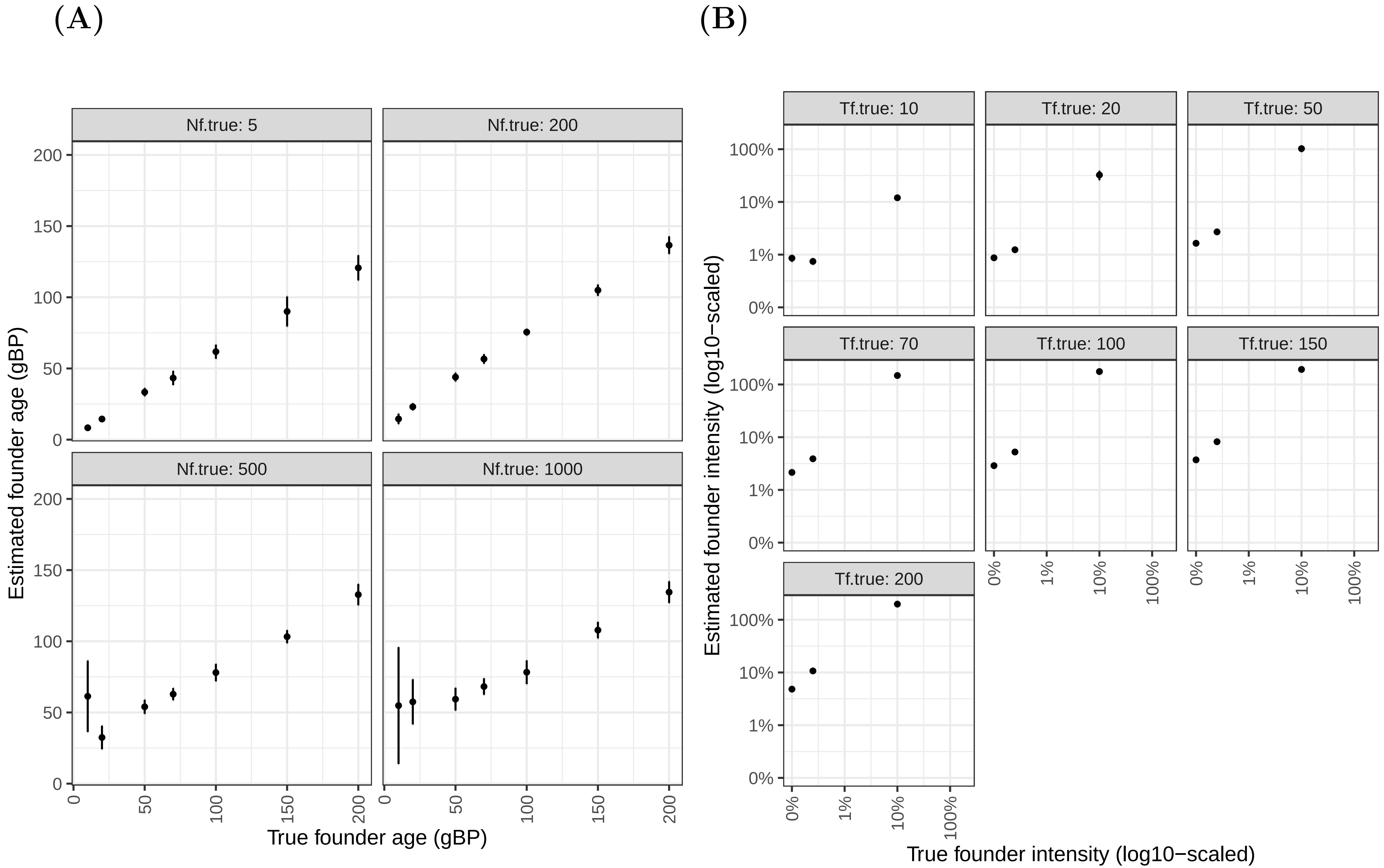

**Figure S2.5.2** - Performance of *ASCEND* for the gradual exponential growth model: **(A)** Founder age $T_f$ and **(B)** Founder intensity $I_f$.
The x-axis shows the true simulated parameter values and the y-axis shows the parameter values estimated by *ASCEND*.
$N_f$ refers to the population size during the bottleneck.

#### S2.6 No recovery founder event model

In this model, we assumed that the target population $A$ experienced a founder event $T_f$ generations ago (from 10 to 200 generations) where population size reduced to $N_f$ (from 5 to 1,000). Unlike the previous simulations, the target population $A$ did not recover after the bottleneck and maintained a low effective population size of $N_f$ till present (Figure 2.6.1).

Applying *ASCEND*, we inferred that the founder age was systematically underestimated. Defining the true founder intensity $I_f$ as $(T_f)/(2N_f)$ since in this model, the bottleneck period extends over $T_f$ generations to present, we found that $I_f$ was reliably estimated (Figure S2.6.2B).

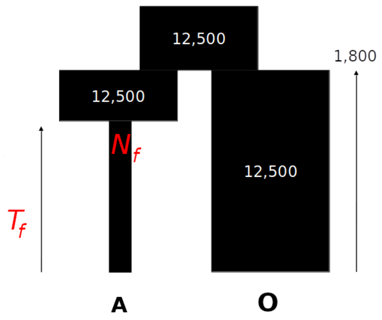

**Figure S2.6.1** - Demographic model for no recovery founder event model. The target population $A$ experienced a founder event $T_f$ generations ago where population size reduced to $N_f$. This historically low population size of $N_f$ is maintained to present.

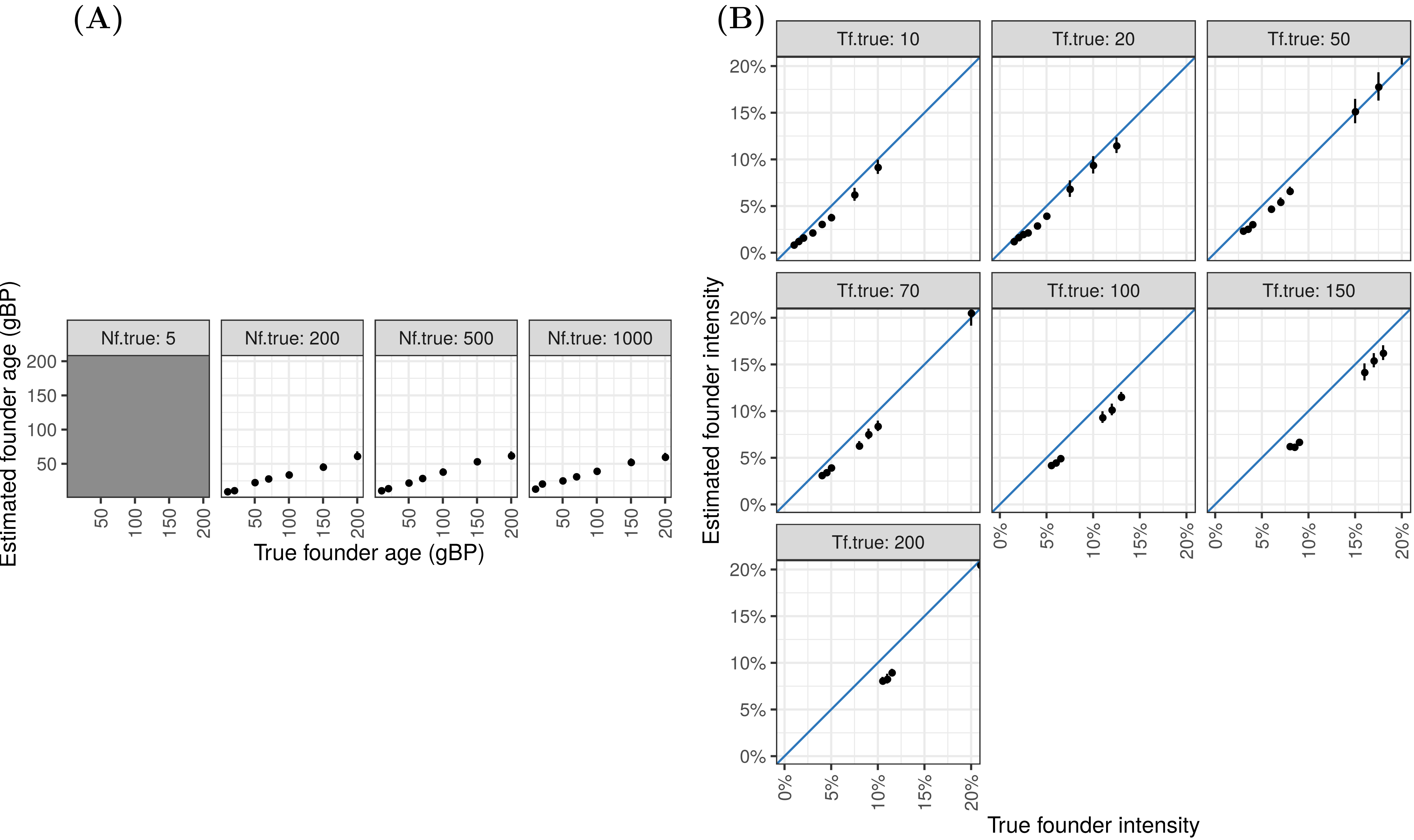

**Figure S2.6.2** - Performance of *ASCEND* for the no recovery founder event model: **(A)** Founder age $T_f$ and **(B)** Founder intensity $I_f$.
The x-axis shows the true simulated parameter values and the y-axis shows the parameter values estimated by *ASCEND*.
The diagonal represents the expectation under the model. $N_f$ refers to the population size during the bottleneck and $I_f$ is equal to $(T_f)/(2N_f)$.

#### S2.7 Robustness of the inference to data quality issues

##### S2.7.1 Impact of sample size

To investigate the impact of sample size of the target population on the inference of the founder event parameters, we simulated data for the single-generation epoch model (described in S2.1, Figure S2.1.1). We varied the sample size ($n$) of the target population between $5 - 30$ diploid individuals. The sample size of the outgroup population $O$ remained unchanged ($n = 15$) as generally there is less constraint for data from reference populations.

We observed that we could reliably estimate the age of the founder event in the target population for all sample sizes. We note however that the estimation of founder intensity can be slightly overestimated for very strong founder events ($N_f = 5$) in case of low sample sizes ($n = 5$) (Figure S2.7.1.2). Results were unbiased for larger samples sizes ($n > 10$).

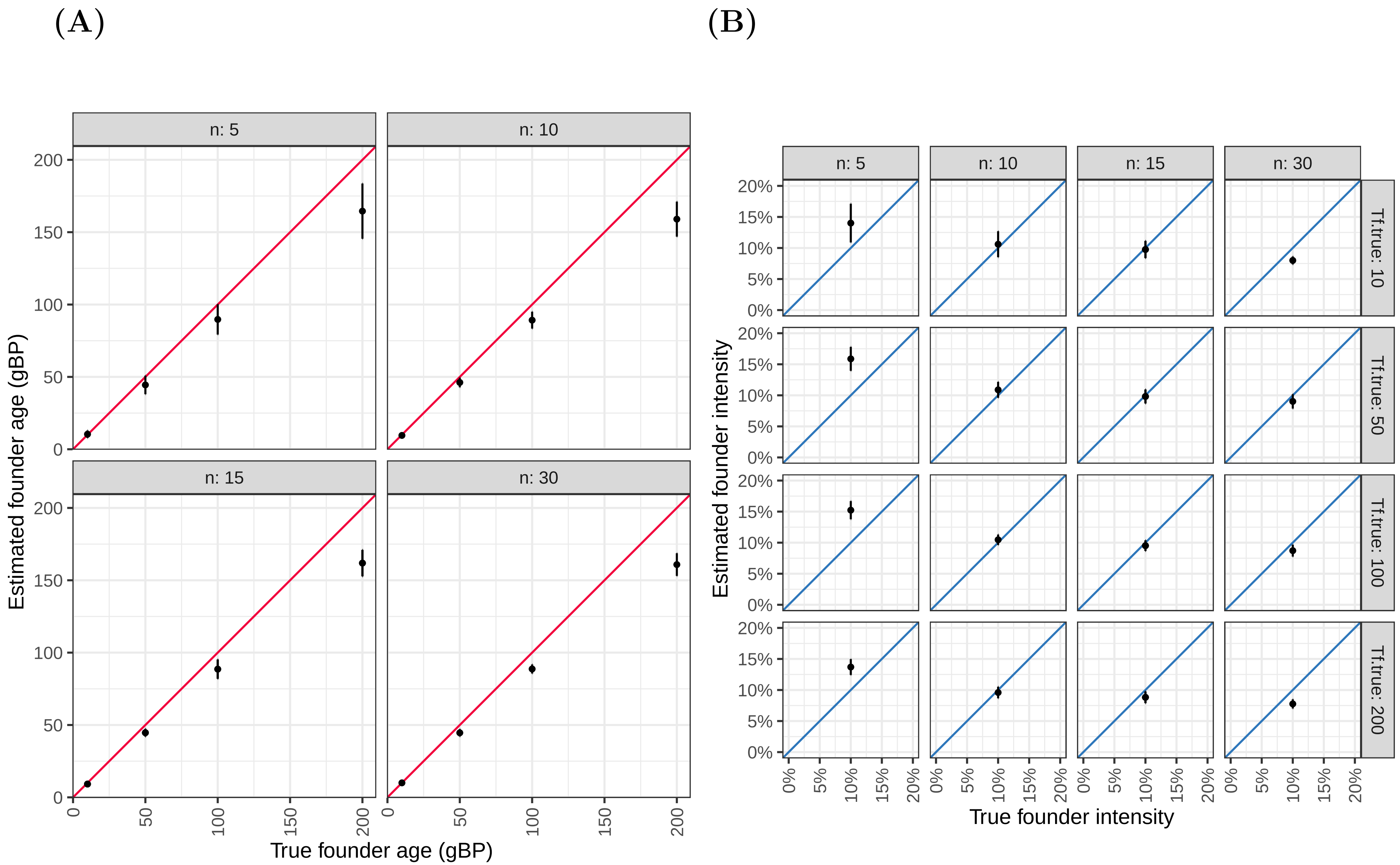

**Figure S2.7.1.2** - Impact of sample size on the inference of founder parameters: **(A)** Founder age $T_f$ and **(B)** Founder intensity $I_f$.
The x-axis shows the true simulated parameter values and the y-axis shows the parameter values estimated by *ASCEND*.
The diagonal represents the expectation under the model. $n$ refers to the sample size of the target population.

##### S2.7.2 Impact of missing data

To investigate the impact of missing genotypes in the target and outgroup population on the inference of the founder event parameters, we simulated data for the single-generation epoch model (described in S2.1, Figure S2.1.1) and set a proportion ($\theta$) of genotypes to missing, with a range of *theta* ranging from 20% - 90%.

We observed that the inference was robust to large proportion of missing data ($< 90\%$) and both age and intensity were accurately inferred (Figure S2.7.2.2). With greater that 90% missing genotypes, the inference is unstable and founder intensity can be overestimated (Figure S2.7.2.2).

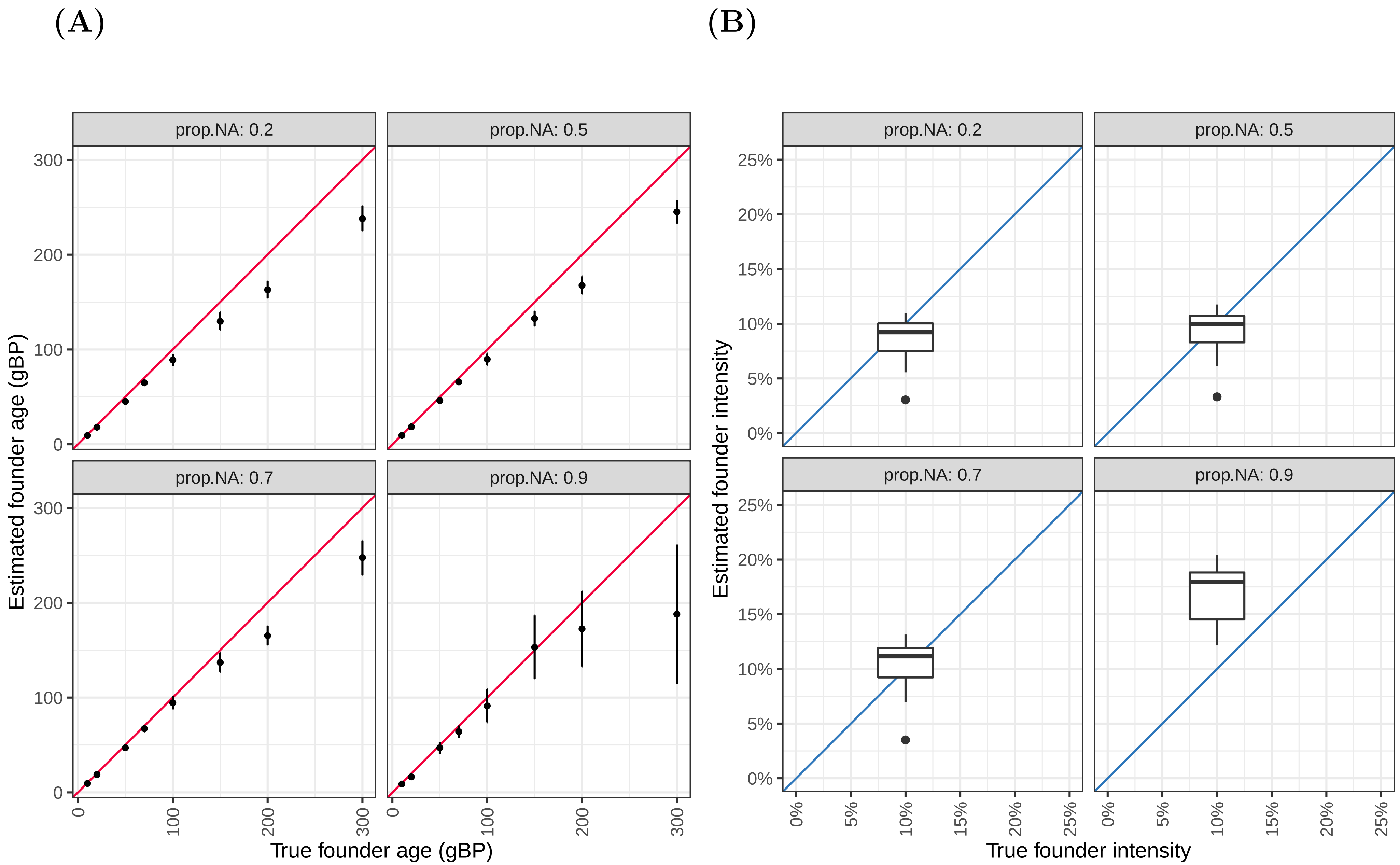

**Figure S2.7.2.2** - Impact of missing data on the inference of founder parameters: **(A)** Founder age $T_f$ and **(B)** Founder intensity $I_f$.
The x-axis shows the true simulated parameter values and the y-axis shows the parameter values estimated by *ASCEND*.
The diagonal represents the expectation under the model. *prop.NA* refers to the proportion of missing data.

##### S2.7.3 Impact of missing data: special case of ancient DNA

Ancient DNA specimens tend to have low quality of data due to poor preservation and high degradation of DNA with time. Thus, ancient DNA datasets often have high proportion of missing data and limited or low coverage. To avoid bias, it is common practice to make pseudo-homozygous genotype calls using a random allele observed in the reads at each site in the genome (Harney et al. 2018).

To test the robustness of our inference to ancient DNA datasets, we simulated data for the single-generation epoch model (described in S2.1, Figure S2.1.1) and then resampled this dataset to mimic the features of ancient genomes, namely (i) sample size was set to 5 samples in the target population as often only few ancient genomes are available; (ii) number of SNPs was set to 500,000; (iii) out of the 500,000 SNPs, a large proportion were set to missing with the rate ranging from 20% to 90%; (iv) for all sites, we used pseudo-homozygous genotype calls (i.e., for every heterozygous site, we randomly assigned one of the alleles as the homozygous genotype at that site). As described in the main text, it is difficult to find a reliable outgroup for ancient DNA samples (as it depends on matching samples based on timescale and location and currently is infeasible due to the limited number of ancient genomes available), and hence we based the inference on the within-population correlation ($z_w$) in the target population only, without subtracting the cross-population correlation ($z_c$).

Our simulations showed that ancient DNA features have minimal impact on the inference and we can reliably estimate the age and intensity of the founder event even with pseudo-homozygous genotype calls and with large amounts of missing data (Figure S2.7.3.2). These results highlight a major strength of our method, in that it works reliably even with limited data and hence is applicable to sparse datasets and to ancient genomes.

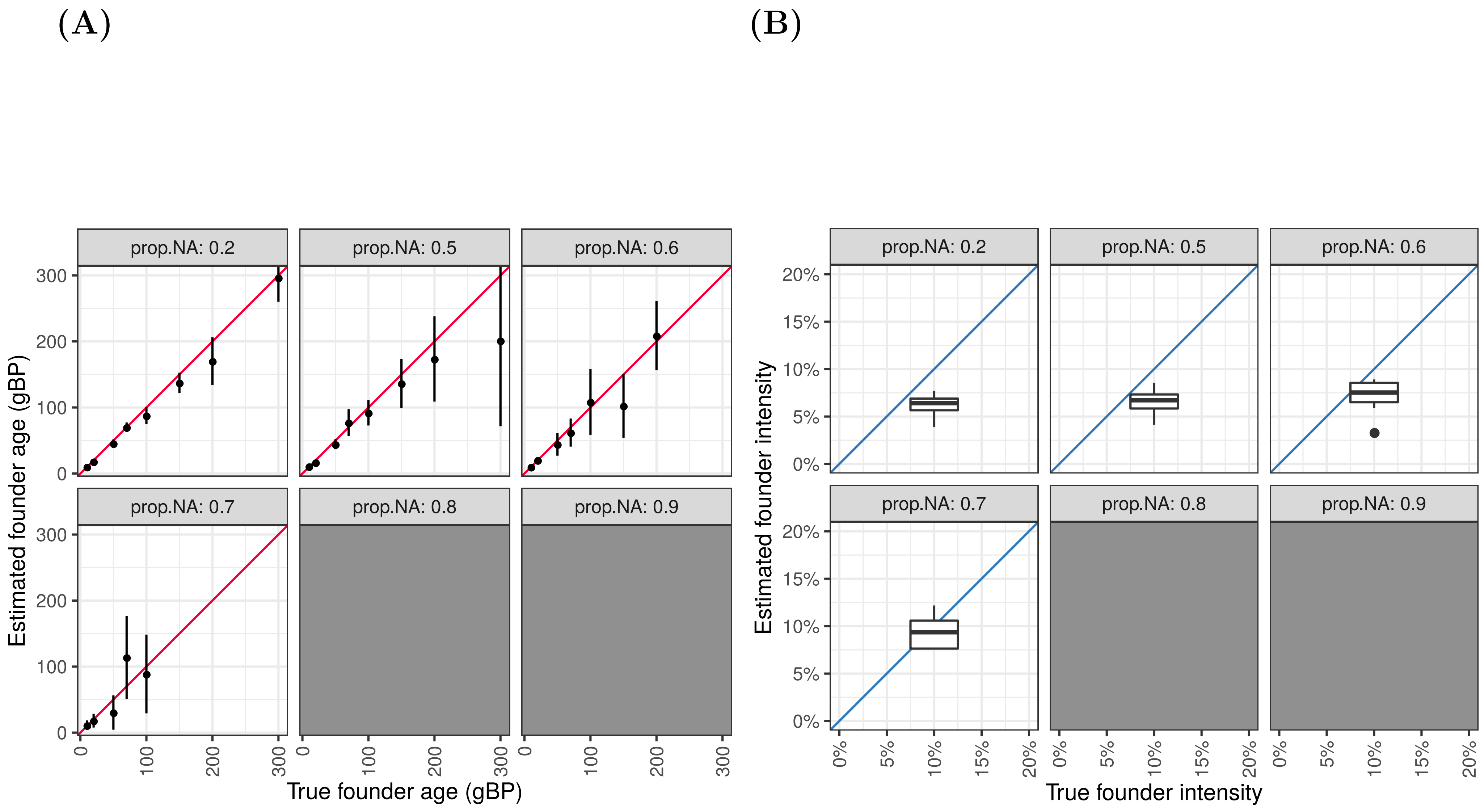

**Figure S2.7.3.2** - Impact of features of ancient DNA on the inference of founder parameters: **(A)** Founder age $T_f$ and **(B)** Founder intensity $I_f$
The x-axis shows the true simulated parameter values and the y-axis shows the parameter values estimated by *ASCEND*. The diagonal shows the expectation under the model. *prop.NA* refers to the proportion of missing data.
Grey boxes indicate cases where founder events failed to be detected by *ASCEND* due to high proportion of missing data.

#### S2.8. Comparison between Naive and FFT implementation

Using the data simulated under the S2.1 and S2.2 models, we estimated the founder parameters using both the Naive and the FFT approaches. The results are provided in Table S2.8.1 and show the nearly identical estimates provided by the two approaches. However, the speedup gain using FFT is of around a hundred-fold.

**Table S2.8.1** - Comparison between the estimates of the founder parameters (founder age $T_f$, founder intensity $I_f$) based on the Naive and the FFT approaches, using data simulated under the single-pulse and multi-generation epoch models.

| Founder age ($T_f$) (in generations) | | | $N_f$ | $D_f$ | Founder intensity ($I_f$) | | |
|---|---|---|---|---|---|---|---|
| True | FFT estimate (mean ± SE) | NAIVE estimate (mean ± SE) | True | True | True | FFT estimate (mean ± SE) | NAIVE estimate (mean ± SE) |
| 10 | 9 ± 1 | 9 ± 1 | 5 | 1 | 10% | 9.8% ± 0.7% | 9.8% ± 0.7% |
| 10 | 9 ± 1 | 9 ± 1 | 5 | 1 | 10% | 9.8% ± 0.7% | 9.8% ± 0.7% |
| 100 | 89 ± 3 | 89 ± 3 | 5 | 1 | 10% | 9.5% ± 0.4% | 9.5% ± 0.4% |
| 100 | 89 ± 3 | 89 ± 3 | 5 | 1 | 10% | 9.5% ± 0.4% | 9.5% ± 0.4% |
| 150 | 128 ± 4 | 128 ± 4 | 5 | 1 | 10% | 9% ± 0.5% | 9% ± 0.5% |
| 150 | 128 ± 4 | 128 ± 4 | 5 | 1 | 10% | 9% ± 0.5% | 9% ± 0.5% |
| 200 | 162 ± 5 | 163 ± 4 | 5 | 1 | 10% | 8.8% ± 0.5% | 8.8% ± 0.5% |
| 200 | 162 ± 5 | 163 ± 4 | 5 | 1 | 10% | 8.8% ± 0.5% | 8.8% ± 0.5% |
| 50 | 45 ± 1 | 45 ± 1 | 5 | 1 | 10% | 9.8% ± 0.6% | 9.8% ± 0.6% |
| 50 | 45 ± 1 | 45 ± 1 | 5 | 1 | 10% | 9.8% ± 0.6% | 9.8% ± 0.6% |
| 10 | 31 ± 6 | 32 ± 6 | 1000 | 10 | 0.5% | 0.8% ± 0% | 0.8% ± 0% |
| 10 | 33 ± 3 | 33 ± 2 | 1000 | 20 | 1% | 1.1% ± 0% | 1.2% ± 0% |
| 10 | 31 ± 2 | 31 ± 2 | 1000 | 30 | 1.5% | 1.5% ± 0% | 1.5% ± 0% |
| 10 | 17 ± 1 | 17 ± 1 | 200 | 10 | 2.5% | 2.2% ± 0.1% | 2.3% ± 0.1% |
| 10 | 19 ± 1 | 18 ± 1 | 200 | 20 | 5% | 4.3% ± 0.2% | 4.3% ± 0.2% |
| 10 | 22 ± 1 | 22 ± 1 | 200 | 30 | 7.5% | 6.8% ± 0.3% | 6.9% ± 0.3% |
| 10 | 23 ± 2 | 23 ± 2 | 500 | 10 | 1% | 1% ± 0% | 1.1% ± 0% |
| 10 | 25 ± 1 | 25 ± 1 | 500 | 20 | 2% | 1.9% ± 0% | 1.9% ± 0% |
| 10 | 27 ± 1 | 26 ± 1 | 500 | 30 | 3% | 2.8% ± 0.1% | 2.8% ± 0.1% |
| 100 | 125 ± 8 | 125 ± 8 | 1000 | 10 | 0.5% | 1.3% ± 0.1% | 1.3% ± 0.1% |
| 100 | 118 ± 7 | 115 ± 6 | 1000 | 20 | 1% | 1.7% ± 0% | 1.7% ± 0% |
| 100 | 134 ± 7 | 132 ± 7 | 1000 | 30 | 1.5% | 2% ± 0.1% | 2% ± 0.1% |
| 100 | 112 ± 3 | 112 ± 2 | 200 | 10 | 2.5% | 2.8% ± 0.1% | 2.9% ± 0.1% |
| 100 | 107 ± 2 | 107 ± 2 | 200 | 20 | 5% | 4.9% ± 0.1% | 4.9% ± 0.1% |
| 100 | 108 ± 2 | 108 ± 2 | 200 | 30 | 7.5% | 7.3% ± 0.2% | 7.4% ± 0.2% |
| 100 | 114 ± 5 | 116 ± 5 | 500 | 10 | 1% | 1.6% ± 0% | 1.7% ± 0% |
| 100 | 118 ± 3 | 119 ± 4 | 500 | 20 | 2% | 2.4% ± 0.1% | 2.5% ± 0.1% |
| 100 | 117 ± 3 | 116 ± 3 | 500 | 30 | 3% | 3.2% ± 0.1% | 3.2% ± 0.1% |
| 150 | 142 ± 8 | 142 ± 7 | 1000 | 10 | 0.5% | 1.4% ± 0% | 1.4% ± 0% |

| | | | | | | | |
|---|---|---|---|---|---|---|---|
| 150 | 149 ± 7 | 147 ± 6 | 1000 | 20 | 1% | 1.7% ± 0.1% | 1.7% ± 0.1% |
| 150 | 158 ± 7 | 156 ± 7 | 1000 | 30 | 1.5% | 2.1% ± 0% | 2.1% ± 0% |
| 150 | 138 ± 5 | 137 ± 5 | 200 | 10 | 2.5% | 2.8% ± 0.1% | 2.9% ± 0.1% |
| 150 | 152 ± 4 | 150 ± 4 | 200 | 20 | 5% | 4.8% ± 0.1% | 4.9% ± 0.1% |
| 150 | 142 ± 4 | 139 ± 4 | 200 | 30 | 7.5% | 7.3% ± 0.3% | 7.3% ± 0.3% |
| 150 | 151 ± 6 | 154 ± 7 | 500 | 10 | 1% | 1.8% ± 0.1% | 1.8% ± 0.1% |
| 150 | 146 ± 5 | 147 ± 6 | 500 | 20 | 2% | 2.4% ± 0.1% | 2.4% ± 0.1% |
| 150 | 157 ± 4 | 155 ± 4 | 500 | 30 | 3% | 3.2% ± 0.1% | 3.3% ± 0.1% |
| 200 | 152 ± 8 | 147 ± 8 | 1000 | 10 | 0.5% | 1.4% ± 0% | 1.4% ± 0% |
| 200 | 166 ± 9 | 167 ± 8 | 1000 | 20 | 1% | 1.7% ± 0% | 1.8% ± 0% |
| 200 | 183 ± 9 | 184 ± 9 | 1000 | 30 | 1.5% | 2.1% ± 0.1% | 2.1% ± 0.1% |
| 200 | 174 ± 7 | 171 ± 6 | 200 | 10 | 2.5% | 2.8% ± 0.1% | 2.8% ± 0.1% |
| 200 | 183 ± 4 | 180 ± 4 | 200 | 20 | 5% | 4.5% ± 0.1% | 4.5% ± 0.1% |
| 200 | 185 ± 5 | 184 ± 5 | 200 | 30 | 7.5% | 6.8% ± 0.2% | 6.8% ± 0.2% |
| 200 | 163 ± 7 | 165 ± 7 | 500 | 10 | 1% | 1.7% ± 0.1% | 1.8% ± 0.1% |
| 200 | 174 ± 4 | 170 ± 4 | 500 | 20 | 2% | 2.4% ± 0% | 2.4% ± 0% |
| 200 | 178 ± 5 | 177 ± 6 | 500 | 30 | 3% | 3% ± 0% | 3% ± 0.1% |
| 50 | 100 ± 7 | 97 ± 6 | 1000 | 10 | 0.5% | 1.3% ± 0% | 1.3% ± 0% |
| 50 | 73 ± 4 | 73 ± 4 | 1000 | 20 | 1% | 1.6% ± 0% | 1.6% ± 0% |
| 50 | 76 ± 3 | 77 ± 2 | 1000 | 30 | 1.5% | 1.8% ± 0% | 1.9% ± 0% |
| 50 | 59 ± 2 | 59 ± 2 | 200 | 10 | 2.5% | 2.8% ± 0.1% | 2.8% ± 0.1% |
| 50 | 58 ± 1 | 58 ± 2 | 200 | 20 | 5% | 4.7% ± 0.1% | 4.8% ± 0.1% |
| 50 | 61 ± 2 | 62 ± 2 | 200 | 30 | 7.5% | 7.3% ± 0.2% | 7.4% ± 0.2% |
| 50 | 81 ± 4 | 80 ± 3 | 500 | 10 | 1% | 1.5% ± 0% | 1.5% ± 0% |
| 50 | 71 ± 2 | 71 ± 2 | 500 | 20 | 2% | 2.4% ± 0.1% | 2.4% ± 0.1% |
| 50 | 66 ± 3 | 66 ± 2 | 500 | 30 | 3% | 2.9% ± 0.1% | 2.9% ± 0.1% |

Note: The column labeled "**True**" indicates the true value of the founder event parameters that were used in the simulations;
$N_f$ = population size during the founder event;
$D_f$ = duration of founder event.

### S2.9. *msprime* commands

#### S2.1 Single-generation epoch model

***$T_f$***: parameter set by the user (variable). Note that we simulate four populations, but only the first one and the last one are used (respectively the target and outgroup populations) for the analysis.

```
L = 50e6
recomb_rate = 1e-8
mut_rate = 1.2e-8
No = 12500
Nf = 5
Df = 1
```

```
pop_conf = [
    msprime.PopulationConfiguration(sample_size=30, initial_size = No),
    msprime.PopulationConfiguration(sample_size=30, initial_size = No),
    msprime.PopulationConfiguration(sample_size=30, initial_size = No),
    msprime.PopulationConfiguration(sample_size=30, initial_size = No)]

migr_matrix = [[0, 0, 0, 0],
               [0, 0, 0, 0],
               [0, 0, 0, 0],
               [0, 0, 0, 0]]

demo = [
       msprime.PopulationParametersChange(time = Tf, initial_size = Nf, population_id = 0),
       msprime.PopulationParametersChange(time = Tf+Df, initial_size = No, population_id =
0),
       msprime.MassMigration(time = 200, source = 1, destination = 0, proportion = 1),
       msprime.MassMigration(time = 1000, source = 2, destination = 0, proportion = 1),
       msprime.MassMigration(time = 1800, source = 3, destination = 0, proportion = 1)]

tree = msprime.simulate(population_configurations = pop_conf,
                            migration_matrix = migr_matrix,
                            length = L,
                            recombination_rate = recomb_rate,
                            mutation_rate = mut_rate,
                            demographic_events = demo)
```

#### S2.2 Multi-generation epoch model

**Tf**, **Nf**, **Df**: parameters set by the user (variable). Note that we simulate four populations, but only the first one and the last one are used (respectively the target and outgroup populations) for the analysis.

```
L = 50e6
recomb_rate = 1e-8
mut_rate = 1.2e-8
No = 12500

pop_conf = [
    msprime.PopulationConfiguration(sample_size=30, initial_size = No),
    msprime.PopulationConfiguration(sample_size=30, initial_size = No),
    msprime.PopulationConfiguration(sample_size=30, initial_size = No),
    msprime.PopulationConfiguration(sample_size=30, initial_size = No)]

migr_matrix = [[0, 0, 0, 0],
               [0, 0, 0, 0],
               [0, 0, 0, 0],
               [0, 0, 0, 0]]

demo = [
       msprime.PopulationParametersChange(time = Tf, initial_size = Nf, population_id = 0),
       msprime.PopulationParametersChange(time = Tf+Df, initial_size = No, population_id =
0),
       msprime.MassMigration(time = 200, source = 1, destination = 0, proportion = 1),
       msprime.MassMigration(time = 1000, source = 2, destination = 0, proportion = 1),
       msprime.MassMigration(time = 1800, source = 3, destination = 0, proportion = 1)]

tree = msprime.simulate(population_configurations = pop_conf,
                            migration_matrix = migr_matrix,
                            length = L,
                            recombination_rate = recomb_rate,
                            mutation_rate = mut_rate,
                            demographic_events = demo)
```

#### S2.3 Two-epoch bottleneck model

**Tf**, **Nf**, **Df**: parameters set by the user (variable).

```
L = 50e6
recomb_rate = 1e-8
mut_rate = 1.2e-8
No = 12500

pop_conf = [
    msprime.PopulationConfiguration(sample_size=30, initial_size = No),
    msprime.PopulationConfiguration(sample_size=30, initial_size = No)]

migr_matrix = [[0, 0], [0, 0]]

demo = [
    msprime.PopulationParametersChange(time = Tf, initial_size = Nf, population_id = 0),
    msprime.PopulationParametersChange(time = Tf+1, initial_size = No, population_id = 0),
    msprime.PopulationParametersChange(time = Tf+Df, initial_size = 5, population_id = 0),
    msprime.PopulationParametersChange(time = Tf+Df+1, initial_size = No, population_id =
0),
    msprime.MassMigration(time = 1800, source = 1, destination = 0, proportion = 1)]

tree = msprime.simulate(population_configurations = pop_conf,
                            migration_matrix = migr_matrix,
                            length = L,
                            recombination_rate = recomb_rate,
                            mutation_rate = mut_rate,
                            demographic_events = demo)
```

#### S2.4 Model with founder event and admixture

**Tf**, **Ta**: parameters set by the user (variable). Note that we simulate four populations, but only the first one and the last one are used (respectively the target and outgroup populations) for the analysis.

```
L = 50e6
recomb_rate = 1e-8
mut_rate = 1.2e-8
No = 12500
prop_admix = 0.4
Nf = 5
Df = 1

pop_conf = [
    msprime.PopulationConfiguration(sample_size=30, initial_size = No),
    msprime.PopulationConfiguration(sample_size=30, initial_size = No),
    msprime.PopulationConfiguration(sample_size=0, initial_size = No),
    msprime.PopulationConfiguration(sample_size=30, initial_size = No)]

migr_matrix = [[0, 0, 0, 0],
               [0, 0, 0, 0],
               [0, 0, 0, 0],
               [0, 0, 0, 0]]

demo = [
     msprime.PopulationParametersChange(time = Tf, initial_size = Nf, population_id = 0),
     msprime.PopulationParametersChange(time = Tf+Df, initial_size = No, population_id =
0),

     msprime.MassMigration(time = Ta, source = 0, destination = 2, proportion =
prop_admix),
     msprime.MassMigration(time = Ta, source = 1, destination = 2, proportion =
prop_admix),
```

```
     msprime.MassMigration(time = 1800, source = 0, destination = 3, proportion = 1),
     msprime.MassMigration(time = 1800, source = 1, destination = 3, proportion = 1),
     msprime.MassMigration(time = 1800, source = 2, destination = 3, proportion = 1)]

tree = msprime.simulate(population_configurations = pop_conf,
                            migration_matrix = migr_matrix,
                            length = L,
                            recombination_rate = recomb_rate,
                            mutation_rate = mut_rate,
                            demographic_events = demo)
```

#### S2.5 Gradual exponential growth model

**Tf**, **Nf**: parameters set by the user (variable).

```
L = 50e6
recomb_rate = 1e-8
mut_rate = 1.2e-8
No = 12500
Df = 1
alpha = 1/(Tf*1.) * np.log(No/(Nf*1.))

pop_conf = [
    msprime.PopulationConfiguration(sample_size=30, initial_size = No, growth_rate =
alpha),
    msprime.PopulationConfiguration(sample_size=30, initial_size = No)]

migr_matrix = [[0, 0],
               [0, 0]]

demo = [
    msprime.PopulationParametersChange(time = Tf, initial_size = No, population_id = 0,
growth_rate = 0),
    msprime.MassMigration(time = 1800, source = 1, destination = 0, proportion = 1)]

tree = msprime.simulate(population_configurations = pop_conf,
                            migration_matrix = migr_matrix,
                            length = L,
                            recombination_rate = recomb_rate,
                            mutation_rate = mut_rate,
                            demographic_events = demo)
```

#### S2.6 No recovery founder event model

**Tf**, **Nf**, **Df**: parameters set by the user (variable).

Note that we simulate four populations, but only the first one and the last one are used (respectively the target and outgroup populations).

```
L = 50e6
recomb_rate = 1e-8
mut_rate = 1.2e-8
No = 12500
Df = 1

pop_conf = [
    msprime.PopulationConfiguration(sample_size=30, initial_size = Nf),
    msprime.PopulationConfiguration(sample_size=30, initial_size = No),
    msprime.PopulationConfiguration(sample_size=30, initial_size = No),
    msprime.PopulationConfiguration(sample_size=30, initial_size = No)]
```

```
migr_matrix = [[0, 0, 0, 0],
               [0, 0, 0, 0],
               [0, 0, 0, 0],
               [0, 0, 0, 0]]

demo = [
       msprime.PopulationParametersChange(time = Tf, initial_size = Nf, population_id = 0),
       msprime.PopulationParametersChange(time = Tf+Df, initial_size = No, population_id =
0),
       msprime.MassMigration(time = 200, source = 1, destination = 0, proportion = 1),
       msprime.MassMigration(time = 1000, source = 2, destination = 0, proportion = 1),
       msprime.MassMigration(time = 1800, source = 3, destination = 0, proportion = 1)]

tree = msprime.simulate(population_configurations = pop_conf,
                            migration_matrix = migr_matrix,
                            length = L,
                            recombination_rate = recomb_rate,
                            mutation_rate = mut_rate,
                            demographic_events = demo)
```

#### S2.7.1 Impact of sample size

**Tf**, **sample_size**: parameters set by the user (variable).

```
L = 50e6
recomb_rate = 1e-8
mut_rate = 1.2e-8
No = 12500
Nf = 5
Df = 1

pop_conf = [
    msprime.PopulationConfiguration(sample_size=sample_size*2, initial_size = No),
    msprime.PopulationConfiguration(sample_size=0, initial_size = No),
    msprime.PopulationConfiguration(sample_size=0, initial_size = No),
    msprime.PopulationConfiguration(sample_size=30, initial_size = No)]

migr_matrix = [[0, 0, 0, 0],
               [0, 0, 0, 0],
               [0, 0, 0, 0],
               [0, 0, 0, 0]]

demo = [
       msprime.PopulationParametersChange(time = Tf, initial_size = Nf, population_id = 0),
       msprime.PopulationParametersChange(time = Tf+Df, initial_size = No, population_id =
0),
       msprime.MassMigration(time = 200, source = 1, destination = 0, proportion = 1),
       msprime.MassMigration(time = 1000, source = 2, destination = 0, proportion = 1),
       msprime.MassMigration(time = 1800, source = 3, destination = 0, proportion = 1)]

tree = msprime.simulate(population_configurations = pop_conf,
                            migration_matrix = migr_matrix,
                            length = L,
                            recombination_rate = recomb_rate,
                            mutation_rate = mut_rate,
                            demographic_events = demo)
```

#### S2.7.2 Impact of missing data

We use the same data as generated from **2.1**, but randomly convert some genotypes of the *EIGENSTRAT .geno* files into missing genotypes (i.e. with the value 9) using a custom script with a certain proportion of missingness set by the user.

#### S2.7.3 Impact of ancient DNA data features

**Tf**: parameter set by the user (variable).

```
L = 50e6
recomb_rate = 1e-8
mut_rate = 1.2e-8
No = 12500
Df = 1
Nf = 5

pop_conf = [
    msprime.PopulationConfiguration(sample_size=10, initial_size = No),
    msprime.PopulationConfiguration(sample_size=0, initial_size = No),
    msprime.PopulationConfiguration(sample_size=0, initial_size = No),
    msprime.PopulationConfiguration(sample_size=30, initial_size = No)]

migr_matrix = [[0, 0, 0, 0],
               [0, 0, 0, 0],
               [0, 0, 0, 0],
               [0, 0, 0, 0]]

demo = [
       msprime.PopulationParametersChange(time = Tf, initial_size = Nf, population_id = 0),
       msprime.PopulationParametersChange(time = Tf+Df, initial_size = No, population_id =
0),
       msprime.MassMigration(time = 200, source = 1, destination = 0, proportion = 1),
       msprime.MassMigration(time = 1000, source = 2, destination = 0, proportion = 1),
       msprime.MassMigration(time = 1800, source = 3, destination = 0, proportion = 1)]

tree = msprime.simulate(population_configurations = pop_conf,
                            migration_matrix = migr_matrix,
                            length = L,
                            recombination_rate = recomb_rate,
                            mutation_rate = mut_rate,
                            demographic_events = demo)
```

The *EIGENSTRAT .geno* file is further processed using a custom script in order to convert all genotypes into pseudo-homozygous genotypes by choosing one random allele at each site and doubling into into a diploid call. Also, some genotypes are converted into missing genotypes (i.e. with the value 9) using the same custom script with a certain proportion of missingness set by the user.

Ramasubramanian, Karthik, and Abhishek Singh. 2016. *Machine Learning Using R*. Apress.

Reich, David, Kumarasamy Thangaraj, Nick Patterson, Alkes L. Price, and Lalji Singh. 2009. “Reconstructing Indian Population History.” *Nature* 461 (7263): 489–94.

Sams, Aaron J., and Adam R. Boyko. 2019. “Fine-Scale Resolution of Runs of Homozygosity Reveal Patterns of Inbreeding and Substantial Overlap with Recessive Disease Genotypes in Domestic Dogs.” *G3* 9 (1): 117–23.

Sankararaman, Sriram, Nick Patterson, Heng Li, Svante Pääbo, and David Reich. 2012. “The Date of Interbreeding between Neandertals and Modern Humans.” *PLoS Genetics* 8 (10): e1002947.

Scally, Aylwyn, and Richard Durbin. 2012. “Revising the Human Mutation Rate: Implications for Understanding Human Evolution.” *Nature Reviews. Genetics* 13 (10): 745–53.

### S3. Data curation for human datasets

We applied *ASCEND* to four human datasets that we describe below:
**Human Origins Dataset (HO37):** This dataset comprises of 7,744 individuals (5,637 present-day from ~900 groups and 2,104 ancient individuals from ~300 groups) genotyped on the Human Origins array at 597,573 SNPs. The downloaded version 37.2 (https://reich.hms.harvard.edu/) was released in Narasimham et al. (2019).
**IndiaHO:** This dataset comprises of 1,662 individuals from 249 ethno-linguistic groups from India genotyped on the Human Origins array at 499,158 SNPs and was released in Nakatsuka et al. (2017).
**Behar:** We used the data for 21 Ashkenazi Jews genotyped on the Illumina 610K and 660K bead arrays from Behar et al. (2010).
**Human Origins Dataset (HO42):** This dataset comprises 253 individuals from 27 ancient Hunter-Gatherer, Neolithic Farmer and Steppe Pastoralist populations and was published in Narasimhan et al. (2019). This version can also be downloaded from the Reich Lab website (https://reich.hms.harvard.edu/).

Note that we excluded from the analysis individuals that were annotated as questionable (except for HO42 where we considered also samples with questionable assessment in order to enrich the sample sizes in relevant groups), contaminated, outliers or with “low coverage” annotation in the original *.anno* file. We additionally filtered these datasets to remove 547 groups (1,543 individuals) based on the following criteria (Supplementary Table S1):

(1) To reliably calculate allele sharing correlation, we remove groups with less than 5 individuals (1,254 individuals, 510 groups),
(2) Groups with post-filtering sample sizes of less than 5 individuals (71 individuals, 15 groups),
(3) Groups which are labeled as “Ignore” in the original dataset and usually includes duplicates, relatives or low quality samples (126 individuals, 16 groups),
(4) Groups with samples generated by whole genome amplification, indicated with suffix “.WGA” (52 individuals, 15 groups),
(5) Groups with samples indicated as low coverage or outliers (48 individuals, 14 groups).

For the groups that are retained post-filtering, we removed 317 individuals from all the datasets identified as close relatives because they have either:

(i) a pairwise genomic sharing ($\pi$) greater than 0.45 with another individual in the dataset expected for first-degree relatives. We note $\pi$ is an estimator of the proportion of genome shared identical-by-descent (IBD), or
(ii) both $\pi > 0.125$ (as expected for third-degree relatives) and at least one segment of IBD that is greater than 65 cM (almost half the length of an average chromosome) with another individual in the dataset.

For the calculation of pairwise $\pi$, we used PLINK v1.90b6.2 *genome* module (Chang et al. 2015) with the following command line:

```
plink --file XX --genome --out XX
```

For the detection of IBD segments, we first phased the samples from each dataset using EAGLE 2.4.1 with default parameters (Loh et al. 2016) and using the 1000 Genomes Project III phased samples as a reference panel to increase the phasing power (1000 Genomes Project Consortium et al. 2015):

```
eagle \
     --vcfTarget=XX.QQ.vcf.gz \
     --vcfRef=1000Genomes.QQ.vcf.gz \
     --outPrefix=XX.QQ.phased \
     --geneticMapFile=genetic_map_hg19.txt.gz \
     --chrom=QQ \
     --numThreads=4 \
     --allowRefAltSwap \
     2>&1 | tee XX.QQ.phasing.log
```

The IBD segments were called using GERMLINE 1.5.3 (Gusev et al. 2009) with default parameters and the *-genotype* extension mode.

```
germline  -input  XX.QQ.phased.012.ped  XX.QQ.phased.012.cM.map
\
     -output XX.QQ.phased.012 \
     -bits 75 \
     -err_hom 0 \
     -err_het 0 \
     -min_m 3 \
     -g_extend
```

We applied the HaploScore algorithm (Durand, Eriksson, and McLean 2014) to remove false positive IBD segments with the recommended genotype error of 0.75%, switch error of 0.3% and the threshold matrix for a mean overlap of 80% as suggested by Durand et al. (2014).

```
python2.7 haploscore.py \
     XX.QQ.phased.012.match \
     XX.QQ.phased.012.ped
     XX.QQ.phased.012.cM.map \
     XX.QQ.phased.012.haploscore \
     --genotype_error 0.0075 \
     --switch_error 0.003
     --filter 0.8 \
     --threshold_file chr21.scorethresh.txt
```

After filtering, our dataset contained 1,253 individuals from 116 populations in the IndiaHO dataset (we also excluded the 12 patients affected by Progressive Pseudorheumatoid Dysplasia (PPD) from two groups due to their disease status); 3,285 present-day and ancient individuals from 274 groups for the HO37 dataset and 253 ancient Hunter-Gatherers,

Neolithic Farmers and Steppe Pastoralists from 27 groups for the HO42 dataset. None of the individuals from the Behar dataset was removed (Supplementary Table S1).

**References**

Reich *1240K+HO* dataset. https://reich.hms.harvard.edu/downloadable-genotypes-present-day-and-ancient-dna-data-compiled-published-papers

1000 Genomes Project Consortium, Adam Auton, Lisa D. Brooks, Richard M. Durbin, Erik P. Garrison, Hyun Min Kang, Jan O. Korbel, et al. 2015. "A Global Reference for Human Genetic Variation." Nature 526 (7571): 68–74.

Behar, Doron M., Bayazit Yunusbayev, Mait Metspalu, Ene Metspalu, Saharon Rosset, Jüri Parik, Siiri Rootsi, et al. 2010. "The Genome-Wide Structure of the Jewish People." Nature 466 (7303): 238–42.

Chang, Christopher C., Carson C. Chow, Laurent Cam Tellier, Shashaank Vattikuti, Shaun M. Purcell, and James J. Lee. 2015. "Second-Generation PLINK: Rising to the Challenge of Larger and Richer Datasets." GigaScience 4 (February): 7.

Durand, Eric Y., Nicholas Eriksson, and Cory Y. McLean. 2014. "Reducing Pervasive False-Positive Identical-by-Descent Segments Detected by Large-Scale Pedigree Analysis." Molecular Biology and Evolution 31 (8): 2212–22.

Gusev, Alexander, Jennifer K. Lowe, Markus Stoffel, Mark J. Daly, David Altshuler, Jan L. Breslow, Jeffrey M. Friedman, and Itsik Pe'er. 2009. "Whole Population, Genome-Wide Mapping of Hidden Relatedness." Genome Research 19 (2): 318–26.

Loh, Po-Ru, Petr Danecek, Pier Francesco Palamara, Christian Fuchsberger, Yakir A Reshef, Hilary K Finucane, Sebastian Schoenherr, et al. 2016. "Reference-Based Phasing Using the Haplotype Reference Consortium Panel." Nature Genetics 48 (11): 1443–48.

Nakatsuka, Nathan, Priya Moorjani, Niraj Rai, Biswanath Sarkar, Arti Tandon, Nick Patterson, Gandham Srilakshmi Bhavani, et al. 2017. "The Promise of Discovering Population-Specific Disease-Associated Genes in South Asia." Nature Genetics 49 (9): 1403–7.

Narasimhan, Vagheesh M., Nick Patterson, Priya Moorjani, Nadin Rohland, Rebecca Bernardos, Swapan Mallick, Iosif Lazaridis, et al. 2019. "The Formation of Human Populations in South and Central Asia." Science 365 (6457). https://doi.org/10.1126/science.aat7487.

### S4. Comparison of overlapping groups across human datasets

For present-day humans, we used three datasets–IndiaHO, HO37 and Behar (Supplementary Note S3). Across these datasets, we found that a total of 19 groups present in both the IndiaHO and HO37 datasets. These included: Balochi, Bengali, Brahmin from Uttar Pradesh, Brahui, Burusho, Gujarati from West India (GujaratiB, GujaratiC, GujaratiD), Hazara, Irula, Ashkenazi Jews, Cochin Jews, Kalash, Kusunda, Makrani, Onge, Pathan, Punjabi and Sindhi_Pakistan.

For each of these populations, we compared the founder event parameters estimated in each dataset independently. Considering the groups where we were able to estimate reliable parameters in both datasets, we found that most results were statistically consistent across the two datasets (with overlapping 95% confidence intervals) except in case of three populations (Irula, Onge and Ashkenazi Jews) where the founder intensity or the founder age estimates differed significantly perhaps due to differences in population structure or SNP ascertainment (Table S4.1).

**Table S4.1** - Comparison of the estimated founder age and intensity obtained for populations present in multiple datasets (IndiaHO, HO37 and Behar).

| | Sample sizes+ | | Founder age (in generations) | | | Founder intensity | | |
|---|---|---|---|---|---|---|---|---|
| | IndiaHO | HO37 | IndiaHO | HO37 | Behar | IndiaHO | HO37 | Behar |
| **Balochi** | 18 | 20 | 34 ± 5 | 34 ± 4 | | 0.88% ± 0.08% | 0.86% ± 0.07% | |
| **Bengali** | 7 | 7 | -- | -- | | -- | -- | |
| **Bahmin_UP** | 5 | 10 | -- | -- | | -- | -- | |
| **Brahui** | 19 | 21 | 22 ± 1 | 20 ± 1 | | 1.16% ± 0.04% | 1.17% ± 0.05% | |
| **Burusho** | 20 | 23 | 12 ± 1 | 13 ± 1 | | 0.82% ± 0.03% | 0.81% ± 0.04% | |
| **GujaratiB** | 5 | 5 | -- | -- | | -- | -- | |
| **GujaratiC** | 5 | 5 | -- | -- | | -- | -- | |
| **GujaratiD** | 5 | 5 | 75 ± 24 | 123 ± 30 | | 2.19% ± 0.42% | 3.62% ± 0.52% | |
| **Hazara** | 11 | 16 | 15 ± 3 | -- | | 0.64% ± 0.08% | -- | |
| **Irula** | 15 | 10 | 34 ± 2 | 18 ± 1 | | 3.05% ± 0.11% | 1.95% ± 0.09% | |
| **Jew_Ashkenazi** | 7 | 7 | 43 ± 8 | 37 ± 7 | 31 ± 2 | 2.01% ± 0.23% | 1.7% ± 0.19% | 1.04% ± 0.05% |
| **Jew_Cochin** | 13 | 5 | -- | 10 ± 1 | | -- | 2.39% ± 0.18% | |
| **Kalash** | 15 | 16 | 17 ± 1 | 16 ± 1 | | 5.62% ± 0.13% | 5.62% ± 0.14% | |
| **Kusunda** | 7 | 7 | 6 ± 1 | 7 ± 1 | | 2.62% ± 0.16% | 3.43% ± 0.24% | |
| **Makrani** | 18 | 19 | 20 ± 2 | 15 ± 2 | | 0.84% ± 0.05% | 0.94% ± 0.05% | |
| **Onge** | 7 | 6 | 20 ± 1 | 21 ± 2 | | 20.63% ±0.54% | 8.12% ± 0.4% | |

| | | | | | | | |
|---|---|---|---|---|---|---|---|
| **Pathan** | 18 | 17 | -- | -- | | -- | -- |
| **Punjabi** | 8 | 9 | 41 ± 15 | 68 ± 16 | | 0.89% ± 0.18% | 1.28% ± 0.18% |
| **Sindhi_Pakistan** | 17 | 14 | -- | -- | | -- | -- |

Notes.
(in red) Results shown in red significantly differ across the two datasets compared.
\+ Indicates the post-filtering sample size.
-- Indicates that we were unable to obtain a reliable fit due to one of the criteria: the 95% confidence interval of either $T_f$ or $I_f$ included 0; NRMSD>0.29; $T_f$>200 generations; $I_f$<0.5% or the standard error of $T_f$>50 generations.
(blank) For Behar, we only had estimates for Ashkenazi Jews, the rest of the entries in the table are thus blank.

To investigate if the cause of discrepancy in results for Irula and Onge is driven by population structure, we merged the IndiaHO and HO37 datasets with the 1000 Genomes Project Phase 3 dataset (1000 Genomes Project Consortium et al. 2015) and performed a Principal Component Projection Analysis (PCA) using *smartpca* (Patterson, Price, and Reich 2006). We computed the eigenvectors using three populations from the 1000 Genomes Project: Han Chinese in Beijing, China (CHB) as a proxy for East-Asian ancestry, Northern and Western Europeans (CEU) as a proxy to West Eurasian ancestry and Indian Telugu (ITU) as a proxy for South Asian ancestry, and projected our target populations (Irula and Onge) onto these eigenvectors. We also investigated the population structure in Ashkenazi Jews using the same procedure but after merging the IndiaHO dataset with the 1000 Genomes Project Phase 3 dataset and the Behar dataset (we did not include the HO37 dataset as the Ashkenazi Jew individuals from HO37 are the same as in IndiaHO).

We found that the Irula population and the Ashkenazi Jews are fairly heterogeneous but there were no major clusters seen on PC1 and PC2 (Figure S4.1 and Figure S4.2 respectively). However, for Onge, the individuals do seem cluster into two distinct groups on the PC2, suggesting that the results could be driven by population structure (Figure S4.1).

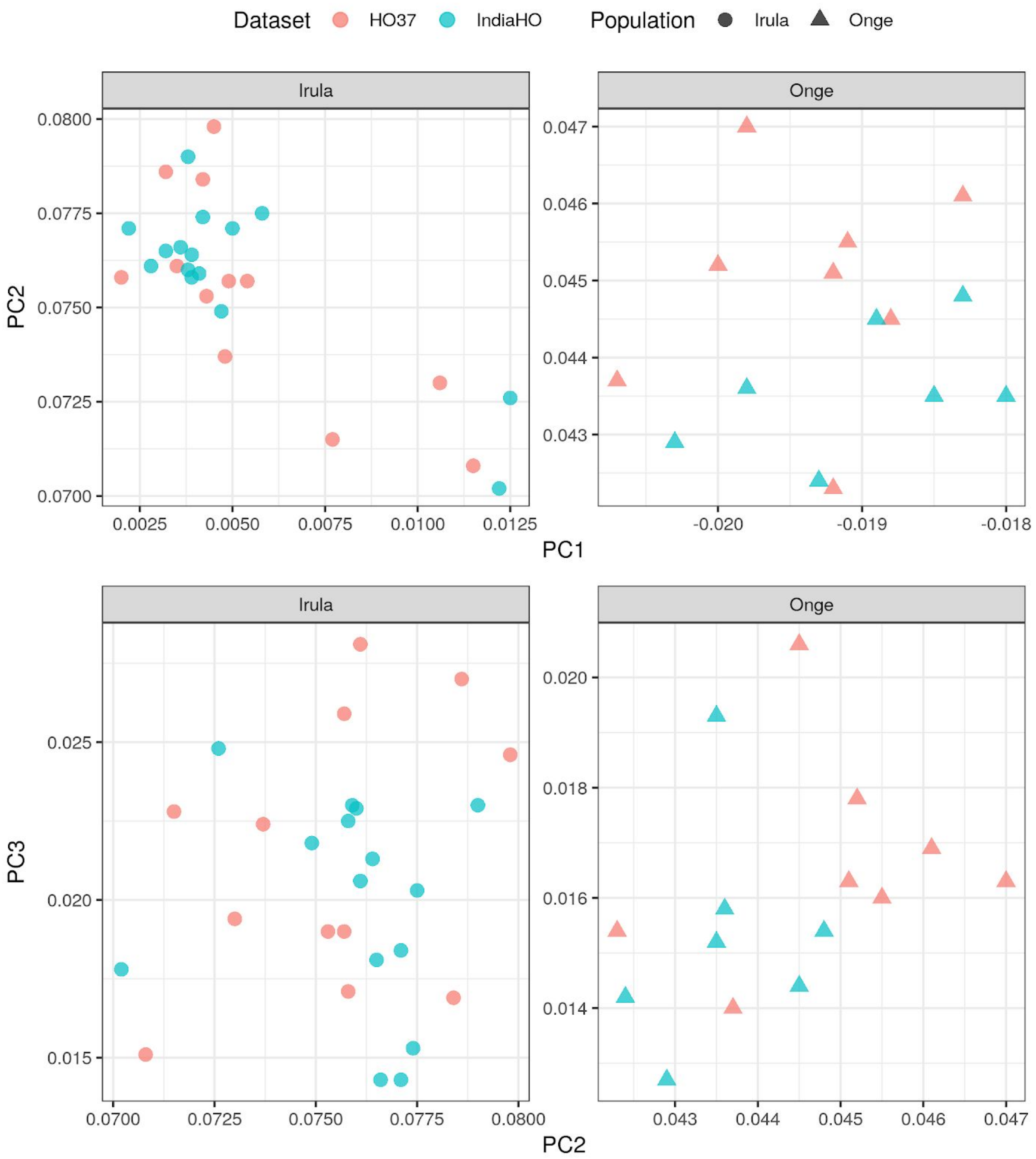

**Figure S4.1** - Distribution of the Onge (left panel) and Irula (right) individuals in the space of the first three principal components trained on the Telugu (ITU), Han (CHB) and North-Western Europeans (CEU) individuals, using the IndiaHO (blue) and HO37 (red) datasets.

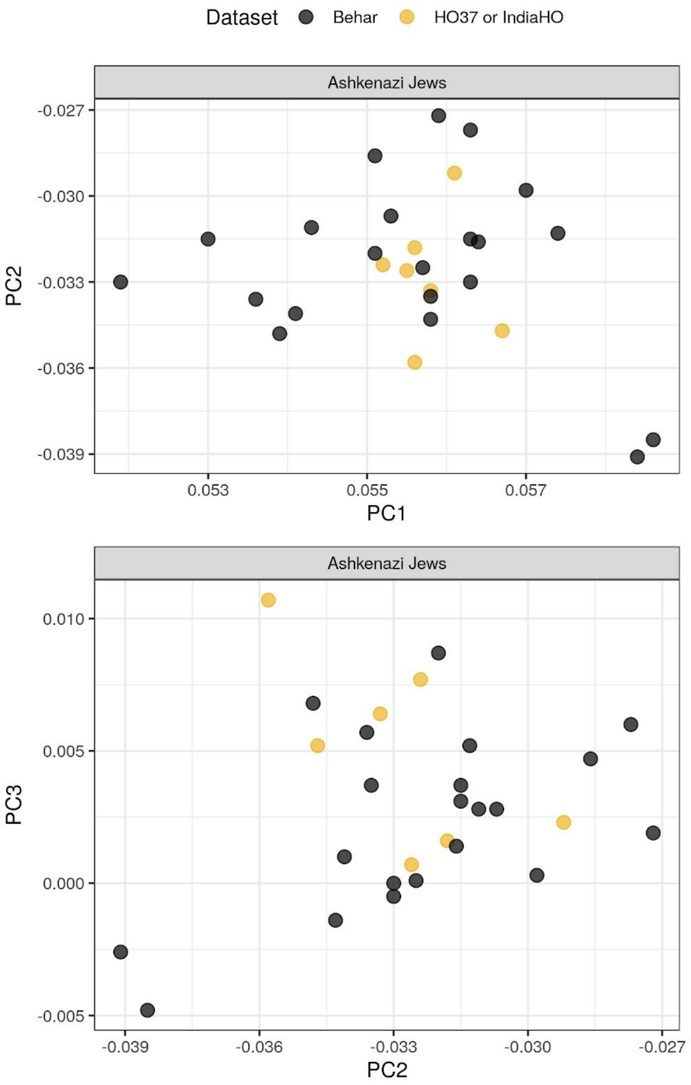

**Figure S4.2** - Distribution of the Ashkenazi Jew individuals in the space of the first three principal components trained on the Telugu (ITU), Han (CHB) and North-Western Europeans (CEU) individuals from the 1000 Genomes Project, using the IndiaHO (golden points) and Behar (black points) datasets.

### S5. Comparison of our results with published estimates

#### **S5.1** Comparison of dates of founder event with Reich et al. 2009

*ASCEND* is an extension of the allele sharing correlation statistic introduced in Reich et al. (2009) that was applied to date founder events in 25 Indian groups. To compare the performance of *ASCEND* with the previous study, we applied *ASCEND* to the original dataset from Reich et al. with the exception of one group (Siddis) for which some samples were missing.

To reliably compare the results, we matched the setup in Reich et al. and separately analyzed the populations on the Indian Cline (the cline formed by the relatedness of Indian groups to West Eurasian populations) and those not belonging to the Indian Cline (mostly Austro-Asiatic and Tibeto-Burmese speaking groups). Most populations on the Indian Cline can be modeled as a mixture of Ancestral North Indian (ANI) related to West Eurasians and Ancestral South Indian (ASI) distantly related to Andamanese Islanders, Onge (Reich et al. (2009)). Specifically, we ran the analysis as follows:

(i) For each population on the Indian Cline, we computed the within population allele sharing correlation across individuals in the target population and then subtracted the cross-population correlation computed between the individuals in the target and individuals on the Indian Cline that best matched the proportion of ANI ancestry estimated in the target population, using the following categories similar to Reich et al. (2009):

- 65% ± 5% ANI ancestry: Meghawal, Vaish and Kashmiri Pandit;
- 58% ± 5% ANI ancestry: Velama, Srivastava, Meghawal and Vaish;
- 53% ± 5% ANI ancestry: Lodi, Naidu, Tharu, Velama and Srivastava;
- 47% ± 5% ANI ancestry: Bhil, Satnami, Kurumba, Kamsali, Vysya, Lodi, Naidu and Tharu;
- 42% ± 5% ANI ancestry: Mala, Madiga, Chenchu, Bhil, Satnami, Kurumba, Kamsali and Vysya.

If a target population was present in two categories, then we used the reference populations present in both the categories of outgroups.
(ii) For each population not belonging to the Indian Cline (Aonaga, Great_Andamanese, Hallaki, Kharia, Nysha, Onge, Sahariya, Santhal), we report estimates for just the within-population allele sharing correlation. Reich et al. do not describe how the cross-population correlation was computed for the off-cline populations or if it was subtracted at all and hence we do not include a correction either.
(iii) We removed any monomorphic SNPs and computed allele sharing correlations between genetic distances from 0.1 to 30 cM with a bin size of 0.1 cM (the default in *ASCEND*).

Table S5.1.1 shows the comparison of the founder age estimates across the two methods. We were able to reliably infer the parameters for 11 populations using *ASCEND*, compared to 12 in Reich et al (this does not include Siddis that was missing in our dataset). For the overlapping 11 groups, we found that results were consistent for all Indian Cline populations, except Bhil. Our dates of founder event were very recent in Bhil (19 [4–35]), unlike reported in Reich et al. that estimated it at 40 generations ago. We note that the results for Bhil were very noisy as indicated by the NRMSD of 0.285 (Figure S.5.1.1). Interestingly, a recent study suggested that the founder event in Bhil occurred ~15 generations ago using a different approach, *IBDNe* (Browning & Browning (2015)) for inference (Debortoli et al. 2020).

For the non-Indian Cline populations, there were two groups for which founder age estimates differed significantly between Reich et al. and *ASCEND*: Great_Andamanese and Onge, both from the Andamanese Islands (Table S5.1.1). Visual inspection of the allele sharing correlation decay curves suggests that the maximum distance used in the analysis could impact the results (Figure S.5.1.1). This is likely exacerbated in groups with a history of multiple or continuous founder events, where we may expect that the decay curve has not fully captured all the IBD segments if the maximum distance threshold is not large enough (which is set as 10 cM in Reich et al. and 30 cM in our analysis). As that the Andamanese populations likely have a history of maintaining a small population size (as evidenced from their low census size), we believe this is likely contributing to the difference in the results we see across the two methods.

**Table S5.1.1** - Comparison of founder age estimates (in generations) based on Reich et al. (2009) and *ASCEND*.

| Population | Cline* | Reich et al. (2009)+ | | This study (using *ASCEND-FFT*) | |
|---|---|---|---|---|---|
| | | Sample size | Founder age+ | Sample size | Founder age Mean [95% CI] |
| Aonaga | NIC | 4 | 120 | 4 | 140 [98–181] |
| Bhil | IC | 7 | 40 | 7 | 19 [4–35] |
| Chenchu | IC | 6 | 10 | 6 | 9 [7–10] |
| Great_Andamanese | NIC | 7 | 14 | 7 | 6 [5–8] |
| Hallaki | NIC | 7 | 32 | 7 | -- |
| Kamsali | IC | 4 | -- | 4 | -- |
| Kashmiri_Pandit | IC | 5 | -- | 5 | -- |
| Kharia | NIC | 6 | 42 | 6 | 69 [4–134] |
| Kurumba | IC | 9 | -- | 9 | -- |
| Lodi | IC | 5 | -- | 5 | -- |
| Madiga | IC | 4 | -- | 4 | -- |
| Mala | IC | 3 | -- | 3 | -- |
| Meghawal | IC | 5 | 59 | 5 | 53 [28–77] |
| Naidu | IC | 4 | -- | 4 | -- |
| Nysha | NIC | 4 | 134 | 4 | 145 [97–192] |

| | | | | | |
|---|---|---|---|---|---|
| Onge | NIC | 9 | 39 | 9 | 31 [28–34] |
| Sahariya | NIC | 4 | 108 | 4 | 73 [34–112] |
| Santhal | NIC | 7 | -- | 7 | -- |
| Satnami | IC | 4 | -- | 4 | -- |
| Siddi | IC | 4 | 8 | 2 | NA |
| Srivastava | IC | 2 | -- | 2 | -- |
| Tharu | IC | 9 | -- | 9 | -- |
| Vaish | IC | 4 | -- | 4 | -- |
| Velama | IC | 4 | 88 | 4 | 59 [17–101] |
| Vysya | IC | 5 | 108 | 5 | 72 [23–121] |

Notes.
(in red) Results shown in red significantly differ across the two datasets compared.
-- Indicates that we were unable to obtain a reliable fit due to one of the criteria: the 95% confidence interval of either $T_f$ or $I_f$ included 0; NRMSD>0.29; $T_f$>200 generations; $I_f$<0.5% or the standard error of $T_f$>50 generations.
+ Confidence intervals are not shown as none were reported in Reich et al. (2009).
NA Sample size differed from the original study and was less than 4, hence *ASCEND* was not run on this group.
* IC = population is on the Indian Cline; NIC = population is not on the Indian Cline.

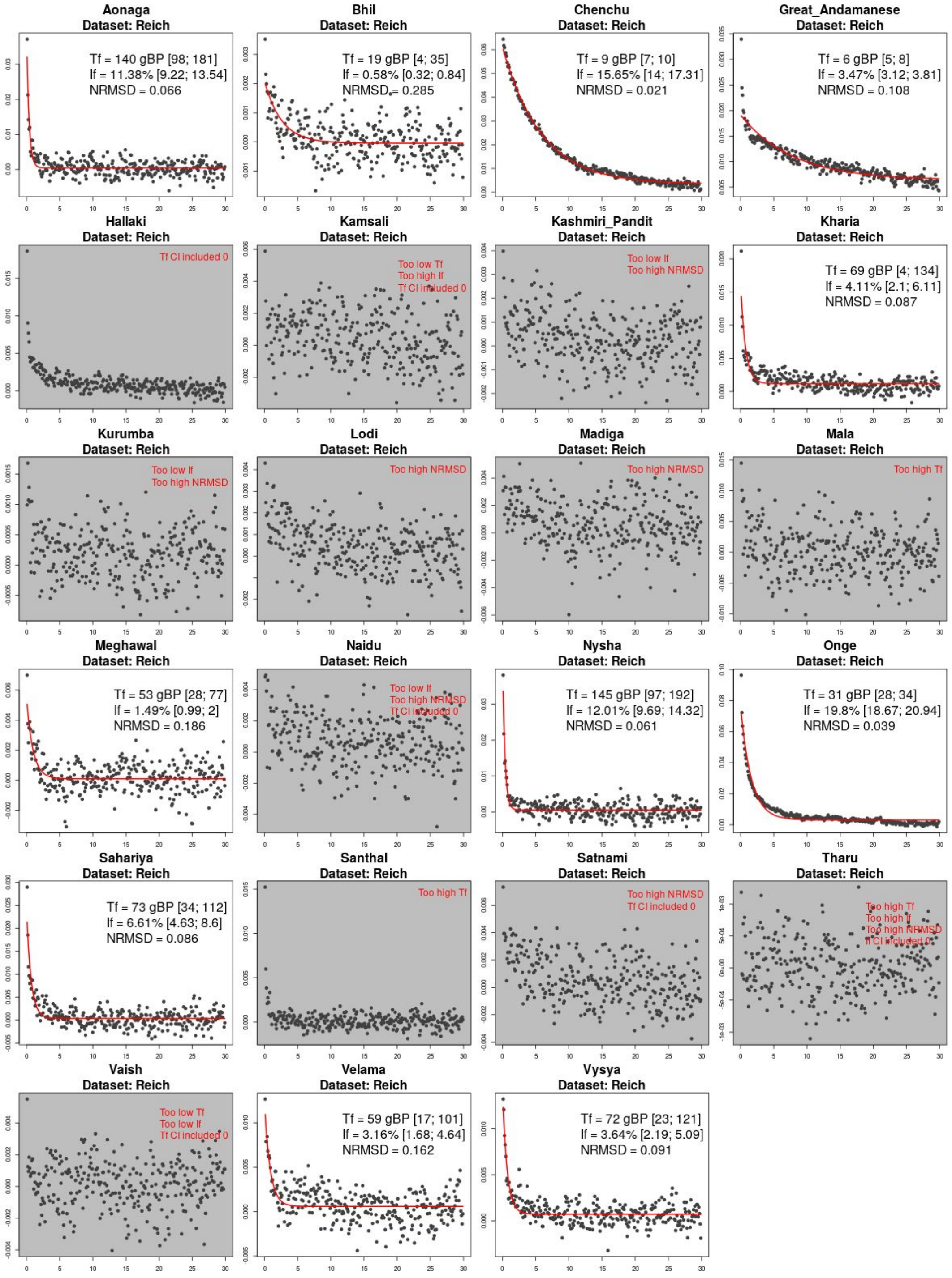

**Figure S5.1.1** - Allele sharing correlation curves for all the samples from Reich et al. that were analyzed with *ASCEND*. The x-axis shows the genetic distance in cM and the y-axis shows the mean allele sharing correlation. The black points represent the values calculated on the empirical dataset. The red curve represents the exponential fit. For each population panel, we report the estimated mean and with the 95% confidence interval for the founder age $T_f$ (in generations) and the founder intensity $I_f$, and the value of the NRMSD (which is used to measure the quality of the fitted exponential). The panel is greyed if the exponential fit failed or if the inference was unreliable (i.e., the estimated age was greater than 200 generations or the standard error of the founder age was greater than 50 generations or the intensity was lower than 0.5% or the NRMSD greater than 0.29 or the 95% confidence interval for either the age or intensity contained negative values).

#### **S5.2.** Comparison of founder event intensity and IBD scores with Nakatsuka et al. (2017)

Nakatsuka et al. (2017) introduced the Identity-By-Descent (IBD) score to measure the strength of the founder events. Specifically, the IBD score measures the average length of IBD segments in the range 3-20 cM shared between pairs of individuals in a population, normalized by the number of pairwise comparisons (related to the sample size). To obtain a relative IBD score, this estimate is divided by the IBD score calculated in Finns or Ashkenazi Jews (Nakatsuka et al. 2017). Using this approach, Nakatsuka et al. identified 81 South Asian groups in the IndiaHO dataset that had an IBD score higher than estimated in Ashkenazi Jews (AJ).

To infer the strength of the founder event, *ASCEND* estimates an intensity parameter ($I_f$) which is related to the population size and the duration of the founder event. This parameter should be qualitatively capturing the same signal as the IBD score. To compare the two estimates, we inferred the founder intensity in the IndiaHO populations using *ASCEND* for 116 groups in the IndiaHO dataset that had a sample size of at least 5 individuals (using the standard *ASCEND* setup, see Methods for details).

Among these 116 groups, the founder intensity $I_f$ was reliably characterized for 66 populations (based on the criteria that they check all of the following conditions: NRMSD < 0.29, $T_f$ < 200 generations, standard error of $T_f$ < 50 generations, $I_f > 5\times10^{-3}$, the 95% confidence intervals of $T_f$ and $I_f$ do not include 0). For the 66 overlapping groups, we observed that the strength of the founder event estimated using $I_f$ in *ASCEND* and the IBD score in Nakatsuka et al. (2017) were strongly correlated (Pearson's r=0.95, $P<10^{-5}$) (Figure S5.2.1, Table S5.2.1).

We note that unlike Nakatsuka et al., we find that out of the 65 populations (excluding the AJ used for comparison), only 16 populations have founder events stronger than in AJ compared to 32 groups suggested based on the IBD score analysis. This discrepancy can be caused by the choice of the threshold used for the estimated intensity in AJ. In our analysis, across three datasets (IndiaHO, HO37 and Behar), we observe that the intensity estimates in AJ vary markedly (1.7% for HO37, 2.0% for IndiaHO and 1.0% for Behar et al.). Using the estimates in Behar (with the largest sample size), we find that 35 South Asian populations have a stronger bottleneck than Ashkenazi Jews, which is more in line with Nakatsuka et al. (2017).

**Table S5.2.1** - Comparison of the IBD scores published in Nakatsuka et al. (2017) with the founder intensities calculated for 67 populations that had reliable estimates of the strength of the founder event based on both methods. The populations are sorted by increasing order of IBD score.

| **Population** | **Nakatsuka et al. (2017)** | **This study (using *ASCEND*)** |
|---|---|---|
| | **IBD Score [95% CI]** | **If [95% CI]** |
| Kondh_TN | 0.05 [0–0.11] | 1.9% [1.3%–2.6%] |
| Thakur | 0.11 [0–0.15] | 0.5% [0.3%–0.7%] |
| Rajbanshi | 0.23 [0–0.32] | 1.2% [0.8%–1.6%] |
| Meena | 0.27 [0–0.48] | 1.2% [0.6%–1.7%] |
| Bhil | 0.28 [0–0.42] | 0.6% [0.4%–0.7%] |
| Kunabi | 0.36 [0–0.65] | 1.6% [0.7%–2.5%] |
| Khairwar | 0.38 [0–0.56] | 1.8% [1.4%–2.2%] |
| Handigodu | 0.45 [0–0.61] | 1.3% [1.1%–1.5%] |
| Panta_Kapu | 0.46 [0–0.53] | 0.6% [0.5%–0.8%] |
| Makrani | 0.56 [0–0.64] | 0.8% [0.7%–0.9%] |
| Patel | 0.64 [0–0.83] | 1.3% [0.9%–1.8%] |
| Bhumij_Orissa | 0.64 [0–0.86] | 1.7% [1.2%–2.2%] |
| Balochi | 0.7 [1–0.81] | 0.9% [0.7%–1%] |
| Magar | 0.71 [1–0.8] | 1.5% [1.4%–1.7%] |
| GujaratiD | 0.78 [1–1] | 2.2% [1.4%–3%] |
| Hazara | 0.84 [1–1.01] | 0.6% [0.5%–0.8%] |
| Kamboj | 0.86 [1–0.94] | 1.1% [1%–1.2%] |
| Jew_Ashkenazi | 0.88 [1–1.16] | 2% [1.6%–2.5%] |
| Minero | 0.88 [1–1.2] | 0.7% [0.6%–0.9%] |
| Brahmin_Catholic_Goa | 0.89 [1–1.04] | 0.6% [0.5%–0.7%] |
| Brahmin_Catholic_Kumta | 0.9 [1–1.09] | 0.6% [0.4%–0.8%] |
| Newar | 0.9 [0–1.4] | 1.7% [1%–2.4%] |
| Nadar | 0.92 [1–1.07] | 1.3% [1.1%–1.4%] |
| Oswal_Jain | 0.98 [1–1.4] | 1.7% [1.5%–2%] |
| Punjabi | 1.05 [1–1.28] | 0.9% [0.5%–1.3%] |
| Kondakamari | 1.12 [1–1.34] | 1.6% [1.4%–1.7%] |
| Hallaki | 1.15 [1–1.43] | 0.8% [0.6%–1%] |
| Manjhi_Jharkhand | 1.21 [1–1.67] | 1.6% [1.1%–2.1%] |
| Vysya | 1.22 [1–1.31] | 2.5% [2.4%–2.7%] |
| Arunthathiyar | 1.26 [1–1.44] | 0.8% [0.6%–0.9%] |
| Garasia | 1.28 [1–1.83] | 1.4% [1.1%–1.8%] |
| Ho_Orissa | 1.32 [1–1.43] | 1.1% [0.9%–1.2%] |
| Jogi | 1.47 [1–1.92] | 1% [0.9%–1.2%] |
| Sahariya_MP | 1.54 [1–1.93] | 1.7% [1.2%–2.1%] |

| | | |
|---|---|---|
| Batudi | 1.55 [1–2.01] | 1.4% [1%–1.7%] |
| Yerukali | 1.62 [1–1.98] | 2% [1.6%–2.3%] |
| Kallar | 1.68 [1–1.88] | 1.2% [1.1%–1.2%] |
| Brahui | 1.82 [2–2.02] | 1.2% [1.1%–1.2%] |
| Burusho | 1.85 [2–2.1] | 0.8% [0.8%–0.9%] |
| Kharia | 1.87 [1–2.25] | 0.8% [0.7%–1%] |
| Reddy_Telangana | 1.96 [1–2.57] | 1% [0.8%–1.2%] |
| Bharia | 2.18 [2–2.78] | 1.5% [1.2%–1.8%] |
| Kumhar | 2.27 [2–2.5] | 3.1% [3%–3.3%] |
| Irula | 2.47 [2–2.79] | 3% [2.8%–3.3%] |
| Havik | 2.98 [2–3.52] | 2.7% [2.2%–3.1%] |
| Juang | 3.07 [3–3.4] | 2.6% [2.4%–2.8%] |
| Hindumalayali | 3.68 [3–4.66] | 2.4% [1.9%–2.9%] |
| Asur | 4.17 [3–4.87] | 2.2% [1.9%–2.5%] |
| Parhaiya | 4.38 [3–5.32] | 4.2% [3.5%–4.9%] |
| Yadav_Pondicherry | 4.4 [4–5.04] | 1.4% [1.2%–1.6%] |
| Gorait | 4.68 [4–5.82] | 1.6% [1.4%–1.8%] |
| Palliyar | 5.11 [5–5.51] | 3% [2.9%–3.2%] |
| Kotwalia | 5.55 [4–6.92] | 2.7% [2.2%–3.3%] |
| Kurumans | 6.49 [5–7.78] | 5.9% [5.4%–6.5%] |
| Bhunjiya | 6.53 [5–7.94] | 3.9% [3.4%–4.4%] |
| Kusunda | 7.29 [6–8.33] | 2.6% [2.3%–2.9%] |
| Kanjad | 7.8 [6–9.27] | 3.2% [2.7%–3.6%] |
| Mohali | 9.12 [8–10.04] | 3.1% [2.9%–3.4%] |
| Gujjar | 11.63 [9–14.59] | 4.6% [3.8%–5.5%] |
| Kalash | 12.45 [11–13.64] | 5.6% [5.4%–5.9%] |
| Pulliyar | 16.71 [15–18.59] | 6.8% [6.5%–7.1%] |
| Hakki_Pikki | 25.56 [22–29.57] | 11.2% [10.3%–12.2%] |
| Narikuravar | 25.71 [23–28.82] | 10.6% [9.3%–12%] |
| Ulladan | 25.89 [23–28.61] | 12.4% [11.7%–13.1%] |
| Onge | 31.1 [28–33.89] | 20.6% [19.6%–21.7%] |
| Malaikuravar | 34.13 [29–38.98] | 14.7% [12.7%–16.7%] |

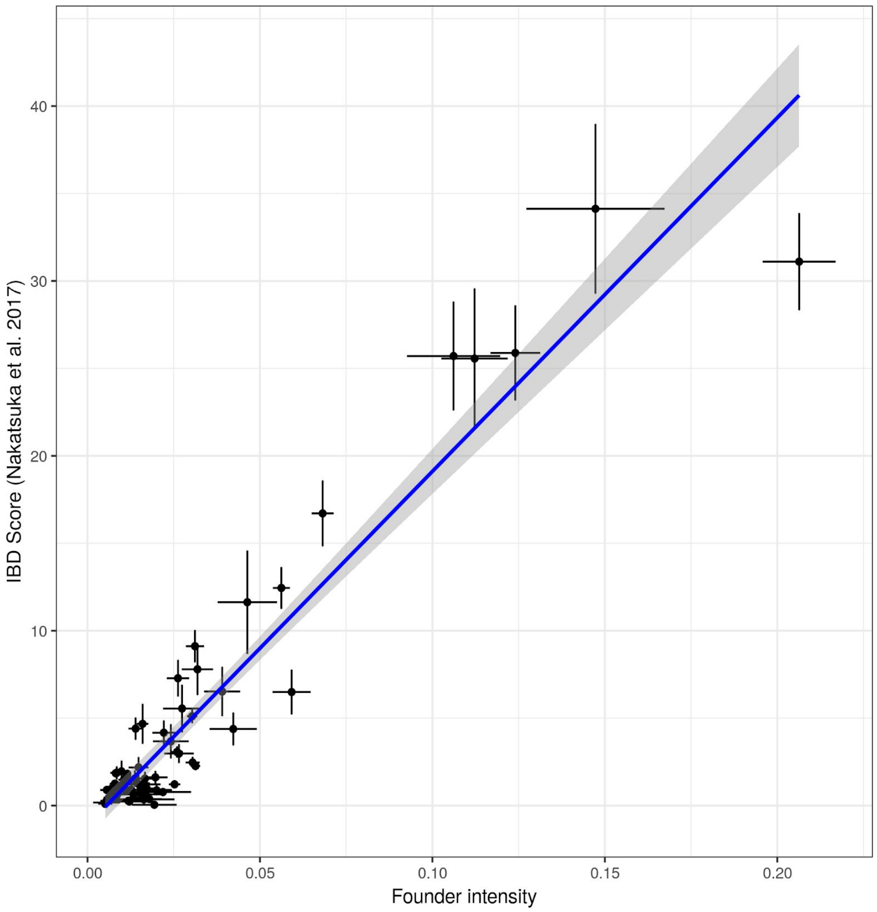

**Figure S5.2.1** - Comparison of the IBD scores published in Nakatsuka et al. (2017) with the founder intensities estimated using *ASCEND*. We report results for the comparison of 67 populations for which reliable estimates were available for both methods. The bars represent the 95% confidence intervals and the blue line is the linear regression line (Pearson's r=0.95, $P<10^{-5}$).

### S5.3. Comparison of founder intensity with the ROH statistic

The inheritance of identical haplotypes from a common ancestor can create long regions of homozygous genotypes known as runs of homozygosity (ROH). ROH are ubiquitous in human populations, and they correlate with pedigree inbreeding, especially consanguinity (Pemberton et al. 2012; Kirin et al. 2010).

To investigate how ROH correlates with the founder intensity estimated using *ASCEND*, we estimated ROH for all groups with at least 5 individuals. For groups with larger sample sizes, we randomly picked 5 individuals for the analysis. We identified ROH on autosomes for all populations using *PLINK* with options *--homozyg group* and *--homozyg-match 0.99*. The distribution of ROH lengths was summarised for each group using: the mean ROH tract length averaged across all individuals in the group ($\mathbb{E}(\bar{L})$), the total sum of ROH tract lengths averaged across all individuals of the population ($\mathbb{E}(L_{tot})$) and the maximum ROH length observed in the group ($max(L_{max})$). The corresponding ROH statistics are included in the Supplementary Table S2.

For the populations in the IndiaHO dataset that have evidence for a significant founder events, we observed that $\mathbb{E}(\bar{L})$ ranged from 1.4 Mbp (Ho_Orissa) to 4.3 Mbp (Malaikuravar) with a mean ~2.1 Mbp, $\mathbb{E}(L_{tot})$ from 23.0 Mb (Nadar) to 349.3 Mbp (Onge) with a mean ~94.2 Mbp and $max(L_{max})$ from 3.5 Mbp (Handigodu) to 69.4 Mbp (Malaikuravar) with a mean of 19.7 Mbp. The three ROH statistics were significantly correlated to the estimated founder intensity (across the correlation with each of the three ROH statistics, Pearson's r are greater than 0.3 with P-values lower than 0.014), with the strongest correlation of $I_f$ with $\mathbb{E}(L_{tot})$ statistic (Pearson r=0.82, P<$10^{-20}$) (Figure S5.2.2). The five groups with the highest $I_f$ and IBD scores in the IndiaHO dataset (Hakki Pikki, Malaikuravar, Narikuravar, Onge, Ulladan) also had the highest $\mathbb{E}(L_{tot})$ values (Figure S5.2.2, red points).

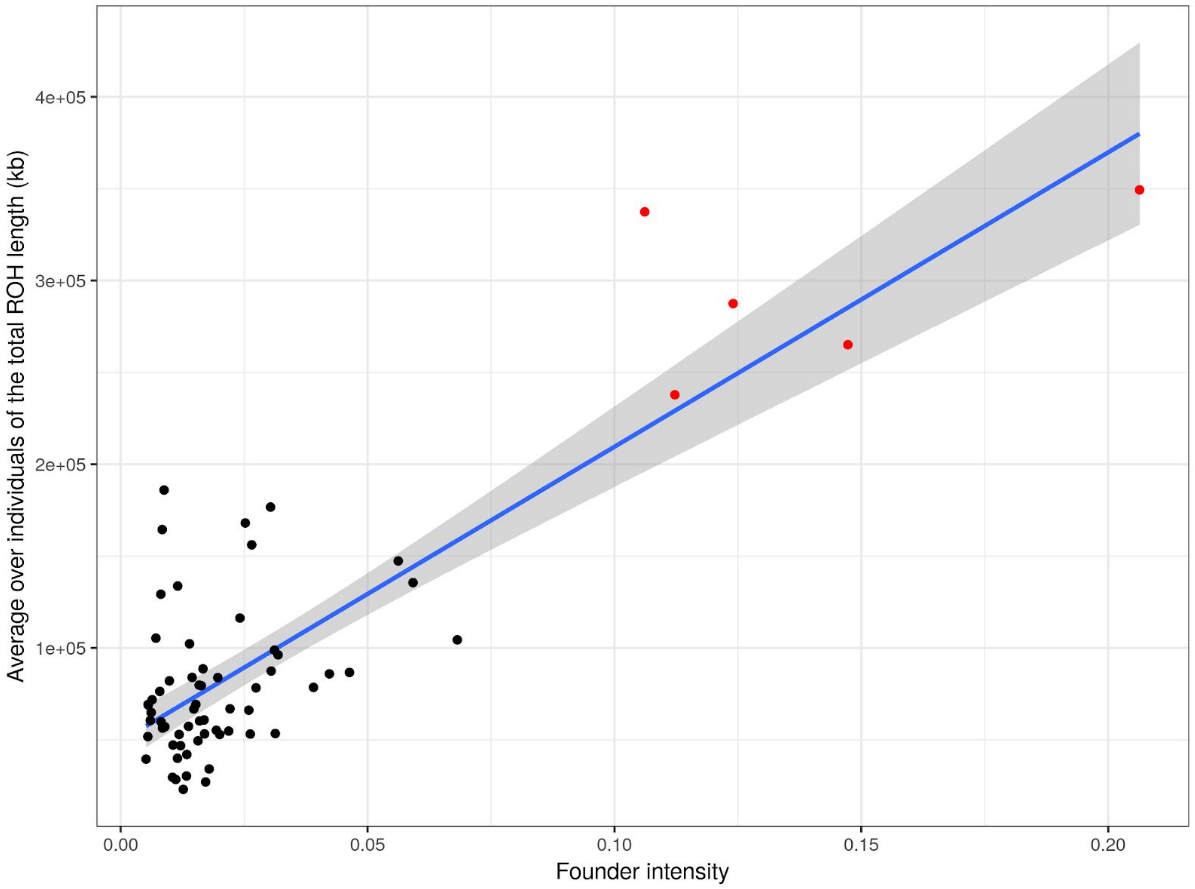

**Figure S5.2.2** - Distribution of the $\mathbb{E}(L_{tot})$ statistics in kilobases as a function of the estimated founder intensities for the founder populations from the curated IndiaHO dataset. We colored in red the five populations with extreme measures of $I_f$, IBD score and $\mathbb{E}(L_{tot})$ (Hakki Pikki, Malaikuravar, Narikuravar, Onge, Ulladan). The blue line is the best linear fit (Pearson's r=0.82, P<$10^{-20}$).

Pemberton, Trevor J., Devin Absher, Marcus W. Feldman, Richard M. Myers, Noah A. Rosenberg, and Jun Z. Li. 2012. “Genomic Patterns of Homozygosity in Worldwide Human Populations.” The American Journal of Human Genetics.

### S6. History of founder events in dogs

To study founder events in dogs, we applied *ASCEND* to two publicly available datasets of genome-wide data from dogs: (i) **Sams dataset** comprising of 1,792 individuals from 11 breeds genotyped on 175,123 SNPs (Sams and Boyko 2019), and (ii) **Hayward dataset** comprising of 4,342 individuals from 198 breeds genotyped on 160,723 SNPs (Hayward et al. 2016). We used the genetic positions from CanFam3.1 genetic map (Auton et al. 2013).

For both datasets, we filtered out SNPs and individuals with missingness greater than 1% and 5% respectively. To remove close relatives, we computed the pairwise genetic sharing π between all pairs of individuals using PLINK v1.90b6.2 *genome* module and we excluded one individual for each pair of individuals with π greater than 45%.

```
plink --vcf XX.vcf --genome --min 0.05 --double-id --dog
```

To control for inbreeding, we excluded any individual with at least one tract of run of homozygosity (ROH) longer than 30.2 Mbp (~30 cM) as this represents half the size of an average chromosome in dogs. To this end, we ran PLINK *homozyg* module:

```
plink --vcf XX.vcf --homozyg group --homozyg-kb 30000
--homozyg-match 0.99 --out XX.ROH --double-id --dog
```

After applying this filter, we considered only those populations with a sample size equal or greater than 5 individuals. We analyzed 42 unique dog breeds (1,097 individuals) across the two datasets (Supplementary Table S5, Supplementary Figure S6 for all the decay curves). The estimated founder ages and founder intensities are illustrated in the Figure 5 of the main text. Overall, we found that 40 out of the 42 dog breeds have evidence of founder events stronger than estimated in Ashkenazi Jews (~1.7%), with a mean at 25.3% [21.1%–29.6%] and a maximum of 77.7% estimated in Boxers. The founder ages range between 6 generations BP (Gordon Setter) to 27 generations BP (Bulldog), with a mean of 15 generations BP across all breeds, translating into 45-75 years BP assuming a generation time of 3-5 years (Wang et al. 2013; Vonholdt et al. 2008).

We compared the founder parameter estimates for 10 overlapping breeds present in both datasets and obtained statistically similar results (Figure S6.1, Supplementary Table S6). Both the founder age and founder intensity were highly correlated across the two datasets (Pearson's $r$>0.90, P<0.0005).

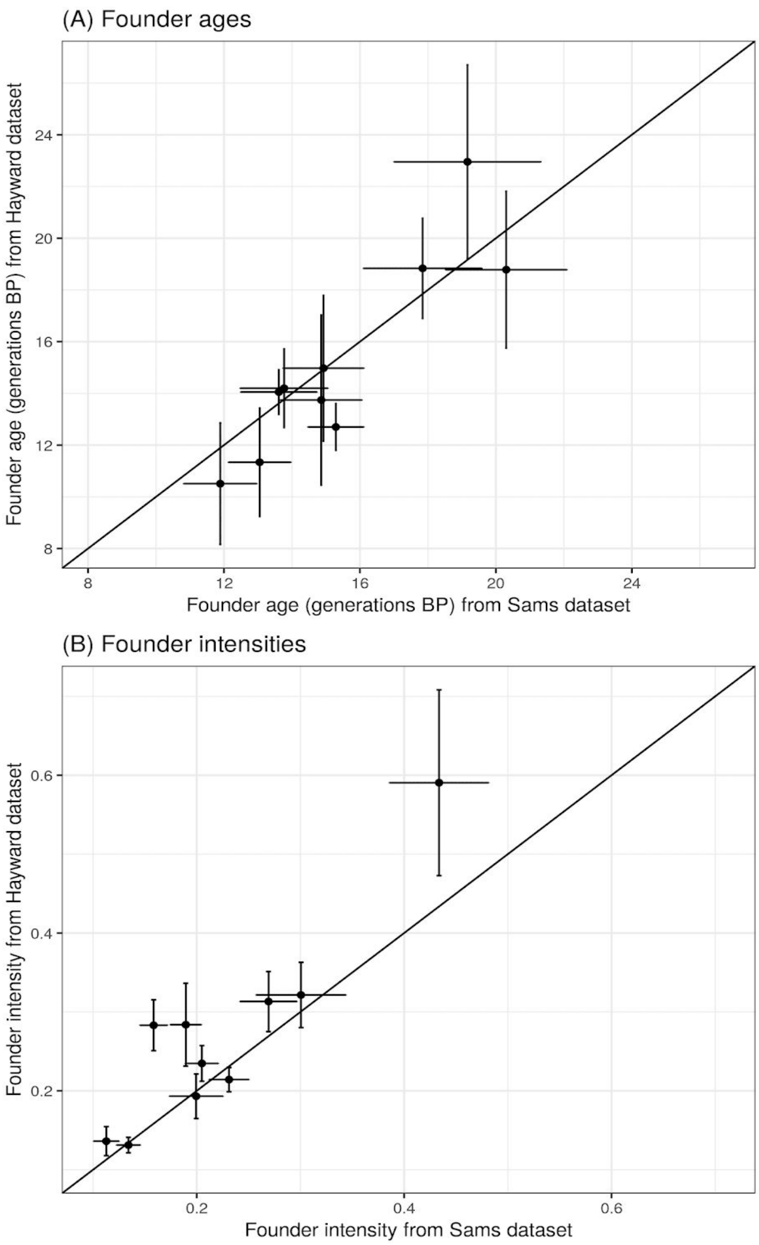

**Figure S6.1** - Comparison in the estimates of founder ages (A) and founder intensities (B) for the 10 dog breeds present in both the Sams (x-axis) and Hayward (y-axis) datasets. The black oblique line is the *y*=*x* diagonal.

**Figure S1 - Decay curves for present-day human populations from the HO37, Behar and IndiaHO datasets.** The x-axis represents the genetic distance (in cM) and the y-axis represents the average allele sharing correlation. The legend shows the mean and 95% confidence interval for the founder age ($T_f$) in generations before present (gBP) and the founder intensity ($I_f$), as well as the NRMSD (see Methods). The panels are greyed when the exponential fitting failed or when the evidence for the founder event was not significant (see Methods). The specific reason is highlighted in red in the legend.

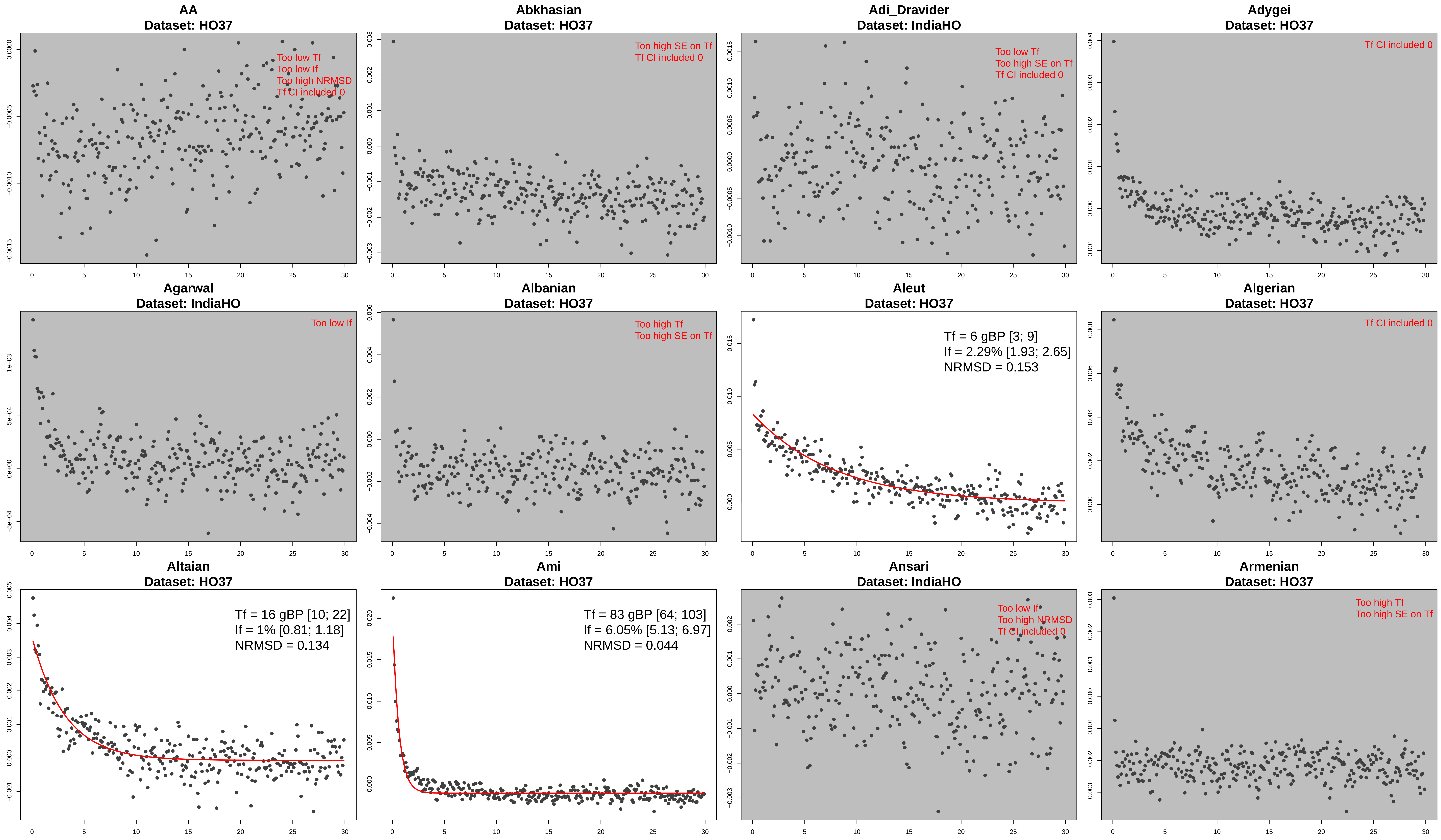
AA
Dataset: HO37
Too low Tf
Too low If
Too high NRMSD
Tf CI included 0
Abkhasian
Dataset: HO37
Too high SE on Tf
Tf CI included 0
Adi_Dravider
Dataset: IndiaHO
Too low Tf
Too high SE on Tf
Tf CI included 0
Adygei
Dataset: HO37
Tf CI included 0
Agarwal
Dataset: IndiaHO
Too low If
Albanian
Dataset: HO37
Too high Tf
Too high SE on Tf
Aleut
Dataset: HO37
Tf = 6 gBP [3; 9]
If = 2.29% [1.93; 2.65]
NRMSD = 0.153
Algerian
Dataset: HO37
Tf CI included 0
Altaian
Dataset: HO37
Tf = 16 gBP [10; 22]
If = 1% [0.81; 1.18]
NRMSD = 0.134
Ami
Dataset: HO37
Tf = 83 gBP [64; 103]
If = 6.05% [5.13; 6.97]
NRMSD = 0.044
Ansari
Dataset: IndiaHO
Too low If
Too high NRMSD
Tf CI included 0
Armenian
Dataset: HO37
Too high Tf
Too high SE on Tf

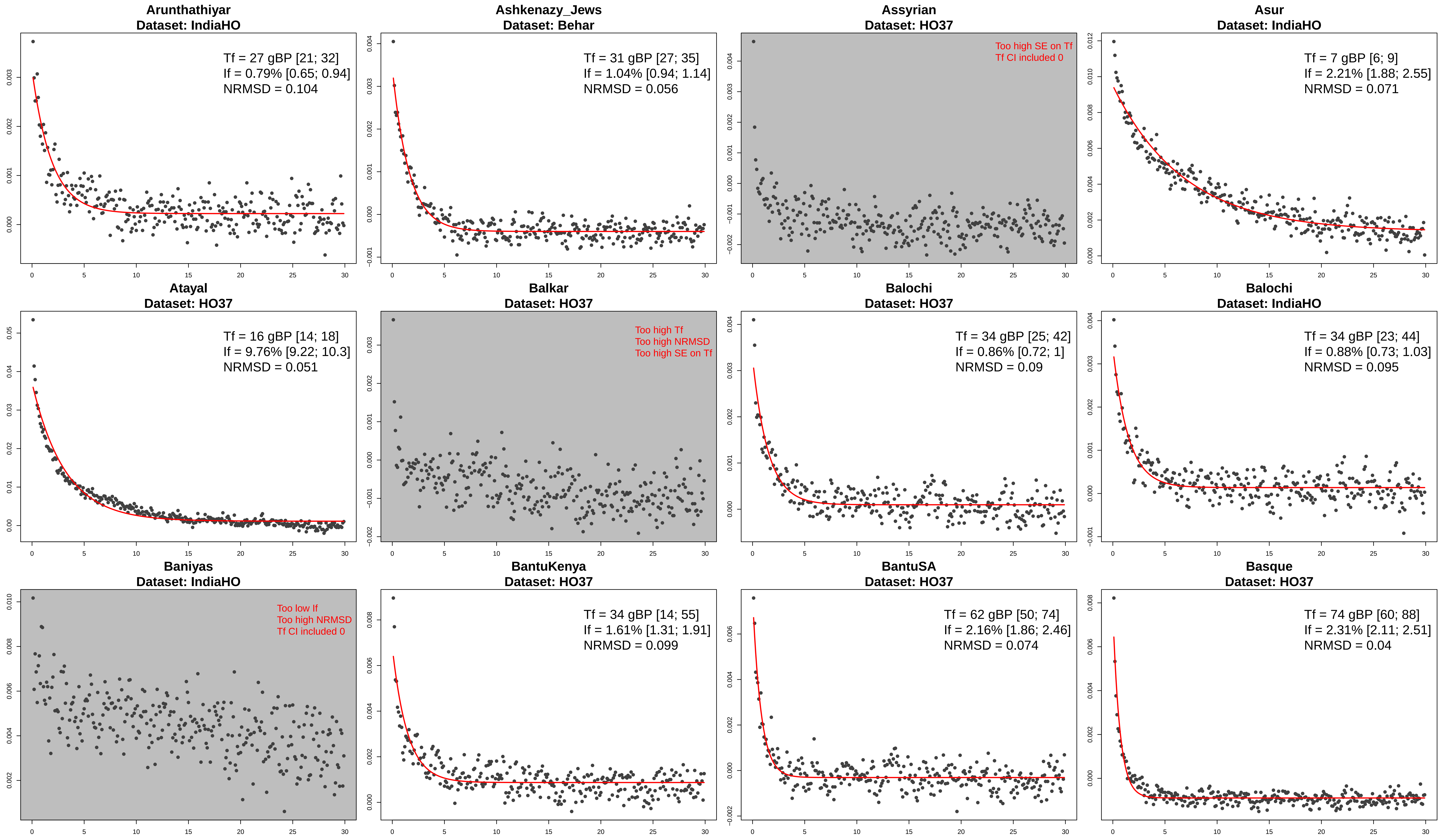
Arunthathiyar
Dataset: IndiaHO
Tf = 27 gBP [21; 32]
If = 0.79% [0.65; 0.94]
NRMSD = 0.104
Ashkenazy_Jews
Dataset: Behar
Tf = 31 gBP [27; 35]
If = 1.04% [0.94; 1.14]
NRMSD = 0.056
Assyrian
Dataset: HO37
Too high SE on Tf
Tf CI included 0
Asur
Dataset: IndiaHO
Tf = 7 gBP [6; 9]
If = 2.21% [1.88; 2.55]
NRMSD = 0.071
Atayal
Dataset: HO37
Tf = 16 gBP [14; 18]
If = 9.76% [9.22; 10.3]
NRMSD = 0.051
Balkar
Dataset: HO37
Too high Tf
Too high NRMSD
Too high SE on Tf
Balochi
Dataset: HO37
Tf = 34 gBP [25; 42]
If = 0.86% [0.72; 1]
NRMSD = 0.09
Balochi
Dataset: IndiaHO
Tf = 34 gBP [23; 44]
If = 0.88% [0.73; 1.03]
NRMSD = 0.095
Baniyas
Dataset: IndiaHO
Too low If
Too high NRMSD
Tf CI included 0
BantuKenya
Dataset: HO37
Tf = 34 gBP [14; 55]
If = 1.61% [1.31; 1.91]
NRMSD = 0.099
BantuSA
Dataset: HO37
Tf = 62 gBP [50; 74]
If = 2.16% [1.86; 2.46]
NRMSD = 0.074
Basque
Dataset: HO37
Tf = 74 gBP [60; 88]
If = 2.31% [2.11; 2.51]
NRMSD = 0.04

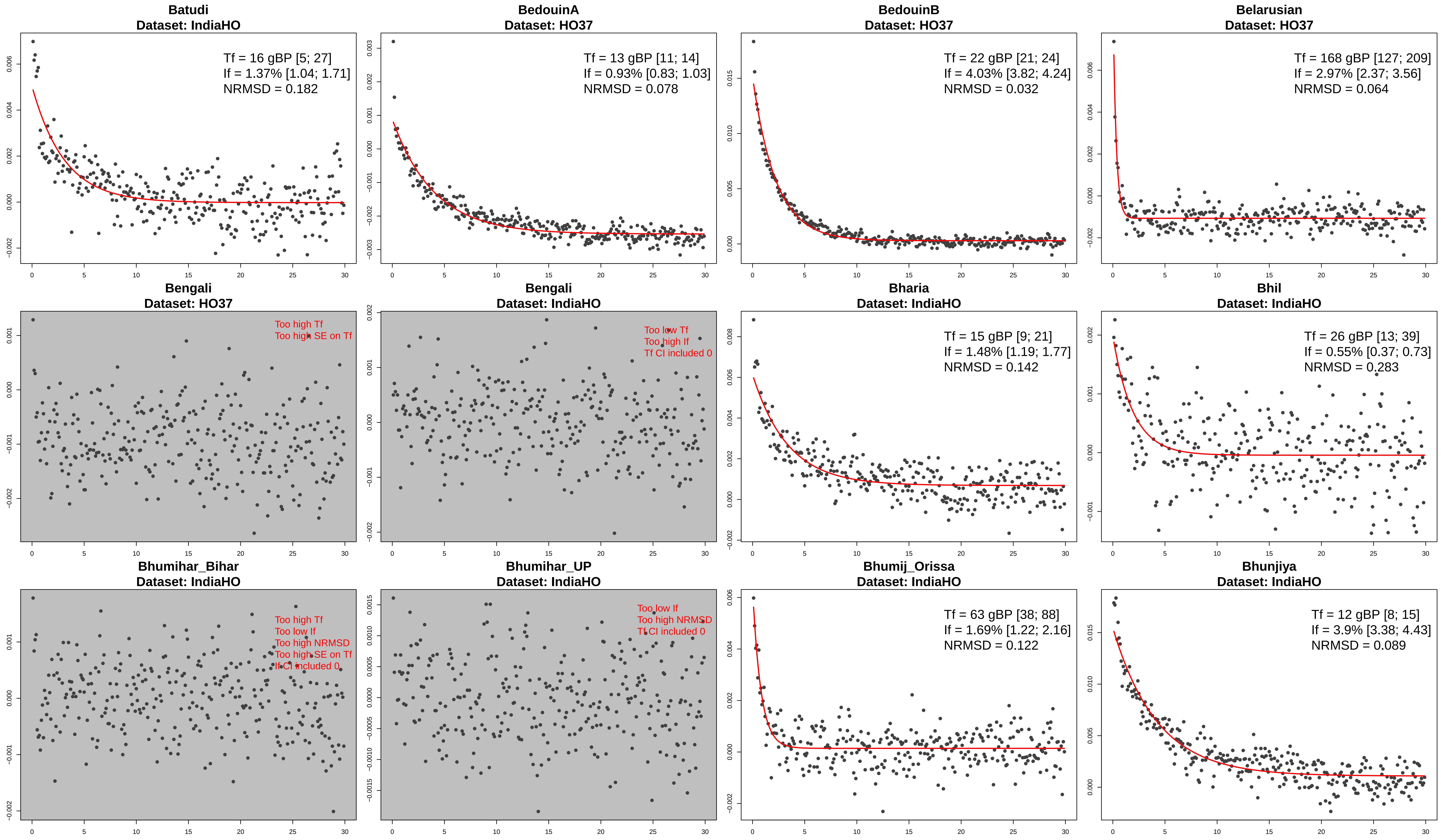
Batudi
Dataset: IndiaHO
Tf = 16 gBP [5; 27]
If = 1.37% [1.04; 1.71]
NRMSD = 0.182
BedouinA
Dataset: HO37
Tf = 13 gBP [11; 14]
If = 0.93% [0.83; 1.03]
NRMSD = 0.078
BedouinB
Dataset: HO37
Tf = 22 gBP [21; 24]
If = 4.03% [3.82; 4.24]
NRMSD = 0.032
Belarusian
Dataset: HO37
Tf = 168 gBP [127; 209]
If = 2.97% [2.37; 3.56]
NRMSD = 0.064
Bengali
Dataset: HO37
Too high Tf
Too high SE on Tf
Bengali
Dataset: IndiaHO
Too low Tf
Too high If
Tf CI included 0
Bharia
Dataset: IndiaHO
Tf = 15 gBP [9; 21]
If = 1.48% [1.19; 1.77]
NRMSD = 0.142
Bhil
Dataset: IndiaHO
Tf = 26 gBP [13; 39]
If = 0.55% [0.37; 0.73]
NRMSD = 0.283
Bhumihar_Bihar
Dataset: IndiaHO
Too high Tf
Too low If
Too high NRMSD
Too high SE on Tf
If CI included 0
Bhumihar_UP
Dataset: IndiaHO
Too low If
Too high NRMSD
Tf CI included 0
Bhumij_Orissa
Dataset: IndiaHO
Tf = 63 gBP [38; 88]
If = 1.69% [1.22; 2.16]
NRMSD = 0.122
Bhunjiya
Dataset: IndiaHO
Tf = 12 gBP [8; 15]
If = 3.9% [3.38; 4.43]
NRMSD = 0.089

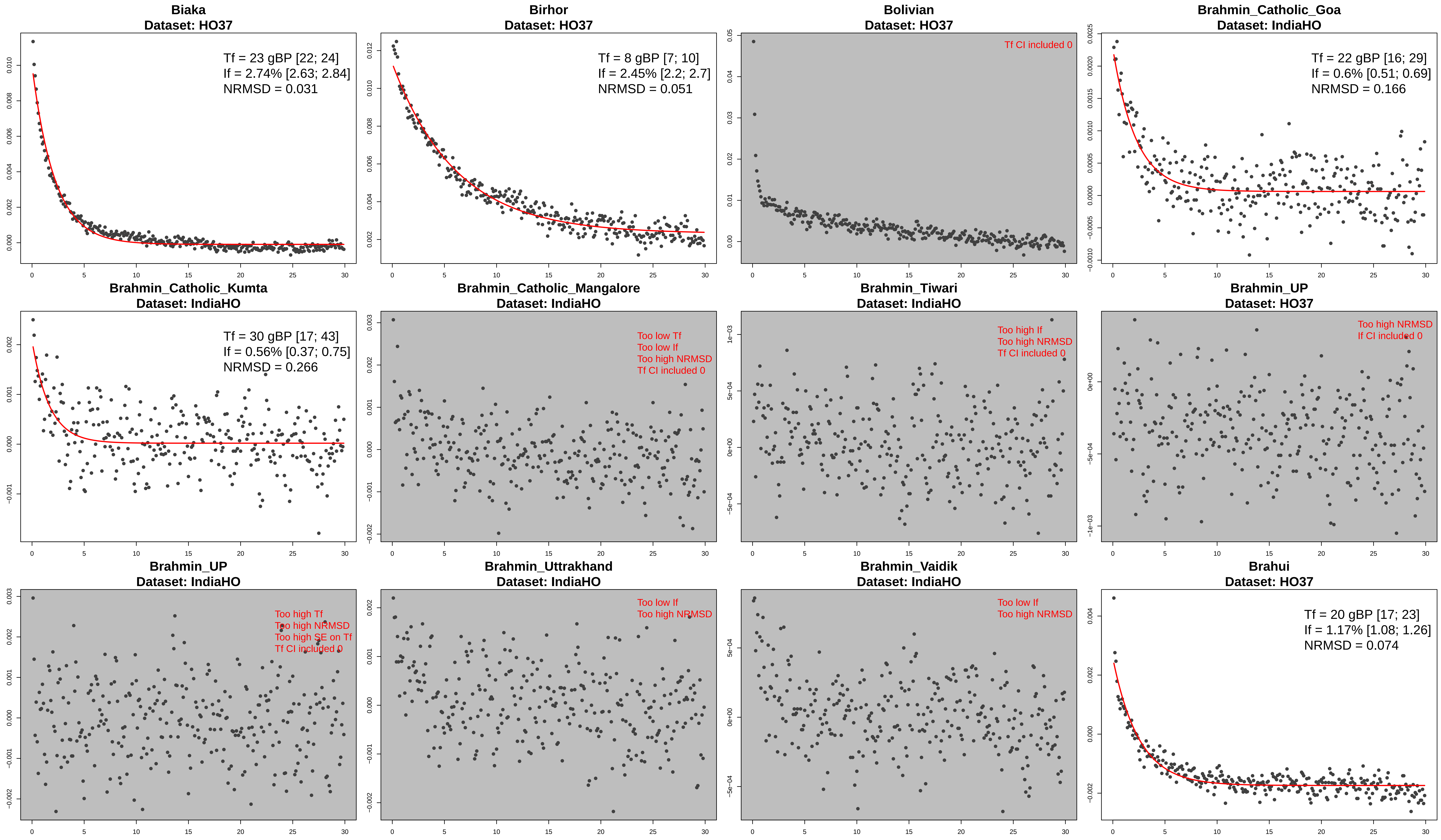

Biaka
Dataset: HO37
Tf = 23 gBP [22; 24]
If = 2.74% [2.63; 2.84]
NRMSD = 0.031
Birhor
Dataset: HO37
Tf = 8 gBP [7; 10]
If = 2.45% [2.2; 2.7]
NRMSD = 0.051
Bolivian
Dataset: HO37
Tf CI included 0
Brahmin_Catholic_Goa
Dataset: IndiaHO
Tf = 22 gBP [16; 29]
If = 0.6% [0.51; 0.69]
NRMSD = 0.166
Brahmin_Catholic_Kumta
Dataset: IndiaHO
Tf = 30 gBP [17; 43]
If = 0.56% [0.37; 0.75]
NRMSD = 0.266
Brahmin_Catholic_Mangalore
Dataset: IndiaHO
Too low Tf
Too low If
Too high NRMSD
Tf CI included 0
Brahmin_Tiwari
Dataset: IndiaHO
Too high If
Too high NRMSD
Tf CI included 0
Brahmin_UP
Dataset: HO37
Too high NRMSD
If CI included 0
Brahmin_UP
Dataset: IndiaHO
Too high Tf
Too high NRMSD
Too high SE on Tf
Tf CI included 0
Brahmin_Uttrakhand
Dataset: IndiaHO
Too low If
Too high NRMSD
Brahmin_Vaidik
Dataset: IndiaHO
Too low If
Too high NRMSD
Brahui
Dataset: HO37
Tf = 20 gBP [17; 23]
If = 1.17% [1.08; 1.26]
NRMSD = 0.074

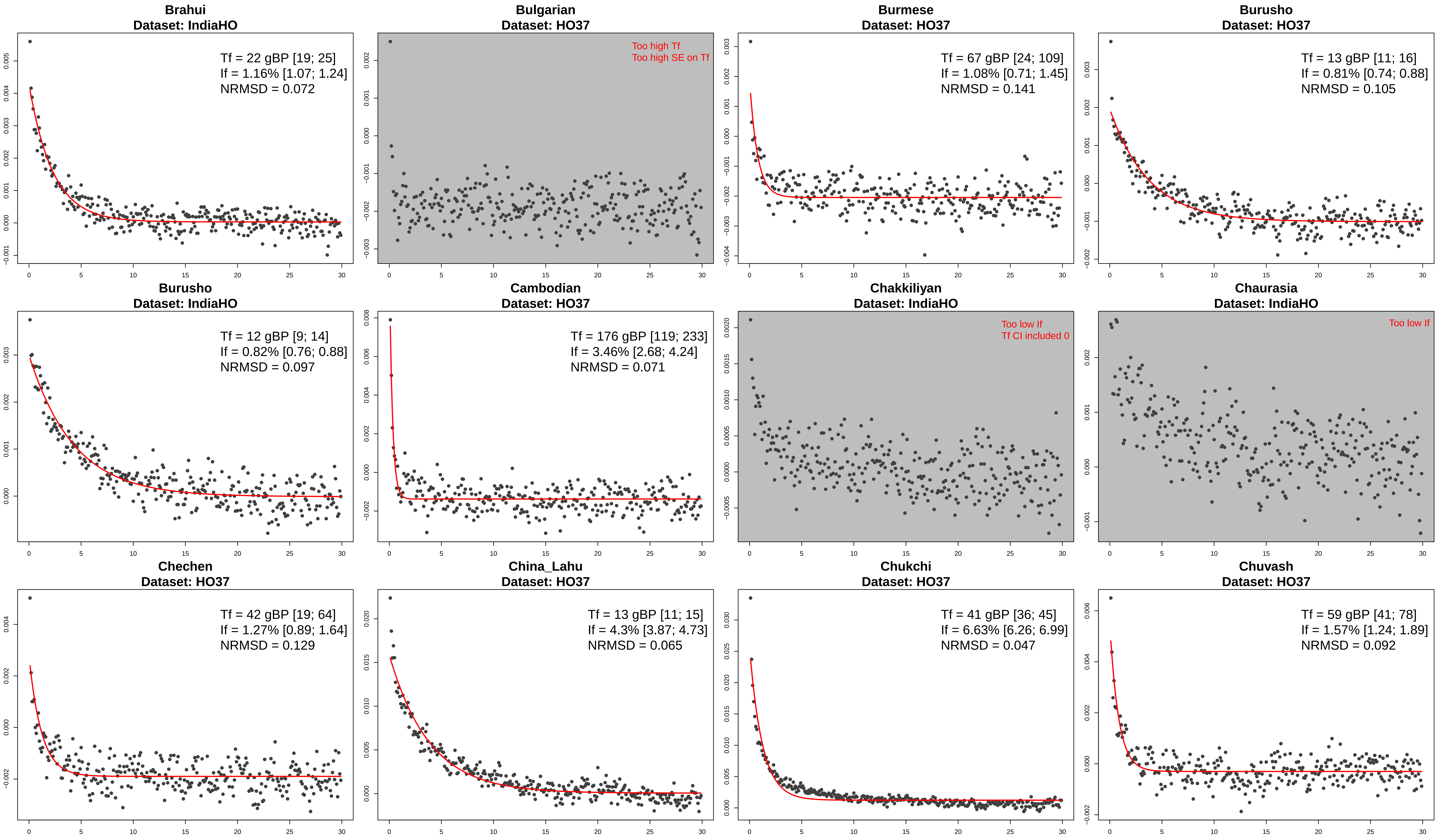
Brahui
Dataset: IndiaHO
Tf = 22 gBP [19; 25]
If = 1.16% [1.07; 1.24]
NRMSD = 0.072
Bulgarian
Dataset: HO37
Too high Tf
Too high SE on Tf
Burmese
Dataset: HO37
Tf = 67 gBP [24; 109]
If = 1.08% [0.71; 1.45]
NRMSD = 0.141
Burusho
Dataset: HO37
Tf = 13 gBP [11; 16]
If = 0.81% [0.74; 0.88]
NRMSD = 0.105
Burusho
Dataset: IndiaHO
Tf = 12 gBP [9; 14]
If = 0.82% [0.76; 0.88]
NRMSD = 0.097
Cambodian
Dataset: HO37
Tf = 176 gBP [119; 233]
If = 3.46% [2.68; 4.24]
NRMSD = 0.071
Chakkiliyan
Dataset: IndiaHO
Too low If
Tf CI included 0
Chaurasia
Dataset: IndiaHO
Too low If
Chechen
Dataset: HO37
Tf = 42 gBP [19; 64]
If = 1.27% [0.89; 1.64]
NRMSD = 0.129
China_Lahu
Dataset: HO37
Tf = 13 gBP [11; 15]
If = 4.3% [3.87; 4.73]
NRMSD = 0.065
Chukchi
Dataset: HO37
Tf = 41 gBP [36; 45]
If = 6.63% [6.26; 6.99]
NRMSD = 0.047
Chuvash
Dataset: HO37
Tf = 59 gBP [41; 78]
If = 1.57% [1.24; 1.89]
NRMSD = 0.092

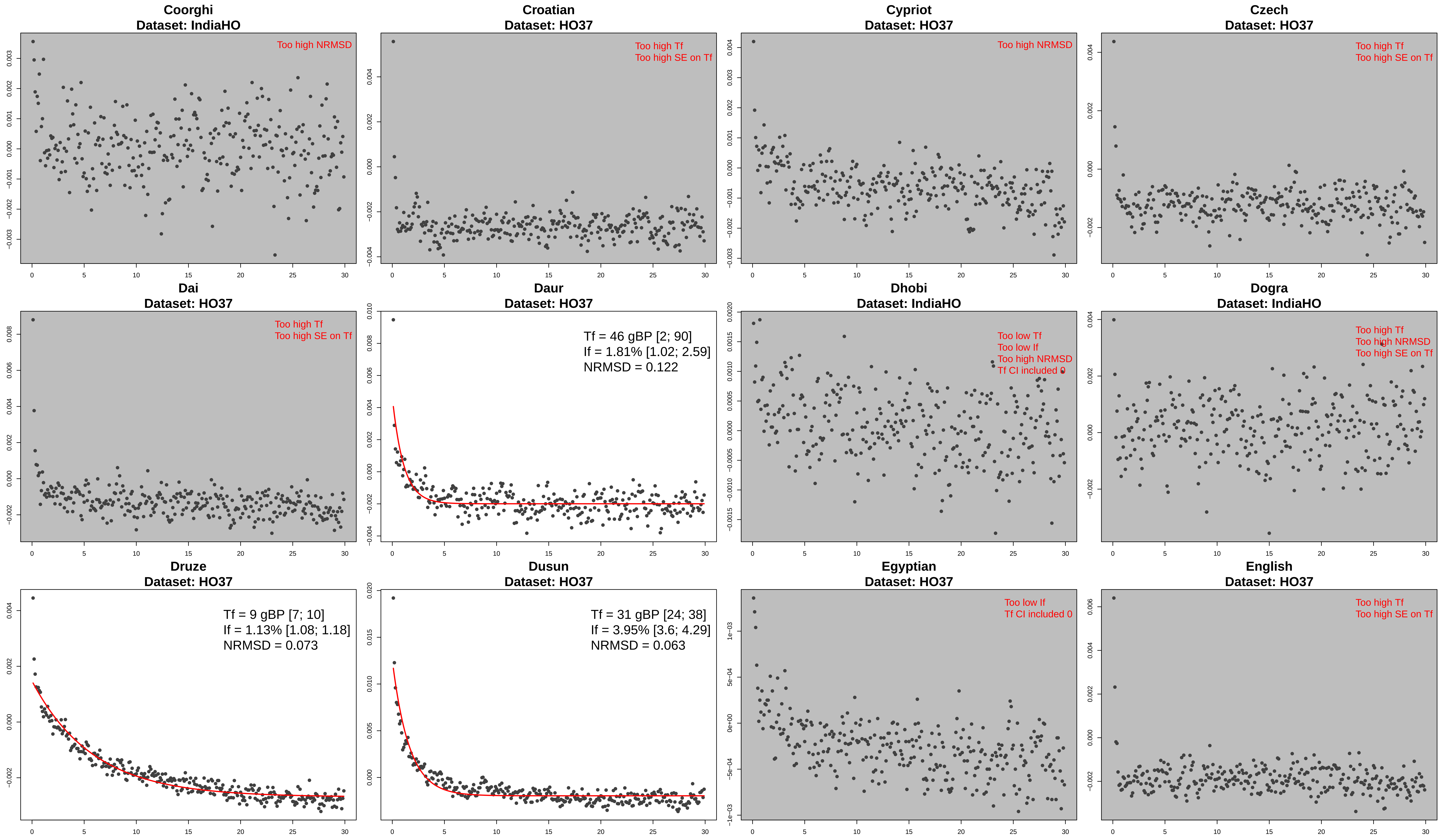

Coorghi
Dataset: IndiaHO
Too high NRMSD
Croatian
Dataset: HO37
Too high Tf
Too high SE on Tf
Cypriot
Dataset: HO37
Too high NRMSD
Czech
Dataset: HO37
Too high Tf
Too high SE on Tf
Dai
Dataset: HO37
Too high Tf
Too high SE on Tf
Daur
Dataset: HO37
Tf = 46 gBP [2; 90]
If = 1.81% [1.02; 2.59]
NRMSD = 0.122
Dhobi
Dataset: IndiaHO
Too low Tf
Too low If
Too high NRMSD
Tf CI included 0
Dogra
Dataset: IndiaHO
Too high Tf
Too high NRMSD
Too high SE on Tf
Druze
Dataset: HO37
Tf = 9 gBP [7; 10]
If = 1.13% [1.08; 1.18]
NRMSD = 0.073
Dusun
Dataset: HO37
Tf = 31 gBP [24; 38]
If = 3.95% [3.6; 4.29]
NRMSD = 0.063
Egyptian
Dataset: HO37
Too low If
Tf CI included 0
English
Dataset: HO37
Too high Tf
Too high SE on Tf

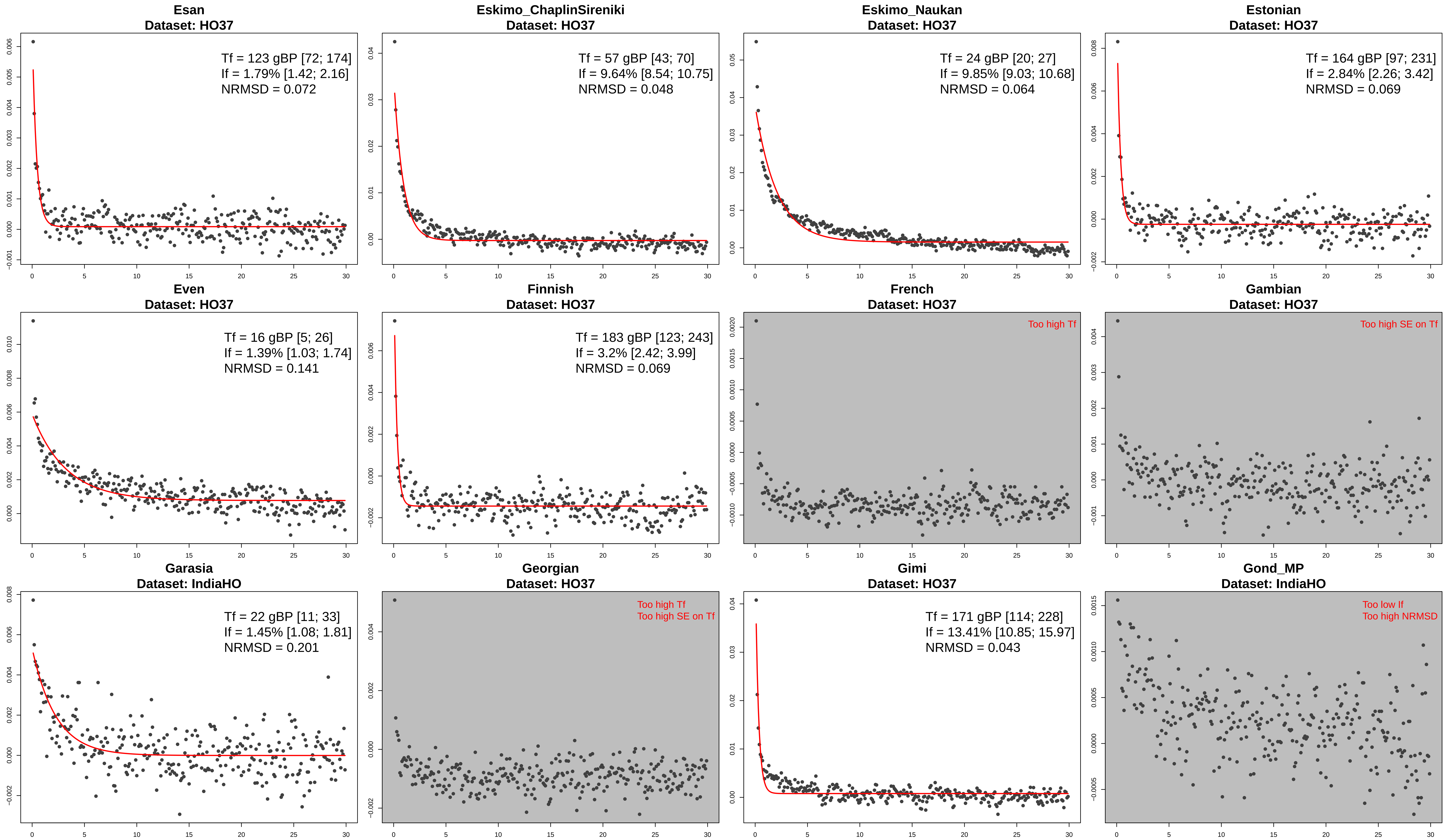
Esan
Dataset: HO37
Tf = 123 gBP [72; 174]
If = 1.79% [1.42; 2.16]
NRMSD = 0.072
Eskimo_ChaplinSireniki
Dataset: HO37
Tf = 57 gBP [43; 70]
If = 9.64% [8.54; 10.75]
NRMSD = 0.048
Eskimo_Naukan
Dataset: HO37
Tf = 24 gBP [20; 27]
If = 9.85% [9.03; 10.68]
NRMSD = 0.064
Estonian
Dataset: HO37
Tf = 164 gBP [97; 231]
If = 2.84% [2.26; 3.42]
NRMSD = 0.069
Even
Dataset: HO37
Tf = 16 gBP [5; 26]
If = 1.39% [1.03; 1.74]
NRMSD = 0.141
Finnish
Dataset: HO37
Tf = 183 gBP [123; 243]
If = 3.2% [2.42; 3.99]
NRMSD = 0.069
French
Dataset: HO37
Too high Tf
Gambian
Dataset: HO37
Too high SE on Tf
Garasia
Dataset: IndiaHO
Tf = 22 gBP [11; 33]
If = 1.45% [1.08; 1.81]
NRMSD = 0.201
Georgian
Dataset: HO37
Too high Tf
Too high SE on Tf
Gimi
Dataset: HO37
Tf = 171 gBP [114; 228]
If = 13.41% [10.85; 15.97]
NRMSD = 0.043
Gond_MP
Dataset: IndiaHO
Too low If
Too high NRMSD

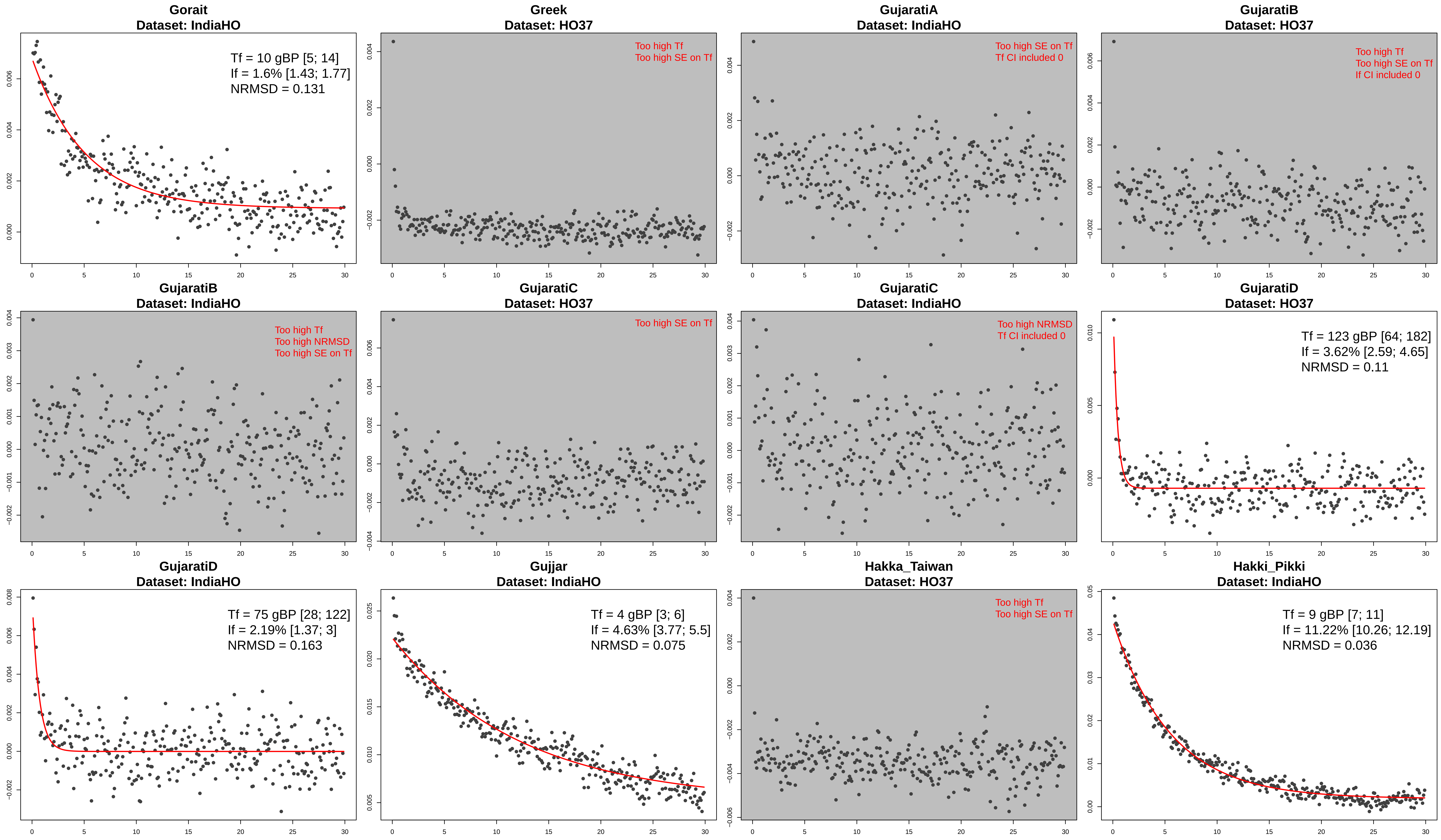
Gorait
Dataset: IndiaHO
Tf = 10 gBP [5; 14]
If = 1.6% [1.43; 1.77]
NRMSD = 0.131
Greek
Dataset: HO37
Too high Tf
Too high SE on Tf
GujaratiA
Dataset: IndiaHO
Too high SE on Tf
Tf CI included 0
GujaratiB
Dataset: HO37
Too high Tf
Too high SE on Tf
If CI included 0
GujaratiB
Dataset: IndiaHO
Too high Tf
Too high NRMSD
Too high SE on Tf
GujaratiC
Dataset: HO37
Too high SE on Tf
GujaratiC
Dataset: IndiaHO
Too high NRMSD
Tf CI included 0
GujaratiD
Dataset: HO37
Tf = 123 gBP [64; 182]
If = 3.62% [2.59; 4.65]
NRMSD = 0.11
GujaratiD
Dataset: IndiaHO
Tf = 75 gBP [28; 122]
If = 2.19% [1.37; 3]
NRMSD = 0.163
Gujjar
Dataset: IndiaHO
Tf = 4 gBP [3; 6]
If = 4.63% [3.77; 5.5]
NRMSD = 0.075
Hakka_Taiwan
Dataset: HO37
Too high Tf
Too high SE on Tf
Hakki_Pikki
Dataset: IndiaHO
Tf = 9 gBP [7; 11]
If = 11.22% [10.26; 12.19]
NRMSD = 0.036

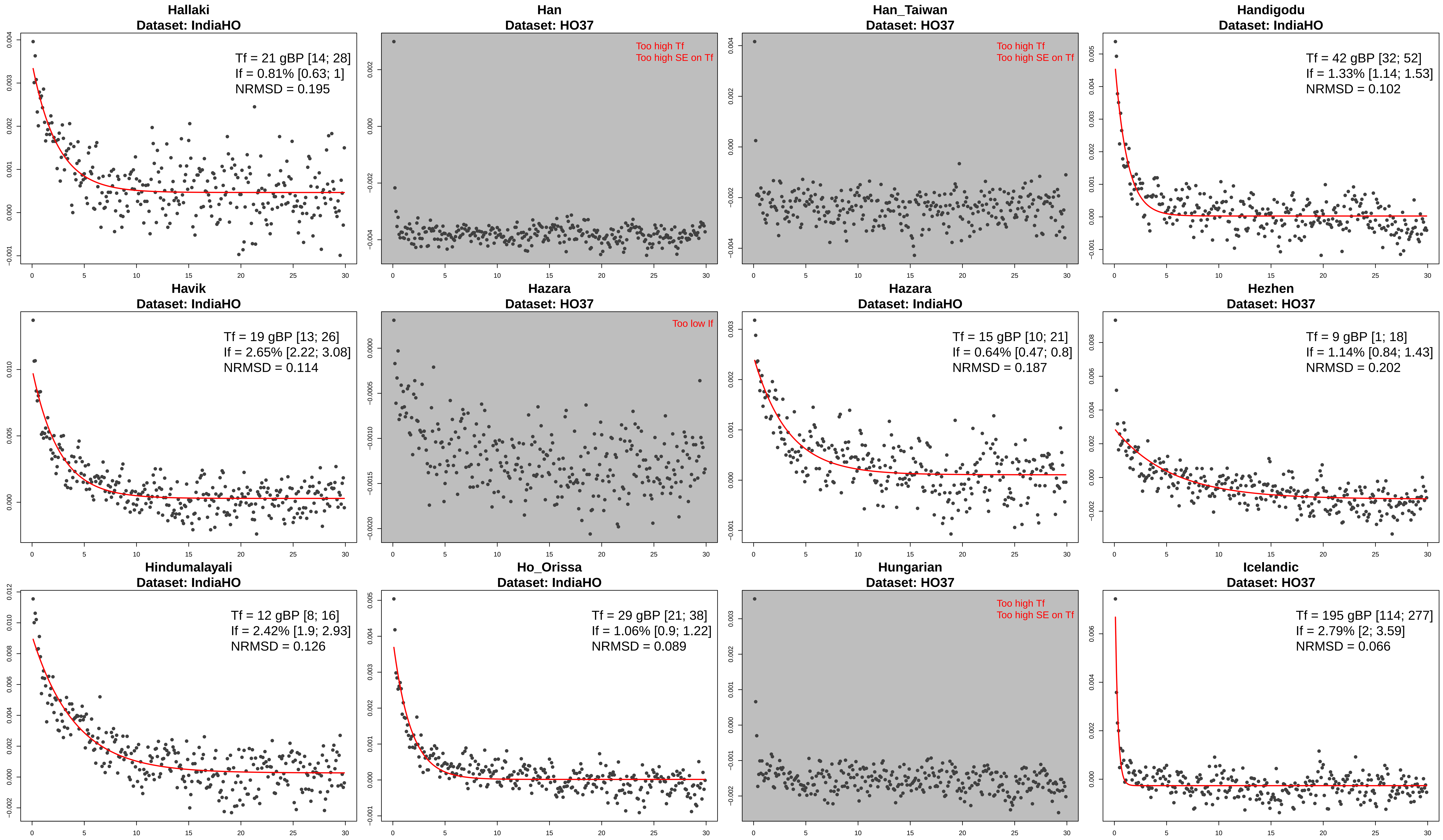
Hallaki
Dataset: IndiaHO
Tf = 21 gBP [14; 28]
If = 0.81% [0.63; 1]
NRMSD = 0.195
Han
Dataset: HO37
Too high Tf
Too high SE on Tf
Han_Taiwan
Dataset: HO37
Too high Tf
Too high SE on Tf
Handigodu
Dataset: IndiaHO
Tf = 42 gBP [32; 52]
If = 1.33% [1.14; 1.53]
NRMSD = 0.102
Havik
Dataset: IndiaHO
Tf = 19 gBP [13; 26]
If = 2.65% [2.22; 3.08]
NRMSD = 0.114
Hazara
Dataset: HO37
Too low If
Hazara
Dataset: IndiaHO
Tf = 15 gBP [10; 21]
If = 0.64% [0.47; 0.8]
NRMSD = 0.187
Hezhen
Dataset: HO37
Tf = 9 gBP [1; 18]
If = 1.14% [0.84; 1.43]
NRMSD = 0.202
Hindumalayali
Dataset: IndiaHO
Tf = 12 gBP [8; 16]
If = 2.42% [1.9; 2.93]
NRMSD = 0.126
Ho_Orissa
Dataset: IndiaHO
Tf = 29 gBP [21; 38]
If = 1.06% [0.9; 1.22]
NRMSD = 0.089
Hungarian
Dataset: HO37
Too high Tf
Too high SE on Tf
Icelandic
Dataset: HO37
Tf = 195 gBP [114; 277]
If = 2.79% [2; 3.59]
NRMSD = 0.066

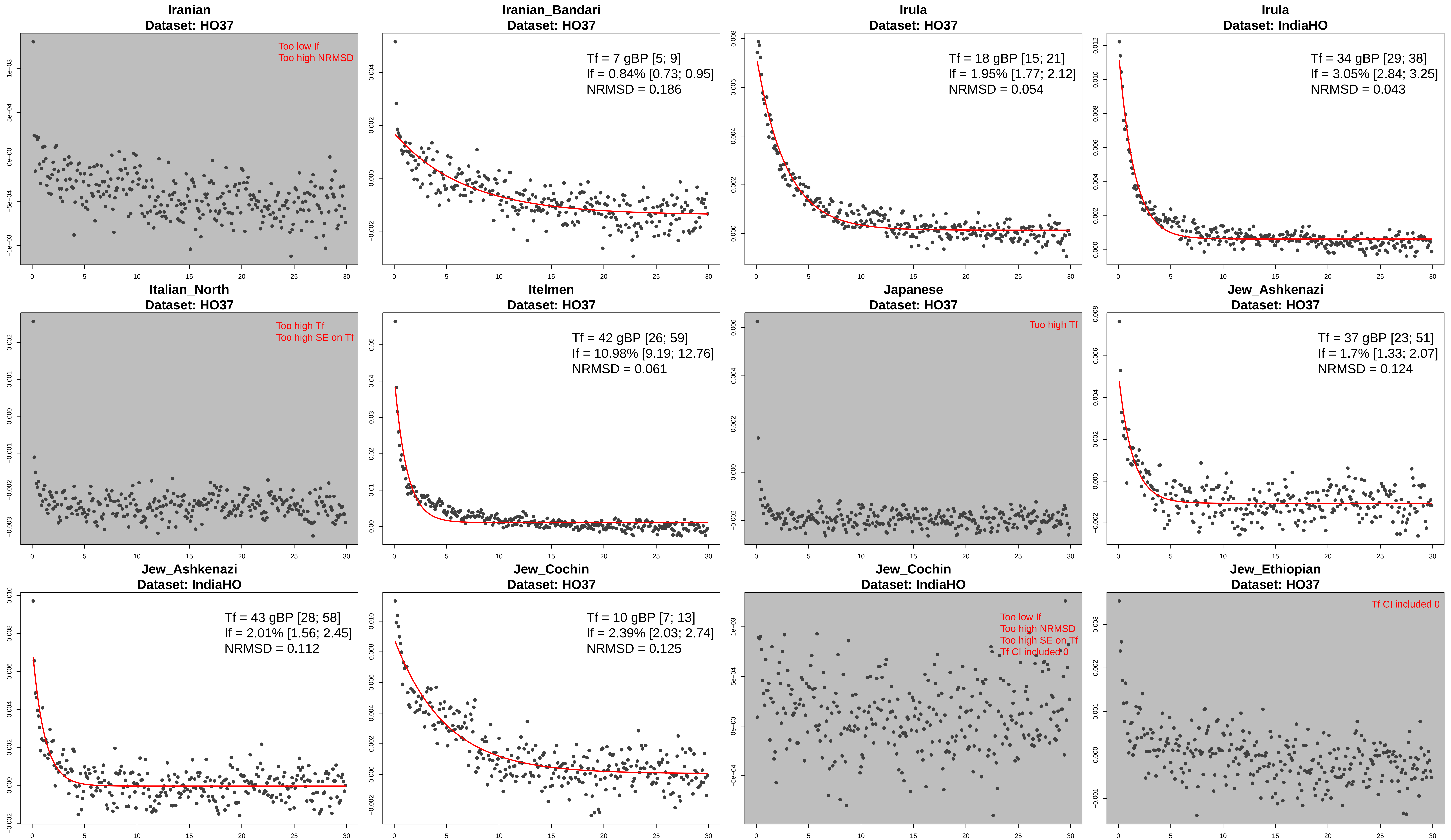
Iranian
Dataset: HO37
Too low If
Too high NRMSD
Iranian_Bandari
Dataset: HO37
Tf = 7 gBP [5; 9]
If = 0.84% [0.73; 0.95]
NRMSD = 0.186
Irula
Dataset: HO37
Tf = 18 gBP [15; 21]
If = 1.95% [1.77; 2.12]
NRMSD = 0.054
Irula
Dataset: IndiaHO
Tf = 34 gBP [29; 38]
If = 3.05% [2.84; 3.25]
NRMSD = 0.043
Italian_North
Dataset: HO37
Too high Tf
Too high SE on Tf
Itelmen
Dataset: HO37
Tf = 42 gBP [26; 59]
If = 10.98% [9.19; 12.76]
NRMSD = 0.061
Japanese
Dataset: HO37
Too high Tf
Jew_Ashkenazi
Dataset: HO37
Tf = 37 gBP [23; 51]
If = 1.7% [1.33; 2.07]
NRMSD = 0.124
Jew_Ashkenazi
Dataset: IndiaHO
Tf = 43 gBP [28; 58]
If = 2.01% [1.56; 2.45]
NRMSD = 0.112
Jew_Cochin
Dataset: HO37
Tf = 10 gBP [7; 13]
If = 2.39% [2.03; 2.74]
NRMSD = 0.125
Jew_Cochin
Dataset: IndiaHO
Too low If
Too high NRMSD
Too high SE on Tf
Tf CI included 0
Jew_Ethiopian
Dataset: HO37
Tf CI included 0

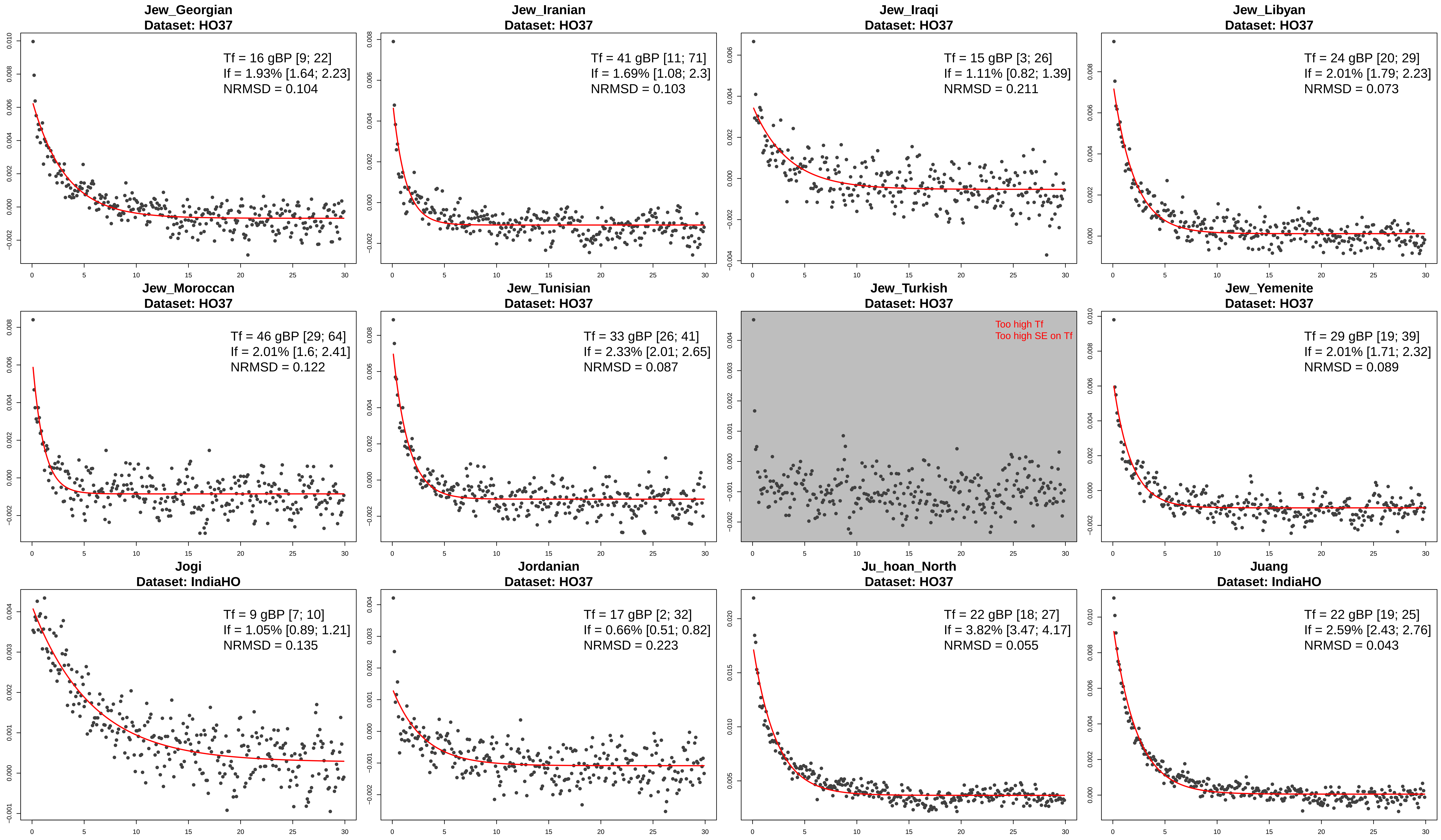
Jew_Georgian
Dataset: HO37
Tf = 16 gBP [9; 22]
If = 1.93% [1.64; 2.23]
NRMSD = 0.104
Jew_Iranian
Dataset: HO37
Tf = 41 gBP [11; 71]
If = 1.69% [1.08; 2.3]
NRMSD = 0.103
Jew_Iraqi
Dataset: HO37
Tf = 15 gBP [3; 26]
If = 1.11% [0.82; 1.39]
NRMSD = 0.211
Jew_Libyan
Dataset: HO37
Tf = 24 gBP [20; 29]
If = 2.01% [1.79; 2.23]
NRMSD = 0.073
Jew_Moroccan
Dataset: HO37
Tf = 46 gBP [29; 64]
If = 2.01% [1.6; 2.41]
NRMSD = 0.122
Jew_Tunisian
Dataset: HO37
Tf = 33 gBP [26; 41]
If = 2.33% [2.01; 2.65]
NRMSD = 0.087
Jew_Turkish
Dataset: HO37
Too high Tf
Too high SE on Tf
Jew_Yemenite
Dataset: HO37
Tf = 29 gBP [19; 39]
If = 2.01% [1.71; 2.32]
NRMSD = 0.089
Jogi
Dataset: IndiaHO
Tf = 9 gBP [7; 10]
If = 1.05% [0.89; 1.21]
NRMSD = 0.135
Jordanian
Dataset: HO37
Tf = 17 gBP [2; 32]
If = 0.66% [0.51; 0.82]
NRMSD = 0.223
Ju_hoan_North
Dataset: HO37
Tf = 22 gBP [18; 27]
If = 3.82% [3.47; 4.17]
NRMSD = 0.055
Juang
Dataset: IndiaHO
Tf = 22 gBP [19; 25]
If = 2.59% [2.43; 2.76]
NRMSD = 0.043

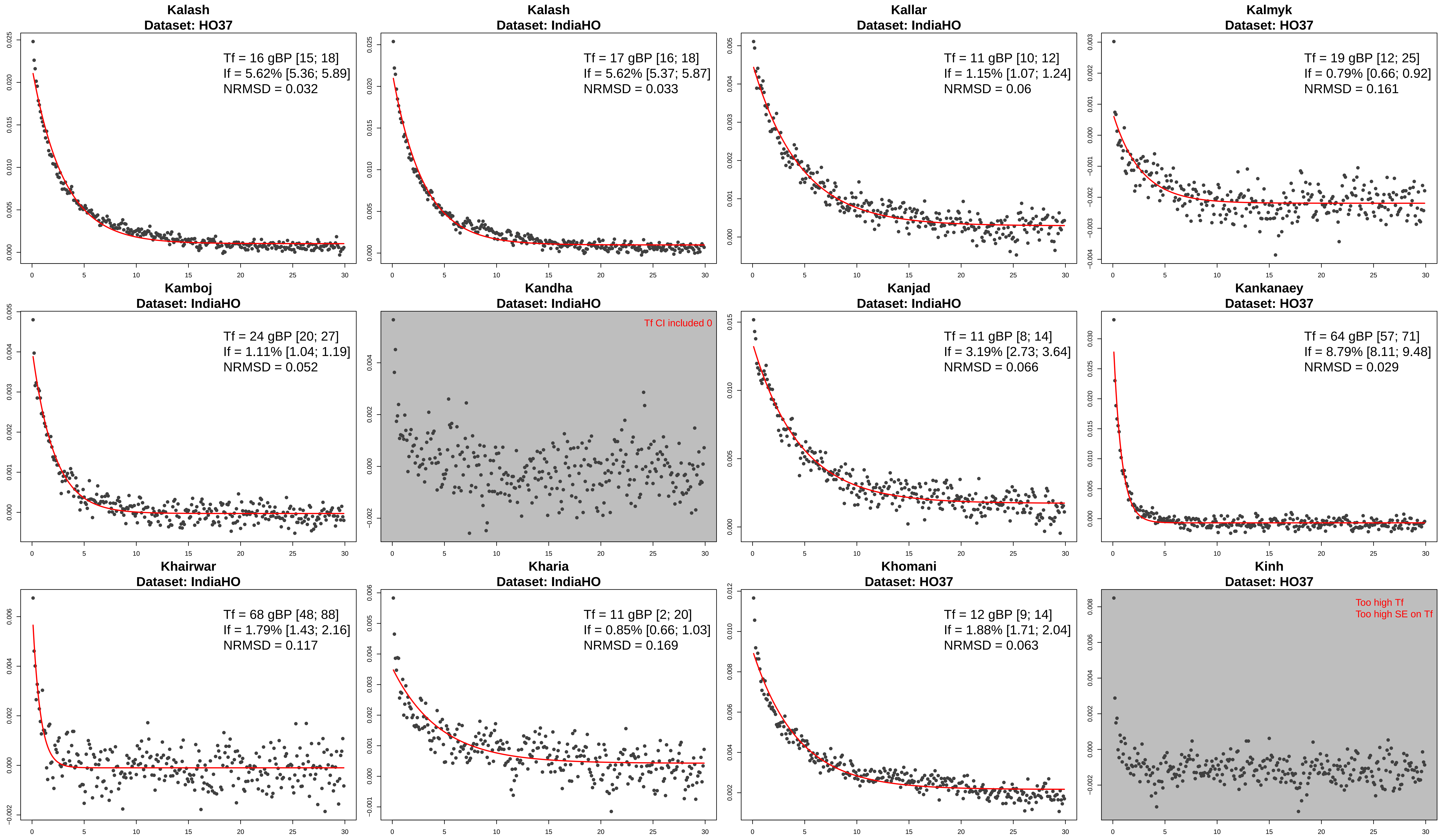
Kalash
Dataset: HO37
Tf = 16 gBP [15; 18]
If = 5.62% [5.36; 5.89]
NRMSD = 0.032
Kalash
Dataset: IndiaHO
Tf = 17 gBP [16; 18]
If = 5.62% [5.37; 5.87]
NRMSD = 0.033
Kallar
Dataset: IndiaHO
Tf = 11 gBP [10; 12]
If = 1.15% [1.07; 1.24]
NRMSD = 0.06
Kalmyk
Dataset: HO37
Tf = 19 gBP [12; 25]
If = 0.79% [0.66; 0.92]
NRMSD = 0.161
Kamboj
Dataset: IndiaHO
Tf = 24 gBP [20; 27]
If = 1.11% [1.04; 1.19]
NRMSD = 0.052
Kandha
Dataset: IndiaHO
Tf CI included 0
Kanjad
Dataset: IndiaHO
Tf = 11 gBP [8; 14]
If = 3.19% [2.73; 3.64]
NRMSD = 0.066
Kankanaey
Dataset: HO37
Tf = 64 gBP [57; 71]
If = 8.79% [8.11; 9.48]
NRMSD = 0.029
Khairwar
Dataset: IndiaHO
Tf = 68 gBP [48; 88]
If = 1.79% [1.43; 2.16]
NRMSD = 0.117
Kharia
Dataset: IndiaHO
Tf = 11 gBP [2; 20]
If = 0.85% [0.66; 1.03]
NRMSD = 0.169
Khomani
Dataset: HO37
Tf = 12 gBP [9; 14]
If = 1.88% [1.71; 2.04]
NRMSD = 0.063
Kinh
Dataset: HO37
Too high Tf
Too high SE on Tf

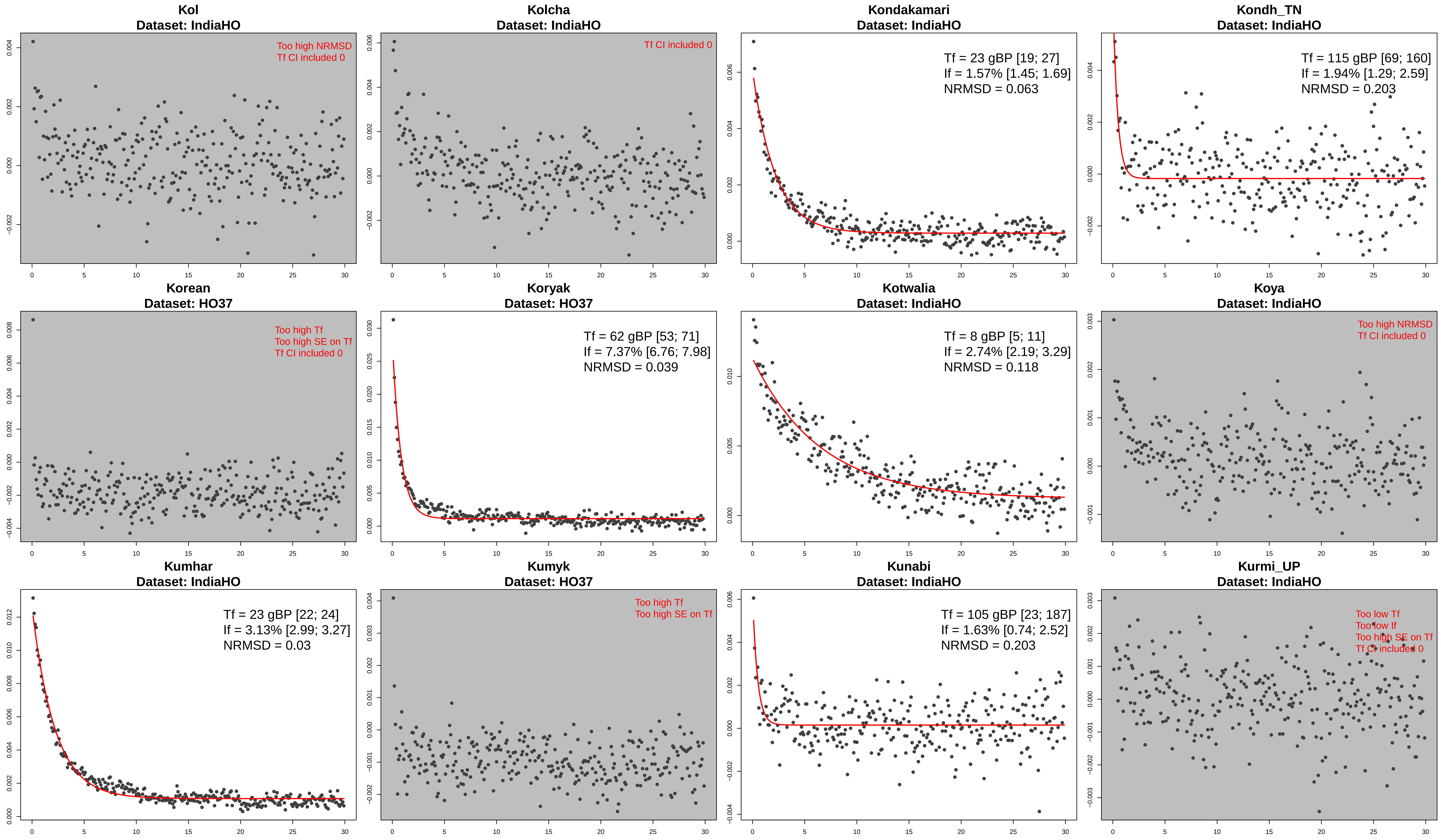
Kol
Dataset: IndiaHO
Too high NRMSD
Tf CI included 0
Kolcha
Dataset: IndiaHO
Tf CI included 0
Kondakamari
Dataset: IndiaHO
Tf = 23 gBP [19; 27]
If = 1.57% [1.45; 1.69]
NRMSD = 0.063
Kondh_TN
Dataset: IndiaHO
Tf = 115 gBP [69; 160]
If = 1.94% [1.29; 2.59]
NRMSD = 0.203
Korean
Dataset: HO37
Too high Tf
Too high SE on Tf
Tf CI included 0
Koryak
Dataset: HO37
Tf = 62 gBP [53; 71]
If = 7.37% [6.76; 7.98]
NRMSD = 0.039
Kotwalia
Dataset: IndiaHO
Tf = 8 gBP [5; 11]
If = 2.74% [2.19; 3.29]
NRMSD = 0.118
Koya
Dataset: IndiaHO
Too high NRMSD
Tf CI included 0
Kumhar
Dataset: IndiaHO
Tf = 23 gBP [22; 24]
If = 3.13% [2.99; 3.27]
NRMSD = 0.03
Kumyk
Dataset: HO37
Too high Tf
Too high SE on Tf
Kunabi
Dataset: IndiaHO
Tf = 105 gBP [23; 187]
If = 1.63% [0.74; 2.52]
NRMSD = 0.203
Kurmi_UP
Dataset: IndiaHO
Too low Tf
Too low If
Too high SE on Tf
Tf CI included 0

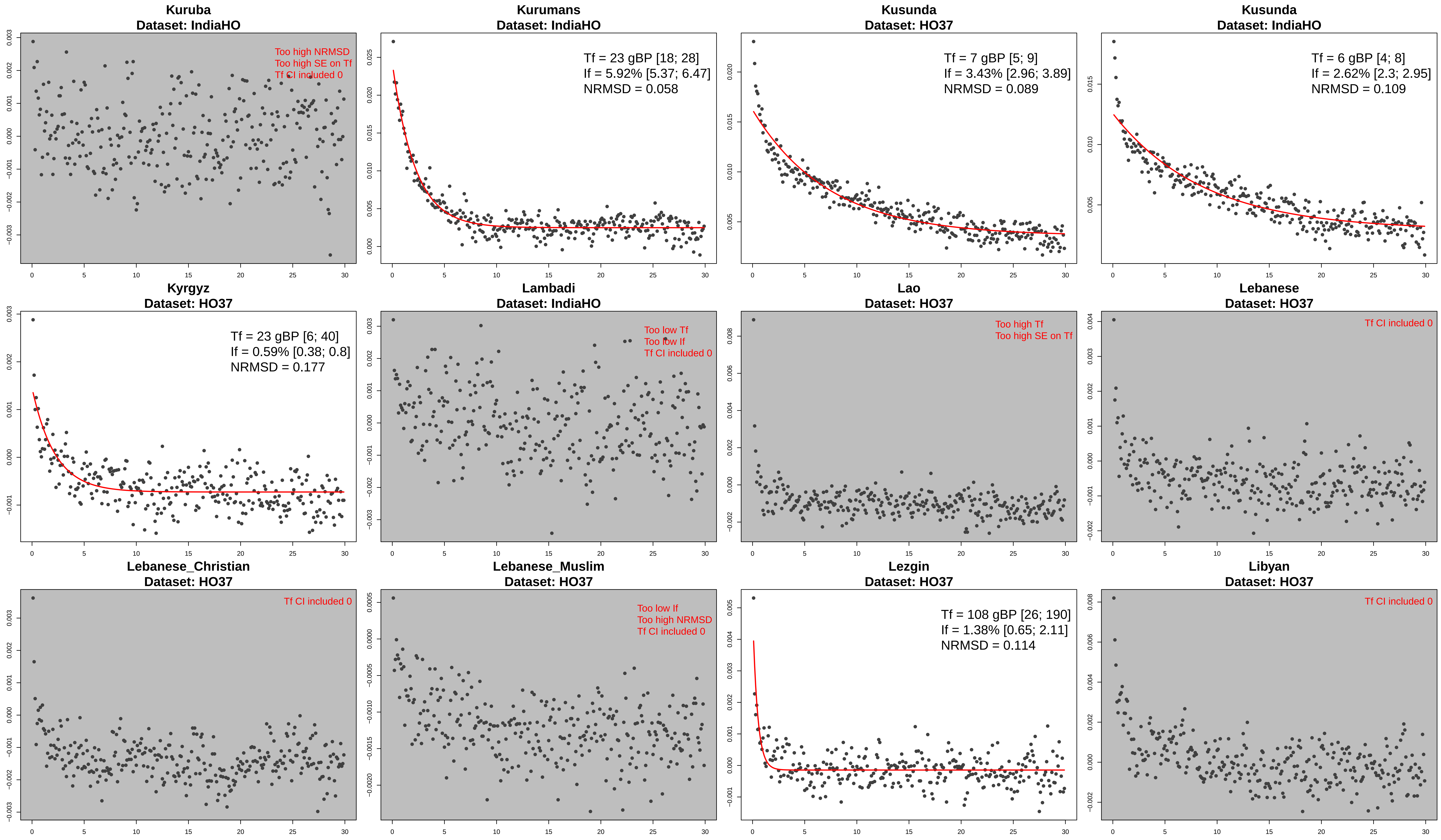
Kuruba
Dataset: IndiaHO
Too high NRMSD
Too high SE on Tf
Tf CI included 0
Kurumans
Dataset: IndiaHO
Tf = 23 gBP [18; 28]
If = 5.92% [5.37; 6.47]
NRMSD = 0.058
Kusunda
Dataset: HO37
Tf = 7 gBP [5; 9]
If = 3.43% [2.96; 3.89]
NRMSD = 0.089
Kusunda
Dataset: IndiaHO
Tf = 6 gBP [4; 8]
If = 2.62% [2.3; 2.95]
NRMSD = 0.109
Kyrgyz
Dataset: HO37
Tf = 23 gBP [6; 40]
If = 0.59% [0.38; 0.8]
NRMSD = 0.177
Lambadi
Dataset: IndiaHO
Too low Tf
Too low If
Tf CI included 0
Lao
Dataset: HO37
Too high Tf
Too high SE on Tf
Lebanese
Dataset: HO37
Tf CI included 0
Lebanese_Christian
Dataset: HO37
Tf CI included 0
Lebanese_Muslim
Dataset: HO37
Too low If
Too high NRMSD
Tf CI included 0
Lezgin
Dataset: HO37
Tf = 108 gBP [26; 190]
If = 1.38% [0.65; 2.11]
NRMSD = 0.114
Libyan
Dataset: HO37
Tf CI included 0

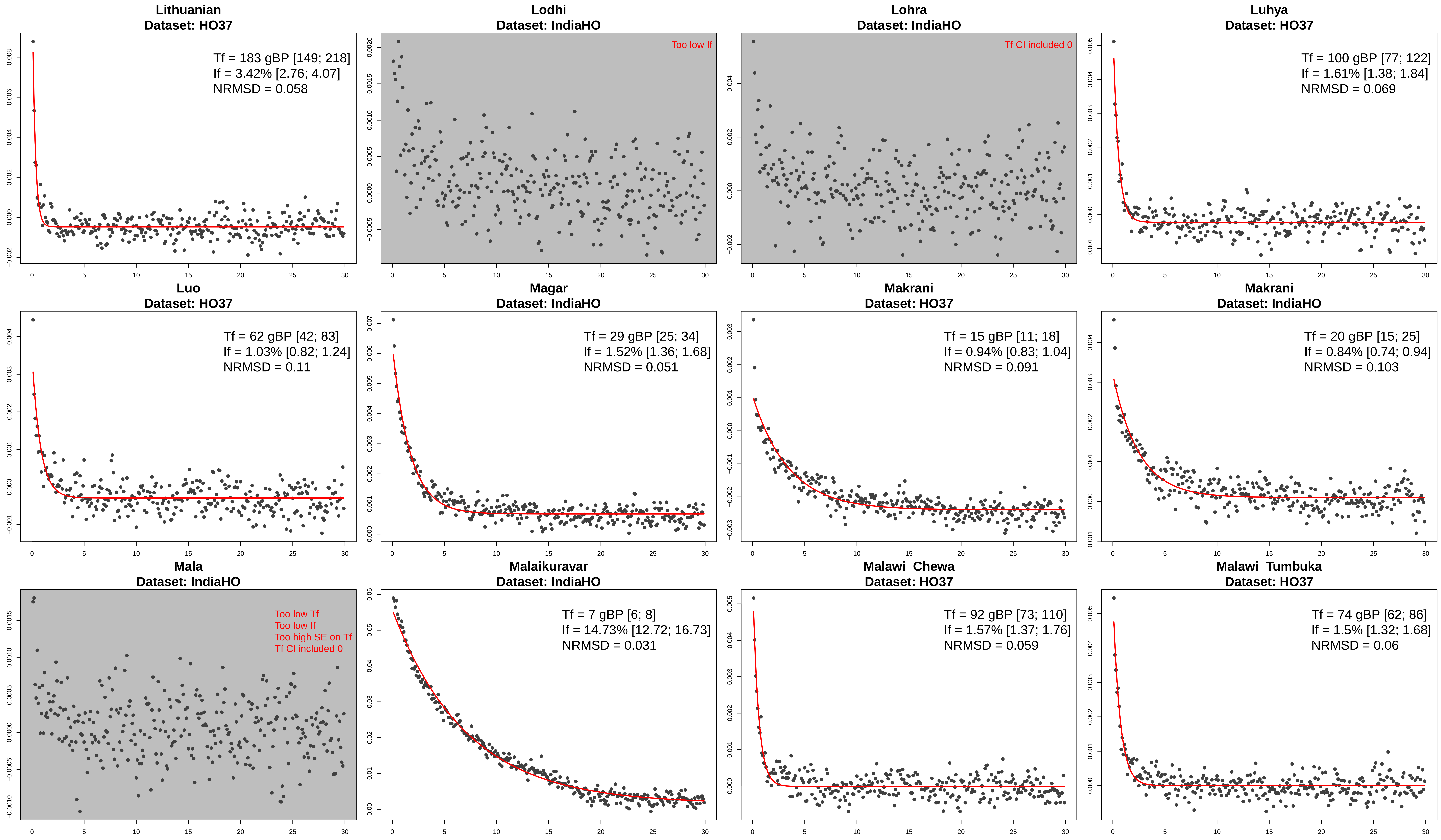
Lithuanian
Dataset: HO37
Tf = 183 gBP [149; 218]
If = 3.42% [2.76; 4.07]
NRMSD = 0.058
Lodhi
Dataset: IndiaHO
Too low If
Lohra
Dataset: IndiaHO
Tf CI included 0
Luhya
Dataset: HO37
Tf = 100 gBP [77; 122]
If = 1.61% [1.38; 1.84]
NRMSD = 0.069
Luo
Dataset: HO37
Tf = 62 gBP [42; 83]
If = 1.03% [0.82; 1.24]
NRMSD = 0.11
Magar
Dataset: IndiaHO
Tf = 29 gBP [25; 34]
If = 1.52% [1.36; 1.68]
NRMSD = 0.051
Makrani
Dataset: HO37
Tf = 15 gBP [11; 18]
If = 0.94% [0.83; 1.04]
NRMSD = 0.091
Makrani
Dataset: IndiaHO
Tf = 20 gBP [15; 25]
If = 0.84% [0.74; 0.94]
NRMSD = 0.103
Mala
Dataset: IndiaHO
Too low Tf
Too low If
Too high SE on Tf
Tf CI included 0
Malaikuravar
Dataset: IndiaHO
Tf = 7 gBP [6; 8]
If = 14.73% [12.72; 16.73]
NRMSD = 0.031
Malawi_Chewa
Dataset: HO37
Tf = 92 gBP [73; 110]
If = 1.57% [1.37; 1.76]
NRMSD = 0.059
Malawi_Tumbuka
Dataset: HO37
Tf = 74 gBP [62; 86]
If = 1.5% [1.32; 1.68]
NRMSD = 0.06

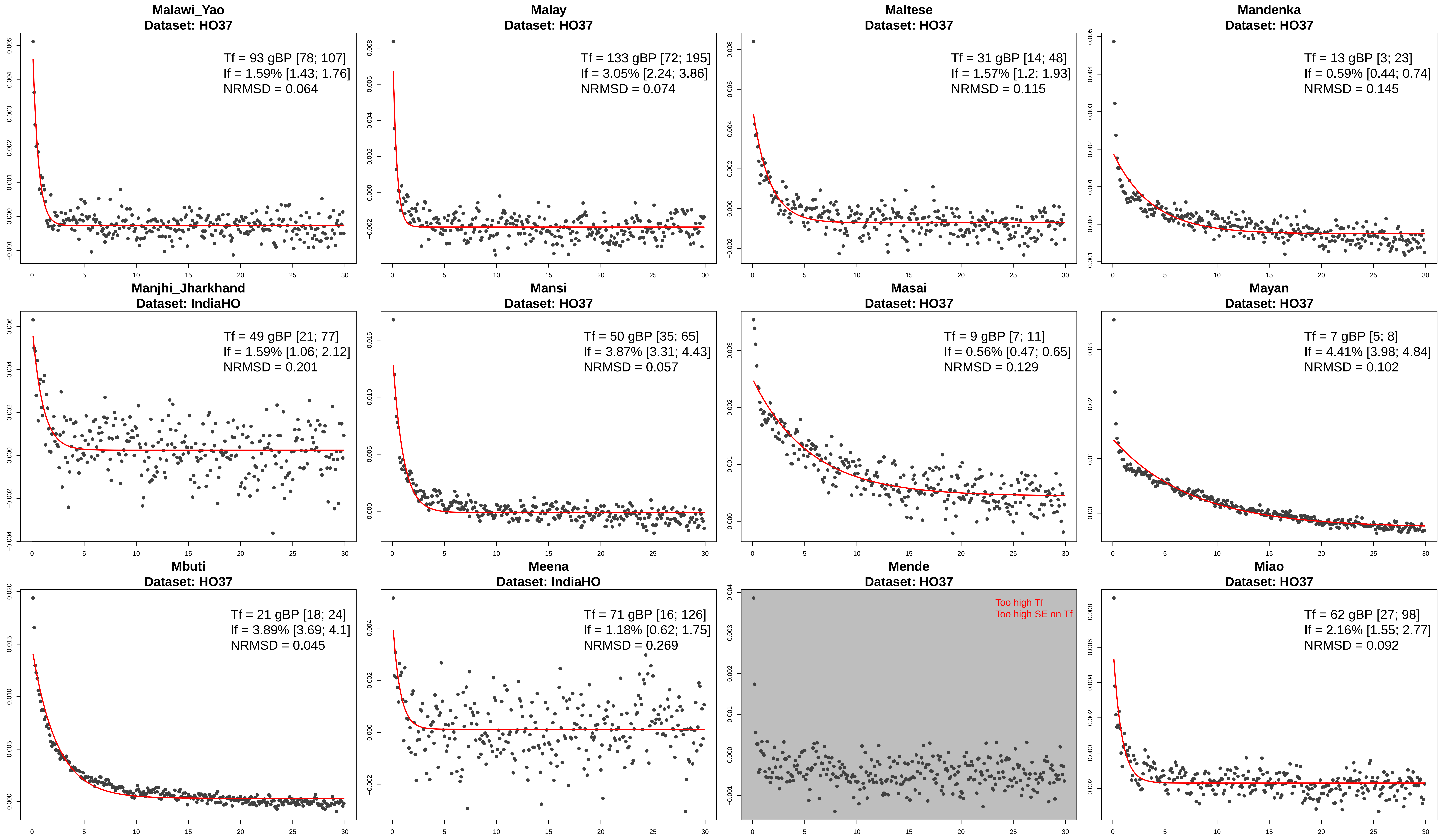
Malawi_Yao
Dataset: HO37
Tf = 93 gBP [78; 107]
If = 1.59% [1.43; 1.76]
NRMSD = 0.064
Malay
Dataset: HO37
Tf = 133 gBP [72; 195]
If = 3.05% [2.24; 3.86]
NRMSD = 0.074
Maltese
Dataset: HO37
Tf = 31 gBP [14; 48]
If = 1.57% [1.2; 1.93]
NRMSD = 0.115
Mandenka
Dataset: HO37
Tf = 13 gBP [3; 23]
If = 0.59% [0.44; 0.74]
NRMSD = 0.145
Manjhi_Jharkhand
Dataset: IndiaHO
Tf = 49 gBP [21; 77]
If = 1.59% [1.06; 2.12]
NRMSD = 0.201
Mansi
Dataset: HO37
Tf = 50 gBP [35; 65]
If = 3.87% [3.31; 4.43]
NRMSD = 0.057
Masai
Dataset: HO37
Tf = 9 gBP [7; 11]
If = 0.56% [0.47; 0.65]
NRMSD = 0.129
Mayan
Dataset: HO37
Tf = 7 gBP [5; 8]
If = 4.41% [3.98; 4.84]
NRMSD = 0.102
Mbuti
Dataset: HO37
Tf = 21 gBP [18; 24]
If = 3.89% [3.69; 4.1]
NRMSD = 0.045
Meena
Dataset: IndiaHO
Tf = 71 gBP [16; 126]
If = 1.18% [0.62; 1.75]
NRMSD = 0.269
Mende
Dataset: HO37
Too high Tf
Too high SE on Tf
Miao
Dataset: HO37
Tf = 62 gBP [27; 98]
If = 2.16% [1.55; 2.77]
NRMSD = 0.092

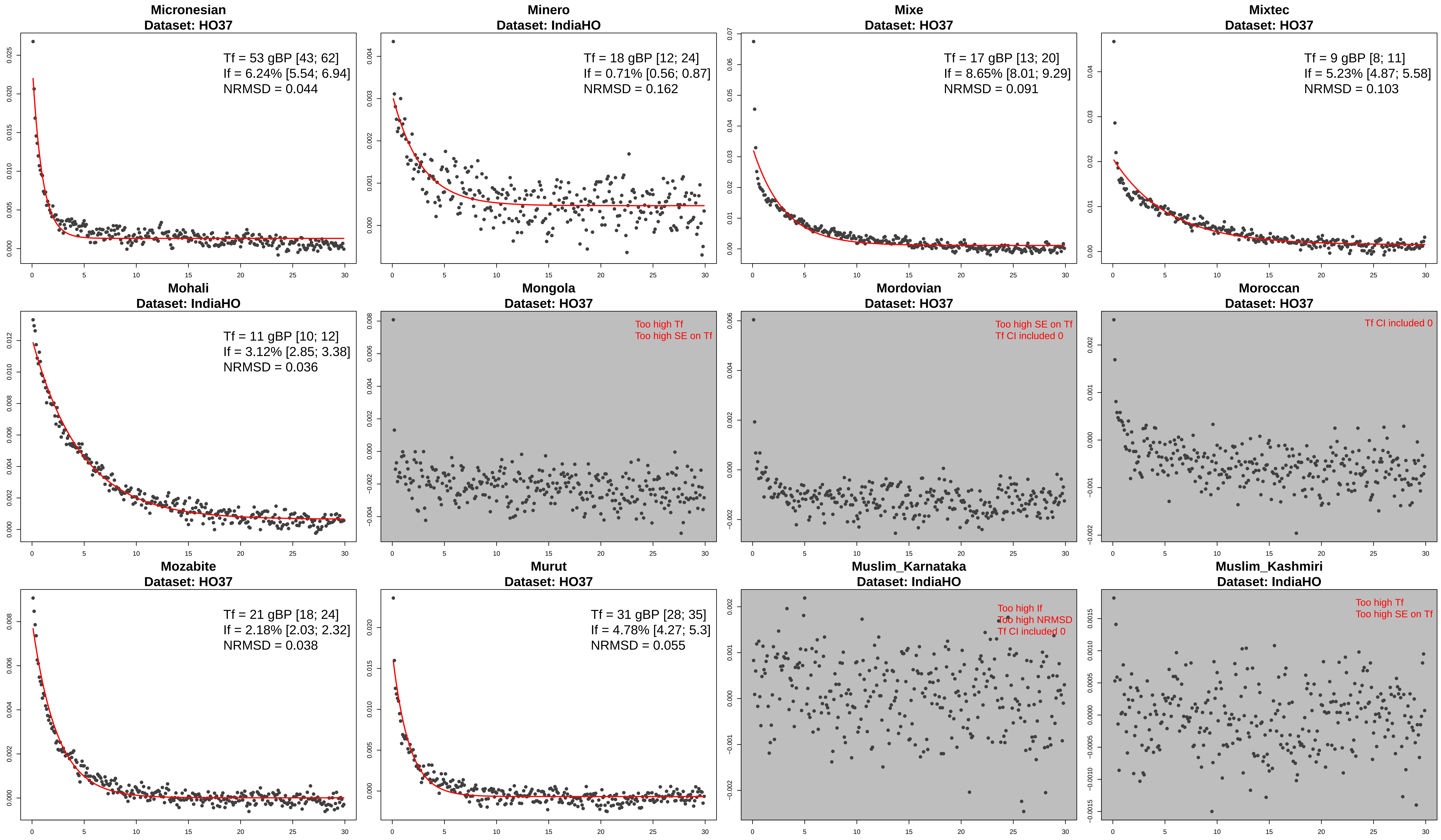
Micronesian
Dataset: HO37
Tf = 53 gBP [43; 62]
If = 6.24% [5.54; 6.94]
NRMSD = 0.044
Minero
Dataset: IndiaHO
Tf = 18 gBP [12; 24]
If = 0.71% [0.56; 0.87]
NRMSD = 0.162
Mixe
Dataset: HO37
Tf = 17 gBP [13; 20]
If = 8.65% [8.01; 9.29]
NRMSD = 0.091
Mixtec
Dataset: HO37
Tf = 9 gBP [8; 11]
If = 5.23% [4.87; 5.58]
NRMSD = 0.103
Mohali
Dataset: IndiaHO
Tf = 11 gBP [10; 12]
If = 3.12% [2.85; 3.38]
NRMSD = 0.036
Mongola
Dataset: HO37
Too high Tf
Too high SE on Tf
Mordovian
Dataset: HO37
Too high SE on Tf
Tf CI included 0
Moroccan
Dataset: HO37
Tf CI included 0
Mozabite
Dataset: HO37
Tf = 21 gBP [18; 24]
If = 2.18% [2.03; 2.32]
NRMSD = 0.038
Murut
Dataset: HO37
Tf = 31 gBP [28; 35]
If = 4.78% [4.27; 5.3]
NRMSD = 0.055
Muslim_Karnataka
Dataset: IndiaHO
Too high If
Too high NRMSD
Tf CI included 0
Muslim_Kashmiri
Dataset: IndiaHO
Too high Tf
Too high SE on Tf

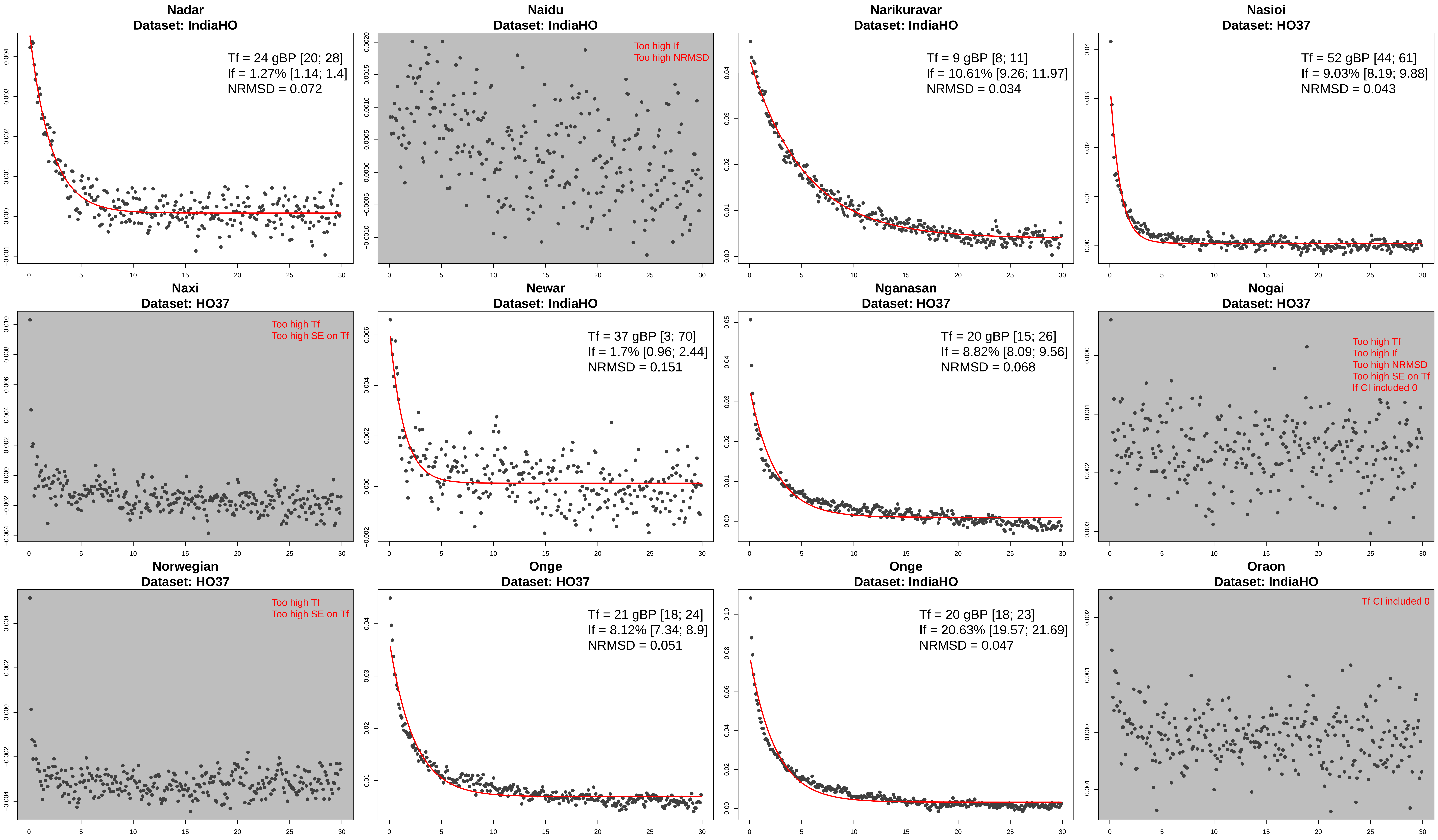
Nadar
Dataset: IndiaHO
Tf = 24 gBP [20; 28]
If = 1.27% [1.14; 1.4]
NRMSD = 0.072
Naidu
Dataset: IndiaHO
Too high If
Too high NRMSD
Narikuravar
Dataset: IndiaHO
Tf = 9 gBP [8; 11]
If = 10.61% [9.26; 11.97]
NRMSD = 0.034
Nasioi
Dataset: HO37
Tf = 52 gBP [44; 61]
If = 9.03% [8.19; 9.88]
NRMSD = 0.043
Naxi
Dataset: HO37
Too high Tf
Too high SE on Tf
Newar
Dataset: IndiaHO
Tf = 37 gBP [3; 70]
If = 1.7% [0.96; 2.44]
NRMSD = 0.151
Nganasan
Dataset: HO37
Tf = 20 gBP [15; 26]
If = 8.82% [8.09; 9.56]
NRMSD = 0.068
Nogai
Dataset: HO37
Too high Tf
Too high If
Too high NRMSD
Too high SE on Tf
If CI included 0
Norwegian
Dataset: HO37
Too high Tf
Too high SE on Tf
Onge
Dataset: HO37
Tf = 21 gBP [18; 24]
If = 8.12% [7.34; 8.9]
NRMSD = 0.051
Onge
Dataset: IndiaHO
Tf = 20 gBP [18; 23]
If = 20.63% [19.57; 21.69]
NRMSD = 0.047
Oraon
Dataset: IndiaHO
Tf CI included 0

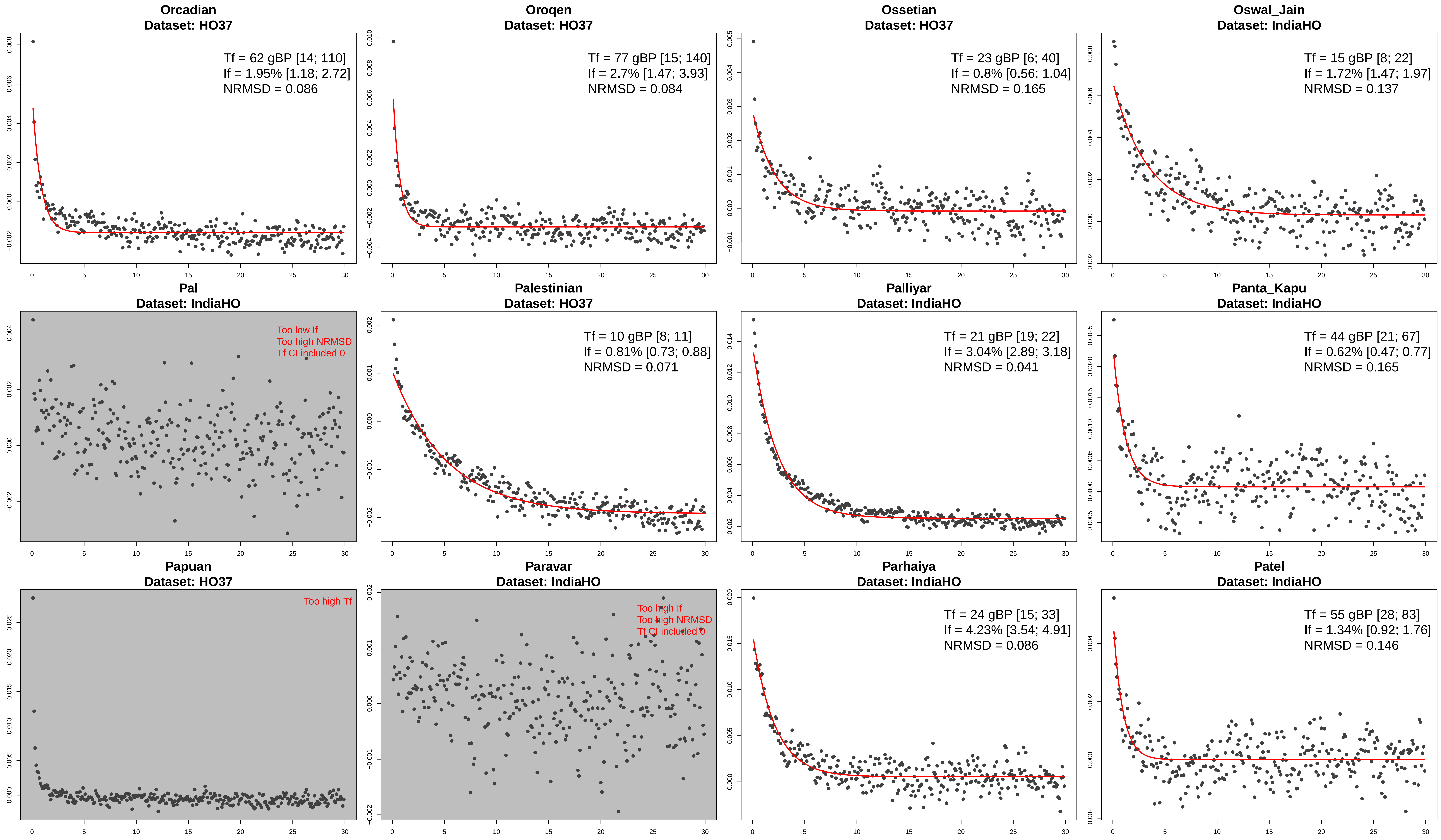
Orcadian
Dataset: HO37
Tf = 62 gBP [14; 110]
If = 1.95% [1.18; 2.72]
NRMSD = 0.086
Oroqen
Dataset: HO37
Tf = 77 gBP [15; 140]
If = 2.7% [1.47; 3.93]
NRMSD = 0.084
Ossetian
Dataset: HO37
Tf = 23 gBP [6; 40]
If = 0.8% [0.56; 1.04]
NRMSD = 0.165
Oswal_Jain
Dataset: IndiaHO
Tf = 15 gBP [8; 22]
If = 1.72% [1.47; 1.97]
NRMSD = 0.137
Pal
Dataset: IndiaHO
Too low If
Too high NRMSD
Tf CI included 0
Palestinian
Dataset: HO37
Tf = 10 gBP [8; 11]
If = 0.81% [0.73; 0.88]
NRMSD = 0.071
Palliyar
Dataset: IndiaHO
Tf = 21 gBP [19; 22]
If = 3.04% [2.89; 3.18]
NRMSD = 0.041
Panta_Kapu
Dataset: IndiaHO
Tf = 44 gBP [21; 67]
If = 0.62% [0.47; 0.77]
NRMSD = 0.165
Papuan
Dataset: HO37
Too high Tf
Paravar
Dataset: IndiaHO
Too high If
Too high NRMSD
Tf CI included 0
Parhaiya
Dataset: IndiaHO
Tf = 24 gBP [15; 33]
If = 4.23% [3.54; 4.91]
NRMSD = 0.086
Patel
Dataset: IndiaHO
Tf = 55 gBP [28; 83]
If = 1.34% [0.92; 1.76]
NRMSD = 0.146

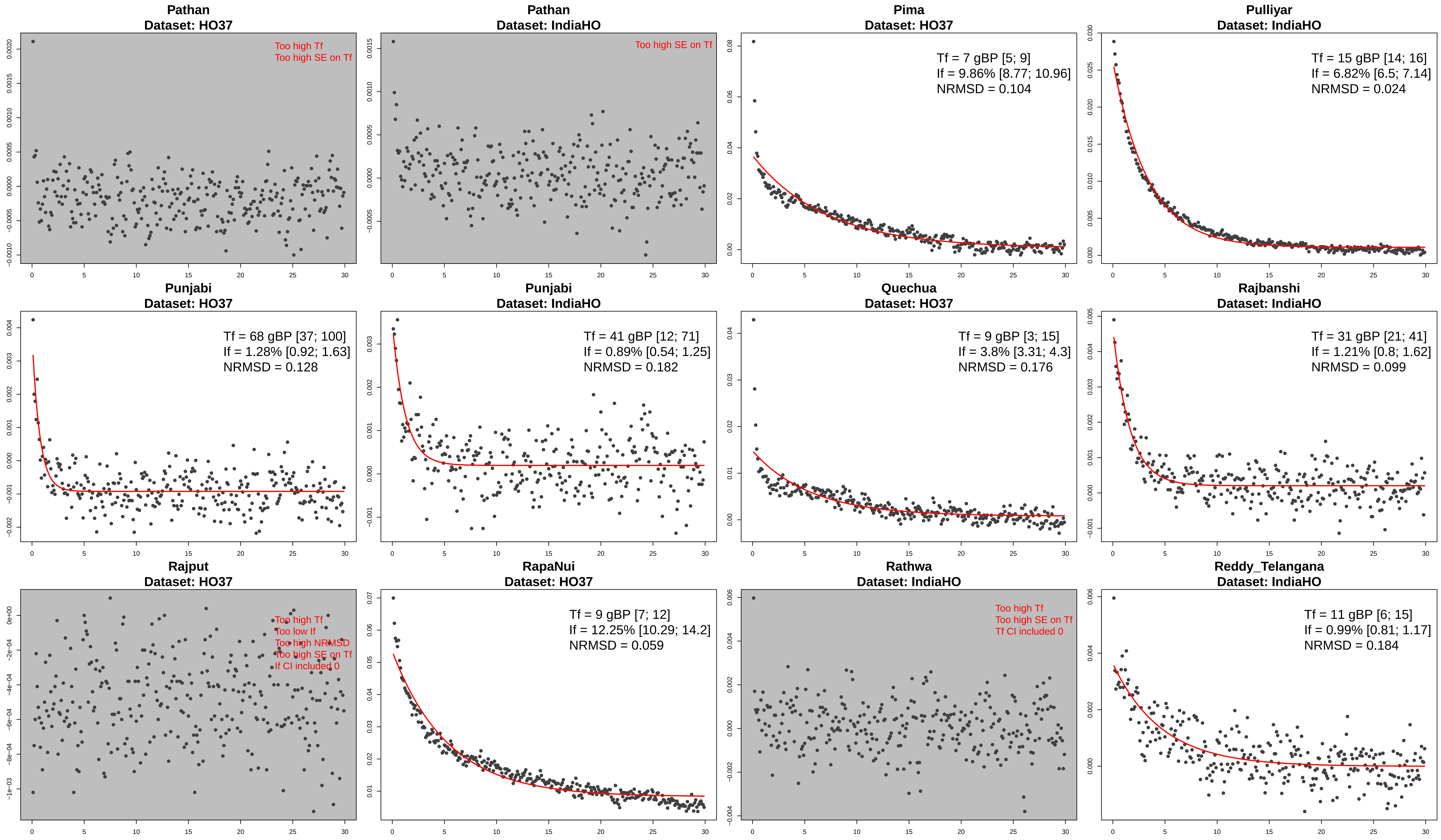
Pathan
Dataset: HO37
Too high Tf
Too high SE on Tf
Pathan
Dataset: IndiaHO
Too high SE on Tf
Pima
Dataset: HO37
Tf = 7 gBP [5; 9]
If = 9.86% [8.77; 10.96]
NRMSD = 0.104
Pulliyar
Dataset: IndiaHO
Tf = 15 gBP [14; 16]
If = 6.82% [6.5; 7.14]
NRMSD = 0.024
Punjabi
Dataset: HO37
Tf = 68 gBP [37; 100]
If = 1.28% [0.92; 1.63]
NRMSD = 0.128
Punjabi
Dataset: IndiaHO
Tf = 41 gBP [12; 71]
If = 0.89% [0.54; 1.25]
NRMSD = 0.182
Quechua
Dataset: HO37
Tf = 9 gBP [3; 15]
If = 3.8% [3.31; 4.3]
NRMSD = 0.176
Rajbanshi
Dataset: IndiaHO
Tf = 31 gBP [21; 41]
If = 1.21% [0.8; 1.62]
NRMSD = 0.099
Rajput
Dataset: HO37
Too high Tf
Too low If
Too high NRMSD
Too high SE on Tf
If CI included 0
RapaNui
Dataset: HO37
Tf = 9 gBP [7; 12]
If = 12.25% [10.29; 14.2]
NRMSD = 0.059
Rathwa
Dataset: IndiaHO
Too high Tf
Too high SE on Tf
Tf CI included 0
Reddy_Telangana
Dataset: IndiaHO
Tf = 11 gBP [6; 15]
If = 0.99% [0.81; 1.17]
NRMSD = 0.184

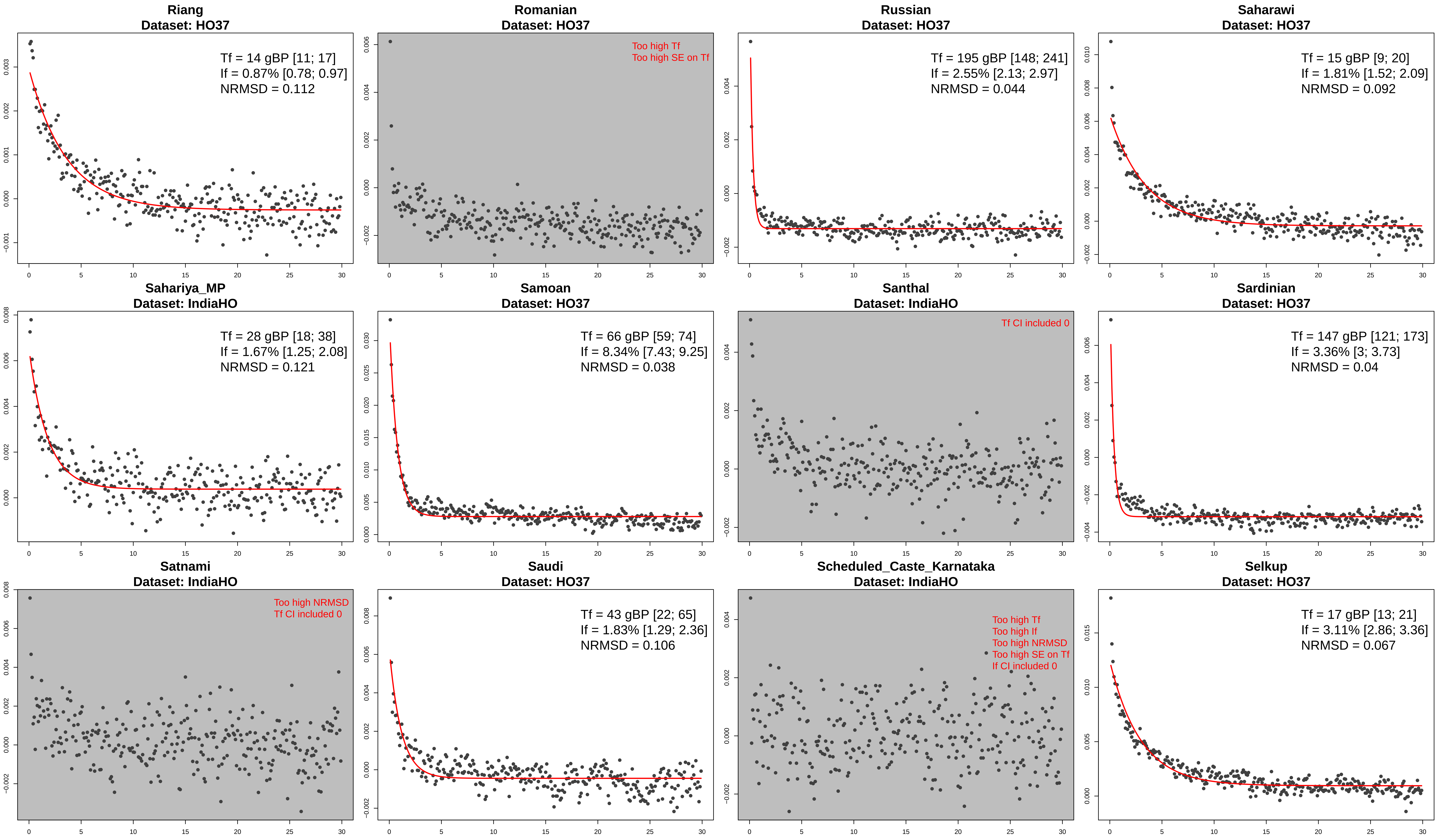
Riang
Dataset: HO37
Tf = 14 gBP [11; 17]
If = 0.87% [0.78; 0.97]
NRMSD = 0.112
Romanian
Dataset: HO37
Too high Tf
Too high SE on Tf
Russian
Dataset: HO37
Tf = 195 gBP [148; 241]
If = 2.55% [2.13; 2.97]
NRMSD = 0.044
Saharawi
Dataset: HO37
Tf = 15 gBP [9; 20]
If = 1.81% [1.52; 2.09]
NRMSD = 0.092
Sahariya_MP
Dataset: IndiaHO
Tf = 28 gBP [18; 38]
If = 1.67% [1.25; 2.08]
NRMSD = 0.121
Samoan
Dataset: HO37
Tf = 66 gBP [59; 74]
If = 8.34% [7.43; 9.25]
NRMSD = 0.038
Santhal
Dataset: IndiaHO
Tf CI included 0
Sardinian
Dataset: HO37
Tf = 147 gBP [121; 173]
If = 3.36% [3; 3.73]
NRMSD = 0.04
Satnami
Dataset: IndiaHO
Too high NRMSD
Tf CI included 0
Saudi
Dataset: HO37
Tf = 43 gBP [22; 65]
If = 1.83% [1.29; 2.36]
NRMSD = 0.106
Scheduled_Caste_Karnataka
Dataset: IndiaHO
Too high Tf
Too high If
Too high NRMSD
Too high SE on Tf
If CI included 0
Selkup
Dataset: HO37
Tf = 17 gBP [13; 21]
If = 3.11% [2.86; 3.36]
NRMSD = 0.067

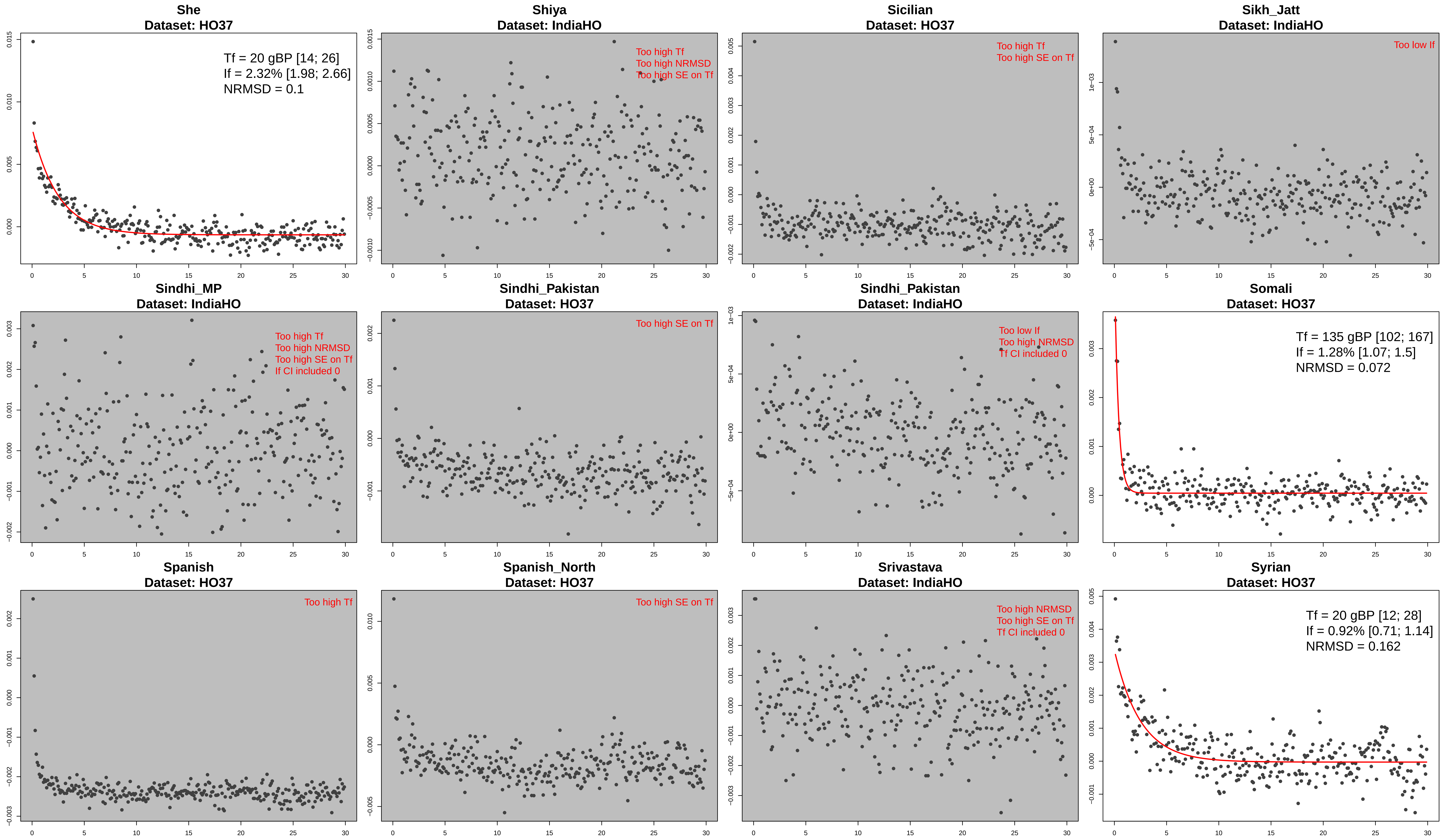

She
Dataset: HO37
Tf = 20 gBP [14; 26]
If = 2.32% [1.98; 2.66]
NRMSD = 0.1
Shiya
Dataset: IndiaHO
Too high Tf
Too high NRMSD
Too high SE on Tf
Sicilian
Dataset: HO37
Too high Tf
Too high SE on Tf
Sikh_Jatt
Dataset: IndiaHO
Too low If
Sindhi_MP
Dataset: IndiaHO
Too high Tf
Too high NRMSD
Too high SE on Tf
If CI included 0
Sindhi_Pakistan
Dataset: HO37
Too high SE on Tf
Sindhi_Pakistan
Dataset: IndiaHO
Too low If
Too high NRMSD
Tf CI included 0
Somali
Dataset: HO37
Tf = 135 gBP [102; 167]
If = 1.28% [1.07; 1.5]
NRMSD = 0.072
Spanish
Dataset: HO37
Too high Tf
Spanish_North
Dataset: HO37
Too high SE on Tf
Srivastava
Dataset: IndiaHO
Too high NRMSD
Too high SE on Tf
Tf CI included 0
Syrian
Dataset: HO37
Tf = 20 gBP [12; 28]
If = 0.92% [0.71; 1.14]
NRMSD = 0.162

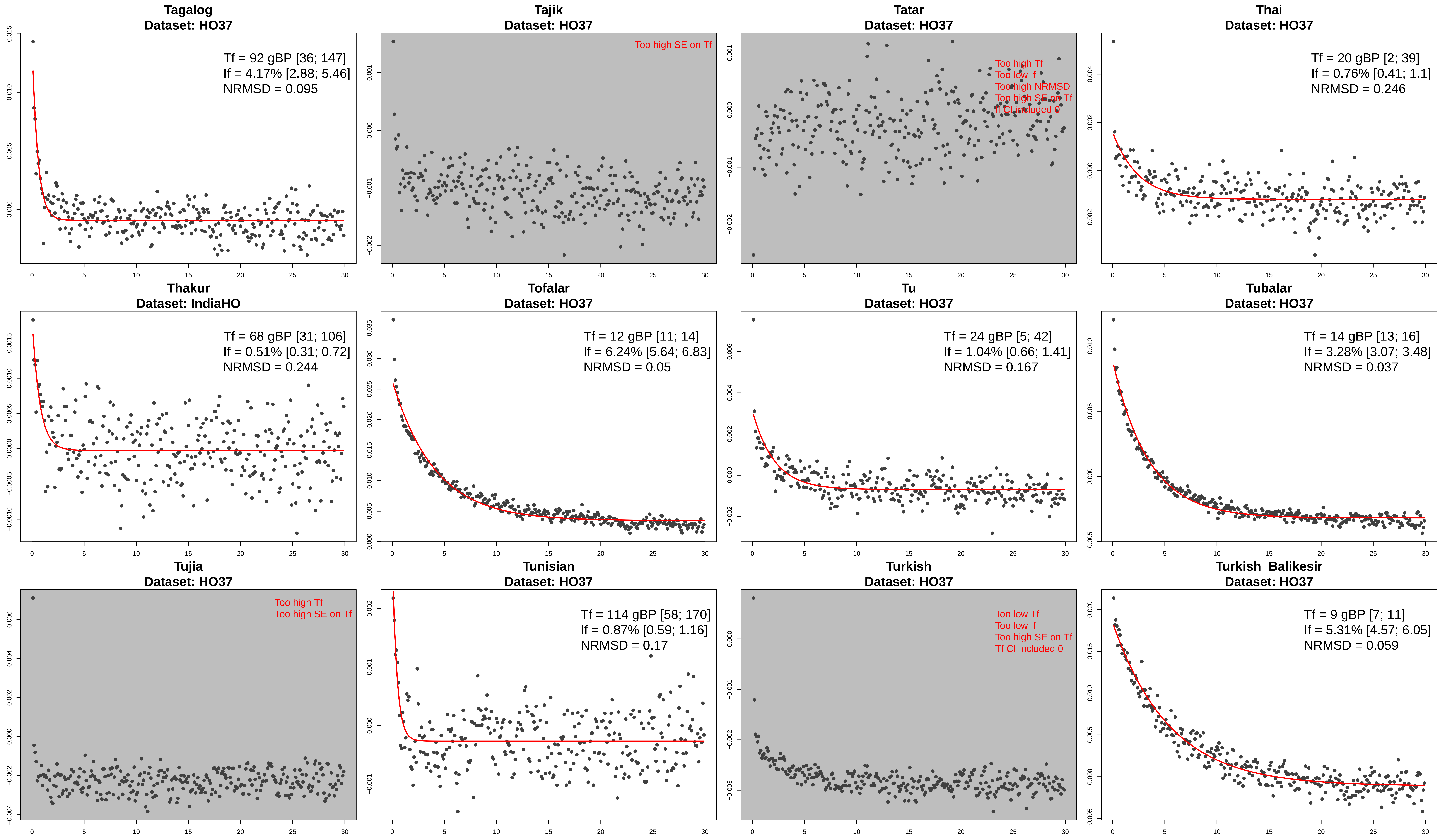

Tagalog
Dataset: HO37
Tf = 92 gBP [36; 147]
If = 4.17% [2.88; 5.46]
NRMSD = 0.095
Tajik
Dataset: HO37
Too high SE on Tf
Tatar
Dataset: HO37
Too high Tf
Too low If
Too high NRMSD
Too high SE on Tf
Tf CI included 0
Thai
Dataset: HO37
Tf = 20 gBP [2; 39]
If = 0.76% [0.41; 1.1]
NRMSD = 0.246
Thakur
Dataset: IndiaHO
Tf = 68 gBP [31; 106]
If = 0.51% [0.31; 0.72]
NRMSD = 0.244
Tofalar
Dataset: HO37
Tf = 12 gBP [11; 14]
If = 6.24% [5.64; 6.83]
NRMSD = 0.05
Tu
Dataset: HO37
Tf = 24 gBP [5; 42]
If = 1.04% [0.66; 1.41]
NRMSD = 0.167
Tubalar
Dataset: HO37
Tf = 14 gBP [13; 16]
If = 3.28% [3.07; 3.48]
NRMSD = 0.037
Tujia
Dataset: HO37
Too high Tf
Too high SE on Tf
Tunisian
Dataset: HO37
Tf = 114 gBP [58; 170]
If = 0.87% [0.59; 1.16]
NRMSD = 0.17
Turkish
Dataset: HO37
Too low Tf
Too low If
Too high SE on Tf
Tf CI included 0
Turkish_Balikesir
Dataset: HO37
Tf = 9 gBP [7; 11]
If = 5.31% [4.57; 6.05]
NRMSD = 0.059

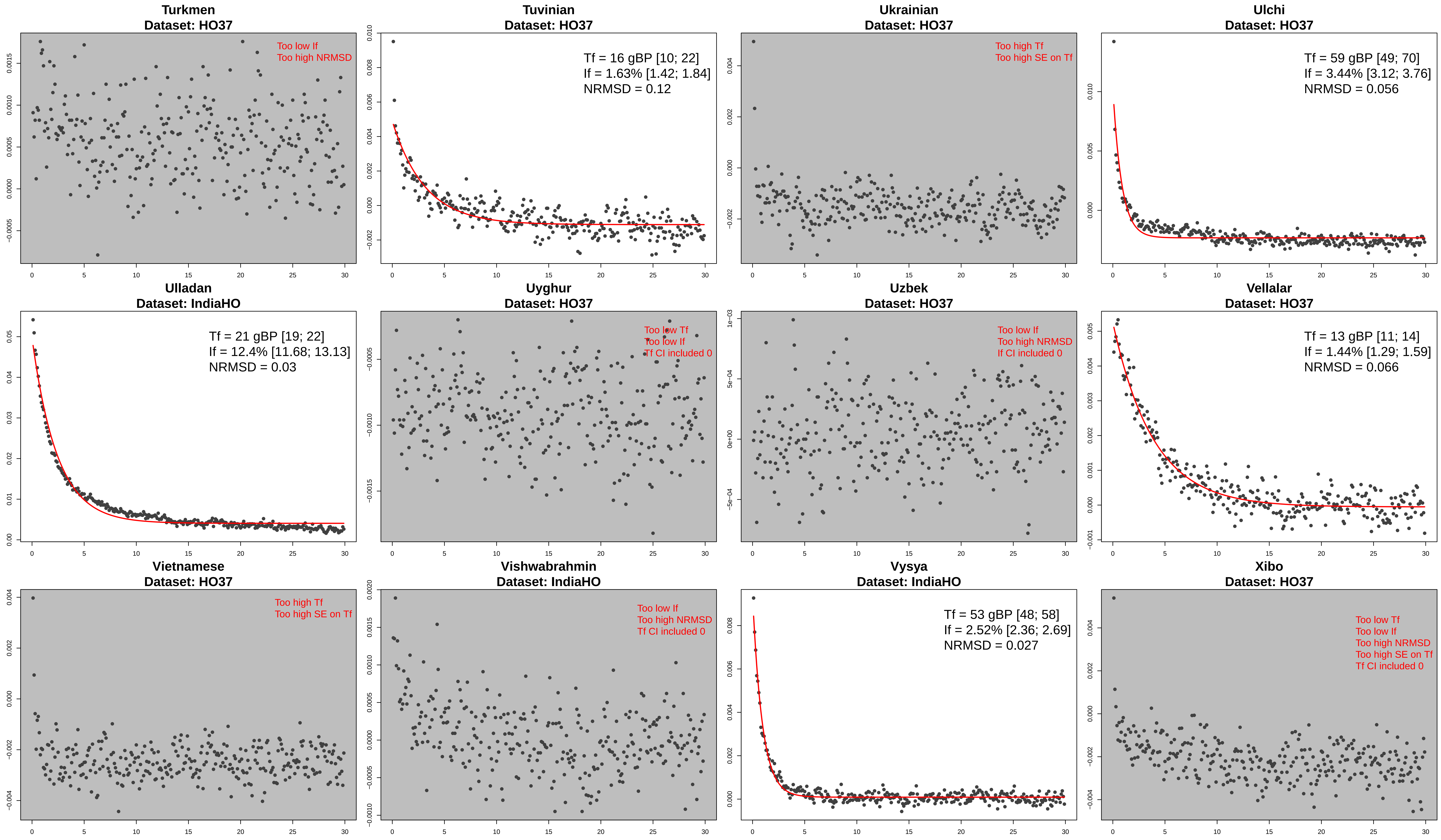

Turkmen
Dataset: HO37
Too low If
Too high NRMSD
Tuvinian
Dataset: HO37
Tf = 16 gBP [10; 22]
If = 1.63% [1.42; 1.84]
NRMSD = 0.12
Ukrainian
Dataset: HO37
Too high Tf
Too high SE on Tf
Ulchi
Dataset: HO37
Tf = 59 gBP [49; 70]
If = 3.44% [3.12; 3.76]
NRMSD = 0.056
Ulladan
Dataset: IndiaHO
Tf = 21 gBP [19; 22]
If = 12.4% [11.68; 13.13]
NRMSD = 0.03
Uyghur
Dataset: HO37
Too low Tf
Too low If
Tf CI included 0
Uzbek
Dataset: HO37
Too low If
Too high NRMSD
If CI included 0
Vellalar
Dataset: HO37
Tf = 13 gBP [11; 14]
If = 1.44% [1.29; 1.59]
NRMSD = 0.066
Vietnamese
Dataset: HO37
Too high Tf
Too high SE on Tf
Vishwabrahmin
Dataset: IndiaHO
Too low If
Too high NRMSD
Tf CI included 0
Vysya
Dataset: IndiaHO
Tf = 53 gBP [48; 58]
If = 2.52% [2.36; 2.69]
NRMSD = 0.027
Xibo
Dataset: HO37
Too low Tf
Too low If
Too high NRMSD
Too high SE on Tf
Tf CI included 0

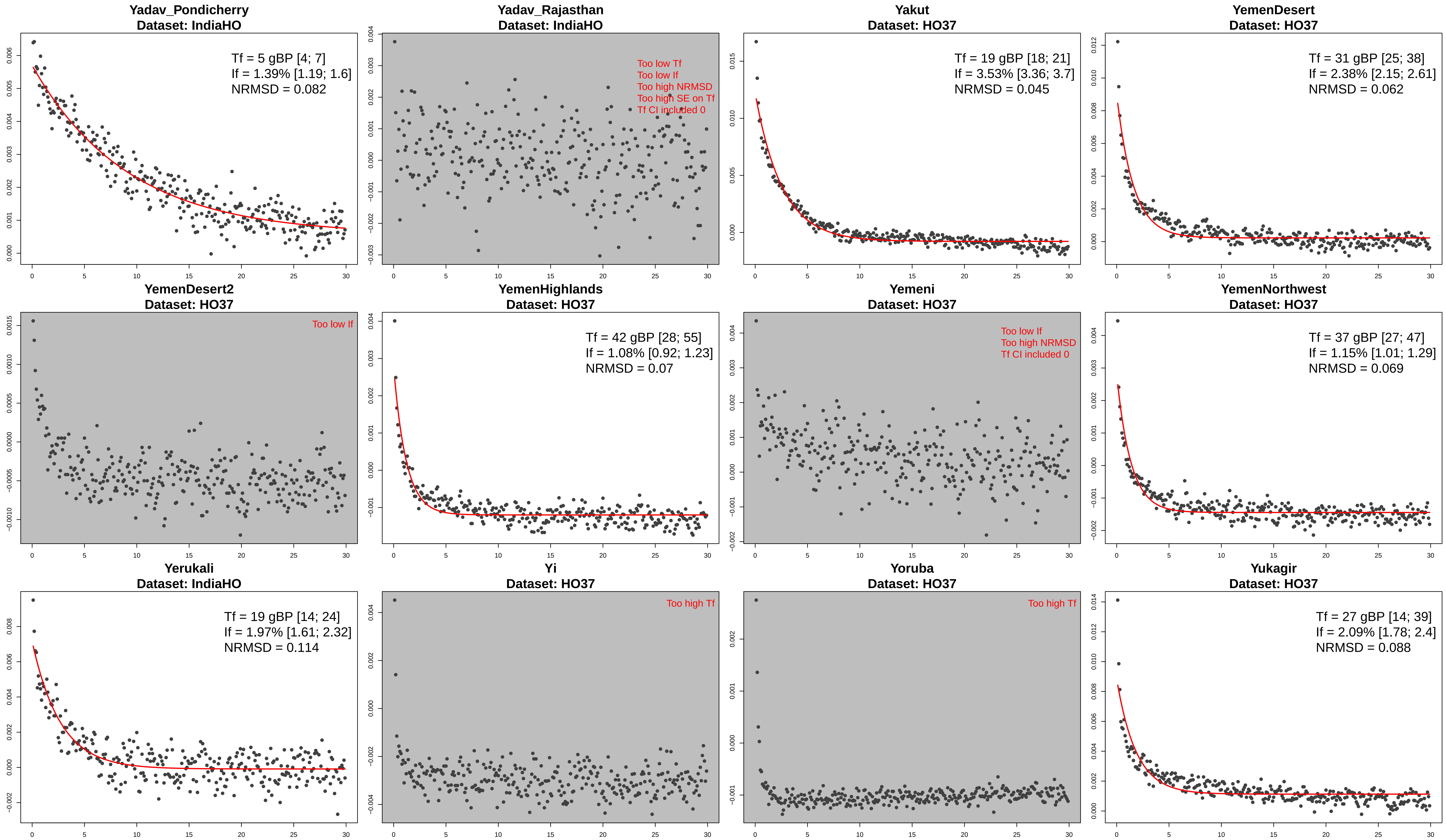
Yadav_Pondicherry
Dataset: IndiaHO
Tf = 5 gBP [4; 7]
If = 1.39% [1.19; 1.6]
NRMSD = 0.082
Yadav_Rajasthan
Dataset: IndiaHO
Too low Tf
Too low If
Too high NRMSD
Too high SE on Tf
Tf CI included 0
Yakut
Dataset: HO37
Tf = 19 gBP [18; 21]
If = 3.53% [3.36; 3.7]
NRMSD = 0.045
YemenDesert
Dataset: HO37
Tf = 31 gBP [25; 38]
If = 2.38% [2.15; 2.61]
NRMSD = 0.062
YemenDesert2
Dataset: HO37
Too low If
YemenHighlands
Dataset: HO37
Tf = 42 gBP [28; 55]
If = 1.08% [0.92; 1.23]
NRMSD = 0.07
Yemeni
Dataset: HO37
Too low If
Too high NRMSD
Tf CI included 0
YemenNorthwest
Dataset: HO37
Tf = 37 gBP [27; 47]
If = 1.15% [1.01; 1.29]
NRMSD = 0.069
Yerukali
Dataset: IndiaHO
Tf = 19 gBP [14; 24]
If = 1.97% [1.61; 2.32]
NRMSD = 0.114
Yi
Dataset: HO37
Too high Tf
Yoruba
Dataset: HO37
Too high Tf
Yukagir
Dataset: HO37
Tf = 27 gBP [14; 39]
If = 2.09% [1.78; 2.4]
NRMSD = 0.088

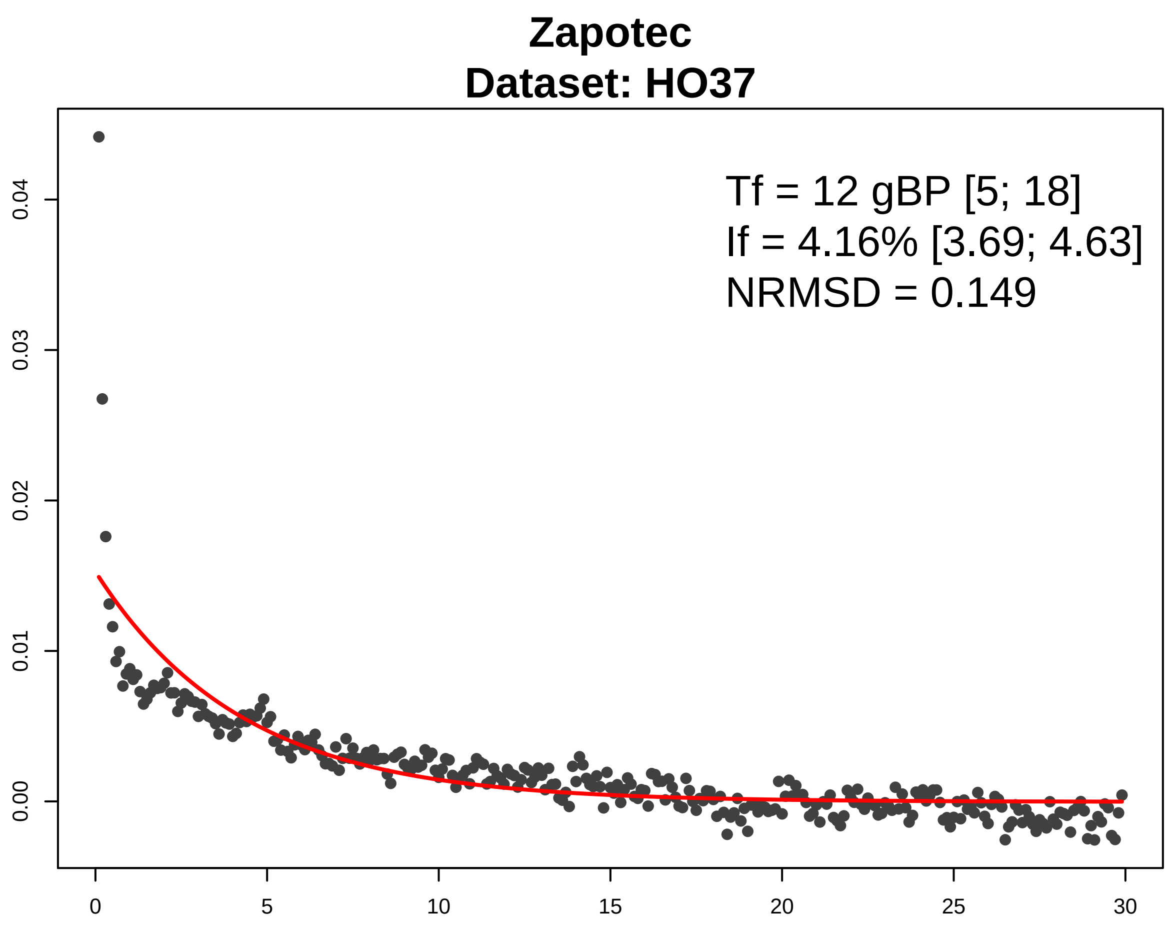

Zapotec
Dataset: HO37
Tf = 12 gBP [5; 18]
If = 4.16% [3.69; 4.63]
NRMSD = 0.149
0.00
0.01
0.02
0.03
0.04
0
5
10
15
20
25
30

**Figure S2 - Distribution of the founder age and intensity estimated in present-day Jewish communities from the HO37 dataset.** The x-axis shows the analyzed Jewish groups, ordered by increasing founder intensity. The y-axis shows the estimated founder intensity. Vertical bars show the 95% confidence intervals. The colors are proportional to the founder age that was estimated in generations.

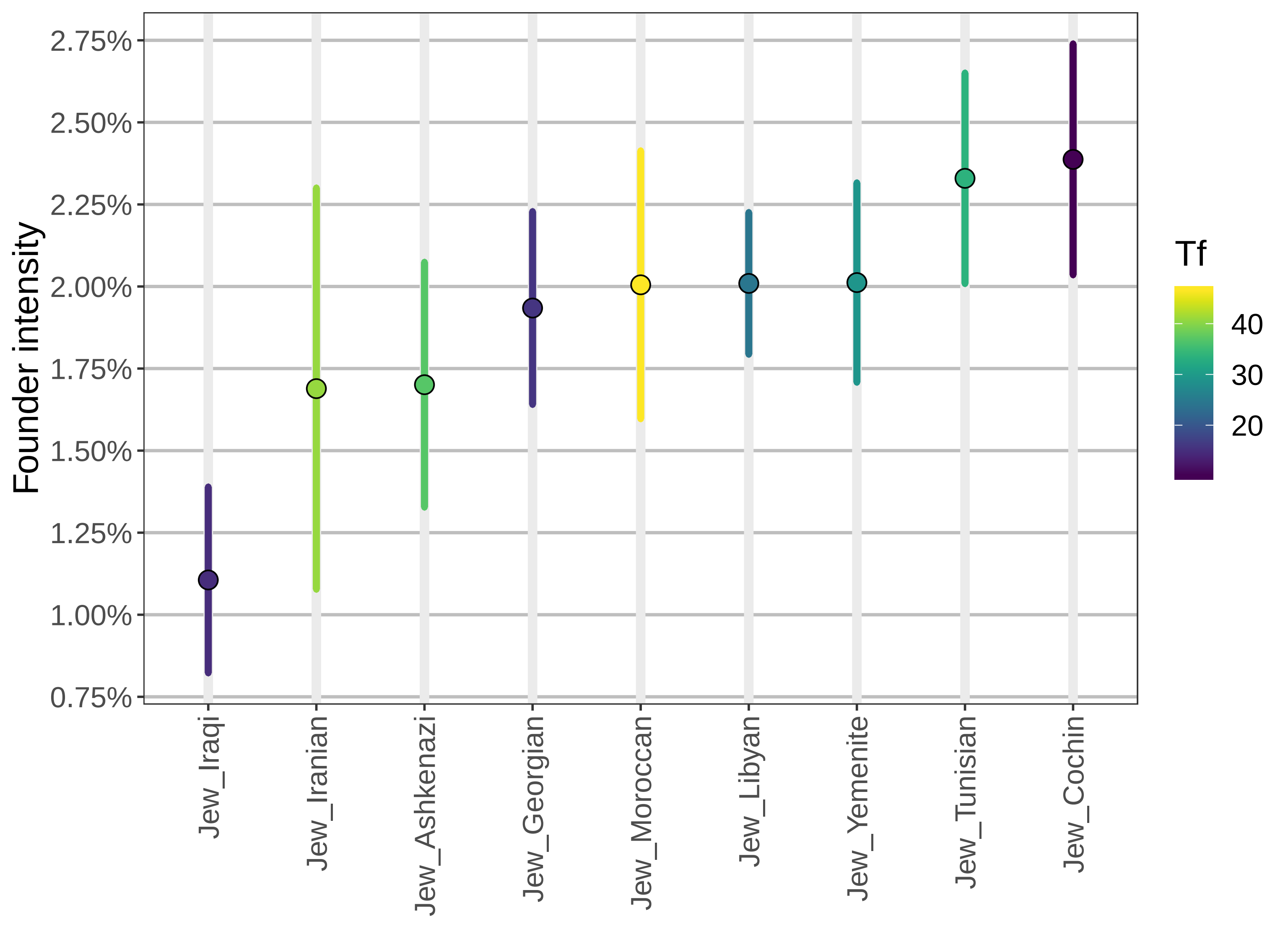

Founder intensity
2.75%
2.50%
2.25%
2.00%
1.75%
1.50%
1.25%
1.00%
0.75%
Jew_Iraqi
Jew_Iranian
Jew_Ashkenazi
Jew_Georgian
Jew_Moroccan
Jew_Libyan
Jew_Yemenite
Jew_Tunisian
Jew_Cochin
Tf
40
30
20

**Figure S3 - History of founder events in present-day South Asian populations from the IndiaHO dataset.** Panel (A) shows the distribution of founder intensity and panel (B) shows the distribution of founder ages for all present-day populations that passed our filtering criteria and showed evidence for a significant founder event in the IndiaHO dataset (see Methods). Each point represents a population and the shape of the points indicates whether the group lives on an island (triangle) or in a continental region (circular). The top colored ribbon represents the linguistic affiliation of the group. *(A) Distribution of the estimated founder intensities.* We show the founder intensity (point) and the associated 95% confidence interval. The populations are ordered by their linguistic affiliation and by increasing order of estimated founder intensity. The black horizontal line shows the inferred founder intensity in Ashkenazi Jews in the IndiaHO dataset (2.0%). Populations that have experienced a founder intensity significantly stronger than in Ashkenazi Jews (i.e. whose founder intensity is greater than the upper-bound of the 95% confidence interval of Ashkenazi Jews) are colored in gold, else they are shown in black. *(B) Distribution of the estimated founder ages*. We show the estimated founder ages and the associated 95% confidence intervals. The estimated ages were converted from generations to years assuming an average generation time of 28 years[20].

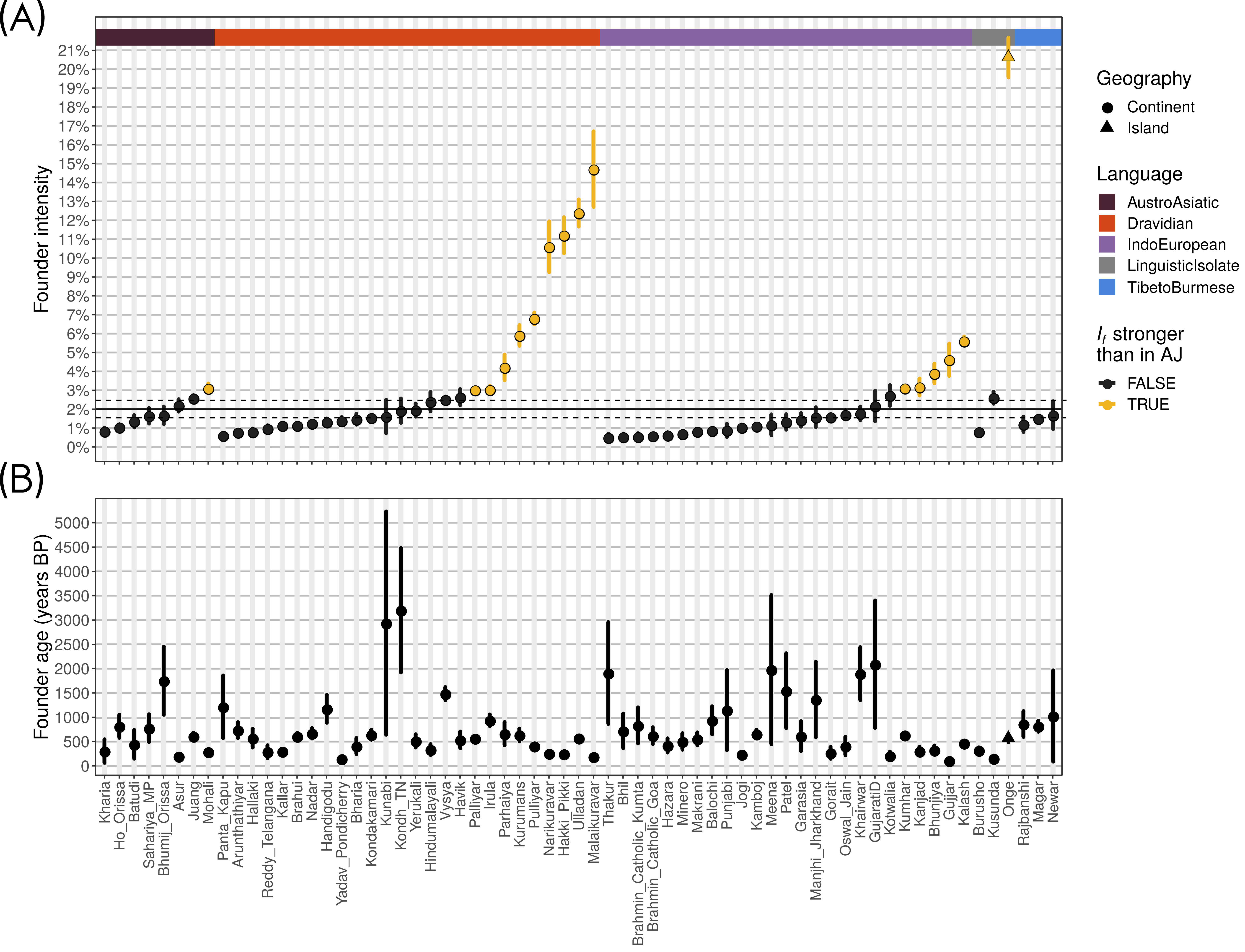
(A)
Founder intensity
21%
20%
19%
18%
17%
16%
15%
14%
13%
12%
11%
10%
9%
8%
7%
6%
5%
4%
3%
2%
1%
0%
Geography
Continent
Island
Language
AustroAsiatic
Dravidian
IndoEuropean
LinguisticIsolate
TibetoBurmese
$I_f$ stronger than in AJ
FALSE
TRUE
(B)
Founder age (years BP)
5000
4500
4000
3500
3000
2500
2000
1500
1000
500
0
Kharia
Ho_Orissa
Batudi
Sahariya_MP
Bhumij_Orissa
Asur
Juang
Mohali
Panta_Kapu
Arunthathiyar
Hallaki
Reddy_Telangana
Kallar
Brahui
Nadar
Handigodu
Yadav_Pondicherry
Bharia
Kondakamari
Kunabi
Kondh_TN
Yerukali
Hindumalayali
Vysya
Havik
Palliyar
Irula
Parhaiya
Kurumans
Pulliyar
Narikuravar
Hakki_Pikki
Ulladan
Malaikuravar
Thakur
Bhil
Brahmin_Catholic_Kumta
Brahmin_Catholic_Goa
Hazara
Minero
Makrani
Balochi
Punjabi
Jogi
Kamboj
Meena
Patel
Garasia
Manjhi_Jharkhand
Gorait
Oswal_Jain
Khairwar
GujaratiD
Kotwalia
Kumhar
Kanjad
Bhunjiya
Gujjar
Kalash
Burusho
Kusunda
Onge
Rajbanshi
Magar
Newar

**Figure S4 - Decay curves for ancient human genomes from the HO37 and HO42 datasets.** The x-axis represents the genetic distance (in cM) and the y-axis represents the average allele sharing correlation. The legend shows the mean and 95% confidence interval for the founder age ($T_f$) in generations before sampling age of the ancient specimen (gBS) and the founder intensity ($I_f$), as well as the NRMSD (see Methods). The panels are greyed when the exponential fitting failed or when the evidence for the founder event was not significant (see Methods). The specific reason is highlighted in red in the legend.

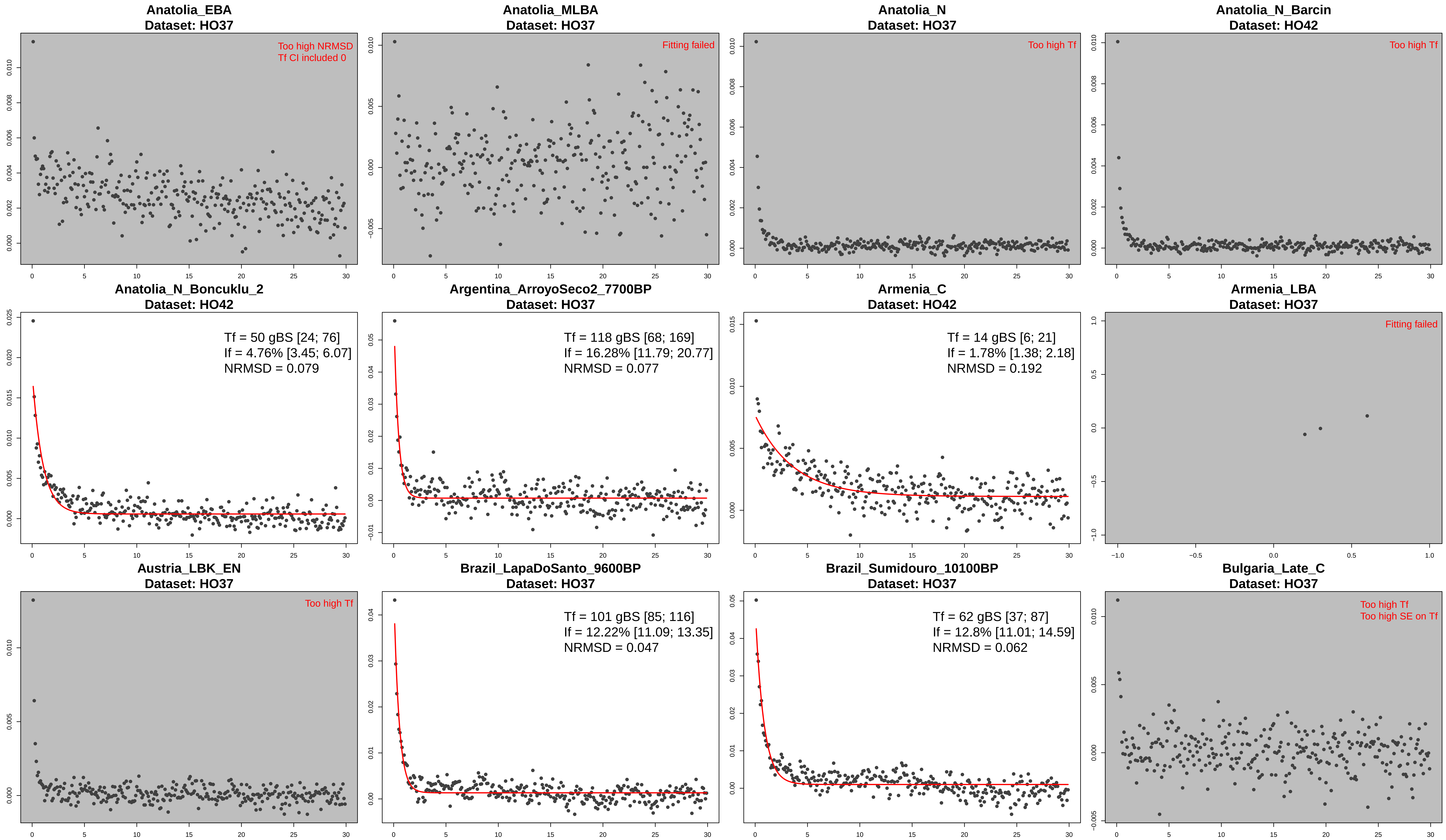
Anatolia_EBA
Dataset: HO37
Too high NRMSD
Tf CI included 0
Anatolia_MLBA
Dataset: HO37
Fitting failed
Anatolia_N
Dataset: HO37
Too high Tf
Anatolia_N_Barcin
Dataset: HO42
Too high Tf
Anatolia_N_Boncuklu_2
Dataset: HO42
Tf = 50 gBS [24; 76]
If = 4.76% [3.45; 6.07]
NRMSD = 0.079
Argentina_ArroyoSeco2_7700BP
Dataset: HO37
Tf = 118 gBS [68; 169]
If = 16.28% [11.79; 20.77]
NRMSD = 0.077
Armenia_C
Dataset: HO42
Tf = 14 gBS [6; 21]
If = 1.78% [1.38; 2.18]
NRMSD = 0.192
Armenia_LBA
Dataset: HO37
Fitting failed
Austria_LBK_EN
Dataset: HO37
Too high Tf
Brazil_LapaDoSanto_9600BP
Dataset: HO37
Tf = 101 gBS [85; 116]
If = 12.22% [11.09; 13.35]
NRMSD = 0.047
Brazil_Sumidouro_10100BP
Dataset: HO37
Tf = 62 gBS [37; 87]
If = 12.8% [11.01; 14.59]
NRMSD = 0.062
Bulgaria_Late_C
Dataset: HO37
Too high Tf
Too high SE on Tf

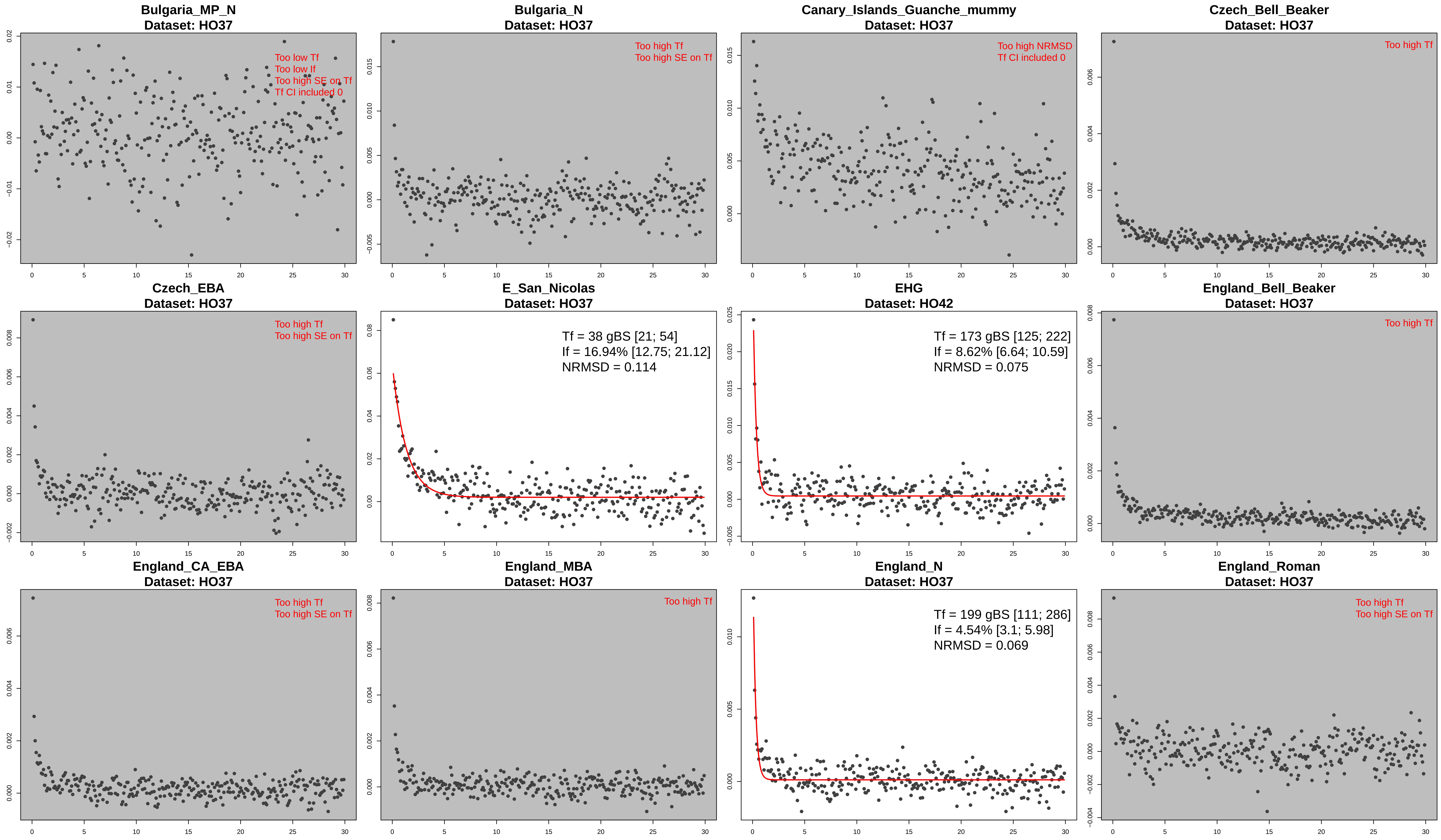

Bulgaria_MP_N
Dataset: HO37
Too low Tf
Too low If
Too high SE on Tf
Tf CI included 0
Bulgaria_N
Dataset: HO37
Too high Tf
Too high SE on Tf
Canary_Islands_Guanche_mummy
Dataset: HO37
Too high NRMSD
Tf CI included 0
Czech_Bell_Beaker
Dataset: HO37
Too high Tf
Czech_EBA
Dataset: HO37
Too high Tf
Too high SE on Tf
E_San_Nicolas
Dataset: HO37
Tf = 38 gBS [21; 54]
If = 16.94% [12.75; 21.12]
NRMSD = 0.114
EHG
Dataset: HO42
Tf = 173 gBS [125; 222]
If = 8.62% [6.64; 10.59]
NRMSD = 0.075
England_Bell_Beaker
Dataset: HO37
Too high Tf
England_CA_EBA
Dataset: HO37
Too high Tf
Too high SE on Tf
England_MBA
Dataset: HO37
Too high Tf
England_N
Dataset: HO37
Tf = 199 gBS [111; 286]
If = 4.54% [3.1; 5.98]
NRMSD = 0.069
England_Roman
Dataset: HO37
Too high Tf
Too high SE on Tf

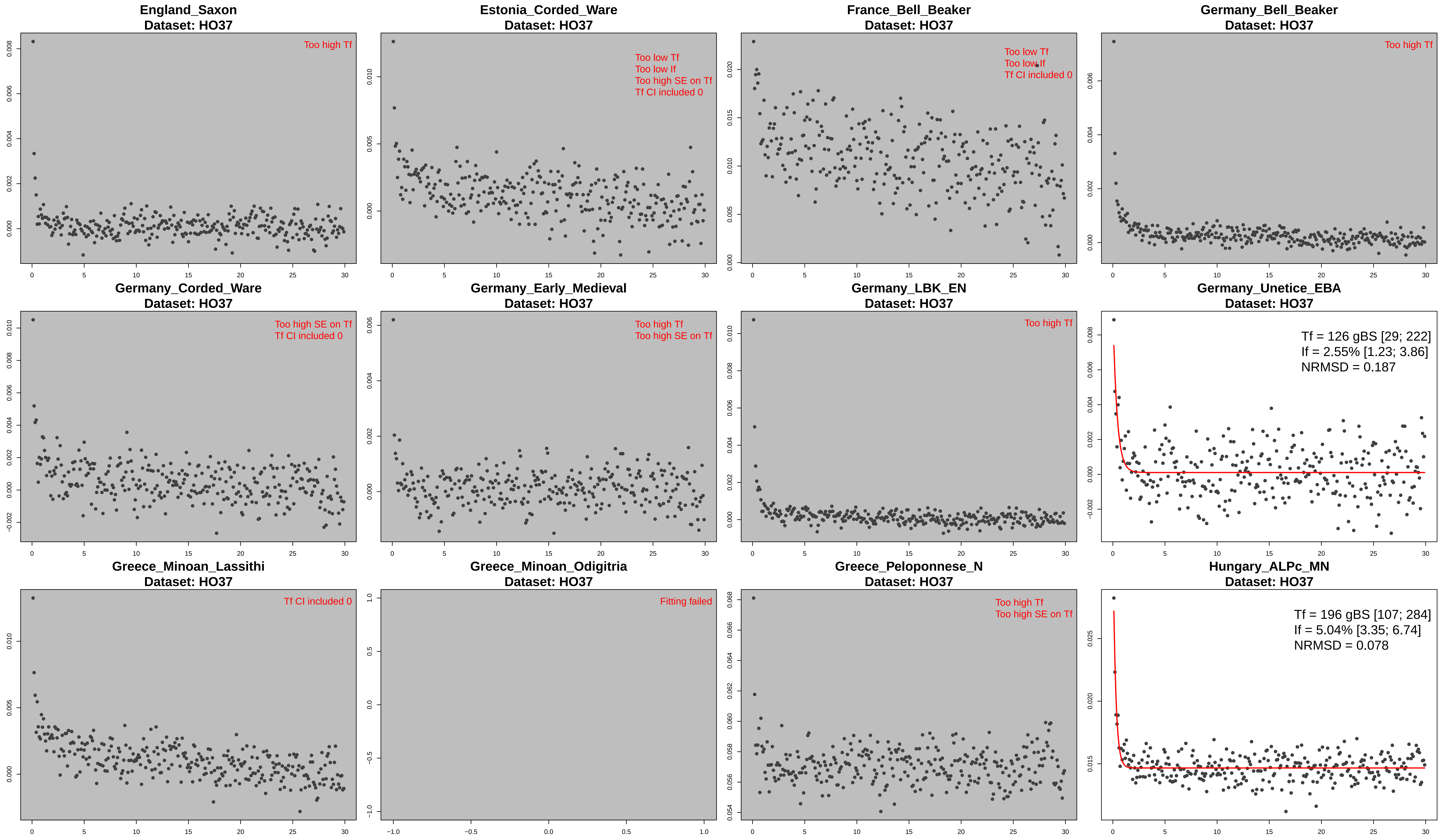
England_Saxon
Dataset: HO37
Too high Tf
Estonia_Corded_Ware
Dataset: HO37
Too low Tf
Too low If
Too high SE on Tf
Tf CI included 0
France_Bell_Beaker
Dataset: HO37
Too low Tf
Too low If
Tf CI included 0
Germany_Bell_Beaker
Dataset: HO37
Too high Tf
Germany_Corded_Ware
Dataset: HO37
Too high SE on Tf
Tf CI included 0
Germany_Early_Medieval
Dataset: HO37
Too high Tf
Too high SE on Tf
Germany_LBK_EN
Dataset: HO37
Too high Tf
Germany_Unetice_EBA
Dataset: HO37
Tf = 126 gBS [29; 222]
If = 2.55% [1.23; 3.86]
NRMSD = 0.187
Greece_Minoan_Lassithi
Dataset: HO37
Tf CI included 0
Greece_Minoan_Odigitria
Dataset: HO37
Fitting failed
Greece_Peloponnese_N
Dataset: HO37
Too high Tf
Too high SE on Tf
Hungary_ALPc_MN
Dataset: HO37
Tf = 196 gBS [107; 284]
If = 5.04% [3.35; 6.74]
NRMSD = 0.078

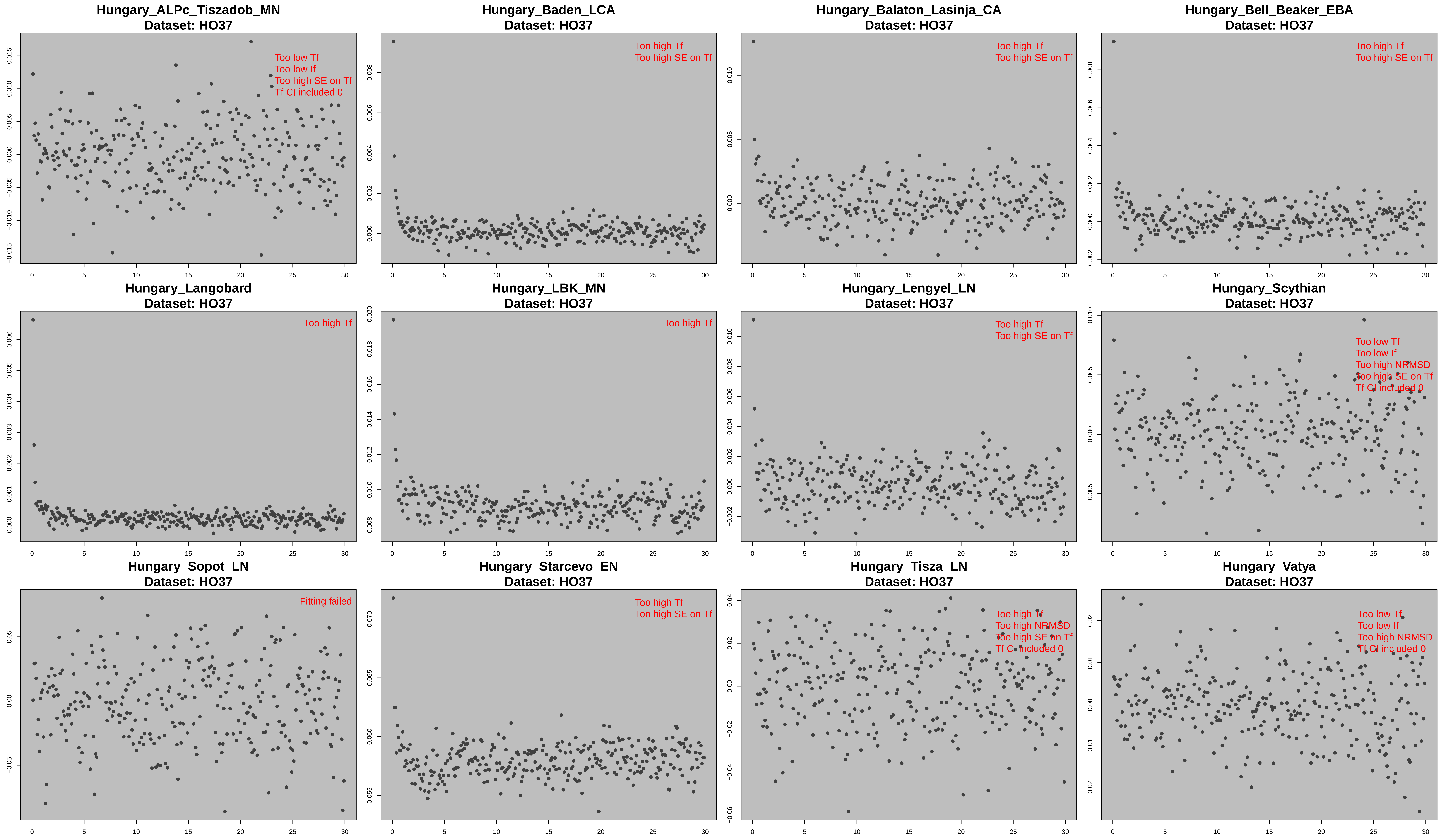
Hungary_ALPc_Tiszadob_MN
Dataset: HO37
Too low Tf
Too low If
Too high SE on Tf
Tf CI included 0
Hungary_Baden_LCA
Dataset: HO37
Too high Tf
Too high SE on Tf
Hungary_Balaton_Lasinja_CA
Dataset: HO37
Too high Tf
Too high SE on Tf
Hungary_Bell_Beaker_EBA
Dataset: HO37
Too high Tf
Too high SE on Tf
Hungary_Langobard
Dataset: HO37
Too high Tf
Hungary_LBK_MN
Dataset: HO37
Too high Tf
Hungary_Lengyel_LN
Dataset: HO37
Too high Tf
Too high SE on Tf
Hungary_Scythian
Dataset: HO37
Too low Tf
Too low If
Too high NRMSD
Too high SE on Tf
Tf CI included 0
Hungary_Sopot_LN
Dataset: HO37
Fitting failed
Hungary_Starcevo_EN
Dataset: HO37
Too high Tf
Too high SE on Tf
Hungary_Tisza_LN
Dataset: HO37
Too high Tf
Too high NRMSD
Too high SE on Tf
Tf CI included 0
Hungary_Vatya
Dataset: HO37
Too low Tf
Too low If
Too high NRMSD
Tf CI included 0

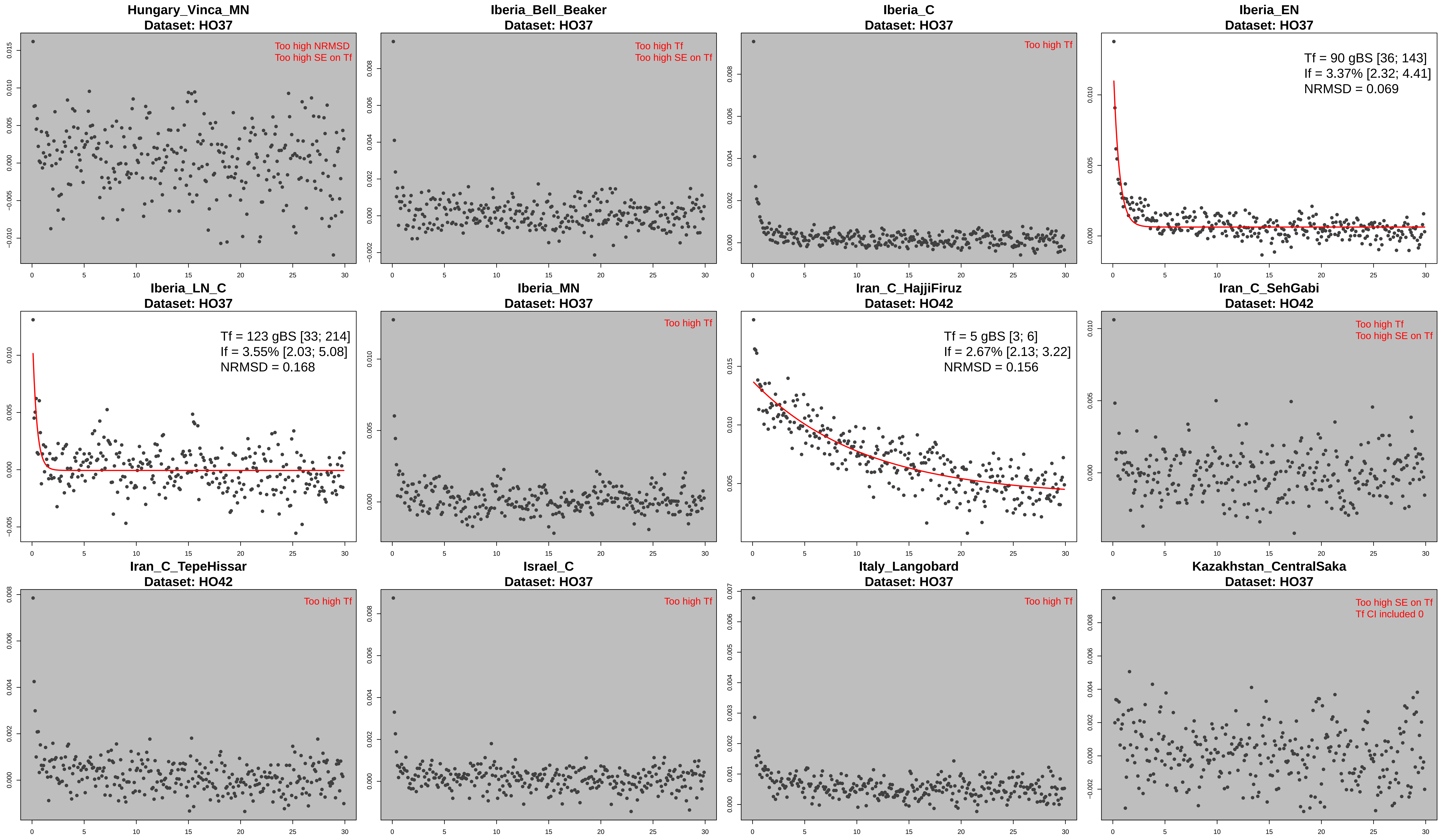
Hungary_Vinca_MN
Dataset: HO37
Too high NRMSD
Too high SE on Tf
Iberia_Bell_Beaker
Dataset: HO37
Too high Tf
Too high SE on Tf
Iberia_C
Dataset: HO37
Too high Tf
Iberia_EN
Dataset: HO37
Tf = 90 gBS [36; 143]
If = 3.37% [2.32; 4.41]
NRMSD = 0.069
Iberia_LN_C
Dataset: HO37
Tf = 123 gBS [33; 214]
If = 3.55% [2.03; 5.08]
NRMSD = 0.168
Iberia_MN
Dataset: HO37
Too high Tf
Iran_C_HajjiFiruz
Dataset: HO42
Tf = 5 gBS [3; 6]
If = 2.67% [2.13; 3.22]
NRMSD = 0.156
Iran_C_SehGabi
Dataset: HO42
Too high Tf
Too high SE on Tf
Iran_C_TepeHissar
Dataset: HO42
Too high Tf
Israel_C
Dataset: HO37
Too high Tf
Italy_Langobard
Dataset: HO37
Too high Tf
Kazakhstan_CentralSaka
Dataset: HO37
Too high SE on Tf
Tf CI included 0

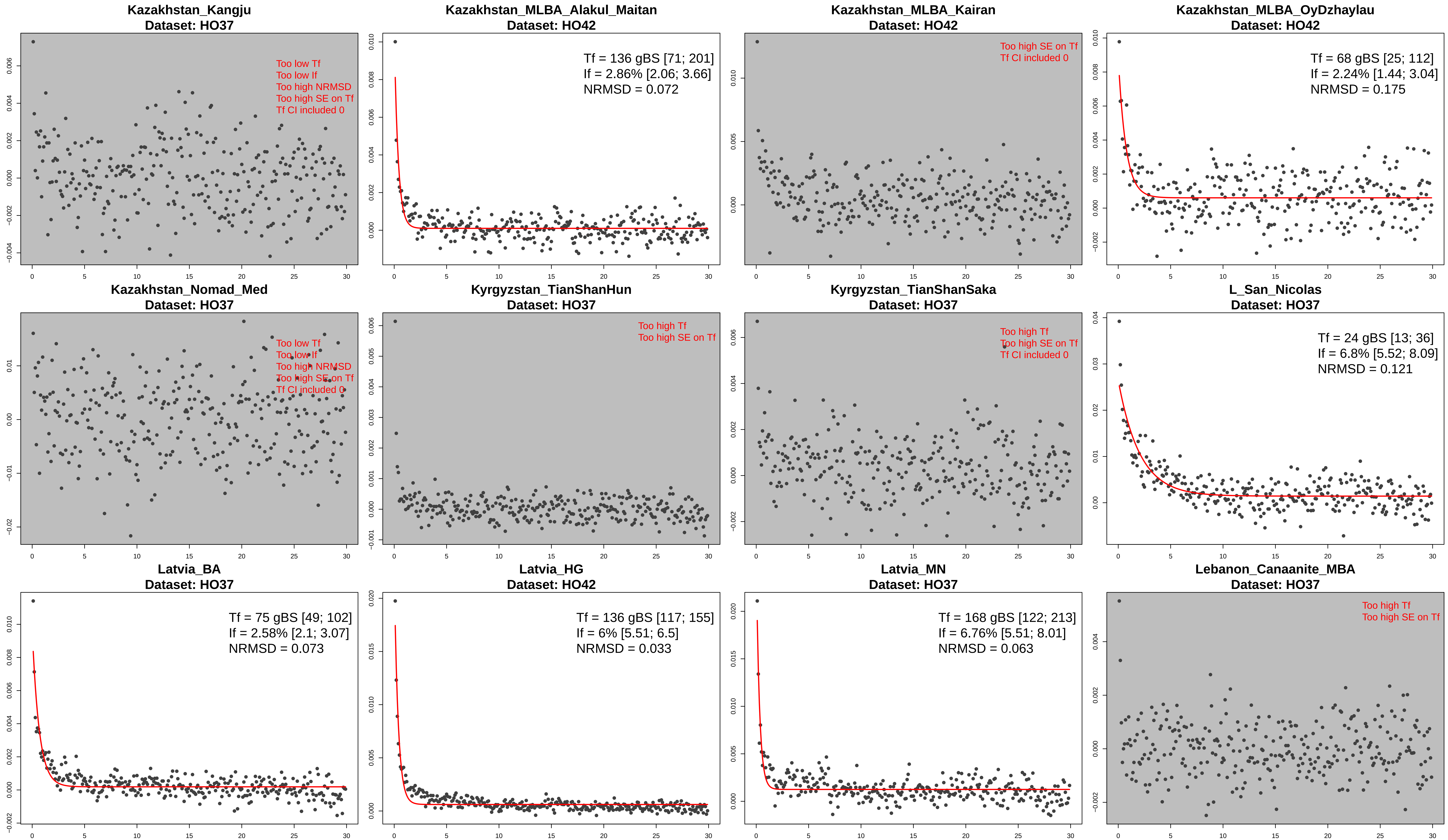

Kazakhstan_Kangju
Dataset: HO37
Too low Tf
Too low If
Too high NRMSD
Too high SE on Tf
Tf CI included 0
Kazakhstan_MLBA_Alakul_Maitan
Dataset: HO42
Tf = 136 gBS [71; 201]
If = 2.86% [2.06; 3.66]
NRMSD = 0.072
Kazakhstan_MLBA_Kairan
Dataset: HO42
Too high SE on Tf
Tf CI included 0
Kazakhstan_MLBA_OyDzhaylau
Dataset: HO42
Tf = 68 gBS [25; 112]
If = 2.24% [1.44; 3.04]
NRMSD = 0.175
Kazakhstan_Nomad_Med
Dataset: HO37
Too low Tf
Too low If
Too high NRMSD
Too high SE on Tf
Tf CI included 0
Kyrgyzstan_TianShanHun
Dataset: HO37
Too high Tf
Too high SE on Tf
Kyrgyzstan_TianShanSaka
Dataset: HO37
Too high Tf
Too high SE on Tf
Tf CI included 0
L_San_Nicolas
Dataset: HO37
Tf = 24 gBS [13; 36]
If = 6.8% [5.52; 8.09]
NRMSD = 0.121
Latvia_BA
Dataset: HO37
Tf = 75 gBS [49; 102]
If = 2.58% [2.1; 3.07]
NRMSD = 0.073
Latvia_HG
Dataset: HO42
Tf = 136 gBS [117; 155]
If = 6% [5.51; 6.5]
NRMSD = 0.033
Latvia_MN
Dataset: HO37
Tf = 168 gBS [122; 213]
If = 6.76% [5.51; 8.01]
NRMSD = 0.063
Lebanon_Canaanite_MBA
Dataset: HO37
Too high Tf
Too high SE on Tf

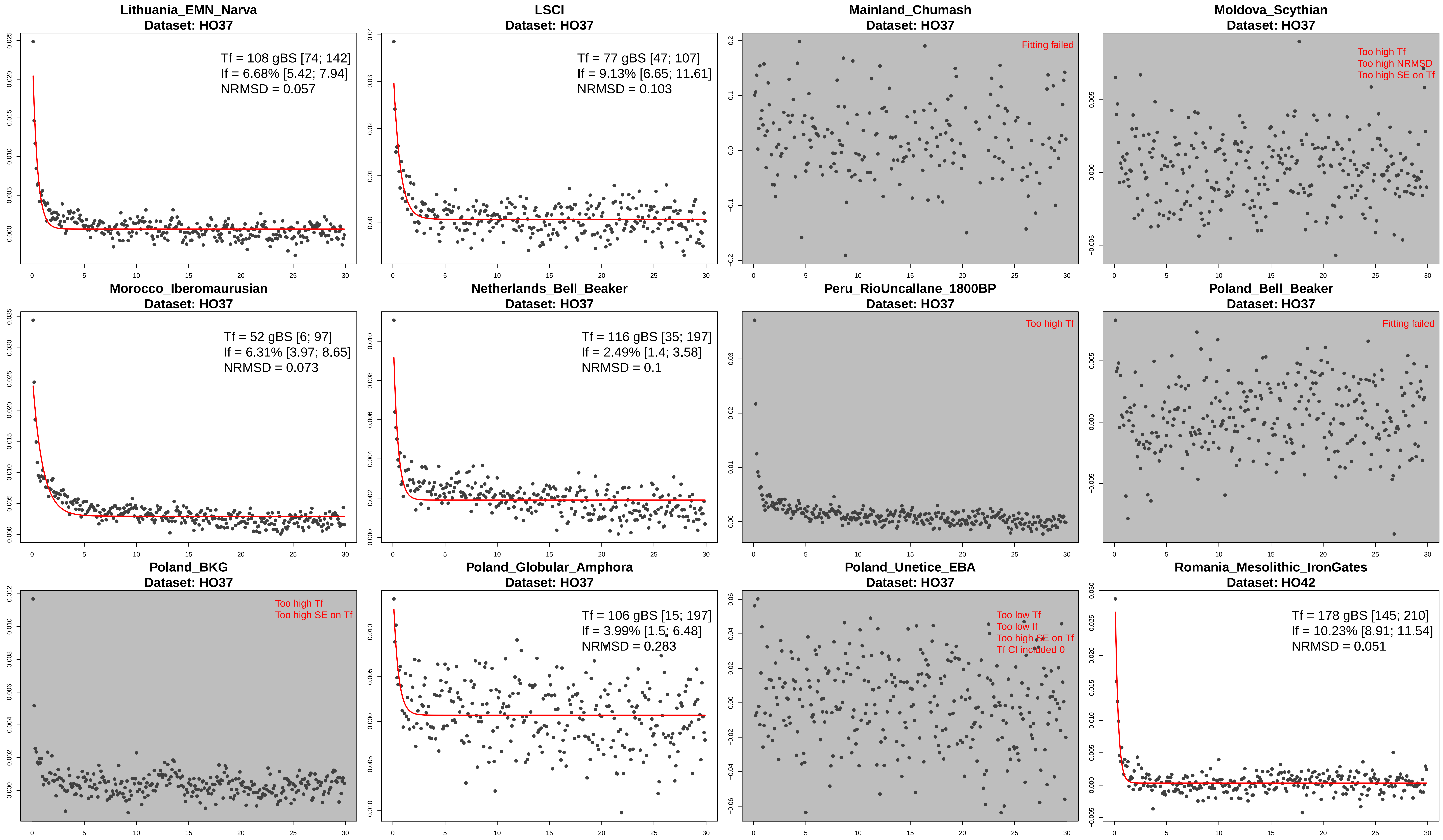

Lithuania_EMN_Narva
Dataset: HO37
Tf = 108 gBS [74; 142]
If = 6.68% [5.42; 7.94]
NRMSD = 0.057
LSCI
Dataset: HO37
Tf = 77 gBS [47; 107]
If = 9.13% [6.65; 11.61]
NRMSD = 0.103
Mainland_Chumash
Dataset: HO37
Fitting failed
Moldova_Scythian
Dataset: HO37
Too high Tf
Too high NRMSD
Too high SE on Tf
Morocco_Iberomaurusian
Dataset: HO37
Tf = 52 gBS [6; 97]
If = 6.31% [3.97; 8.65]
NRMSD = 0.073
Netherlands_Bell_Beaker
Dataset: HO37
Tf = 116 gBS [35; 197]
If = 2.49% [1.4; 3.58]
NRMSD = 0.1
Peru_RioUncallane_1800BP
Dataset: HO37
Too high Tf
Poland_Bell_Beaker
Dataset: HO37
Fitting failed
Poland_BKG
Dataset: HO37
Too high Tf
Too high SE on Tf
Poland_Globular_Amphora
Dataset: HO37
Tf = 106 gBS [15; 197]
If = 3.99% [1.5; 6.48]
NRMSD = 0.283
Poland_Unetice_EBA
Dataset: HO37
Too low Tf
Too low If
Too high SE on Tf
Tf CI included 0
Romania_Mesolithic_IronGates
Dataset: HO42
Tf = 178 gBS [145; 210]
If = 10.23% [8.91; 11.54]
NRMSD = 0.051

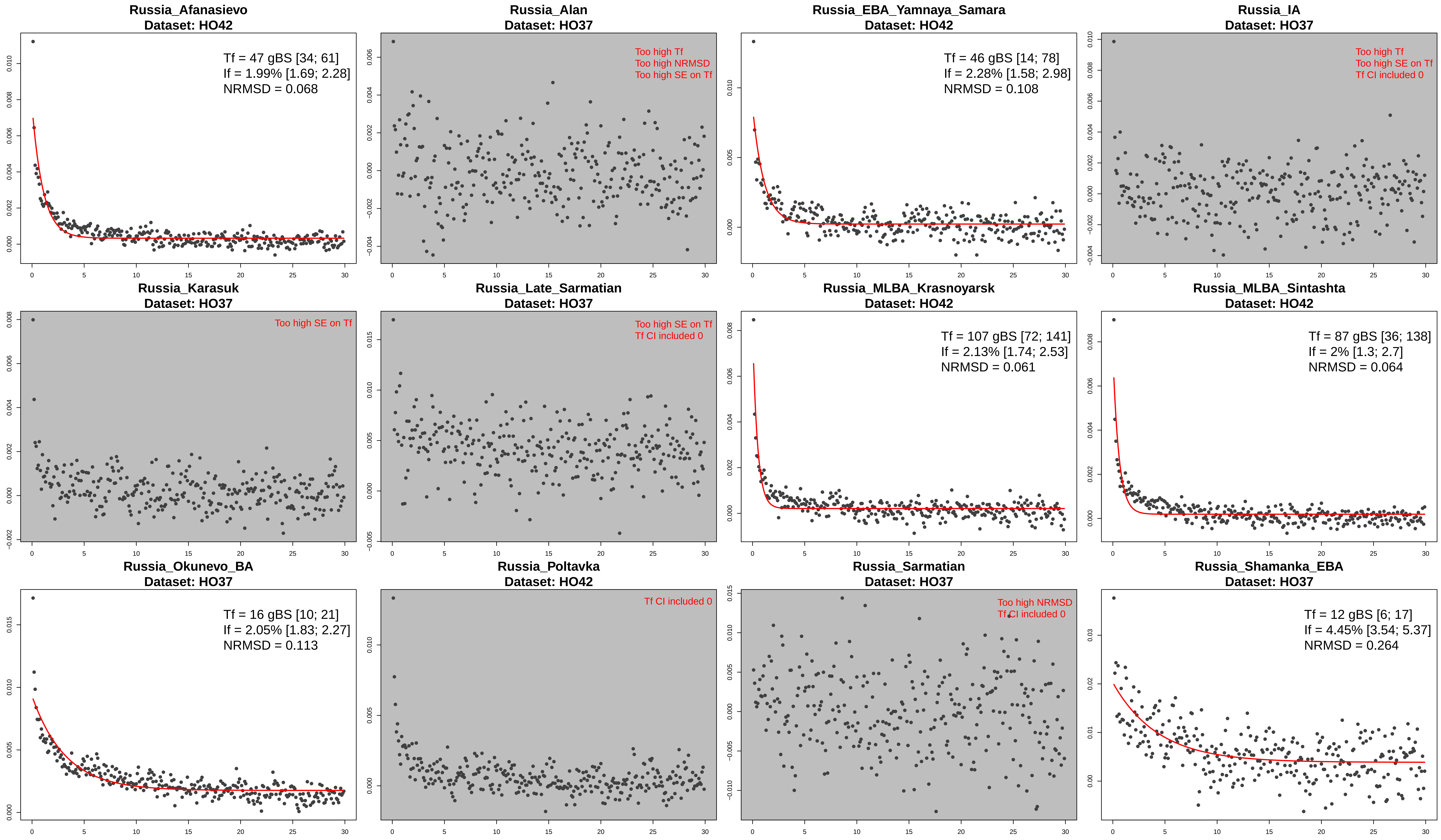
Russia_Afanasievo
Dataset: HO42
Tf = 47 gBS [34; 61]
If = 1.99% [1.69; 2.28]
NRMSD = 0.068
Russia_Alan
Dataset: HO37
Too high Tf
Too high NRMSD
Too high SE on Tf
Russia_EBA_Yamnaya_Samara
Dataset: HO42
Tf = 46 gBS [14; 78]
If = 2.28% [1.58; 2.98]
NRMSD = 0.108
Russia_IA
Dataset: HO37
Too high Tf
Too high SE on Tf
Tf CI included 0
Russia_Karasuk
Dataset: HO37
Too high SE on Tf
Russia_Late_Sarmatian
Dataset: HO37
Too high SE on Tf
Tf CI included 0
Russia_MLBA_Krasnoyarsk
Dataset: HO42
Tf = 107 gBS [72; 141]
If = 2.13% [1.74; 2.53]
NRMSD = 0.061
Russia_MLBA_Sintashta
Dataset: HO42
Tf = 87 gBS [36; 138]
If = 2% [1.3; 2.7]
NRMSD = 0.064
Russia_Okunevo_BA
Dataset: HO37
Tf = 16 gBS [10; 21]
If = 2.05% [1.83; 2.27]
NRMSD = 0.113
Russia_Poltavka
Dataset: HO42
Tf CI included 0
Russia_Sarmatian
Dataset: HO37
Too high NRMSD
Tf CI included 0
Russia_Shamanka_EBA
Dataset: HO37
Tf = 12 gBS [6; 17]
If = 4.45% [3.54; 5.37]
NRMSD = 0.264

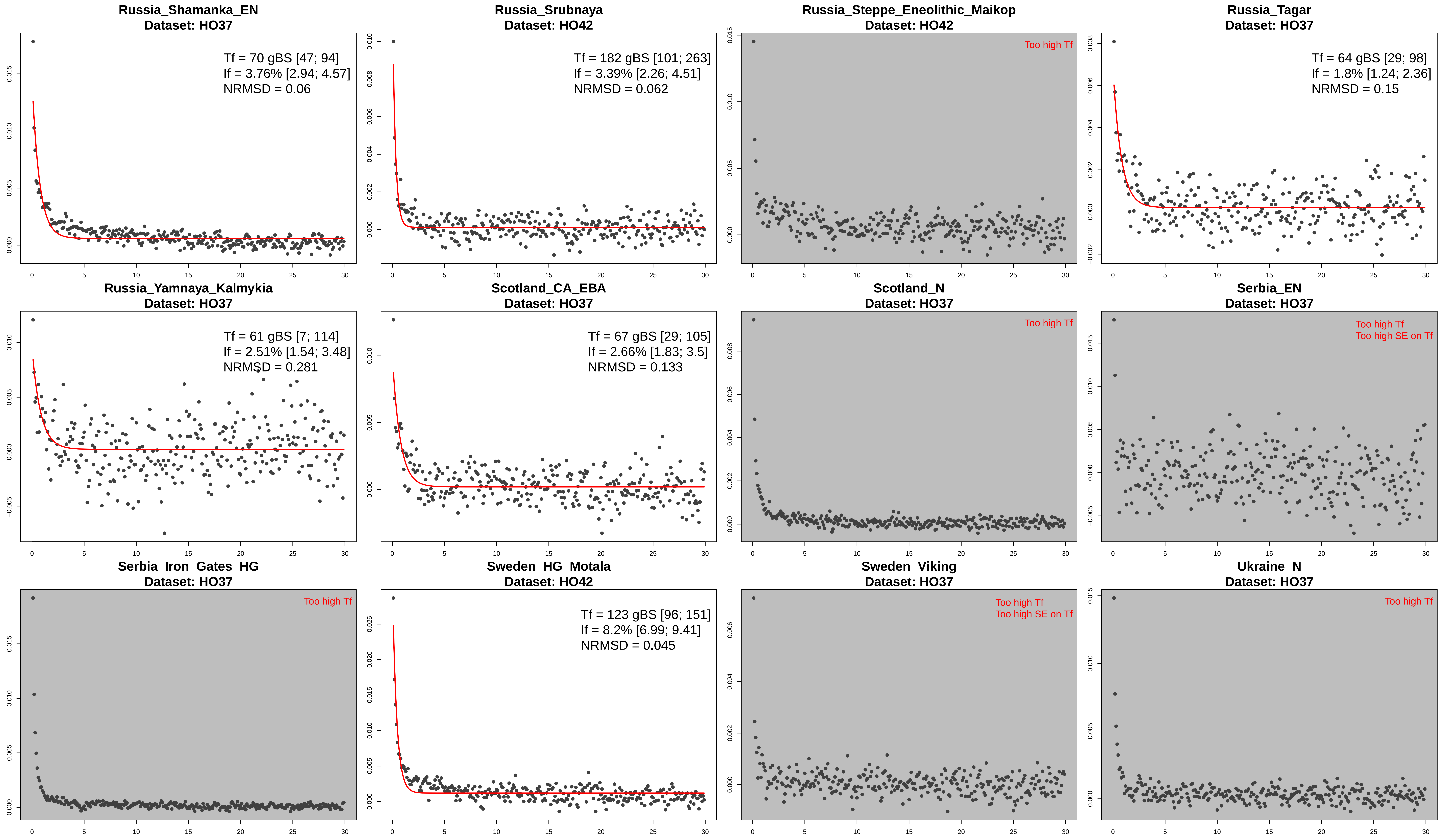
Russia_Shamanka_EN
Dataset: HO37
Tf = 70 gBS [47; 94]
If = 3.76% [2.94; 4.57]
NRMSD = 0.06
Russia_Srubnaya
Dataset: HO42
Tf = 182 gBS [101; 263]
If = 3.39% [2.26; 4.51]
NRMSD = 0.062
Russia_Steppe_Eneolithic_Maikop
Dataset: HO42
Too high Tf
Russia_Tagar
Dataset: HO37
Tf = 64 gBS [29; 98]
If = 1.8% [1.24; 2.36]
NRMSD = 0.15
Russia_Yamnaya_Kalmykia
Dataset: HO37
Tf = 61 gBS [7; 114]
If = 2.51% [1.54; 3.48]
NRMSD = 0.281
Scotland_CA_EBA
Dataset: HO37
Tf = 67 gBS [29; 105]
If = 2.66% [1.83; 3.5]
NRMSD = 0.133
Scotland_N
Dataset: HO37
Too high Tf
Serbia_EN
Dataset: HO37
Too high Tf
Too high SE on Tf
Serbia_Iron_Gates_HG
Dataset: HO37
Too high Tf
Sweden_HG_Motala
Dataset: HO42
Tf = 123 gBS [96; 151]
If = 8.2% [6.99; 9.41]
NRMSD = 0.045
Sweden_Viking
Dataset: HO37
Too high Tf
Too high SE on Tf
Ukraine_N
Dataset: HO37
Too high Tf

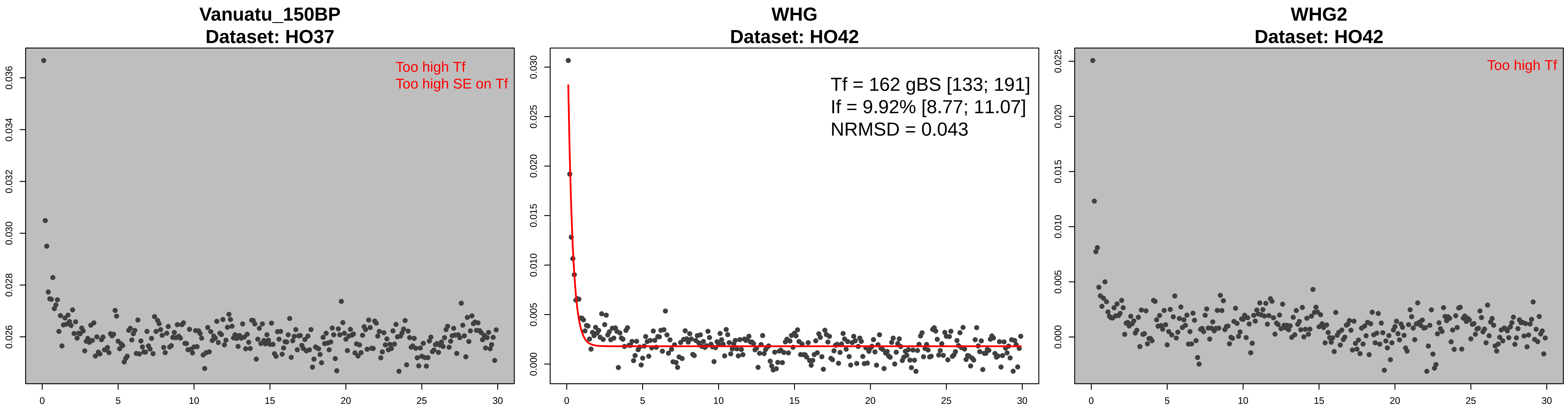
Vanuatu_150BP
Dataset: HO37
Too high Tf
Too high SE on Tf
WHG
Dataset: HO42
Tf = 162 gBS [133; 191]
If = 9.92% [8.77; 11.07]
NRMSD = 0.043
WHG2
Dataset: HO42
Too high Tf

**Figure S5 - History of founder events in ancient human populations from the Human Origins v37 (HO37) and v42 (HO42) datasets.** Panel (A) shows the distribution of founder intensity, and panel (B) shows the distribution of founder ages for all ancient populations that passed our filtering criteria in the HO37 and HO42 datasets and showed evidence for a significant founder event (see Methods). Each point represents a population and the colour of the points indicates the geographical location of the population. *(A) Distribution of the estimated founder intensities.* We show the mean founder intensity (point) and the associated 95% confidence interval. The populations are ordered by their subcontinent and in increasing order of founder intensity. The black horizontal line shows the inferred founder intensity in Ashkenazi Jews (1.7% [1.3%–2.1%]). *(B) Distribution of the estimated founder ages*. We show the estimated founder ages and the associated 95% confidence intervals. The estimated ages were converted from generations to years before present by using a generation time of 28 years[20] and by adding the sampling age of the specimens (shown as the black diamond-shaped points).

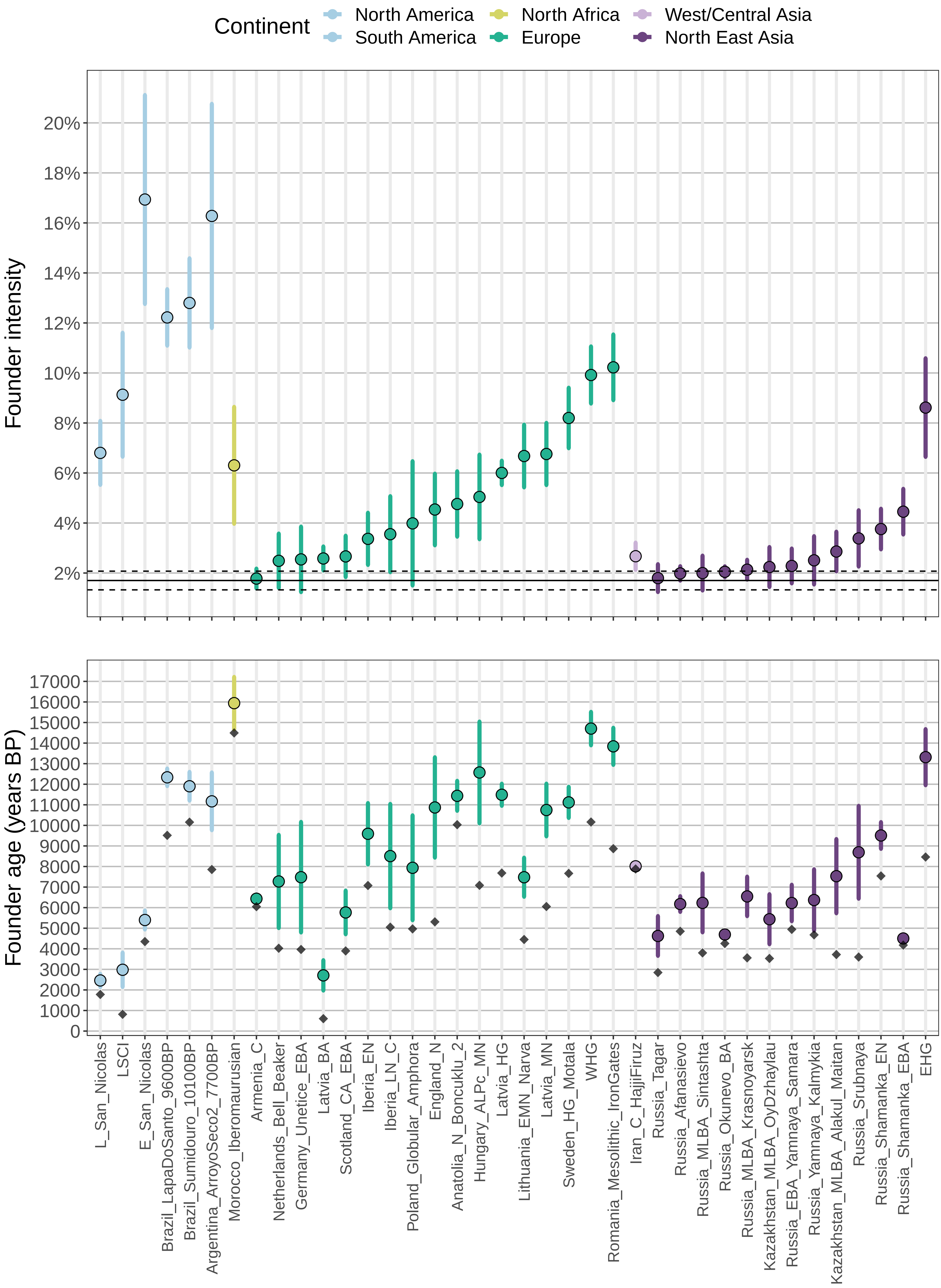
Continent
North America
South America
North Africa
Europe
West/Central Asia
North East Asia
Founder intensity
20%
18%
16%
14%
12%
10%
8%
6%
4%
2%
Founder age (years BP)
17000
16000
15000
14000
13000
12000
11000
10000
9000
8000
7000
6000
5000
4000
3000
2000
1000
0
L_San_Nicolas
LSCI
E_San_Nicolas
Brazil_LapaDoSanto_9600BP
Brazil_Sumidouro_10100BP
Argentina_ArroyoSeco2_7700BP
Morocco_Iberomaurusian
Armenia_C
Netherlands_Bell_Beaker
Germany_Unetice_EBA
Latvia_BA
Scotland_CA_EBA
Iberia_EN
Iberia_LN_C
Poland_Globular_Amphora
England_N
Anatolia_N_Boncuklu_2
Hungary_ALPc_MN
Latvia_HG
Lithuania_EMN_Narva
Latvia_MN
Sweden_HG_Motala
WHG
Romania_Mesolithic_IronGates
Iran_C_HajjiFiruz
Russia_Tagar
Russia_Afanasievo
Russia_MLBA_Sintashta
Russia_Okunevo_BA
Russia_MLBA_Krasnoyarsk
Kazakhstan_MLBA_OyDzhaylau
Russia_EBA_Yamnaya_Samara
Russia_Yamnaya_Kalmykia
Kazakhstan_MLBA_Alakul_Maitan
Russia_Srubnaya
Russia_Shamanka_EN
Russia_Shamanka_EBA
EHG

**Figure S6 - Decay curves for all dog breeds surveyed in the Sams and Hayward datasets.** The x-axis represents the genetic distance (in cM) and the y-axis represents the average allele sharing correlation. The legend shows the mean and 95% confidence interval for the founder age ($T_f$) and the founder intensity ($I_f$), as well as the NRMSD (see Methods). The panels are greyed when the exponential fitting failed or when the evidence for the founder event was not significant (see Methods). The specific reason is highlighted in red in the legend.

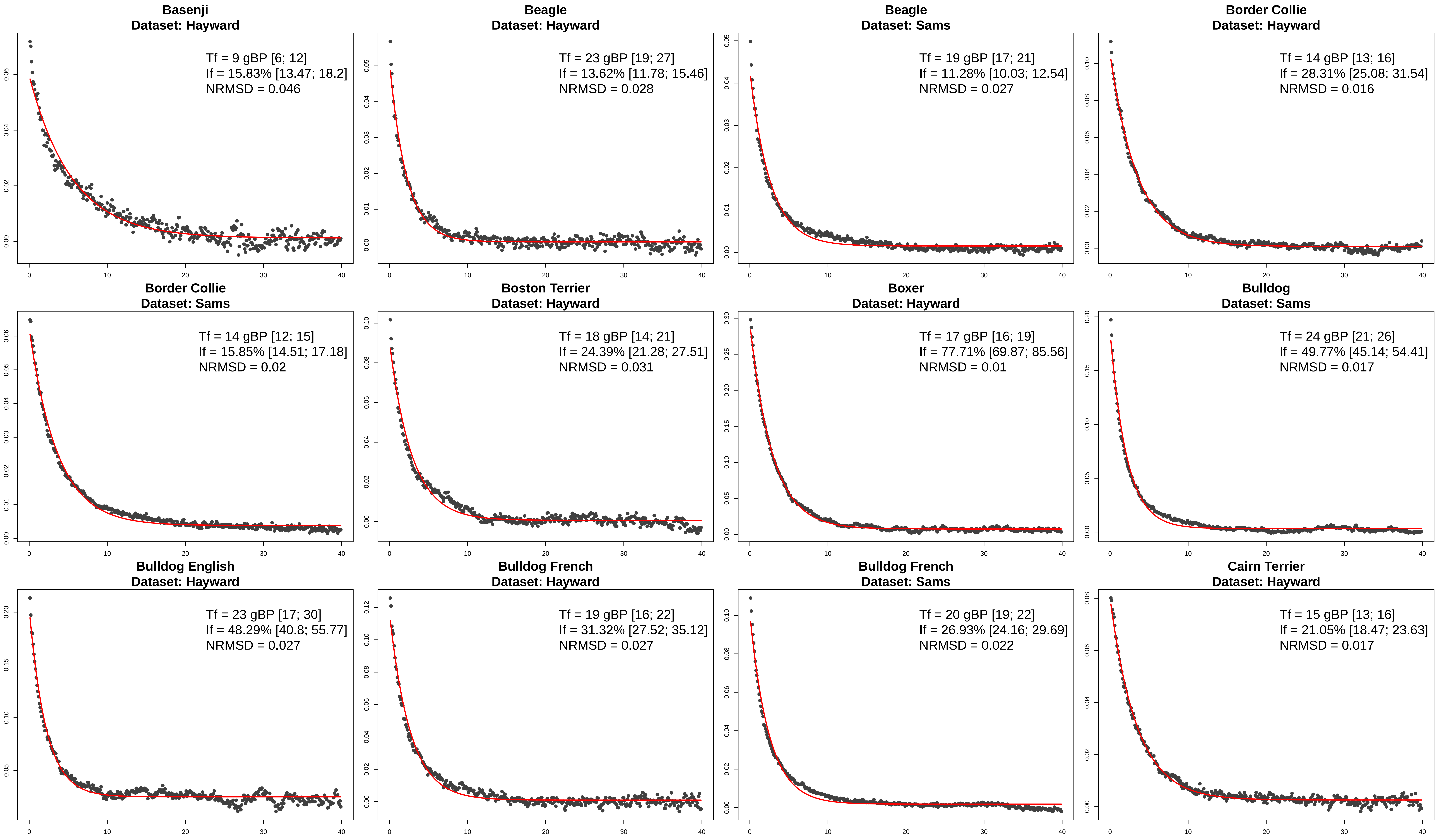
Basenji
Dataset: Hayward
Tf = 9 gBP [6; 12]
If = 15.83% [13.47; 18.2]
NRMSD = 0.046
Beagle
Dataset: Hayward
Tf = 23 gBP [19; 27]
If = 13.62% [11.78; 15.46]
NRMSD = 0.028
Beagle
Dataset: Sams
Tf = 19 gBP [17; 21]
If = 11.28% [10.03; 12.54]
NRMSD = 0.027
Border Collie
Dataset: Hayward
Tf = 14 gBP [13; 16]
If = 28.31% [25.08; 31.54]
NRMSD = 0.016
Border Collie
Dataset: Sams
Tf = 14 gBP [12; 15]
If = 15.85% [14.51; 17.18]
NRMSD = 0.02
Boston Terrier
Dataset: Hayward
Tf = 18 gBP [14; 21]
If = 24.39% [21.28; 27.51]
NRMSD = 0.031
Boxer
Dataset: Hayward
Tf = 17 gBP [16; 19]
If = 77.71% [69.87; 85.56]
NRMSD = 0.01
Bulldog
Dataset: Sams
Tf = 24 gBP [21; 26]
If = 49.77% [45.14; 54.41]
NRMSD = 0.017
Bulldog English
Dataset: Hayward
Tf = 23 gBP [17; 30]
If = 48.29% [40.8; 55.77]
NRMSD = 0.027
Bulldog French
Dataset: Hayward
Tf = 19 gBP [16; 22]
If = 31.32% [27.52; 35.12]
NRMSD = 0.027
Bulldog French
Dataset: Sams
Tf = 20 gBP [19; 22]
If = 26.93% [24.16; 29.69]
NRMSD = 0.022
Cairn Terrier
Dataset: Hayward
Tf = 15 gBP [13; 16]
If = 21.05% [18.47; 23.63]
NRMSD = 0.017

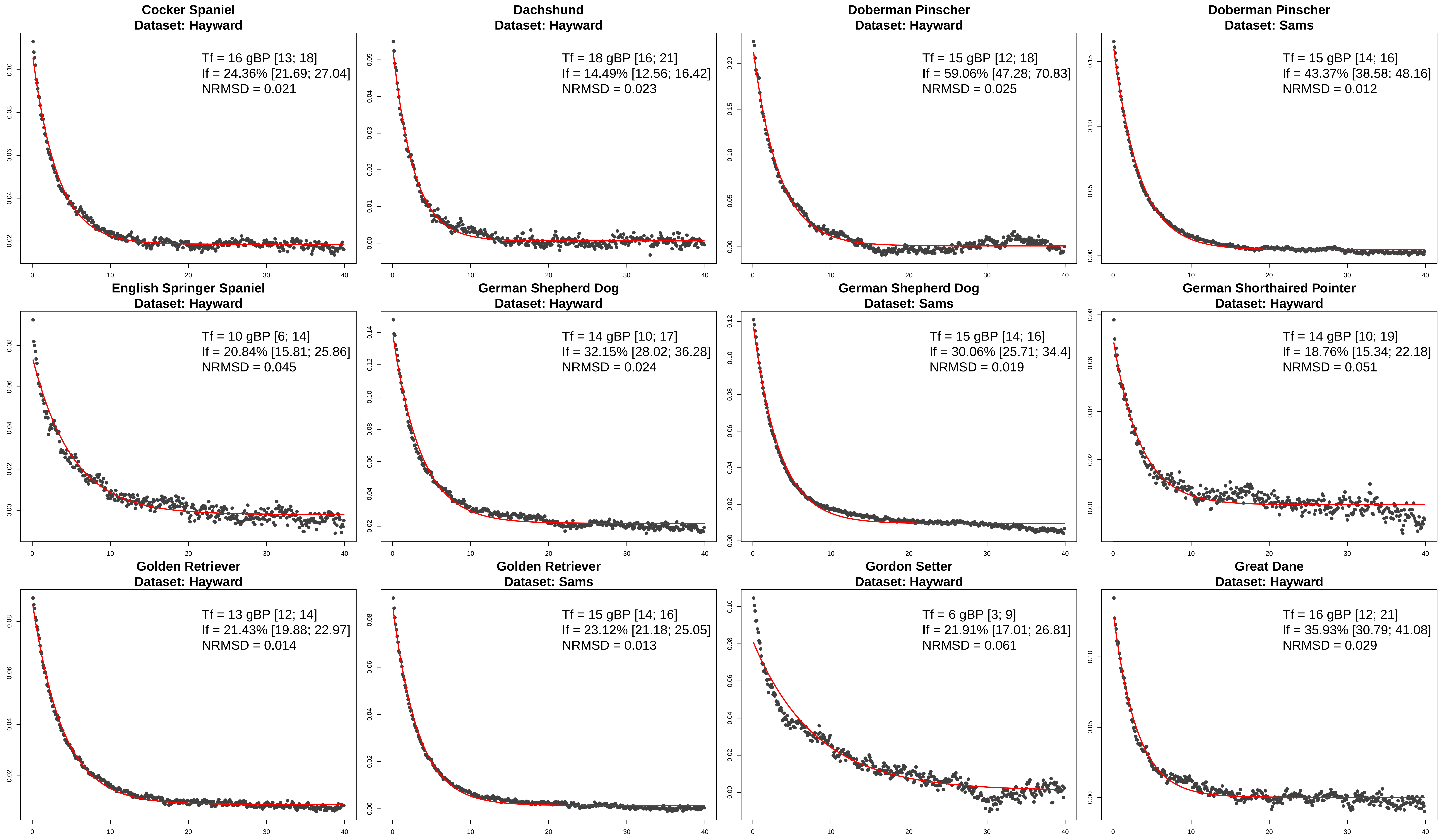
Cocker Spaniel
Dataset: Hayward
Tf = 16 gBP [13; 18]
If = 24.36% [21.69; 27.04]
NRMSD = 0.021
Dachshund
Dataset: Hayward
Tf = 18 gBP [16; 21]
If = 14.49% [12.56; 16.42]
NRMSD = 0.023
Doberman Pinscher
Dataset: Hayward
Tf = 15 gBP [12; 18]
If = 59.06% [47.28; 70.83]
NRMSD = 0.025
Doberman Pinscher
Dataset: Sams
Tf = 15 gBP [14; 16]
If = 43.37% [38.58; 48.16]
NRMSD = 0.012
English Springer Spaniel
Dataset: Hayward
Tf = 10 gBP [6; 14]
If = 20.84% [15.81; 25.86]
NRMSD = 0.045
German Shepherd Dog
Dataset: Hayward
Tf = 14 gBP [10; 17]
If = 32.15% [28.02; 36.28]
NRMSD = 0.024
German Shepherd Dog
Dataset: Sams
Tf = 15 gBP [14; 16]
If = 30.06% [25.71; 34.4]
NRMSD = 0.019
German Shorthaired Pointer
Dataset: Hayward
Tf = 14 gBP [10; 19]
If = 18.76% [15.34; 22.18]
NRMSD = 0.051
Golden Retriever
Dataset: Hayward
Tf = 13 gBP [12; 14]
If = 21.43% [19.88; 22.97]
NRMSD = 0.014
Golden Retriever
Dataset: Sams
Tf = 15 gBP [14; 16]
If = 23.12% [21.18; 25.05]
NRMSD = 0.013
Gordon Setter
Dataset: Hayward
Tf = 6 gBP [3; 9]
If = 21.91% [17.01; 26.81]
NRMSD = 0.061
Great Dane
Dataset: Hayward
Tf = 16 gBP [12; 21]
If = 35.93% [30.79; 41.08]
NRMSD = 0.029

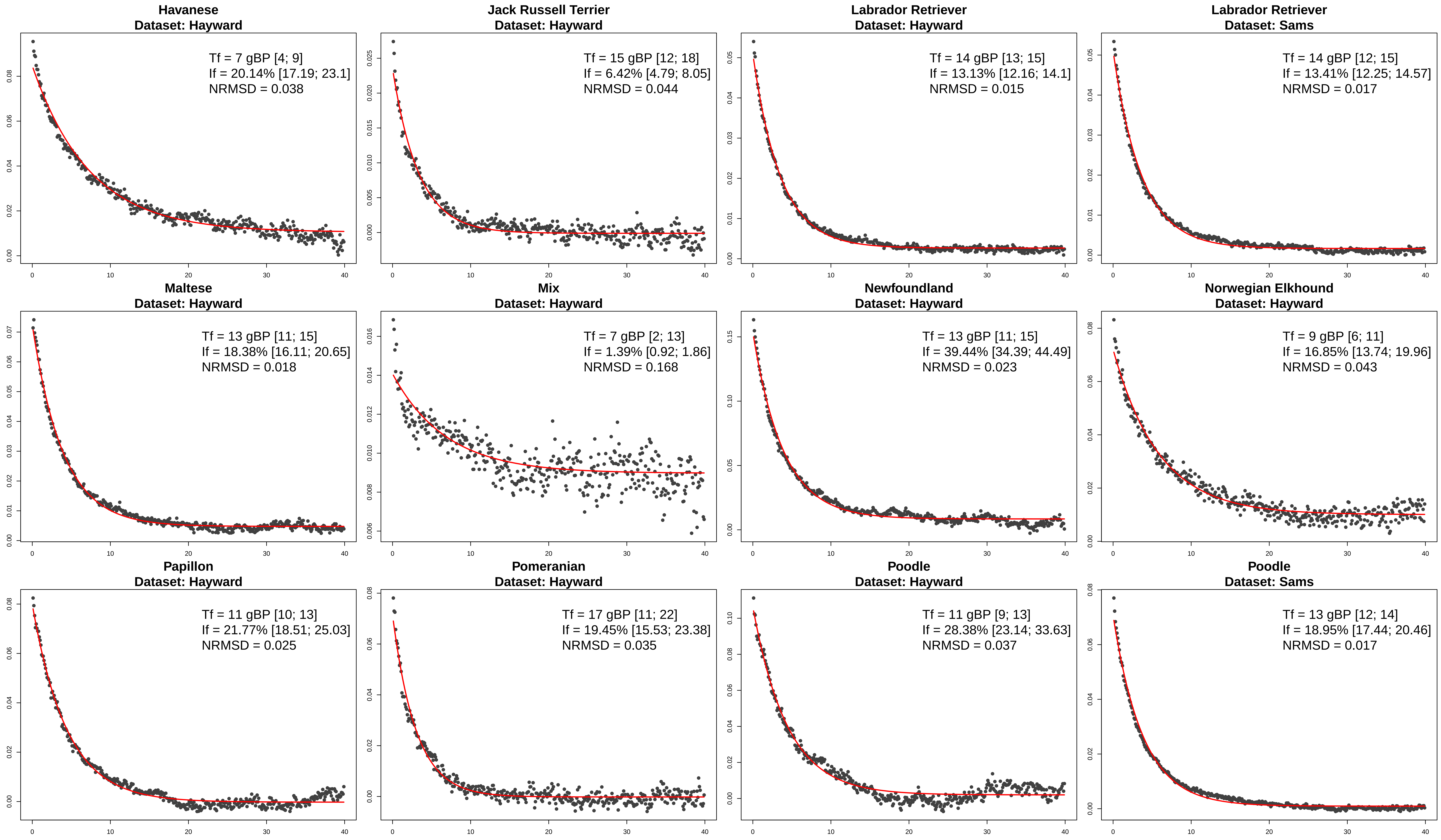
Havanese
Dataset: Hayward
Tf = 7 gBP [4; 9]
If = 20.14% [17.19; 23.1]
NRMSD = 0.038
Jack Russell Terrier
Dataset: Hayward
Tf = 15 gBP [12; 18]
If = 6.42% [4.79; 8.05]
NRMSD = 0.044
Labrador Retriever
Dataset: Hayward
Tf = 14 gBP [13; 15]
If = 13.13% [12.16; 14.1]
NRMSD = 0.015
Labrador Retriever
Dataset: Sams
Tf = 14 gBP [12; 15]
If = 13.41% [12.25; 14.57]
NRMSD = 0.017
Maltese
Dataset: Hayward
Tf = 13 gBP [11; 15]
If = 18.38% [16.11; 20.65]
NRMSD = 0.018
Mix
Dataset: Hayward
Tf = 7 gBP [2; 13]
If = 1.39% [0.92; 1.86]
NRMSD = 0.168
Newfoundland
Dataset: Hayward
Tf = 13 gBP [11; 15]
If = 39.44% [34.39; 44.49]
NRMSD = 0.023
Norwegian Elkhound
Dataset: Hayward
Tf = 9 gBP [6; 11]
If = 16.85% [13.74; 19.96]
NRMSD = 0.043
Papillon
Dataset: Hayward
Tf = 11 gBP [10; 13]
If = 21.77% [18.51; 25.03]
NRMSD = 0.025
Pomeranian
Dataset: Hayward
Tf = 17 gBP [11; 22]
If = 19.45% [15.53; 23.38]
NRMSD = 0.035
Poodle
Dataset: Hayward
Tf = 11 gBP [9; 13]
If = 28.38% [23.14; 33.63]
NRMSD = 0.037
Poodle
Dataset: Sams
Tf = 13 gBP [12; 14]
If = 18.95% [17.44; 20.46]
NRMSD = 0.017

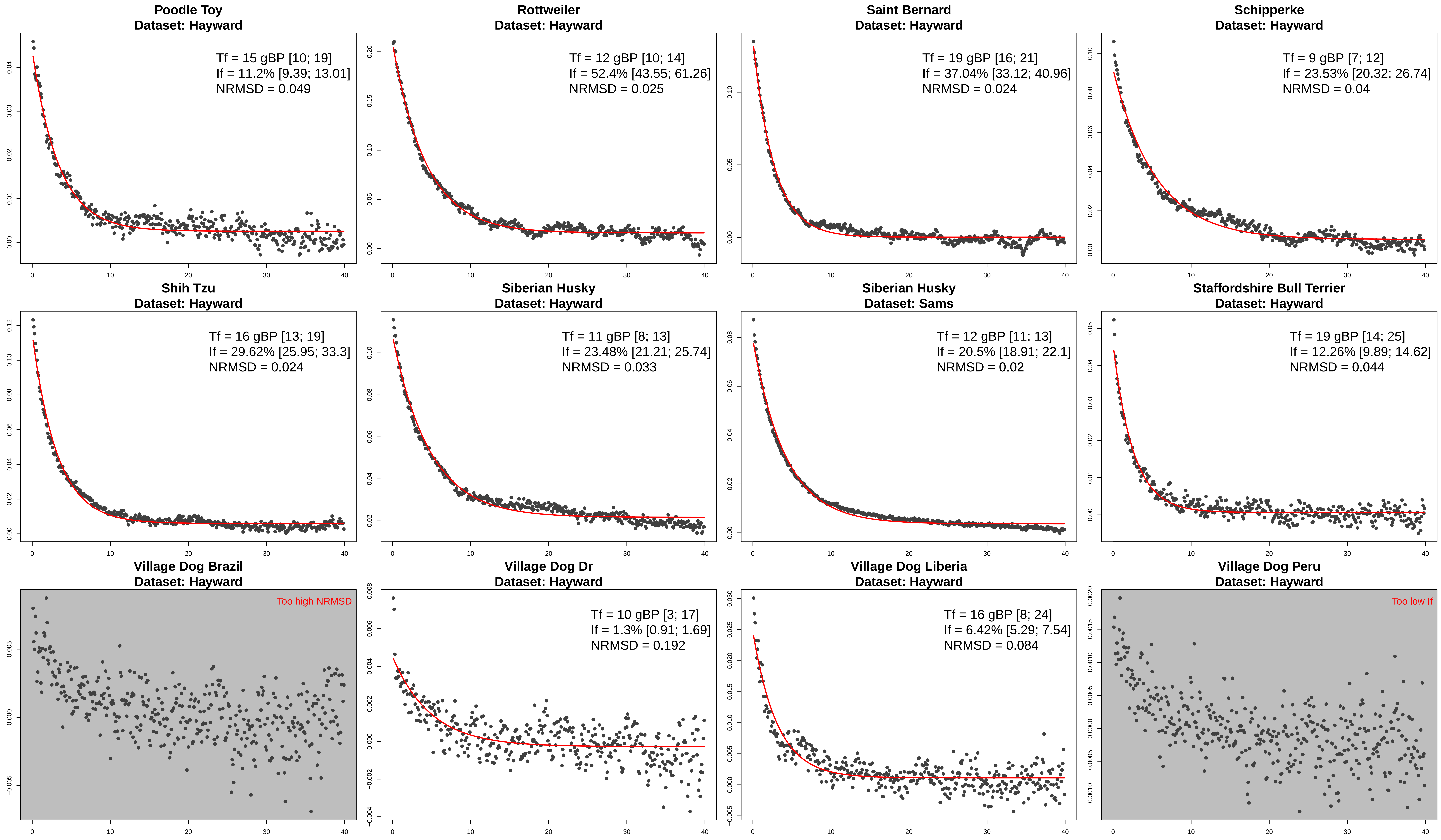

Poodle Toy
Dataset: Hayward
Tf = 15 gBP [10; 19]
If = 11.2% [9.39; 13.01]
NRMSD = 0.049
Rottweiler
Dataset: Hayward
Tf = 12 gBP [10; 14]
If = 52.4% [43.55; 61.26]
NRMSD = 0.025
Saint Bernard
Dataset: Hayward
Tf = 19 gBP [16; 21]
If = 37.04% [33.12; 40.96]
NRMSD = 0.024
Schipperke
Dataset: Hayward
Tf = 9 gBP [7; 12]
If = 23.53% [20.32; 26.74]
NRMSD = 0.04
Shih Tzu
Dataset: Hayward
Tf = 16 gBP [13; 19]
If = 29.62% [25.95; 33.3]
NRMSD = 0.024
Siberian Husky
Dataset: Hayward
Tf = 11 gBP [8; 13]
If = 23.48% [21.21; 25.74]
NRMSD = 0.033
Siberian Husky
Dataset: Sams
Tf = 12 gBP [11; 13]
If = 20.5% [18.91; 22.1]
NRMSD = 0.02
Staffordshire Bull Terrier
Dataset: Hayward
Tf = 19 gBP [14; 25]
If = 12.26% [9.89; 14.62]
NRMSD = 0.044
Village Dog Brazil
Dataset: Hayward
Too high NRMSD
Village Dog Dr
Dataset: Hayward
Tf = 10 gBP [3; 17]
If = 1.3% [0.91; 1.69]
NRMSD = 0.192
Village Dog Liberia
Dataset: Hayward
Tf = 16 gBP [8; 24]
If = 6.42% [5.29; 7.54]
NRMSD = 0.084
Village Dog Peru
Dataset: Hayward
Too low If

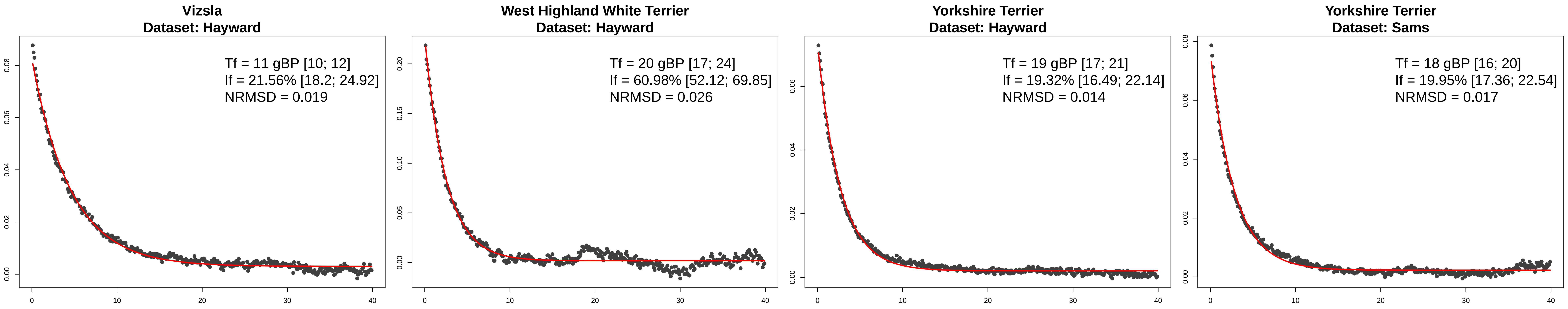
Vizsla
Dataset: Hayward
Tf = 11 gBP [10; 12]
If = 21.56% [18.2; 24.92]
NRMSD = 0.019
West Highland White Terrier
Dataset: Hayward
Tf = 20 gBP [17; 24]
If = 60.98% [52.12; 69.85]
NRMSD = 0.026
Yorkshire Terrier
Dataset: Hayward
Tf = 19 gBP [17; 21]
If = 19.32% [16.49; 22.14]
NRMSD = 0.014
Yorkshire Terrier
Dataset: Sams
Tf = 18 gBP [16; 20]
If = 19.95% [17.36; 22.54]
NRMSD = 0.017
